# Tracking the opening of spike crowns on the surface of coronaviruses

**DOI:** 10.1101/2025.11.02.686173

**Authors:** Di Wu

## Abstract

Circulating over years after the pandemic, SARS-CoV-2 still poses a threat to the human society. The onset of viral infection requires the opening of a trimeric protein, called spike, located on the viral surface essential for binding the host-cell receptors and the subsequent fusion into the host cells. Upon spike-crown opening, one to three Receptor Binding Domains (RBD) rise from the compactly assembled spike head, reaching for the host receptors. Many spike structures were solved that captured RBDs in the different down-to-up trimeric states. These structures not only depict spike’s conformational change but also help design the more efficacious antiviral therapeutics. However, such a dynamic crown-opening pathway is hardly described by only few stationary pictures, and questions yet remain. Do all RBDs rise following the same track in the various studies? If they do, what does this track look like? Is there a common adaptive conformational change of spike as its crown opens? Do all trimeric RBDs rise cooperatively in each spike? And how does it relate to the antibody-binding event? Here, a general RBD-rising pathway, describing the crown-opening dynamics using two angular parameters, was proposed based on analyzing the published spike structures. These analyses describe not only the orientational change of individual RBDs, but also the asymmetric rising preference of the trimeric RBDs in a single-spike entity. In addition, the quantified map clearly describes RBD’s spatial change upon antibody binding, which is often accompanied by an enlarged crown-opening scale. These findings may provide additional clues to develop therapeutics targeting viral spikes in the future.

**Highlights:**

- 778 spike structures were analyzed that revealed a general RBD-rising pathway.
- More intermediate-state RBD structures were identified, one of which might be related to the high FRET-signal state.
- RBD twisted upon rotating up.
- Antibodies elicited larger RBD-rising scales.

**In summary:** *α* and *β* rotation-parameters were introduced that outlined a general RBD-rising pathway, along which the DOWN-, intermediate-, and UP-state RBDs were easily detected and compared that helped explain unresolved issues and provided additional clues in developing therapeutics targeting viral spikes.

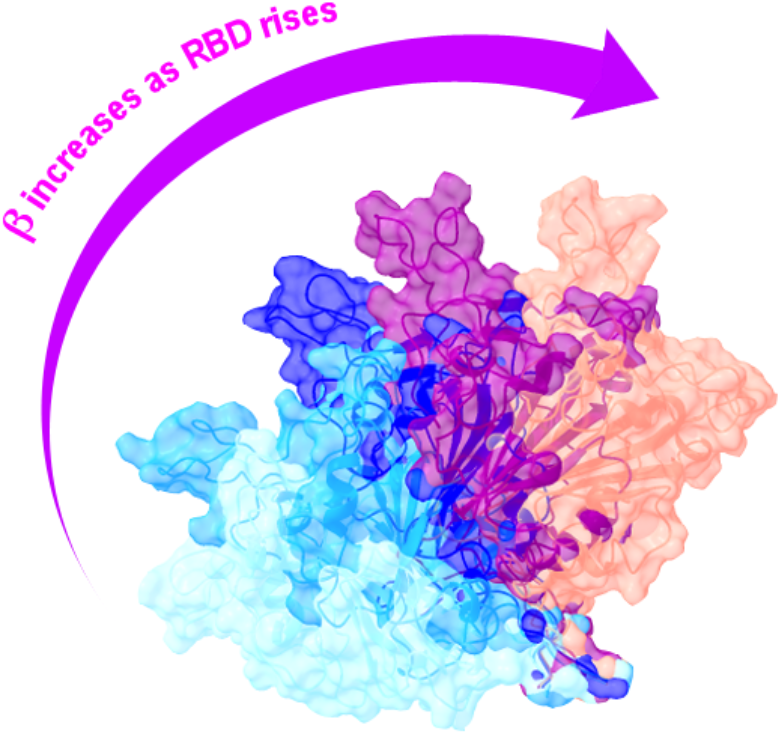

## Introduction

SARS-CoV-2, an enveloped RNA virus belonging to betacoronavirus genus^1^, appeared in 2019^2,3^, quickly spread globally, claimed over 7.1 million lives by October 2025^4^, is still circulating in human society with the unforeseen variant strains emerging each year^5,6^. Some of these variant strains had successfully evaded the then-efficacious neutralizing antibodies^7-12^. This raised serious concerns that the upcoming ones, equipped with more gain-of-function mutations, might one day develop into the more virulent strains. Thus, research on coronaviruses goes on, with the oriented efforts focusing on predicting the potential mutations on viral spikes^13^, and the subsequently altered binding-affinities on neutralizing antibodies^14-20^. In parallel, antiviral therapeutic development continues^21-31^, exploring new sites on viral spikes^32-38^ and employing cocktail therapies^39-43^, aiming to capture the likely evolving mutant strains. Meanwhile, spikes bearing more stable structures were designed in order to improve the efficacy of vaccines^44-50^. Yet all these efforts point to the same viral protein: spike.

Spike is located on viral surface and is assembled by three protomers, each composed of the S1 and S2 subunits. The S1 subunit is further divided into the N-terminal domain (NTD), the receptor-binding domain (RBD), and two subunit domains SD1 and SD2 (also called CTD1 and CTD2)^51-53^. Spikes are highly mobile on viral surfaces. They swing freely around three hinges located near the bottom of S2 subunit^54^, facilitating their reaching to host cells. Meanwhile, on the top, spike’s crown opens occasionally^55,56^, extending its trimeric RBDs for binding to the host-cell receptor angiotensin-converting enzyme 2 (ACE2)^57-60^. Once bound, it helps anchor the associated S2 subunit, which subsequently fuses into the host cell after reshaping and shedding the S1 subunit^51,61-63.^

Binding to ACE2 requires one to three RBDs rise from the well-packed spike head. This produces a pronounced structural change that is easily detected in experiments^52,64-68^, often called an UP-state RBD, compared to a DOWN-state RBD observed in a crown-closed spike. This DOWN to UP transition of RBD is vital for viral infections, and hence was described in detail via many simulation^69-78^ and experimental studies using techniques such as single-molecule fluorescence (Förster) resonance energy transfer (smFRET) imaging^56,79,80^, hydrogen-deuterium exchange coupled to mass spectrometry (HDX-MS)^81,82^, high-speed atomic force microscopy (HS-AFM)^83,84^, etc. In addition, glycans, covering specific regions on spike crowns (22 N-linked glycosylation sites on each spike protomer were predicted^68,85,86^), may also shield and mediate this crown-opening process^69,75,77^. All these studies were valuable in that they described the features of spikes along its crown-opening pathway. And accurately illustrating this pathway can help develop the more efficacious therapeutics targeting viral spikes.

However, questions yet remain. In FRET experiments, at least four states were identified during the RBD-rising event^56^. Besides the typical DOWN and UP states, an intermediate state and an unrecognized high FRET-signal (0.8) state were detected as well^56^. Simulation studies had successfully described the intermediate states and suggested the high FRET-signal state spike structures^70^. These structures were valuable because they were seldom sampled in experiments. The high FRET-signal state, as usual, was assumed to link with a more compactly assembled crown-closed spike, which was in accordance with common sense. But can there be other possibilities?

Currently, many spike structures were solved, and they may help clarify the issue. Sometimes, the overlooked structural features or some hidden information may be unveiled or extracted by comparing similar structures. In this article, various spike structures were analyzed and compared by RBD-oriented parameters that outlined a general RBD-rising track. On this track, one can detect not only the canonical DOWN- and UP-RBD states, but also the pre-rising, the intermediate, the over-rising, and the outlier RBD states. Therefore, more rarely sampled RBD structures were identified on this pathway that may help explain the unresolved issues. The conclusions made by analyzing these results show additional features of RBDs, which may help design therapeutics targeting viral spikes in the future.

## Results

### Four domains’ adaptive motions

778 spike structures were selected (see Methods). They were brought to the same frame by aligning (see Methods) to a common reference structure (PDB: 8DLW, Figs. 1A and 1B). The rising of RBD (Fig. 1C) can induce sequential adaptive motions in NTD, SD1, and SD2 domains on the same S1 subunit, but affect little to the S2 subunit beneath (although some S2 residues may be involved^82^). And hence S2 subunit (Fig. S1) was omitted from the analyses here. Figures 1D-1G show the center-of-mass (COM) motions of the four domains (see Methods) in Cartesian coordinates. The DOWN and UP (both denoting RBD states) centers were clearly separated for RBD (Fig. 1D), but overlapped for NTD (Fig. 1E) and SD1 (Fig. 1F), and obscure for SD2 (Fig. 1G). Thus, the rising of RBD affects SD1 most, which, contrary to our intuition, shifts downward in the z-direction while RBD moves up.

**Figure 1.**
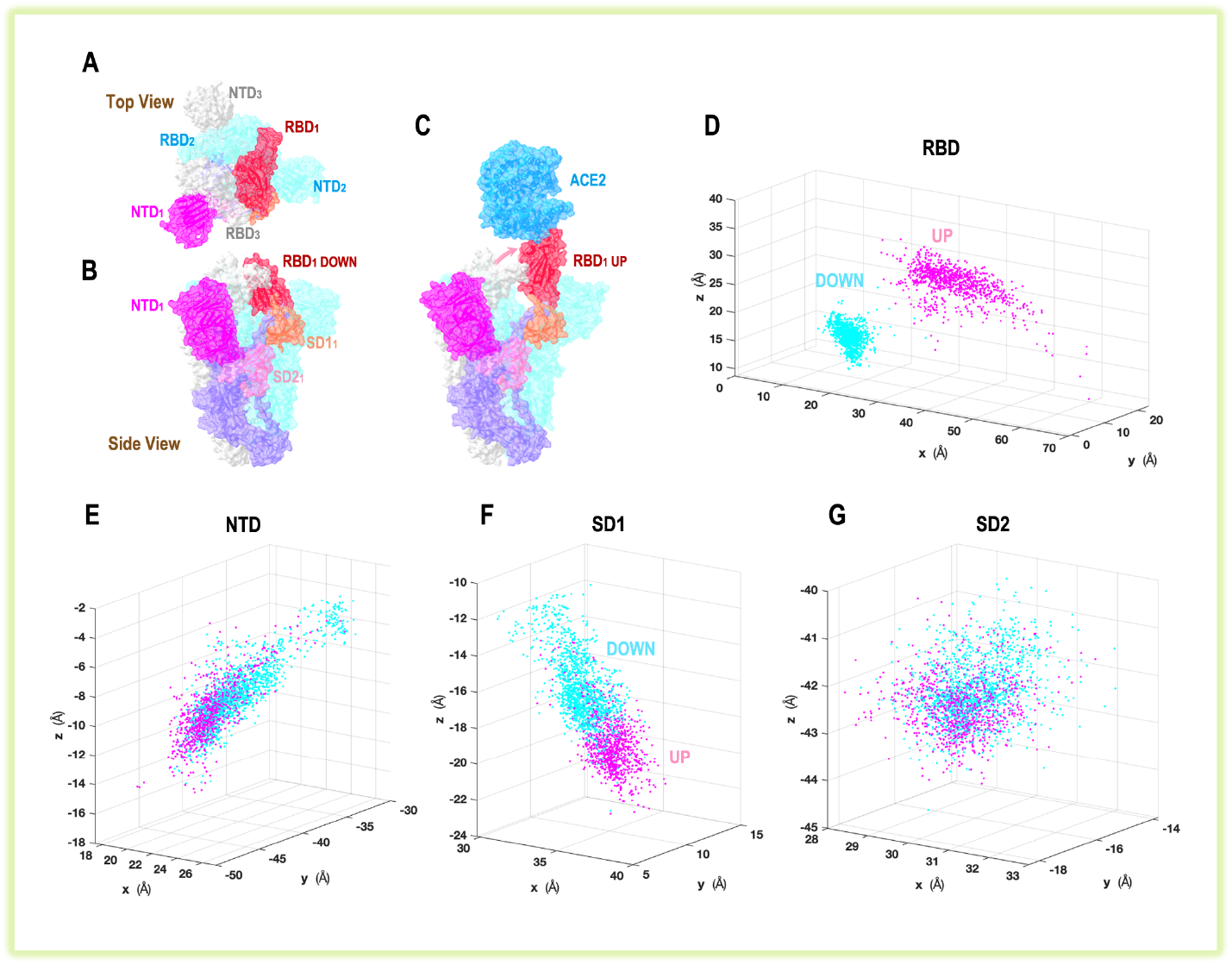
Spike structure and motions of S1 subunit upon crown opening. (A) Top view of a closed crown with trimeric RBDs all in DOWN states packed in the center, surrounded by NTDs (PDB: 8DLW). (B) Side view of the same spike assembled by three protomers colored purple, cyan, and gray. The four domains (used for analyses) on one S1 subunit were colored differently: magenta for NTD, red for RBD, orange for SD1, and pink for SD2. (C) An UP-state RBD binding to a host receptor ACE2 (PDB: 7W98). (D-G) The center-of-mass movements of four domains on each protomer, grouped into the DOWN (cyan dots) or UP (magenta dots) clusters based on the RBD-state on that protomer.

Figure 2 shows NTD, SD1, and SD2 motions relative to the RBD motion on the same protomer, using each domain’s characteristic distance *d* (see Methods). Clearly, SD1 showed the most perceptible movement, where the DOWN and UP cluster-centers were apparently shifted in the *d*_SD1_ direction (y-axis in Fig. 2B). However, such a shift was less obvious in NTD motions (Fig. 2C), albeit *d*_NTD_ showed large fluctuations due to the great flexibility of NTD’s location at the spike-head border. Because RBD is close to the NTD on next protomer (e.g., RBD_1_ is close to NTD_2_, see Fig. 1A), the next-protomer NTD motions were examined as well (Fig. S2), but no apparent difference was found. In contrast, SD2 motions were greatly restricted, showing only tiny fluctuations (Fig. 2D).

**Figure 2.**
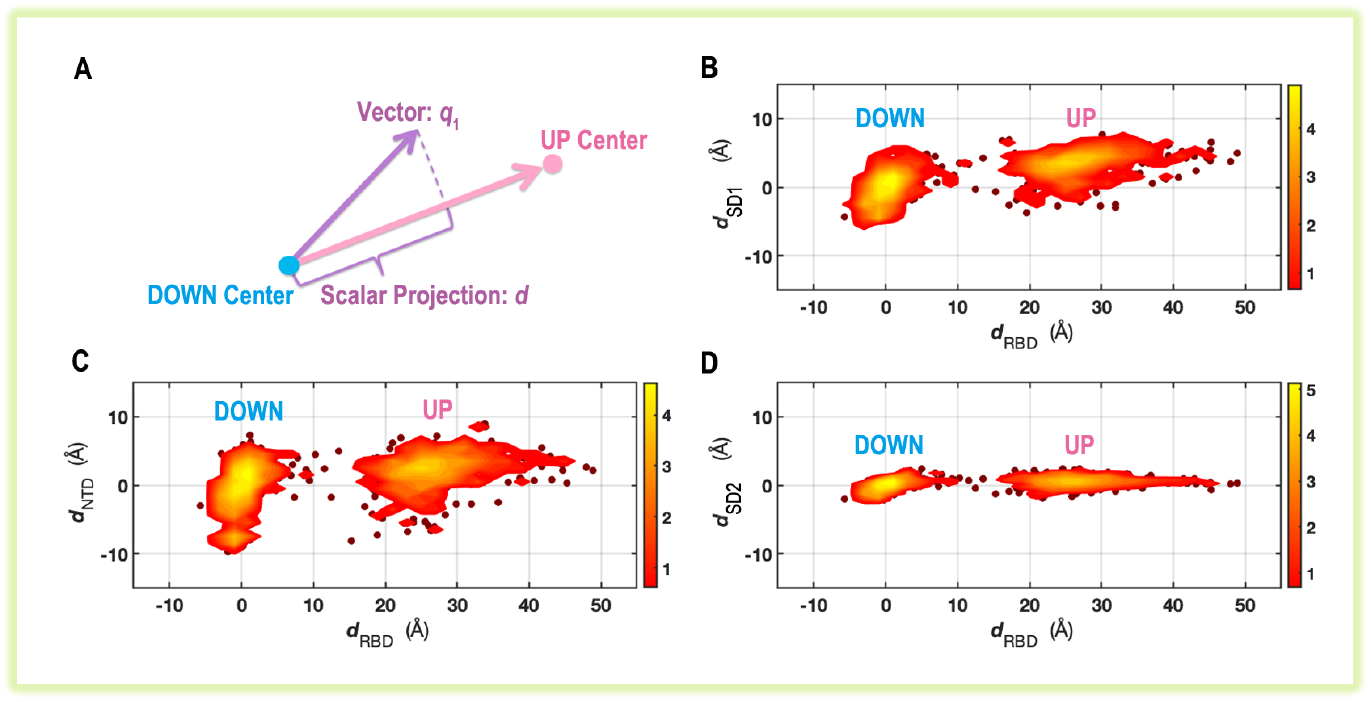
Motions of four domains in the RBD-rising event. (A) The characteristic distance *d* was defined as the scalar projection of a domain’s COM-derived vector *q*_1_ onto a unit vector directed from the DOWN to UP center calculated for that domain (see Methods). Changes of *d* in SD1, NTD, and SD2, with respect to *d*_RBD_ on the same protomer were plotted in (B-D). The color bar on the right-hand side shows the populations labeled in the natural logarithmic scale. See Methods for drawing the contour plots.

Now, drawing these domain motions relative to a crown-closed spike structure (PDB: 8DLW), we see a twisted out-spiraled trajectory of RBD (Fig. 3), accompanied by the adaptive small-scale motions in the associated domains (Fig. S3). Here, parameters describing domain rotations (around spike’s central axis) and elongations were employed (Fig. 3A). They described a particular RBD-rising trajectory, i.e., RBD stretched out not only axially (∼14 Å along z-axis, Δ*z*) but also horizontally (∼14 Å in x-y plane, Δ*d*_xy_), and rotated about 28º (Δ*θ*) around spike’s central axis clockwise when viewed above. In comparison, the movements of SD1 and NTD were greatly reduced, and those of SD2 were very small (Fig. S3). Among them, detectable changes in Δ*z* were observed in SD1 (Fig. S3A), demonstrating its downward movement while RBD moved up.

**Figure 3.**
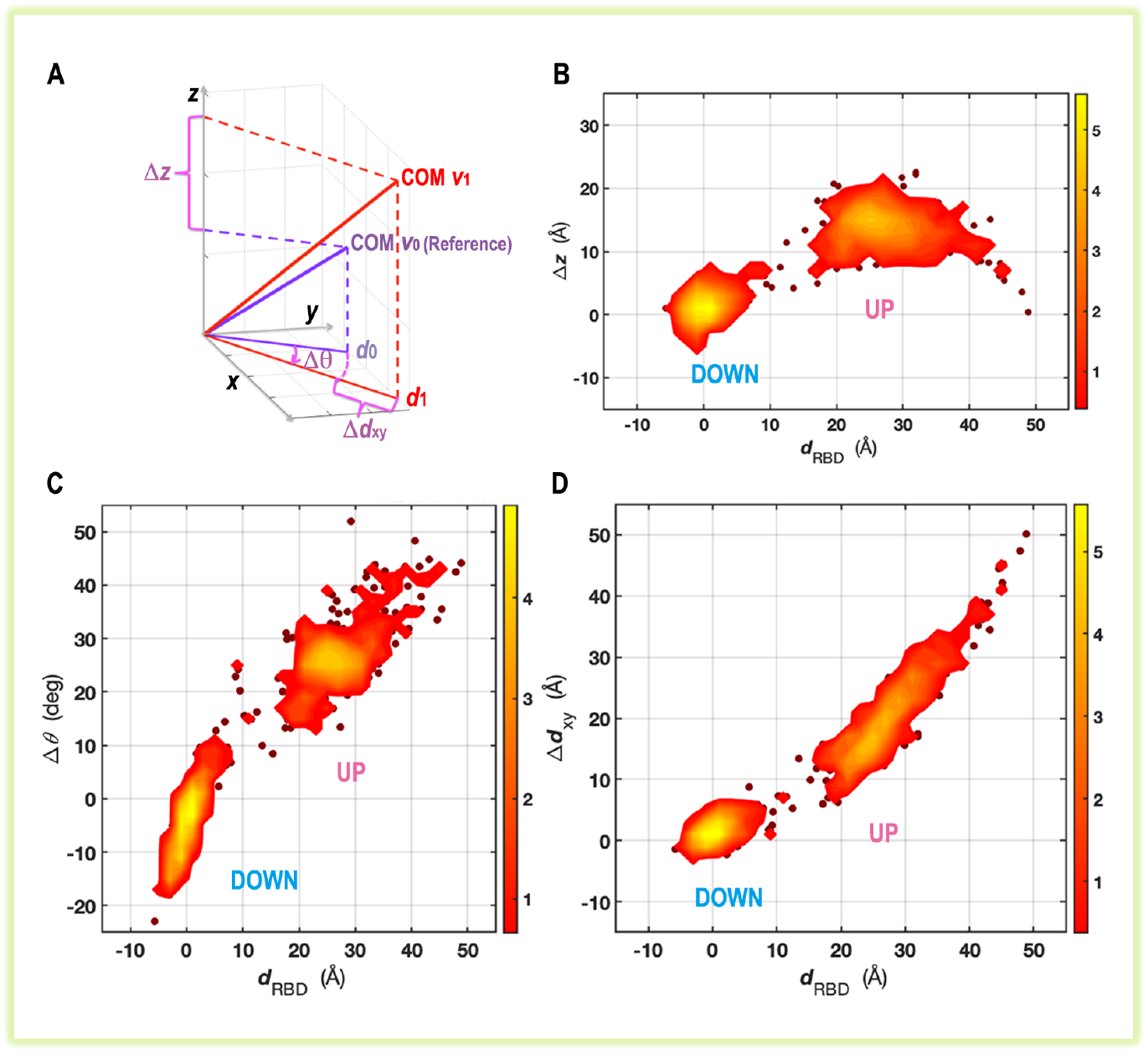
RBD motions relative to the reference spike. (A) Spatial parameters Δ*z*, Δ*θ*, and Δ*d*_xy_ were defined based on vector *v*_1_, relative to vector *v*_0_ of the same domain in the reference structure (see Methods). (B-D) Changes of the parameters Δ*z*, Δ*θ*, and Δ*d*_xy_ of RBD domain were plotted against *d*_RBD_.

### Two angular parameters designed to describe RBD motions

The rising of RBD not only elevates and rotates its COM position relative to a crown-closed spike, but also rotates RBD itself that was not shown in Figs.1-3. This well-recognized hinge-like rotation of RBD can produce COM elevation, but cannot explain COM rotation. To illustrate these rotations, two orientational parameters, *α* and *β*, were introduced (Figs. 4A and 4B, see Methods). Figure 4C shows that *β* linearly correlates with *d*_RBD_, a proper parameter used to describe COM motions (Fig. 2). And hence *β*, in addition to describing RBD rotation, can also substitute for *d*_RBD_ in many cases to describe the crown-opening trajectory.

**Figure 4.**
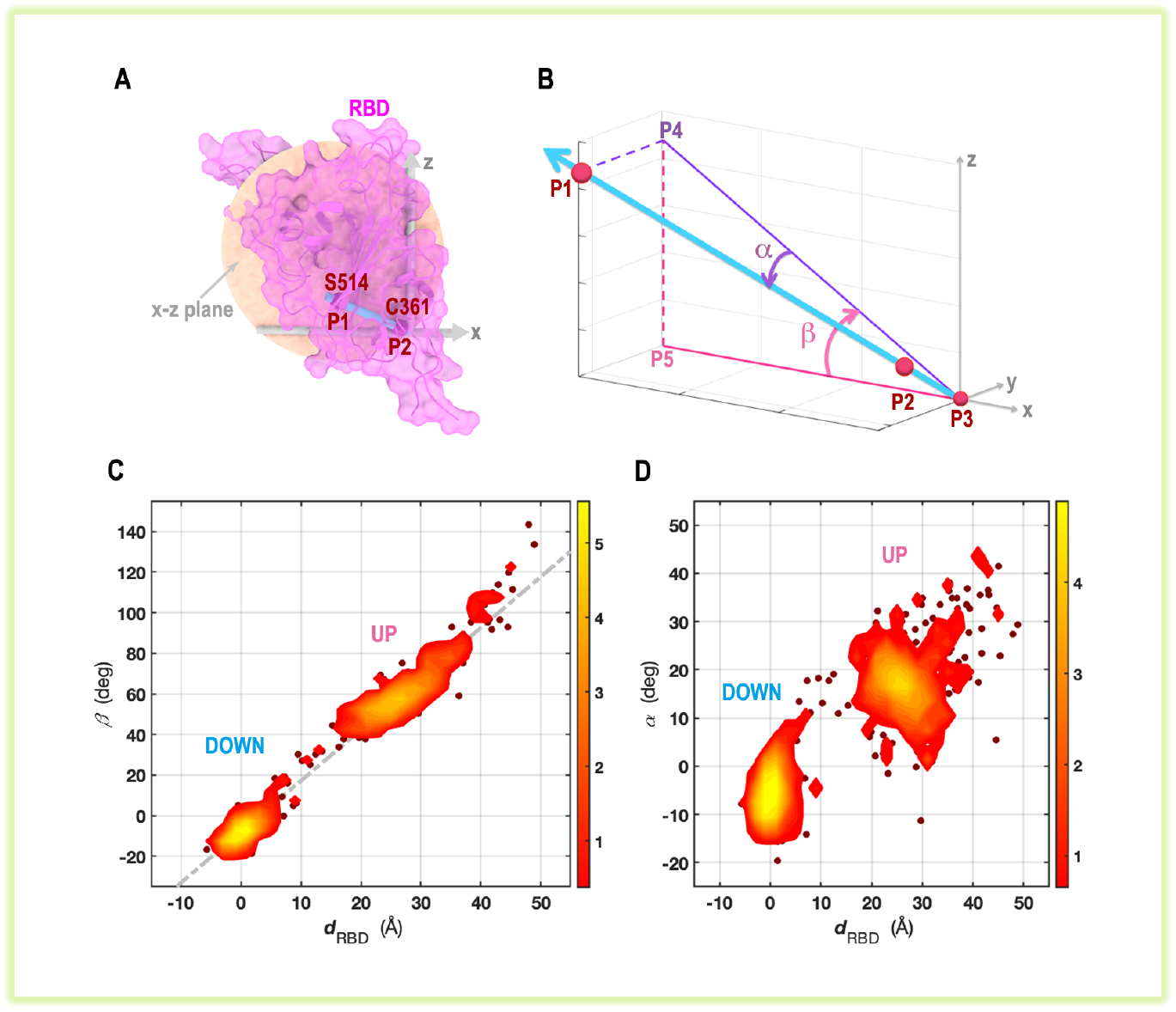
RBD rotations described by two angular parameters. (A). P1 (C*α* of S514) and P2 (C*α* of C361) were selected and drawn as red balls in an RBD structure (PDB: 8D5A). (B) P1 and P2 were used to define angles *α* and *β* (see Methods for the definitions of P3, P4, and P5). (C) Angle *β* correlates linearly with *d*_RBD_. (D) Angle *α* correlates linearly with *d*_RBD_ approximately.

Parameter *α*, on the other hand, shows the twist angle as RBD rises (Fig. 4B). Like parameter *β, α* also correlates with *d*_RBD_, but bearing larger fluctuations (Fig. 4D). Nevertheless, this shows the trend of RBD rotation, that it tends to swing its head away from the original orientation upon rising (a twist motion), as demonstrated by an increase in *α*.

The definitions of *α* and *β* involve a dummy atom P3, whose position does not affect either *α* or *β* values. Inclusion of P3 is only for an illustrative purpose that RBD seems to rotate around a “same” hinge point P3 upon rising. But this description is only approximate since P3 position changed as RBD rose, albeit in small scales (see Fig. S4).

### The RBD-rising pathway

In the different scenarios examined (Figs. 5A-5D), the RBD-rising pathways were regular and consistent, with two well-defined DOWN and UP centers connected by a narrowed neck (red dots) that showed the obligatory transition path via *β* ≈ 16º ∼ 35º and *α* ≈ 10º ∼ 22º. This is exactly the intermediate RBD-state that many studies tried to describe. And excitingly enough, there were experimental structures found in this region, represented by the red dots connecting the DOWN and UP clusters in Figs. 5A-5C. Some of these intermediate states were immediately recognized along with the published spike structures^87-89^, whereas others were not. In this regard, mapping this pathway is useful in that it helps identify more intermediate-state RBDs and label them by distinguishable *α* and *β* values easier for further comparison and analysis. One of these spike structures, characterized by one intermediate-state RBD (*β* = 30.4º, *α* = 19º) and two DOWN-state RBDs, was drawn explicitly and compared with RBDs in a closed crown in Figs. 5E and 5F.

**Figure 5.**
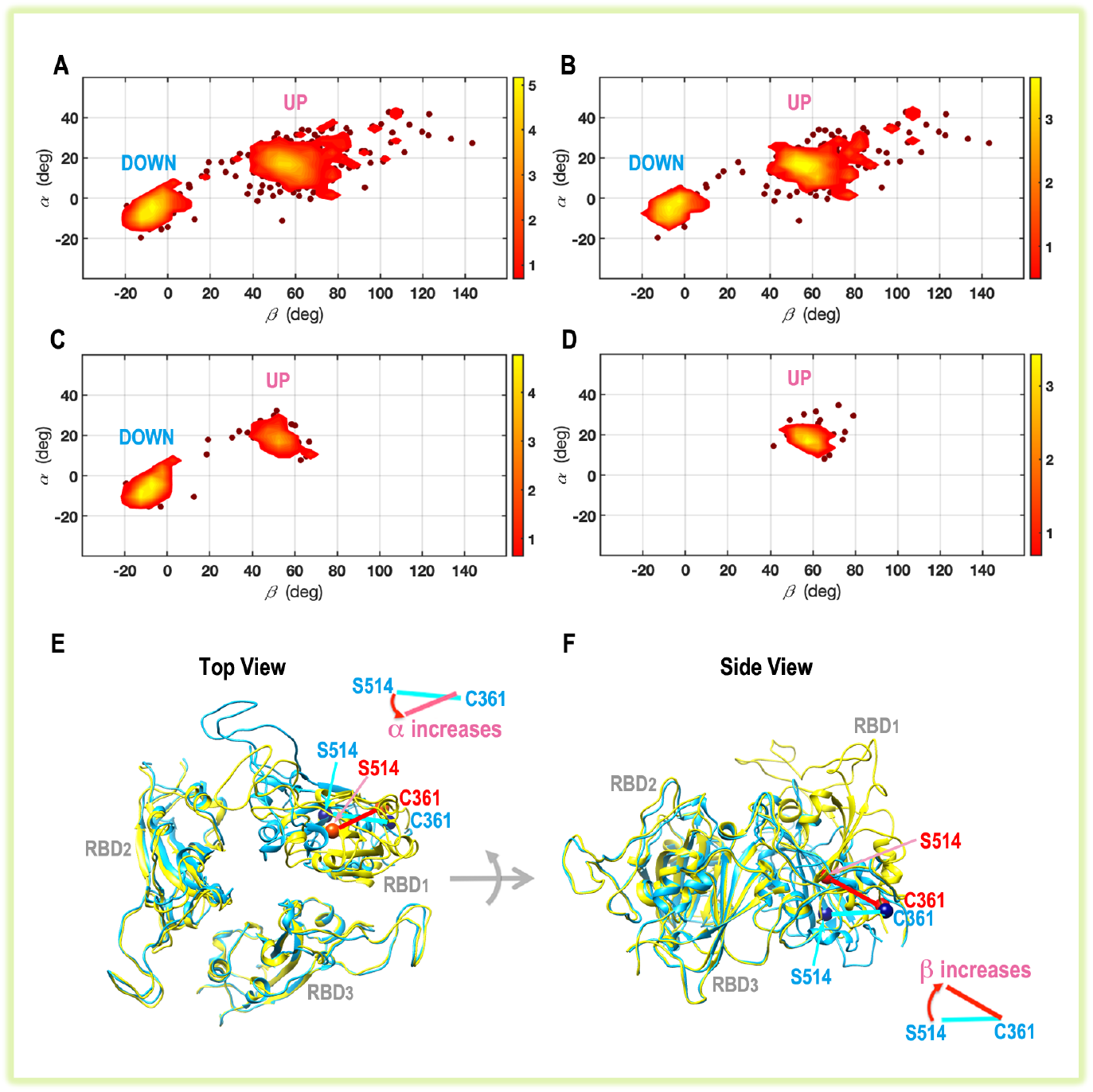
A general RBD-rising pathway outlined by angles *α* and *β*. Four circumstances were examined, where the pathways were calculated using all RBDs selected in this study (A), RBDs bound with antibodies (B), RBDs in the unbound-spikes (C), and RBDs bound with ACE2 (D). A closed spike (three RBDs all in DOWN-states, PDB: 8D55, blue color) was aligned to a preopen spike (one RBD in intermediate-state, two in DOWN-states, PDB: 8D5A, yellow color), showing that the rising of RBD was accompanied by an increase in *α* (E) and an increase in *β* (F). C*α* atoms of C361 and S514 were drawn in ball-representations, blue in closed spike and red in preopen spike.

Examining the pathways, *α* value falls down immediately after RBD transitioning to UP state, indicating the involuntary increase in *α* during this transition, followed by a quick and spontaneous relax right after crossing an energetic barrier. This phenomenon agreed with the findings that a small activation free-energy barrier existed, which was ∼0.6 kcal/mol in FRET experiments^56^ and ∼1 kcal/mol in simulation studies^74^. Comparing the RBD structures (Fig. 5E), a closed spike-crown tends to lock its trimeric RBDs all in the DOWN conformation via the interactions with their neighbors (including NTDs, see Fig. 1A). This may force RBD to swing its head during rising (*α* increases), as if to liberate itself from the ties made by its neighbors. When RBD rotates up, its *β* value is usually larger than 40º (Figs. 5A-5D). Based on the observation that only few populations existed between *β* = 20º ∼ 35º, in this study, *β* = 35º was chosen as a criterion to divide the DOWN and UP states of RBDs.

Compared to the unbound spikes (Fig. 5C), *β* of an UP-state RBD increased in both the antibody-bound (Fig. 5B) and ACE2-bound spikes (Fig. 5D). Apparently, antibodies can elicit greatly diversified *β* values, extending over 140º in space (Fig. 5B). In addition, perhaps unexpectedly, antibody binding also increased the DOWN-state RBD *β* values (comparing Fig. 5B and Fig. 5C).

### The crown-opening pathway

The trimeric assembly of RBDs increases the complexity of tracking the spike-crown opening event, which requires capturing not one but three RBDs in their individual states. The spike structures examined in this study show that the three RBDs seldom synchronize during rising, and these are easily measured by their distinguishable *β* values. To track spike-crown opening, three RBDs on each spike were ordered counterclockwise when viewed above (Fig. 6A), and the first RBD was defined to have the largest *β* value.

**Figure 6.**
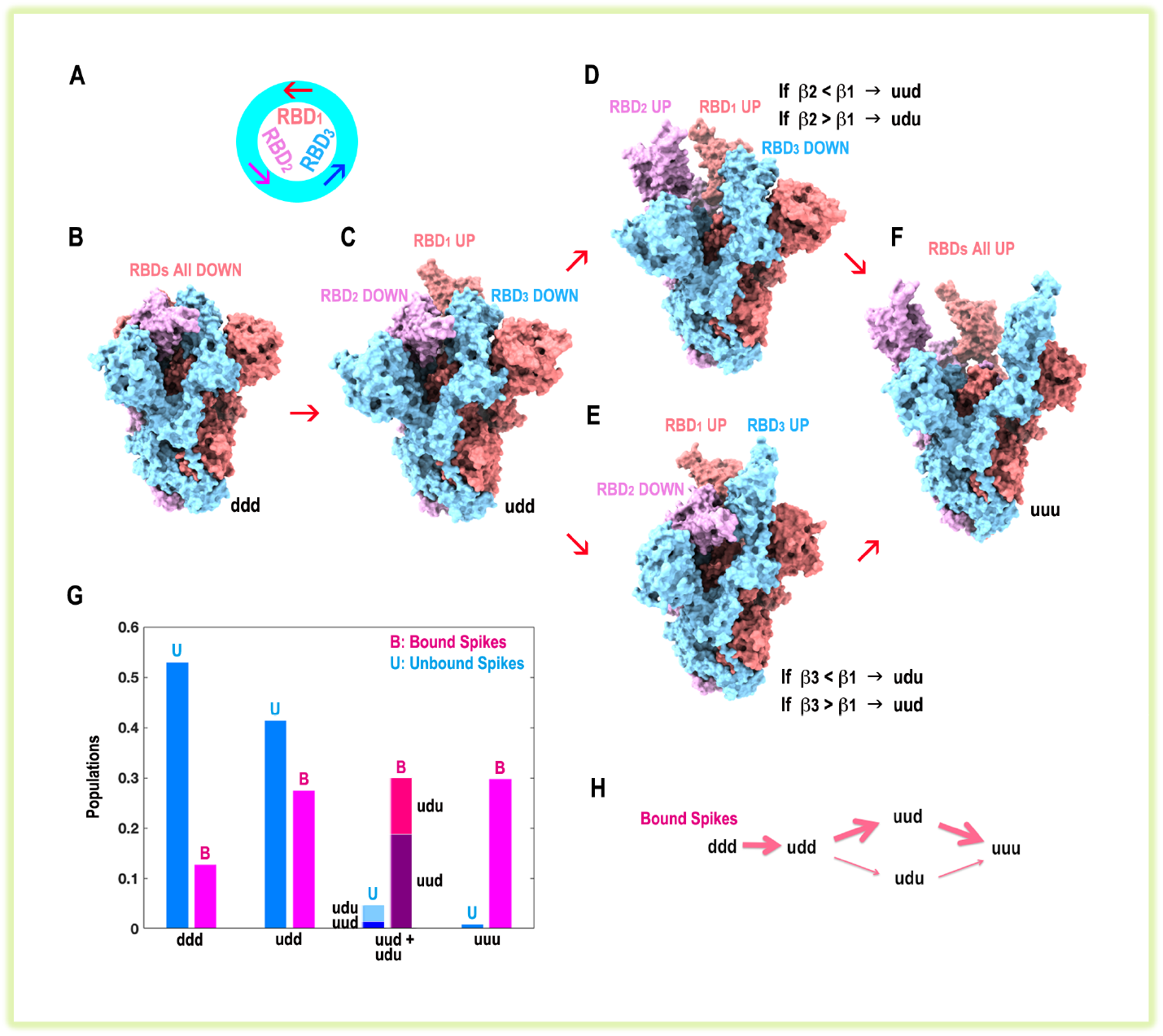
Spike-crown opening pathway described by sequential cases. (A) The trimeric RBDs on a spike were ordered counterclockwise. (B) A closed spike (PDB: 8DLW) represents a *ddd* case. (C) A partly closed spike (PDB: 7W98) represents a *udd* case. (D) A partly open spike with RBD_1_ and RBD_2_ in UP-states. This can represent a *uud* case (PDB: 7DK4) if *β*_2_ < *β*_1_ or a *udu* case if *β*_2_ > *β*_1_. (E) A partly open spike with RBD_1_ and RBD_3_ in UP-states. This can represent a *udu* case (PDB: 7WOR) if *β*_3_ < *β*_1_ or a *uud* case if *β*_3_ > *β*_1_. (F) An open spike (PDB: 7JVC) represents a *uuu* case. (G) Populations of different cases were grouped and normalized in either the bound spikes (magenta) or the unbound spikes (blue). (H) A crown-opening pathway was proposed for the bound spikes, where the majority of them went through the *uud* case.

Following these definitions, the crown-opening track involves at least four cases composed of five instances (Figs. 6B-6F): the closed case (all RBDs are in DOWN states, denoted as *ddd*), the partly closed case (one RBD is UP, denoted as *udd*), the partly open case (two RBDs are UP, denoted as *uud* and *udu*, depending on the location of the DOWN-RBD), and the fully open case (all RBDs are UP, denoted as *uuu*). Note that *uud* and *udu* cases are non-interchangeable due to the definition that the first RBD always has the largest *β* value.

Now counting each case over all structures examined (Fig. 6G), an outline of the crown-opening pathway is sketched in Fig. 6H for the bound spikes (binding either antibodies or ACE2). Apparently, two routes are viable (Fig. 6H). They distinguish in the branches through either the *uud* case (more structures, major route) or the *udu* case (fewer structures, minor route). To eliminate the cases where the two UP-RBD *β* values were similar, the populations of *uud* and *udu* cases were compared further by imposing a large difference in the UP-RBD *β* values (Fig. S5), which proved that the *uud* case always exceeded the *udu* case in the bound-spikes.

The *uud* case is likely evolved from RBD_3_ rising from the *udd* case, whereas the *udu* case is likely evolved from RBD_2_ rising (maybe mixed with RBD_3_ rising) from the *udd* case (Figs. 7A and 7B). This proposal was made assuming that the rising of RBD_2_ did not affect the UP-RBD *β*_1_ value. Comparing the UP-RBD *β* values in three cases (Fig. 7A), one can easily see that *β*_1_ of *u*_1_ (*u*_1_ denotes RBD_1_ in UP state) in *u*_1_*dd* case, *β*_2_ of *u*_2_ in *u*_1_*u*_2_*d* case, and *β*_3_ of *u*_3_ in *u*_1_*du*_3_ case were almost same. Combining the results of Fig. 6G, this means that RBD_3_ is more likely to rise ahead of RBD_2_ while RBD_1_ is in UP state, so that the *uud* case is more abundant subsequently. This proposal was supported by an energetic feature found in an intermediate-state RBD structure (Fig. 7C) that a temporary but strong electrostatic repulsion existed during RBD_1_’s rising that tended to push RBD_3_ away or facilitate RBD_3_’s rising. Another evidence, in accordance with this proposal, was found while analyzing the trimeric RBD *β* values in the *udd* case (Fig. 8), where a rare bound-spike bearing *β*_3_ ≈ 27º and *β*_2_ ≈ −3º was observed and another rare unbound-spike bearing *β*_3_ ≈ 13º and *β*_2_ ≈ −10º was observed, both indicating that RBD_3_ may rise ahead of RBD_2_.

**Figure 7.**
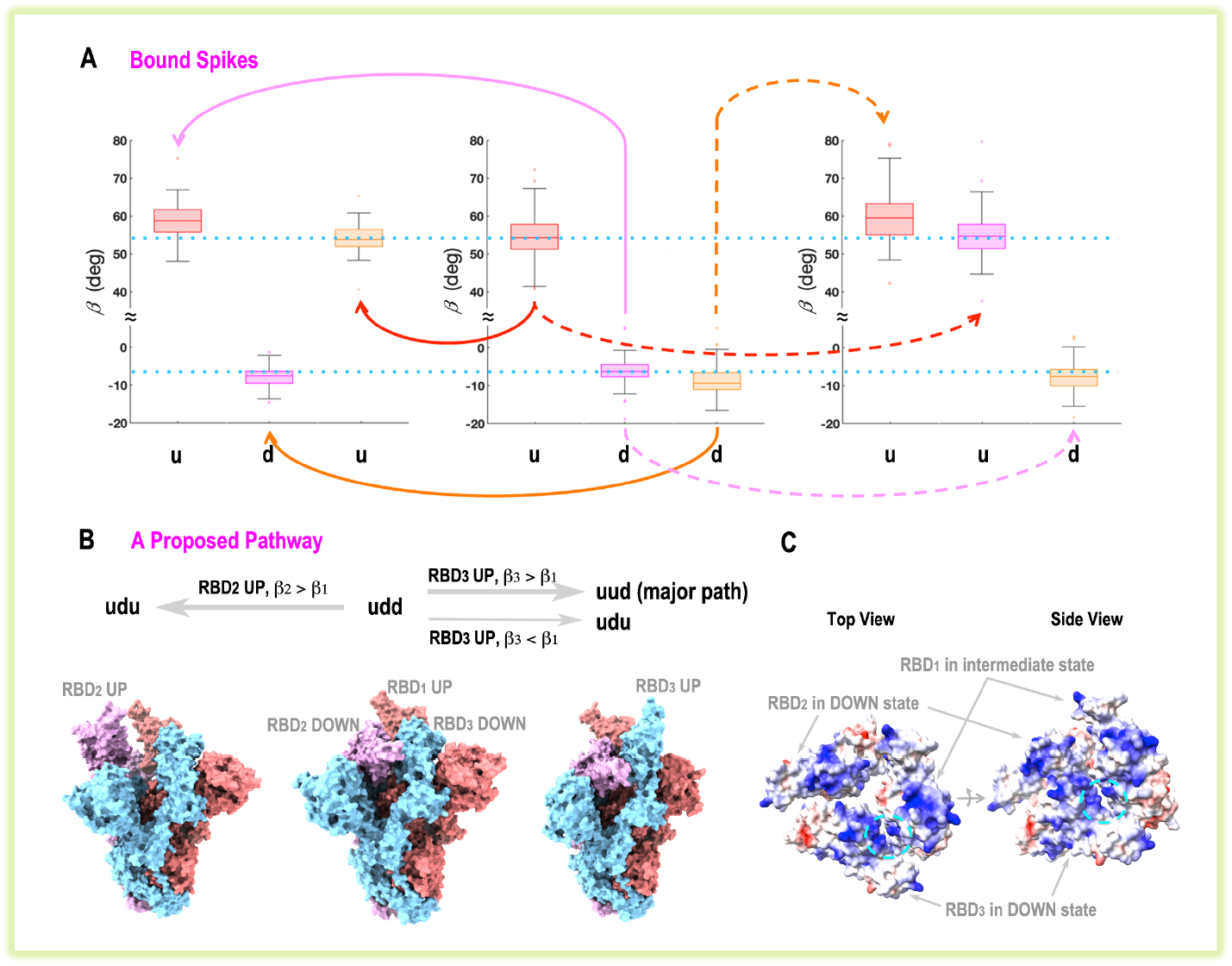
Proposed pathways from a partly closed to a partly open spike. (A) Box plots of *β* values of the trimeric RBDs in *udd, uud*, and *udu* cases, where the proposed major pathways were sketched by solid lines (RBD_2_ rising from *udd* to *udu* case, *β*_2_ > *β*_1_) and dashed lines (RBD_3_ rising from *udd* to *uud* case, *β*_3_ > *β*_1_). Data having *β* > 80º in UP-RBD or *β* > 10º in DOWN-RBD were excluded from averaging. An uncertain minor-pathway formed by RBD_3_ rising from *udd* to *udu* case (*β*_3_ < *β*_1_) was not sketched here. Dotted-blue lines were drawn at the *β*-centers of the UP-RBD_1_ (*u*_1_) and the DOWN-RBD_2_ (*d*_2_) of *u*_1_*d*_2_*d*_3_ case to facilitate comparisons across plots. (B) The proposed pathways were schematically illustrated by three spike structures: *udu* (7WOR), *udd* (7W98), and *uud* (7DK4), drawn from left to right. (C) Coulomb-potential surface of the trimeric RBDs in a preopen spike, where only RBD_1_ was in intermediate-state (*β*_1_ = 30.4º), and the other two were in DOWN-states (PDB: 8D5A). The rising of RBD_1_ exerted a strong electrostatic repulsion to RBD_3_, highlighted in the cyan circles. The Coulomb-potential surfaces were calculated and rendered by ChimeraX^90^, where the blue and red colors indicate the two extremes of the positive and negative electrostatic potentials.

**Figure 8.**
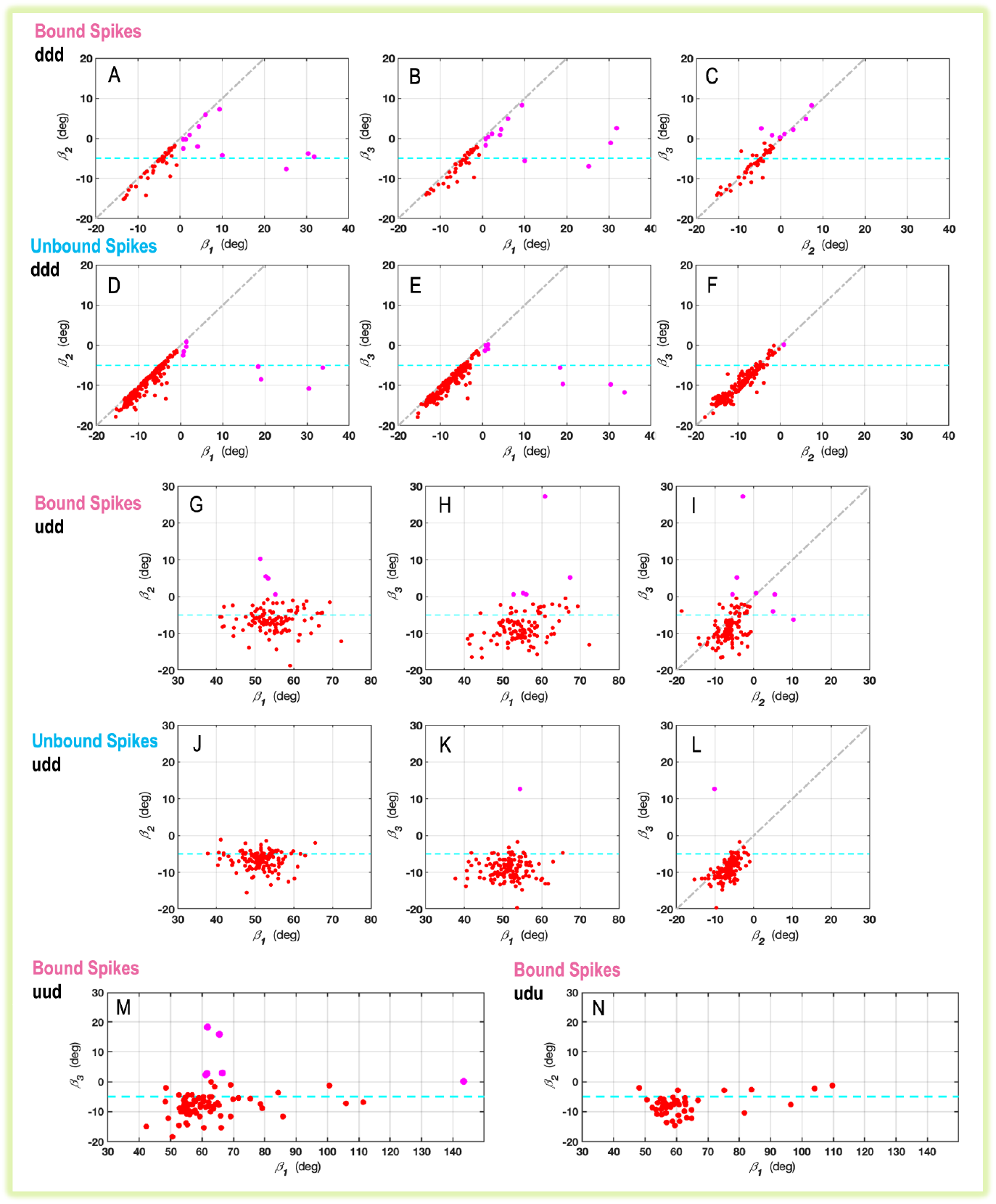
Pairwise correlations of *β* values in different cases. The selected cases were the *ddd* cases for bound (A-C) and unbound (D-F) spikes, the *udd* cases for bound (G-I) and unbound (J-L) spikes, the *uud* case for bound spikes (M), and the *udu* case for bound spikes (N). All cyan lines were drawn at *β* = −5º, merely severing as an aid to compare *β* values across plots. The data were colored magenta when *β* > 0º in a DOWN-state RBD.

In the unbound-spike scenario, there were uncertainties due to the few structures examined in the *uud* and *udu* cases. Nevertheless, based on these few cases, a tentative crown-opening pathway for the unbound-spikes was proposed (Fig. S6), where the *udu* case (11 spike structures) exceeded the *uud* case (5 spike structures).

### Asymmetric rising preference of the trimeric RBDs on individual spikes

The trimeric RBDs were examined in a pairwise manner by comparing their *β* values in adjacent protomers (Fig. 8). In a crown-closed spike (*ddd* case), binding ACE2 or antibodies (bound spikes, Figs. 8A-8C) apparently promotes the rising of RBD with the average *β* increased by at least 4º compared to the unbound spikes (Figs. 8D-8F). Of these crown-closed structures, no spike has two RBDs having *β* larger than 10º concurrently, indicating the stepped rising of RBDs, i.e., three RBDs rise one by one.

In the *udd* case, RBDs of the bound spikes (Figs. 8G-8I) have increased *β* values in both DOWN- and UP-states, compared to those of the unbound spikes (Figs. 8J-8L). In addition, although both RBD_2_ and RBD_3_ are in DOWN states, they have apparently different *β* values (Figs. 8I and 8L), clearly, *β*_2_ > *β*_3_ on average. Under the *udd* case, no concurrent rise of two RBDs at DOWN-state (*β* > 10º) was observed, as in the *ddd* case.

When two RBDs are in UP states (*uud* and *udu* cases, Figs. 8M and 8N; only bound-spike data were plotted here, the unbound-spike plots were omitted because of the few data examined), their opening scales significantly increase compared to the UP-state RBD in *udd* case, where the average *β*_1_ value (averaging only when *β*_1_ ≤ 80º) increases by about 5º in *uud* case and 4º in *udu* case. Within each case, the two UP-state *β* values are distinguishable. <*β*_1_ - *β*_2_> ≈ 5.2º in *uud* case and <*β*_1_ - *β*_3_> ≈ 4.7º in *udu* case (brackets denote the average). These *β* values increase further when all RBDs are in UP states (*uuu* case, Fig. S7), where the three RBDs usually have different *β* values. <*β*_1_ - *β*_2_> ≈ 5.2º and <*β*_1_ - *β*_3_> ≈ 4.5º in *uuu* case (averaging only when *β* ≤ 80º).

## Discussion

Parameters *β* and *α* are **quick** measures to examine the rising scale of the trimeric RDBs on an opening spike-crown. The standard deviations of calculating *β* and *α*, using aligned RBD structures, both were below 2.4º (see Methods). Although this can raise concerns were one to examine any tiny differences in the angles, these errors are usually tolerable compared to the wide spans of *β* and *α* across the entire crown-opening pathway (Fig. 5A). Usually, to minimize the effect of structural flexibility, the center of a group of atoms, but not a single atom, was used to characterize the structural features^65,70^. Here, however, only two C*α* atoms were used. But they were selected carefully that one was located in the beta sheet (S514) and the other was locked by the disulfide bond (C361). Thus, both were minimally susceptible to any structural fluctuations. The viability of this selection was also checked by calculating *α* and *β* using the centers of two groups of C*α* atoms, however, these calculations did not improve the precision but incurred slightly larger standard deviations (see Methods).

The advantage of using *β* and *α* is the easier **quantification** of RBD-rising angles. Visual inspection of a spike structure can easily discern a DOWN- or an UP-state RBD, or even an intermediate-state RBD, but can hardly tell their rising scales, nor differentiate any two RBDs both in the same state bearing similar conformations. In these situations, *β* and *α* can help. An application is to detect a series of intermediate states along the RBD-rising pathway (such as the structures represented by the red dots connecting the DOWN and UP centers in Figs. 5A-5C). Figure 9 shows one instance of these intermediate-state RBDs, easily recognized and compared by its *β* and *α* values. This means that *β* and *α* can help classify and annotate spike structures. This can be useful in several circumstances, for example, suggesting sequential intermediate-state spike structures for the molecular dynamics simulations.

**Figure 9.**
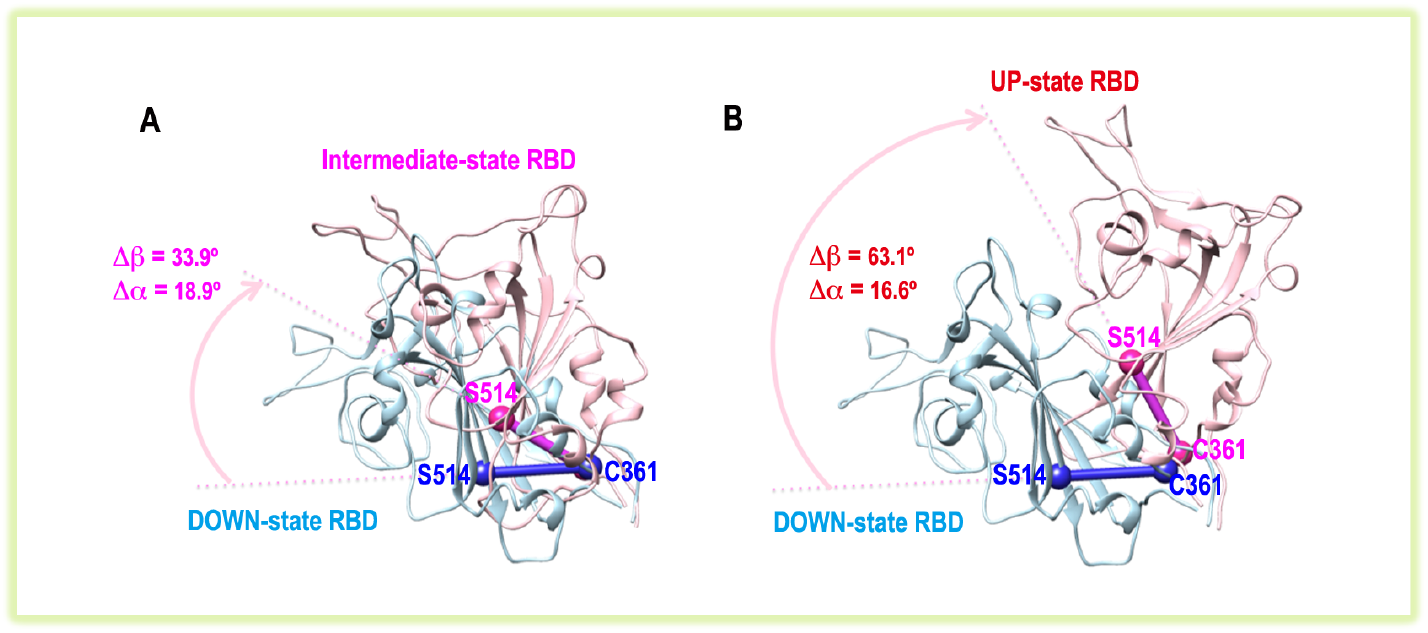
Intermediate-state RBD structures detected by *β* and *α* values. An intermediate-state RBD (on protomer C of PDB: 8D5A, pink color), rotated up from a DOWN-state RBD (on protomer B of PDB: 8DLW, blue color) shown in (A), was compared with an UP-state RBD (on protomer C of PDB: 7W98, pink color) shown in (B).

The unrecognized state in FRET experiments^56^, namely the high FRET-signal (0.8) state, was commonly accepted to associate with a more compactly assembled spike crown, i.e., a DOWN-state RBD having a small *β* value (the ultra-left points in Fig. 5A). Now this can be checked using *β* plotted against *d*, the distance between the donor and acceptor fluorophores inserted into the same spike protomer for conducting the FRET experiment^56^. As RBD rises, this pair distance elongates, resulting in low FRET signals^56^. Conversely, a high FRET signal should correlate with a smaller distance. Indeed, a small *β* value correlates with a small *d* value (Fig. 10A), and hence a large FRET signal, as expected. However, surprisingly enough, an intermediate-state RBD of *β* ≈ 32º was found to associate with an even smaller *d* of around 35 Å (*d* ≈ 42 Å at the DOWN-state RBD population center in Fig. 10A). Checking the spike structure (Figs. 10B-10D), this intermediate-state RBD, although missing residues at the Receptor-Binding-Motif region, clearly showed a large shift in the loop containing D427 (Fig. 10D). This can shorten the donor-acceptor pair distance and may give a high FRET signal. Although this hypothesis needs to be verified by experiments, it highlights another possibility that an intermediate-state RBD with an altered loop conformation may contribute to the high FRET signal.

**Figure 10.**
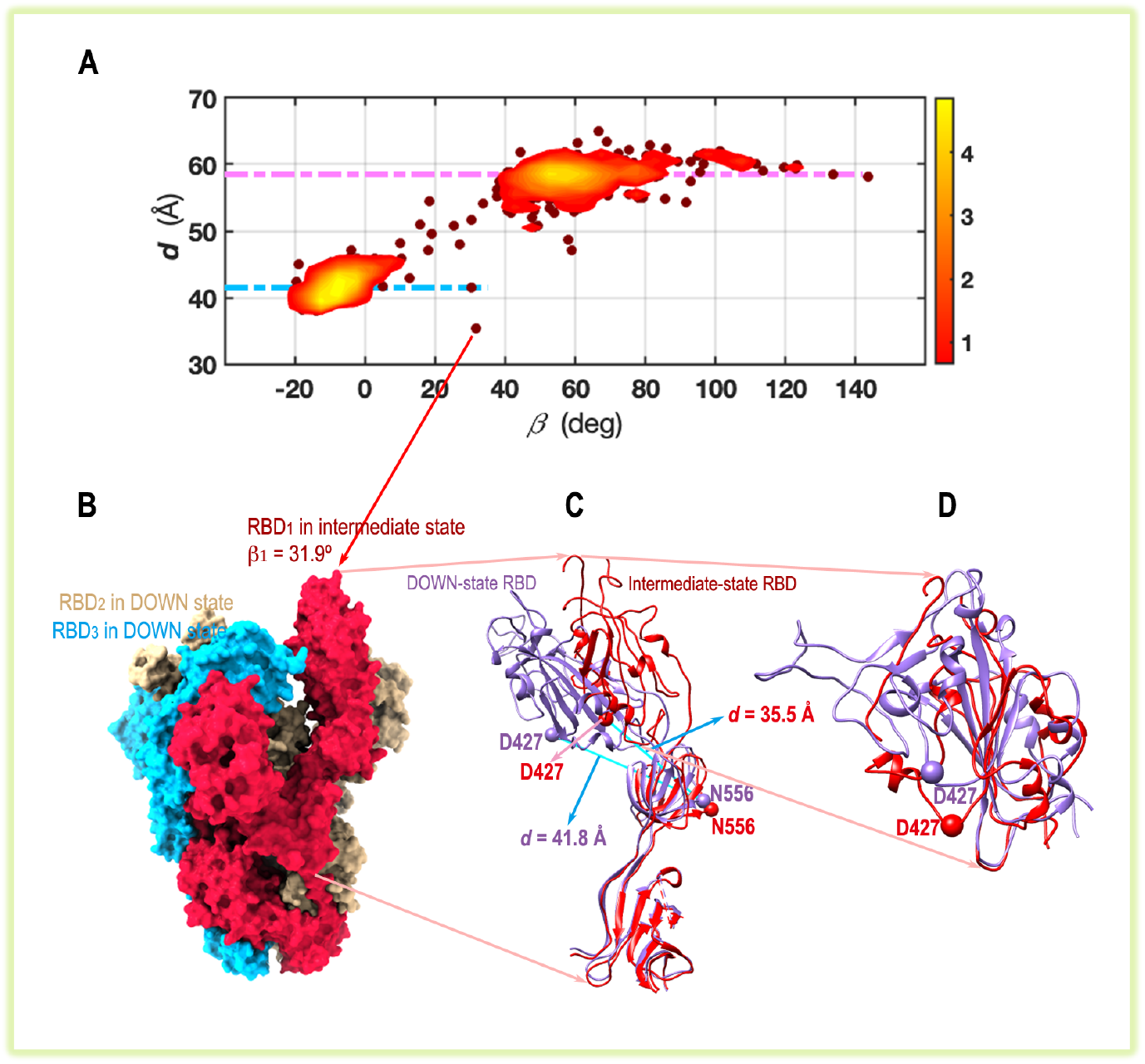
A hypothesis relating an intermediate-state RBD to the high FRET-signal state. (A) The locations of the donor and acceptor fluorophores employed in FRET experiment^56^ were approximately represented by the C*α* atoms of residues D427 and N556, and their distance *d* was plotted against RBD’s *β* value. Dashed-cyan lines marked *d* values at the DOWN- and UP-RBD population centers, spanning a wide range in *β*, especially among the UP-state RBDs. (B) The spike-structure (PDB: 7LJR), having the smallest distance value (*d* = 35.5 Å and *β* = 31.9º) detected in (A), showed that its RBD_1_ was in an intermediate state. (C) This intermediate-state RBD (PDB: 7LJR, red) was compared with a DOWN-state RBD (*d* = 41.8 Å, PDB: 8DLW, purple), that clearly showed a shortened distance between C*α* atoms of D427 and N556. (D) This shortened distance was due to the downward movement of the loop containing D427, clearly seen when the two RBDs were aligned.

While designing therapeutic agents targeting RBDs, one should be aware that RBDs can rise a bit more upon binding substrates (the average *β* of bound-spike-RBDs increased by more than 4º in DOWN-state of *ddd* case and more than 3º in UP-state of *udd* case compared to those of the unbound-spike-RBDs in the same cases, see Fig. 8). Thus, researchers may choose more appropriate spike structures for their design process, and this can be guided by *β* and *α* values. In addition, the asymmetric rising preference of the trimeric RBDs may also be considered, that RBDs tend to rise one by one but not synchronized (at least for the structures examined here).

One limitation of current work is that the calculations of *β* and *α* depend on the quality of RBD structures, particularly, the fluctuations of the C*α* atoms of S514 and C361. Although the motion of the latter is highly restrained by the disulfide bond, it was found that 9 out of 787 structures had RBDs bearing altered loops containing C361. And these structures were removed from the analyses. This issue will be addressed in the future work.

In conclusion, the current work presents a general RBD-rising pathway described by *β* and *α* parameters, which shows a twisted RBD (*α* increases) upon rising (*β* increases). The trimeric RBDs on spike-crown do not rise simultaneously, and usually carry distinguishable rising angles. Antibodies can trigger the opening of spike crowns, preferring the *uud* route and eliciting an increase in RBD’s rising scale. The *β* and *α* parameters introduced in this study may help classify and annotate spike structures and, in particular, detect the intermediate-state RBDs.

## Materials and methods

### Structure selection and alignment

778 coronavirus spike structures were selected from Protein Data Bank (https://www.rcsb.org), all bearing the UniProt accession code P0DTC2. They were all trimers, assembled by full-length or nearly full-length (> 600 residues per protomer) trimeric protomers, solved by either cryo-electron microscopy or x-ray crystallography methods, with resolutions no more than 4 Å.

These structures were then aligned to a reference spike structure (PDB: 8DLW, resolution 2.16 Å), using the software ChimeraX (https://www.rbvi.ucsf.edu/chimera)^90^. Before the alignment, the reference structure was placed at the coordinate center to facilitate the subsequent calculations (the C*α*-atom positions of this structure were provided in the supplementary file). In this coordinate frame, the spike’s trimeric RBDs have these (*α, β*) values: (−0.07º, −3.63º), (−0.08º, −3.56º), and (0.08º, −3.49º) on protomers E, A, and B, respectively. In aligning the structures, only C*α* atoms of the selected helical segments on S2 subunits were used. These were the C*α* atoms of residues 758-783, 946-967, and 986-1032 on each protomer (see their structures in Fig. S1C). All residue numbers presented in this study follow those defined in the reference structure (PDB: 8DLW).

A bound-spike was defined as a spike binding antibody or ACE2. In any spike structure, when any RBD-atom of residues 331-528 was in the vicinity of a non-spike atom (e.g., on antibodies or ACE2) having a distance of less than 4 Å, this RBD was regarded as a bound RBD, otherwise it was considered as an unbound RBD. Here, only non-hydrogen atoms were used in distance calculations. A bound-spike carried at least one bound-RBD, and an unbound-spike had three unbound-RBDs.

All structures presented in this study were visualized by ChimeraX or Chimera (https://www.rbvi.ucsf.edu/chimera)^90^.

### Center-of-mass (COM) calculations

The four domains were defined by four blocks of residues: 15-306 for NTD, 331-528 for RBD, 532-588 for SD1, and 591-675 for SD2. Their COM movements were calculated by C*α* atoms of the corresponding residues. The three spike-protomers need be compared in the same frame, so the COM domain-vectors on the second and third protomers were rotated around z-axis by −120º and 120º, respectively. Each COM vector was then defined as *v*_1_. Meanwhile, the COM vector of the same domain on the first protomer in the reference structure (PDB: 8DLW) was also calculated and was defined as *v*_0_. Here, only the C*α* atoms existing in both the aligned and the reference structures were used in calculating *v*_1_ and *v*_0_. This was to minimize the effect of missing residues during structure comparisons.

Then *v*_1_ was grouped into the DOWN (*β* ≤ 35º) or UP (*β* > 35º) clusters depending on the *β* value of RBD on that protomer. And the results were presented in Fig. 1. Note that the DOWN and UP symbols refer to only the RBD state, although they were used for all domains.

Meanwhile, *v*_1_ – *v*_0_ was also calculated and was defined as *p*_1_, which was subsequently grouped into the DOWN (*β* ≤ 35º) or UP (*β* > 35º) clusters as well. *p*_1_ will be used in the following calculations.

### Calculations of the characteristic distance *d*

TRMEAN method was used in calculating the centers of the DOWN and UP clusters of *p*_1_ vectors (described above) in each domain, where data having any of the three-dimensional coordinate values (x, or y, or z) located at ±3% edge in that dimension were trimmed. Then *p*_1_, subtracting the DOWN-center vector, was redefined as *q*_1_. The characteristic distance *d* of a domain was defined as the scalar projection of each *q*_1_ onto the unit vector directed from the DOWN to the UP center of that domain (one instance was drawn in Fig. 2A).

### Calculations of the spatial parameters Δz, Δ*d*_xy_, and Δ*θ*

The COM vector of each domain *v*_1_ was compared to the COM vector *v*_0_ of the same domain on the first protomer of the reference spike structure (PDB: 8DLW). Δz is the difference between the projections of *v*_1_ and *v*_0_ on the z-axis. Δ*d*_xy_ is the length difference between the projections of *v*_1_ and *v*_0_ onto the x-y plane, i.e., *d*_1_ – *d*_0_. Δ*θ* is the angle difference between the projections of *v*_1_ and *v*_0_ onto the x-y plane, i.e., the angle between *d*_0_ and *d*_1_. Δ*θ* is positive if *d*_0_ rotates to *d*_1_ clockwise when viewed above. An instance was shown in Fig. 3A.

### Definitions of the RBD rotation angles *α* and *β*

To characterize RBD hinge-like rotations, two RBD-atoms were selected, which were the C*α* atoms of S514 and C361. In Fig. 4B, these two C*α* positions were denoted as P1 and P2, based on which P3 was calculated. P3 was on the line connecting P1 and P2, and the distance between P1 and P3 was 18.3 Å. Now the x-z plane (defined in the reference-structure frame for calculating RBD_1_ parameters) passing P3 was defined as *π*_1_. P4 was the projection of P1 onto *π*_1_. P5 (on *π*_1_) was the projection of P4 onto the line (on *π*_1_) passing P3, parallel to the negative x-axis. In this frame, *α* was the angle defined by P4-P3-P1, and *β* was the angle defined by P5-P3-P4. In Fig. 4B, for clarity, all positions were translated relative to P3, so that P3 was located at the origin, and P5 was on the negative x-axis.

Note that P3 serves as a reference position only, and its length to P1 (18.3 Å used here) does not affect *α* or *β* values. This length was chosen simply to make the vectors (directed from P1 to P2) in different RBD structures converge and end at seemingly nearby locations, so that P3 was more like a hinge-point and *β* was more like a rotation-angle around this hinge-point. P3 motions were checked in Fig. S4. Here, the P3 vector was first shifted by the COM vector of its SD1 domain. Then P3_ref_, the P3 vector of the reference structure (also shifted by COM vector of the reference structure’s SD1 domain), was subtracted from P3 to obtain ΔP3. This implementation was to minimize the effect of the SD1 movement during the RBD-rising process. Then ΔP3 vectors were averaged into the DOWN and UP centers using the TRMEAN method described above. Now each ΔP3, after subtracting the DOWN-center vector, was projected onto the unit vector directed from the DOWN to UP center, and the scalar projection was defined as *d*_P3_ that was drawn in Fig. S4.

The precisions of calculating *α* and *β* were checked by further structural alignments. This time, each of the trimeric RBDs on a spike structure was aligned to the same reference RBD structure (located on protomer B of 8DLW, rotated by 120º around z-axis). All the alignments were done using ChimeraX^90^. After the alignments, *α* and *β* values of each RBD were calculated. The standard deviations of *α* and *β* were 2.387º and 2.392º, respectively.

Using these aligned RBD structures, *α* and *β* angles were also redesigned by two groups of C*α* atoms, where the center of C*α* of residues 394-396 and 513-515 were used to represent P1, and the center of C*α* of residues 358-362 were used to represent P2. In this construction, *α* and *β* were recalculated, and their standard deviations were 2.648º and 2.598º, respectively.

### Strategies for drawing the contour plots

The conventional contour plots were drawn in the natural logarithmic scales. But these plots lack the less-sampled data bins, which are essential to show the intermediate and rare states. In this regard, data constructing the bins of no more than 10 data counts were explicitly drawn as red dots below the contour plots, so as to illustrate any transition pathways connecting the DOWN to UP clusters. In all contour plots presented, the color bar on the right-hand side shows the populations, labeled in natural logarithmic scale.

## Supplementary Materials

Supplementary Figures S1 to S7.

Supplementary Data: C*α*-atom positions of the reference spike structure used in this study, which were adapted from the PDB structure 8DLW.pdb.

## Acknowledgments

The author gratefully acknowledges all structural biologists who solved the coronavirus spike structures and published them on PDB website. It was their original work that revealed the features of individual spikes and provided the basis for the current study. This work was supported by Fudan University.

## References

1 Gorbalenya, A. E., Baker, S. C., Baric, R. S., de Groot, R. J., Drosten, C., Gulyaeva, A. A., … Ziebuhr, J. The species Severe acute respiratory syndrome-related coronavirus: classifying 2019-nCoV and naming it SARS-CoV-2. Nat Microbiol 5, 536–544, doi:10.1038/s41564-020-0695-z (2020).

2 Zhou, P., Yang, X. L., Wang, X. G., Hu, B., Zhang, L., Zhang, W., … Shi, Z. L. A pneumonia outbreak associated with a new coronavirus of probable bat origin. Nature 579, 270–273, doi:10.1038/s41586-020-2012-7 (2020).

3 Zhu, N., Zhang, D., Wang, W., Li, X., Yang, B., Song, J., … Tan, W. A Novel Coronavirus from Patients with Pneumonia in China, 2019. N Engl J Med 382, 727–733, doi:10.1056/NEJMoa2001017 (2020).

4 https://www.who.int/emergencies/diseases/novel-coronavirus-2019

5 Gobeil, S. M., Janowska, K., McDowell, S., Mansouri, K., Parks, R., Stalls, V., … Acharya, P. Effect of natural mutations of SARS-CoV-2 on spike structure, conformation, and antigenicity. Science 373, doi:10.1126/science.abi6226 (2021).

6 Mykytyn, A. Z., Fouchier, R. A. & Haagmans, B. L. Antigenic evolution of SARS coronavirus 2. Curr Opin Virol 62, 101349, doi:10.1016/j.coviro.2023.101349 (2023).

7 Chen, Q., Zhang, J., Wang, P. & Zhang, Z. The mechanisms of immune response and evasion by the main SARS-CoV-2 variants. iScience 25, 105044, doi:10.1016/j.isci.2022.105044 (2022).

8 Harvey, W. T., Carabelli, A. M., Jackson, B., Gupta, R. K., Thomson, E. C., Harrison, E. M., … Robertson, D. L. SARS-CoV-2 variants, spike mutations and immune escape. Nat Rev Microbiol 19, 409–424, doi:10.1038/s41579-021-00573-0 (2021).

9 Li, P., Faraone, J. N., Hsu, C. C., Chamblee, M., Liu, Y., Zheng, Y. M., … Liu, S. L. Neutralization and spike stability of JN.1-derived LB.1, KP.2.3, KP.3, and KP.3.1.1 subvariants. mBio 16, e0046425, doi:10.1128/mbio.00464-25 (2025).

10 Liu, C., Zhou, D., Dijokaite-Guraliuc, A., Supasa, P., Duyvesteyn, H. M. E., Ginn, H. M., … Screaton, G. R. A structure-function analysis shows SARS-CoV-2 BA.2.86 balances antibody escape and ACE2 affinity. Cell Rep Med 5, 101553, doi:10.1016/j.xcrm.2024.101553 (2024).

11 Mittal, A., Khattri, A. & Verma, V. Structural and antigenic variations in the spike protein of emerging SARS-CoV-2 variants. PLoS Pathog 18, e1010260, doi:10.1371/journal.ppat.1010260 (2022).

12 Stalls, V., Lindenberger, J., Gobeil, S. M., Henderson, R., Parks, R., Barr, M., … Acharya, P. Cryo-EM structures of SARS-CoV-2 Omicron BA.2 spike. Cell Rep 39, 111009, doi:10.1016/j.celrep.2022.111009 (2022).

13 Dadonaite, B., Brown, J., McMahon, T. E., Farrell, A. G., Figgins, M. D., Asarnow, D., … Bloom, J. D. Spike deep mutational scanning helps predict success of SARS-CoV-2 clades. Nature 631, 617–626, doi:10.1038/s41586-024-07636-1 (2024).

14 Bai, C., Wang, J., Chen, G., Zhang, H., An, K., Xu, P., … Warshel, A. Predicting Mutational Effects on Receptor Binding of the Spike Protein of SARS-CoV-2 Variants. J Am Chem Soc 143, 17646–17654, doi:10.1021/jacs.1c07965 (2021).

15 Dadonaite, B., Crawford, K. H. D., Radford, C. E., Farrell, A. G., Yu, T. C., Hannon, W. W., … Bloom, J. D. A pseudovirus system enables deep mutational scanning of the full SARS-CoV-2 spike. Cell 186, 1263-1278.e1220, doi:10.1016/j.cell.2023.02.001 (2023).

16 Greaney, A. J., Starr, T. N., Barnes, C. O., Weisblum, Y., Schmidt, F., Caskey, M., … Bloom, J. D. Mapping mutations to the SARS-CoV-2 RBD that escape binding by different classes of antibodies. Nat Commun 12, 4196, doi:10.1038/s41467-021-24435-8 (2021).

17 Greaney, A. J., Starr, T. N., Gilchuk, P., Zost, S. J., Binshtein, E., Loes, A. N., … Bloom, J. D. Complete Mapping of Mutations to the SARS-CoV-2 Spike Receptor-Binding Domain that Escape Antibody Recognition. Cell Host Microbe 29, 44-57.e49, doi:10.1016/j.chom.2020.11.007 (2021).

18 Moulana, A., Dupic, T., Phillips, A. M., Chang, J., Roffler, A. A., Greaney, A. J., … Desai, M. M. The landscape of antibody binding affinity in SARS-CoV-2 Omicron BA.1 evolution. eLife 12, e83442, doi:10.7554/eLife.83442 (2023).

19 Starr, T. N., Greaney, A. J., Hilton, S. K., Ellis, D., Crawford, K. H. D., Dingens, S., … Bloom, J. D. Deep Mutational Scanning of SARS-CoV-2 Receptor Binding Domain Reveals Constraints on Folding and ACE2 Binding. Cell 182, 1295-1310.e1220, doi:10.1016/j.cell.2020.08.012 (2020).

20 Starr, T. N., Greaney, A. J., Stewart, C. M., Walls, A. C., Hannon, W. W., Veesler, D. & Bloom, J. D. Deep mutational scans for ACE2 binding, RBD expression, and antibody escape in the SARS-CoV-2 Omicron BA.1 and BA.2 receptor-binding domains. PLoS Pathog 18, e1010951, doi:10.1371/journal.ppat.1010951 (2022).

21 Cao, L., Goreshnik, I., Coventry, B., Case, J. B., Miller, L., Kozodoy, L., … Baker, D. De novo design of picomolar SARS-CoV-2 miniprotein inhibitors. Science 370, 426–431, doi:10.1126/science.abd9909 (2020).

22 Ding, S., Ullah, I., Gong, S. Y., Grover, J. R., Mohammadi, M., Chen, Y., … Baron, C. VE607 stabilizes SARS-CoV-2 Spike in the “RBD-up” conformation and inhibits viral entry. iScience 25, 104528, doi:10.1016/j.isci.2022.104528 (2022).

23 Guan, X., Yang, Y. & Du, L. Advances in SARS-CoV-2 receptor-binding domain-based COVID-19 vaccines. Expert Rev Vaccines 22, 422–439, doi:10.1080/14760584.2023.2211153 (2023).

24 Hunt, A. C., Case, J. B., Park, Y. J., Cao, L., Wu, K., Walls, A. C., … Baker, D. Multivalent designed proteins neutralize SARS-CoV-2 variants of concern and confer protection against infection in mice. Sci Transl Med 14, eabn1252, doi:10.1126/scitranslmed.abn1252 (2022).

25 Kim, J. W., Heo, K., Kim, H. J., Yoo, Y., Cho, H. S., Jang, H. J., … Lee, S. Novel bispecific human antibody platform specifically targeting a fully open spike conformation potently neutralizes multiple SARS-CoV-2 variants. Antiviral Res 212, 105576, doi:10.1016/j.antiviral.2023.105576 (2023).

26 Koenig, P. A., Das, H., Liu, H., Kümmerer, B. M., Gohr, F. N., Jenster, L. M., … Schmidt, F. I. Structure-guided multivalent nanobodies block SARS-CoV-2 infection and suppress mutational escape. Science 371, eabe6230, doi:10.1126/science.abe6230 (2021).

27 Li, S., Ye, F., Zheng, Y., Wang, J., Peng, H., Zhu, L., … Li, H. Dual-Locking the SARS-CoV-2 Spike Trimer: An Amphipathic Molecular “Bolt” Stabilizes Conserved Druggable Interfaces for Coronavirus Inhibition. Adv Sci (Weinh) 12, e2417534, doi:10.1002/advs.202417534 (2025).

28 Mikolajek, H., Weckener, M., Brotzakis, Z. F., Huo, J., Dalietou, E. V., Le Bas, A., … Naismith, J. H. Correlation between the binding affinity and the conformational entropy of nanobody SARS-CoV-2 spike protein complexes. Proc Natl Acad Sci U S A 119, e2205412119, doi:10.1073/pnas.2205412119 (2022).

29 Modhiran, N., Lauer, S. M., Amarilla, A. A., Hewins, P., Lopes van den Broek, S. I., Low, Y. S., … Watterson, D. A nanobody recognizes a unique conserved epitope and potently neutralizes SARS-CoV-2 omicron variants. iScience 26, 107085, doi:10.1016/j.isci.2023.107085 (2023).

30 Parray, H. A., Narayanan, N., Garg, S., Rizvi, Z. A., Shrivastava, T., Kushwaha, S., … Kumar, R. A broadly neutralizing monoclonal antibody overcomes the mutational landscape of emerging SARS-CoV-2 variants of concern. PLoS Pathog 18, e1010994, doi:10.1371/journal.ppat.1010994 (2022).

31 Schoof, M., Faust, B., Saunders, R. A., Sangwan, S., Rezelj, V., Hoppe, N., … Manglik, A. An ultrapotent synthetic nanobody neutralizes SARS-CoV-2 by stabilizing inactive Spike. Science 370, 1473–1479, doi:10.1126/science.abe3255 (2020).

32 Dacon, C., Tucker, C., Peng, L., Lee, C. D., Lin, T. H., Yuan, M., … Tan, J. Broadly neutralizing antibodies target the coronavirus fusion peptide. Science 377, 728–735, doi:10.1126/science.abq3773 (2022).

33 Grunst, M. W., Qin, Z., Dodero-Rojas, E., Ding, S., Prévost, J., Chen, Y., … Li, W. Structure and inhibition of SARS-CoV-2 spike refolding in membranes. Science 385, 757–765, doi:10.1126/science.adn5658 (2024).

34 Guo, L., Chen, Z., Lin, S., Yang, F., Yang, J., Wang, L., … Lu, G. Structural basis and mode of action for two broadly neutralizing nanobodies targeting the highly conserved spike stem-helix of sarbecoviruses including SARS-CoV-2 and its variants. PLoS Pathog 21, e1013034, doi:10.1371/journal.ppat.1013034 (2025).

35 Liu, Z., Xu, W., Chen, Z., Fu, W., Zhan, W., Gao, Y., … Wang, Q. An ultrapotent pan-β-coronavirus lineage B (β-CoV-B) neutralizing antibody locks the receptor-binding domain in closed conformation by targeting its conserved epitope. Protein Cell 13, 655–675, doi:10.1007/s13238-021-00871-6 (2022).

36 Suryadevara, N., Kose, N., Bangaru, S., Binshtein, E., Munt, J., Martinez, D. R., … Crowe, J. E., Jr. Structural characterization of human monoclonal antibodies targeting uncommon antigenic sites on spike glycoprotein of SARS-CoV. J Clin Invest 135, e178880, doi:10.1172/jci178880 (2024).

37 Wang, C., van Haperen, R., Gutiérrez-Álvarez, J., Li, W., Okba, N. M. A., Albulescu, I., … Bosch, B. J. A conserved immunogenic and vulnerable site on the coronavirus spike protein delineated by cross-reactive monoclonal antibodies. Nat Commun 12, 1715, doi:10.1038/s41467-021-21968-w (2021).

38 Zhang, Z., Zhang, Y., Zhang, Y., Cheng, L., Zhang, L., Yan, Q., … Zhao, J. Defining the features and structure of neutralizing antibody targeting the silent face of the SARS-CoV-2 spike N-terminal domain. MedComm 5, e70008, doi:10.1002/mco2.70008 (2024).

39 Baum, A., Fulton, B. O., Wloga, E., Copin, R., Pascal, K. E., Russo, V., … Kyratsous, C. A. Antibody cocktail to SARS-CoV-2 spike protein prevents rapid mutational escape seen with individual antibodies. Science 369, 1014–1018, doi:10.1126/science.abd0831 (2020).

40 Dong, J., Zost, S. J., Greaney, A. J., Starr, T. N., Dingens, A. S., Chen, E. C., … Crowe, J. E., Jr. Genetic and structural basis for SARS-CoV-2 variant neutralization by a two-antibody cocktail. Nat Microbiol 6, 1233–1244, doi:10.1038/s41564-021-00972-2 (2021).

41 Li, X., Pan, Y., Yin, Q., Wang, Z., Shan, S., Zhang, L., … Wang, X. Structural basis of a two-antibody cocktail exhibiting highly potent and broadly neutralizing activities against SARS-CoV-2 variants including diverse Omicron sublineages. Cell Discov 8, 87, doi:10.1038/s41421-022-00449-4 (2022).

42 Su, S. C., Yang, T. J., Yu, P. Y., Liang, K. H., Chen, W. Y., Yang, C. W., … Wu, H. C. Structure-guided antibody cocktail for prevention and treatment of COVID-19. PLoS Pathog 17, e1009704, doi:10.1371/journal.ppat.1009704 (2021).

43 Weinreich, D. M., Sivapalasingam, S., Norton, T., Ali, S., Gao, H., Bhore, R., … Yancopoulos, G. D. REGN-COV2, a Neutralizing Antibody Cocktail, in Outpatients with Covid-19. N Engl J Med 384, 238–251, doi:10.1056/NEJMoa2035002 (2021).

44 Hsieh, C. L., Goldsmith, J. A., Schaub, J. M., DiVenere, A. M., Kuo, H. C., Javanmardi, K., … McLellan, J. S. Structure-based design of prefusion-stabilized SARS-CoV-2 spikes. Science 369, 1501–1505, doi:10.1126/science.abd0826 (2020).

45 Juraszek, J., Rutten, L., Blokland, S., Bouchier, P., Voorzaat, R., Ritschel, T., … Langedijk, J. P. M. Stabilizing the closed SARS-CoV-2 spike trimer. Nat Commun 12, 244, doi:10.1038/s41467-020-20321-x (2021).

46 McCallum, M., Walls, A. C., Bowen, J. E., Corti, D. & Veesler, D. Structureguided covalent stabilization of coronavirus spike glycoprotein trimers in the closed conformation. Nat Struct Mol Biol 27, 942–949, doi:10.1038/s41594-020-0483-8 (2020).

47 Ni, T., Mendonça, L., Zhu, Y., Howe, A., Radecke, J., Shah, P. M., … Zhang, P. ChAdOx1 COVID vaccines express RBD open prefusion SARS-CoV-2 spikes on the cell surface. iScience 26, 107882, doi:10.1016/j.isci.2023.107882 (2023).

48 Nuqui, X., Casalino, L., Zhou, L., Shehata, M., Wang, A., Tse, A. L., … Amaro, R. E. Simulation-driven design of stabilized SARS-CoV-2 spike S2 immunogens. Nat Commun 15, 7370, doi:10.1038/s41467-024-50976-9 (2024).

49 Tan, T. J. C., Mou, Z., Lei, R., Ouyang, W. O., Yuan, M., Song, G., … Wu, N. C. High-throughput identification of prefusion-stabilizing mutations in SARS-CoV-2 spike. Nat Commun 14, 2003, doi:10.1038/s41467-023-37786-1 (2023).

50 Xiong, X., Qu, K., Ciazynska, K. A., Hosmillo, M., Carter, A. P., Ebrahimi, S., … Briggs, J. A. G. A thermostable, closed SARS-CoV-2 spike protein trimer. Nat Struct Mol Biol 27, 934–941, doi:10.1038/s41594-020-0478-5 (2020).

51 Jackson, C. B., Farzan, M., Chen, B. & Choe, H. Mechanisms of SARS-CoV-2 entry into cells. Nat Rev Mol Cell Biol 23, 3–20, doi:10.1038/s41580-021-00418-x (2022).

52 Wrapp, D., Wang, N., Corbett, K. S., Goldsmith, J. A., Hsieh, C. L., Abiona, O., … McLellan, J. S. Cryo-EM structure of the 2019-nCoV spike in the prefusion conformation. Science 367, 1260–1263, doi:10.1126/science.abb2507 (2020).

53 Zhang, J., Xiao, T., Cai, Y. & Chen, B. Structure of SARS-CoV-2 spike protein. Curr Opin Virol 50, 173–182, doi:10.1016/j.coviro.2021.08.010 (2021).

54 Turoňová, B., Sikora, M., Schürmann, C., Hagen, W. J. H., Welsch, S., Blanc, F. E. C., … Beck, M. In situ structural analysis of SARS-CoV-2 spike reveals flexibility mediated by three hinges. Science 370, 203–208, doi:10.1126/science.abd5223 (2020).

55 Ke, Z., Oton, J., Qu, K., Cortese, M., Zila, V., McKeane, L., … Briggs, J. A. G. Structures and distributions of SARS-CoV-2 spike proteins on intact virions. Nature 588, 498–502, doi:10.1038/s41586-020-2665-2 (2020).

56 Lu, M., Uchil, P. D., Li, W., Zheng, D., Terry, D. S., Gorman, J., … Mothes, W. Real-Time Conformational Dynamics of SARS-CoV-2 Spikes on Virus Particles. Cell Host Microbe 28, 880-891.e888, doi:10.1016/j.chom.2020.11.001 (2020).

57 Ge, X. Y., Li, J. L., Yang, X. L., Chmura, A. A., Zhu, G., Epstein, J. H., … Shi, Z. L. Isolation and characterization of a bat SARS-like coronavirus that uses the ACE2 receptor. Nature 503, 535–538, doi:10.1038/nature12711 (2013).

58 Lan, J., Ge, J., Yu, J., Shan, S., Zhou, H., Fan, S., … Wang, X. Structure of the SARS-CoV-2 spike receptor-binding domain bound to the ACE2 receptor. Nature 581, 215–220, doi:10.1038/s41586-020-2180-5 (2020).

59 Li, W., Moore, M. J., Vasilieva, N., Sui, J., Wong, S. K., Berne, M. A., … Farzan, M. Angiotensin-converting enzyme 2 is a functional receptor for the SARS coronavirus. Nature 426, 450–454, doi:10.1038/nature02145 (2003).

60 Shang, J., Ye, G., Shi, K., Wan, Y., Luo, C., Aihara, H., … Li, F. Structural basis of receptor recognition by SARS-CoV-2. Nature 581, 221–224, doi:10.1038/s41586-020-2179-y (2020).

61 Tai, L., Zhu, G., Yang, M., Cao, L., Xing, X., Yin, G., … Zhu, Y. Nanometer-resolution in situ structure of the SARS-CoV-2 postfusion spike protein. Proc Natl Acad Sci U S A 118, e2112703118, doi:10.1073/pnas.2112703118 (2021).

62 Wolf, K. A., Kwan, J. C. & Kamil, J. P. Structural Dynamics and Molecular Evolution of the SARS-CoV-2 Spike Protein. mBio 13, e0203021, doi:10.1128/mbio.02030-21 (2022).

63 Yu, S., Hu, H., Ai, Q., Bai, R., Ma, K., Zhou, M. & Wang, S. SARS-CoV-2 Spike-Mediated Entry and Its Regulation by Host Innate Immunity. Viruses 15, 639, doi:10.3390/v15030639 (2023).

64 Gobeil, S. M., Henderson, R., Stalls, V., Janowska, K., Huang, X., May, A., … Acharya, P. Structural diversity of the SARS-CoV-2 Omicron spike. Mol Cell 82, 2050-2068.e2056, doi:10.1016/j.molcel.2022.03.028 (2022).

65 Henderson, R., Edwards, R. J., Mansouri, K., Janowska, K., Stalls, V., Gobeil, S. M. C., … Acharya, P. Controlling the SARS-CoV-2 spike glycoprotein conformation. Nat Struct Mol Biol 27, 925–933, doi:10.1038/s41594-020-0479-4 (2020).

66 Lv, Z., Deng, Y. Q., Ye, Q., Cao, L., Sun, C. Y., Fan, C., … Wang, X. Structural basis for neutralization of SARS-CoV-2 and SARS-CoV by a potent therapeutic antibody. Science 369, 1505–1509, doi:10.1126/science.abc5881 (2020).

67 Pramanick, I., Sengupta, N., Mishra, S., Pandey, S., Girish, N., Das, A. & Dutta, S. Conformational flexibility and structural variability of SARS-CoV2 S protein. Structure 29, 834-845.e835, doi:10.1016/j.str.2021.04.006 (2021).

68 Walls, A. C., Park, Y. J., Tortorici, M. A., Wall, A., McGuire, A. T. & Veesler, D. Structure, Function, and Antigenicity of the SARS-CoV-2 Spike Glycoprotein. Cell 180, 281–292, doi:10.1016/j.cell.2020.02.058 (2020).

69 Casalino, L., Gaieb, Z., Goldsmith, J. A., Hjorth, C. K., Dommer, A. C., Harbison, M., … Amaro, R. E. Beyond Shielding: The Roles of Glycans in the SARS-CoV-2 Spike Protein. ACS Cent Sci 6, 1722–1734, doi:10.1021/acscentsci.0c01056 (2020).

70 Dokainish, H. M., Re, S., Mori, T., Kobayashi, C., Jung, J. & Sugita, Y. The inherent flexibility of receptor binding domains in SARS-CoV-2 spike protein. eLife 11, e75720, doi:10.7554/eLife.75720 (2022).

71 Fallon, L., Belfon, K. A. A., Raguette, L., Wang, Y., Stepanenko, D., Cuomo, A., … Simmerling, C. Free Energy Landscapes from SARS-CoV-2 Spike Glycoprotein Simulations Suggest that RBD Opening Can Be Modulated via Interactions in an Allosteric Pocket. J Am Chem Soc 143, 11349–11360, doi:10.1021/jacs.1c00556 (2021).

72 Mansbach, R. A., Chakraborty, S., Nguyen, K., Montefiori, D. C., Korber, B. & Gnanakaran, S. The SARS-CoV-2 Spike variant D614G favors an open conformational state. Sci Adv 7, eabf3671, doi:10.1126/sciadv.abf3671 (2021).

73 Mori, T., Jung, J., Kobayashi, C., Dokainish, H. M., Re, S. & Sugita, Y. Elucidation of interactions regulating conformational stability and dynamics of SARS-CoV-2 S-protein. Biophys J 120, 1060–1071, doi:10.1016/j.bpj.2021.01.012 (2021).

74 Ovchinnikov, V. & Karplus, M. Free Energy Simulations of Receptor-Binding Domain Opening of the SARS-CoV-2 Spike Indicate a Barrierless Transition with Slow Conformational Motions. J Phys Chem B 127, 8565–8575, doi:10.1021/acs.jpcb.3c05236 (2023).

75 Pang, Y. T., Acharya, A., Lynch, D. L., Pavlova, A. & Gumbart, J. C. SARS-CoV-2 spike opening dynamics and energetics reveal the individual roles of glycans and their collective impact. Commun Biol 5, 1170, doi:10.1038/s42003-022-04138-6 (2022).

76 Singh, J., Vashishtha, S. & Kundu, B. Spike Protein Mutation-Induced Changes in the Kinetic and Thermodynamic Behavior of Its Receptor Binding Domains Explain Their Higher Propensity to Attain Open States in SARS-CoV-2 Variants of Concern. ACS Cent Sci 9, 1894–1904, doi:10.1021/acscentsci.3c00810 (2023).

77 Sztain, T., Ahn, S. H., Bogetti, A. T., Casalino, L., Goldsmith, J. A., Seitz, E., … Amaro, R. E. A glycan gate controls opening of the SARS-CoV-2 spike protein. Nat Chem 13, 963–968, doi:10.1038/s41557-021-00758-3 (2021).

78 Zimmerman, M. I., Porter, J. R., Ward, M. D., Singh, S., Vithani, N., Meller, A., … Bowman, G. R. SARS-CoV-2 simulations go exascale to predict dramatic spike opening and cryptic pockets across the proteome. Nat Chem 13, 651–659, doi:10.1038/s41557-021-00707-0 (2021).

79 Lu, M. Single-Molecule FRET Imaging of Virus Spike-Host Interactions. Viruses 13, 332, doi:10.3390/v13020332 (2021).

80 Yang, Z., Han, Y., Ding, S., Shi, W., Zhou, T., Finzi, A., … Lu, M. SARS-CoV-2 Variants Increase Kinetic Stability of Open Spike Conformations as an Evolutionary Strategy. mBio 13, e0322721, doi:10.1128/mbio.03227-21 (2021).

81 Calvaresi, V., Wrobel, A. G., Toporowska, J., Hammerschmid, D., Doores, K. J., Bradshaw, R. T., … Politis, A. Structural dynamics in the evolution of SARS-CoV-2 spike glycoprotein. Nat Commun 14, 1421, doi:10.1038/s41467-023-36745-0 (2023).

82 Costello, S. M., Shoemaker, S. R., Hobbs, H. T., Nguyen, A. W., Hsieh, C. L., Maynard, J. A., … Marqusee, S. The SARS-CoV-2 spike reversibly samples an open-trimer conformation exposing novel epitopes. Nat Struct Mol Biol 29, 229–238, doi:10.1038/s41594-022-00735-5 (2022).

83 Lim, K., Nishide, G., Yoshida, T., Watanabe-Nakayama, T., Kobayashi, A., Hazawa, M., … Wong, R. W. Millisecond dynamic of SARS-CoV-2 spike and its interaction with ACE2 receptor and small extracellular vesicles. J Extracell Vesicles 10, e12170, doi:10.1002/jev2.12170 (2021).

84 Saha, P., Fernandez, I., Sumbul, F., Valotteau, C., Kostrz, D., Meola, A., … Rico, F. Modulation of SARS-CoV-2 spike binding to ACE2 through conformational selection. Nat Nanotechnol 20, 926–934, doi:10.1038/s41565-025-01908-1 (2025).

85 Bangaru, S., Ozorowski, G., Turner, H. L., Antanasijevic, A., Huang, D., Wang, X., … Ward, A. B. Structural analysis of full-length SARS-CoV-2 spike protein from an advanced vaccine candidate. Science 370, 1089–1094, doi:10.1126/science.abe1502 (2020).

86 Watanabe, Y., Allen, J. D., Wrapp, D., McLellan, J. S. & Crispin, M. Site-specific glycan analysis of the SARS-CoV-2 spike. Science 369, 330–333, doi:10.1126/science.abb9983 (2020).

87 Wang, Y., Xu, C., Wang, Y., Hong, Q., Zhang, C., Li, Z., … Cong, Y. Conformational dynamics of the Beta and Kappa SARS-CoV-2 spike proteins and their complexes with ACE2 receptor revealed by cryo-EM. Nat Commun 12, 7345, doi:10.1038/s41467-021-27350-0 (2021).

88 Zhang, J., Cai, Y., Xiao, T., Lu, J., Peng, H., Sterling, S. M., … Chen, B. Structural impact on SARS-CoV-2 spike protein by D614G substitution. Science 372, 525–530, doi:10.1126/science.abf2303 (2021).

89 Zhang, J., Tang, W., Gao, H., Lavine, C. L., Shi, W., Peng, H., … Chen, B. Structural and functional characteristics of the SARS-CoV-2 Omicron subvariant BA.2 spike protein. Nat Struct Mol Biol 30, 980–990, doi:10.1038/s41594-023-01023-6 (2023).

90 Pettersen, E. F., Goddard, T. D., Huang, C. C., Couch, G. S., Greenblatt, D. M., Meng, E. C. & Ferrin, T. E. UCSF Chimera--a visualization system for exploratory research and analysis. J Comput Chem 25, 1605–1612, doi:10.1002/jcc.20084 (2004).

