## Supplementary Materials for "Tracking the opening of spike crowns on the surface of coronaviruses"

**Di Wu***

*Department of Physiology and Neurobiology, School of Life Sciences, Fudan University, Shanghai 200438, P. R. China*

### Supplementary Materials

Supplementary Figures S1 to S7.

Supplementary Data: Cα-atom positions of the reference spike structure used in this study, which were adapted from the PDB structure 8DLW.pdb.

### Supplementary Figures

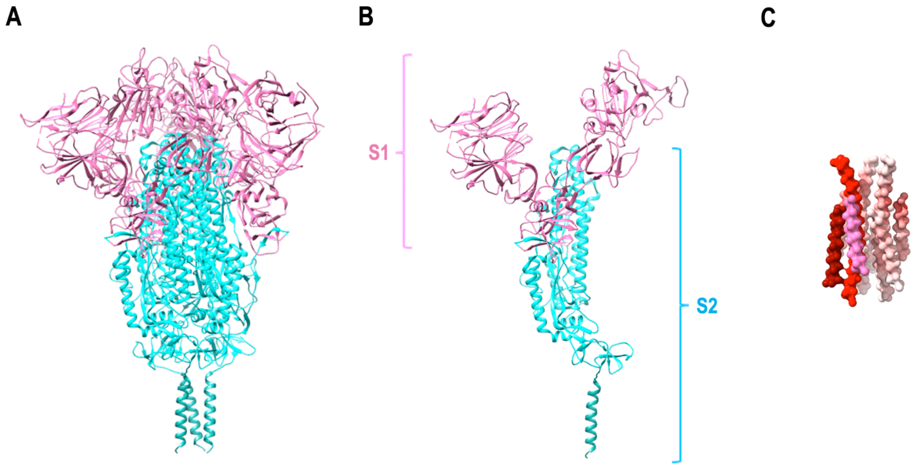


**Figure S1.** A crown-closed spike structure (PDB: 7YBJ). (A) Spike is composed of three protomers, each having S1 (pink) and S2 (cyan) subunits. The bottom triple helical segments sometimes were not solved in experimental spike structures, and hence were omitted in the main-text figures. (B) S1 and S2 subunits on each protomer. (C) The helical-segment bundles on S2 subunits used for the structural alignment in this study. For clarity, on one of these protomers, the three helical segments were colored dark-red, pink, and red, respectively.

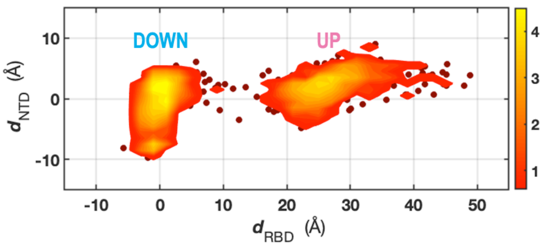


**Figure S2.** Motions of the NTD on the next protomer in relation to the motions of the RBD examined, e.g., $NTD_2$ and $RBD_1$.

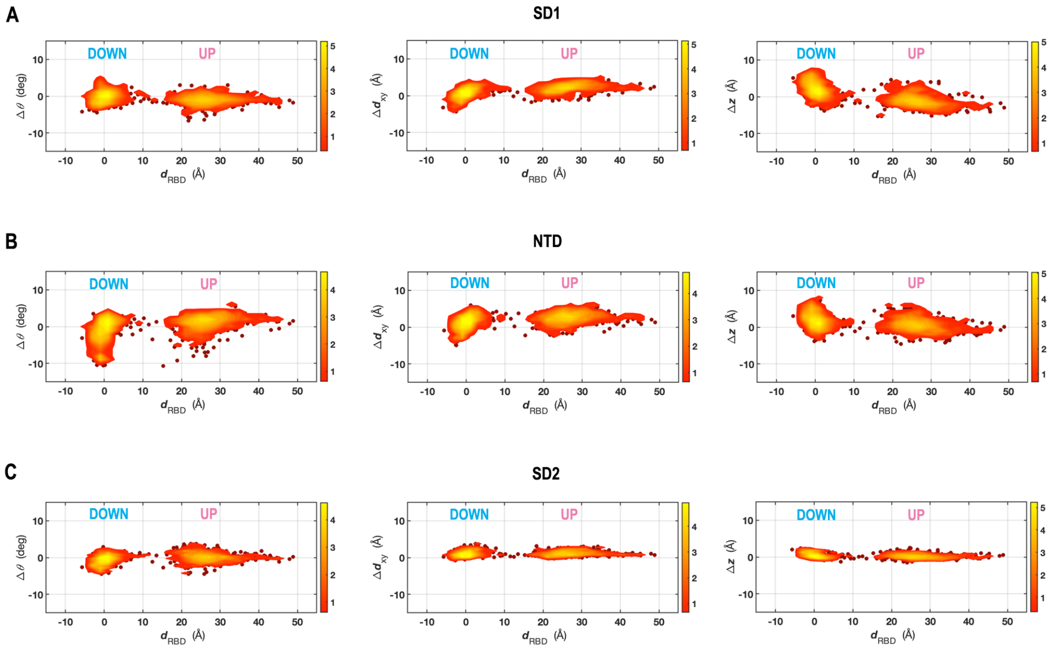


**Figure S3.** Rotation and elongation motions of SD1 (A), NTD (B), and SD2 (C), relative to the crown-closed reference spike.

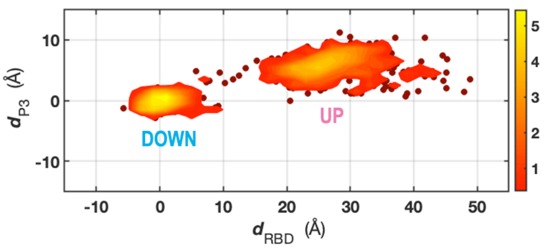


**Figure S4.** Motions of P3 along the RBD-rising pathway.

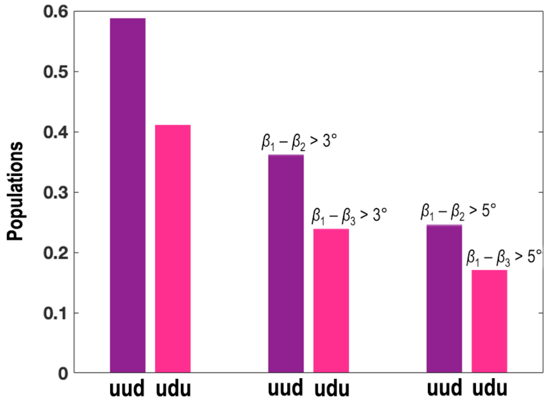


**Figure S5.** Comparisons of the partly open bound-spike populations (*uud* and *udu* cases). The first two columns compare populations of *uud* and *udu* cases of all bound-spikes. The next two columns compare only spikes having two UP-state RBDs' $\beta$-value difference > 3º, i.e., $\beta_1$ - $\beta_2$ > 3º in *uud* case and $\beta_1$ - $\beta_3$ > 3º in *udu* case. The last two columns compare spikes having two UP-state RBDs' $\beta$-value difference > 5º.

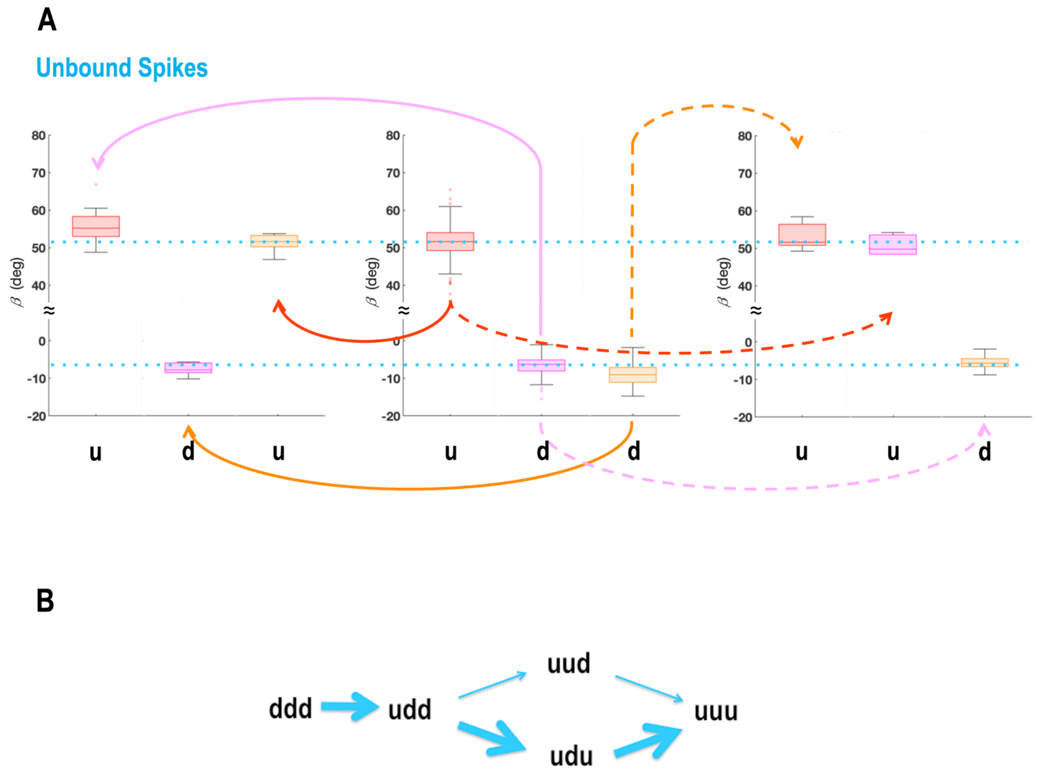


**Figure S6.** Proposals for the crown-opening pathways of unbound spikes. (A) Box plots of $\beta$ values of the trimeric RBDs in the partly closed (*udd* case) and partly open (*uud* and *udu* cases) unbound spikes. The proposed major pathways were sketched by solid lines ($RBD_2$ rising from *udd* to *udu* case, $\beta_2 > \beta_1$) and dashed lines ($RBD_3$ rising from *udd* to *uud* case, $\beta_3 > \beta_1$). An uncertain pathway formed by $RBD_3$ rising from *udd* to *udu* case ($\beta_3 < \beta_1$) was not sketched here. Data having $\beta > 80°$ in UP-RBD or $\beta > 10°$ in DOWN-RBD were excluded from averaging. Dotted-blue lines were drawn at the $\beta$-centers of $RBD_1$ and $RBD_2$ in *udd* case. (B) Proposed crown-opening pathway for the unbound spikes, where the majority of spikes went through the *udu* case based on their populations (see Fig. 6G). There are uncertainties in this proposal due to the few structures examined in the *udu* and *uud* cases of the unbound spikes.

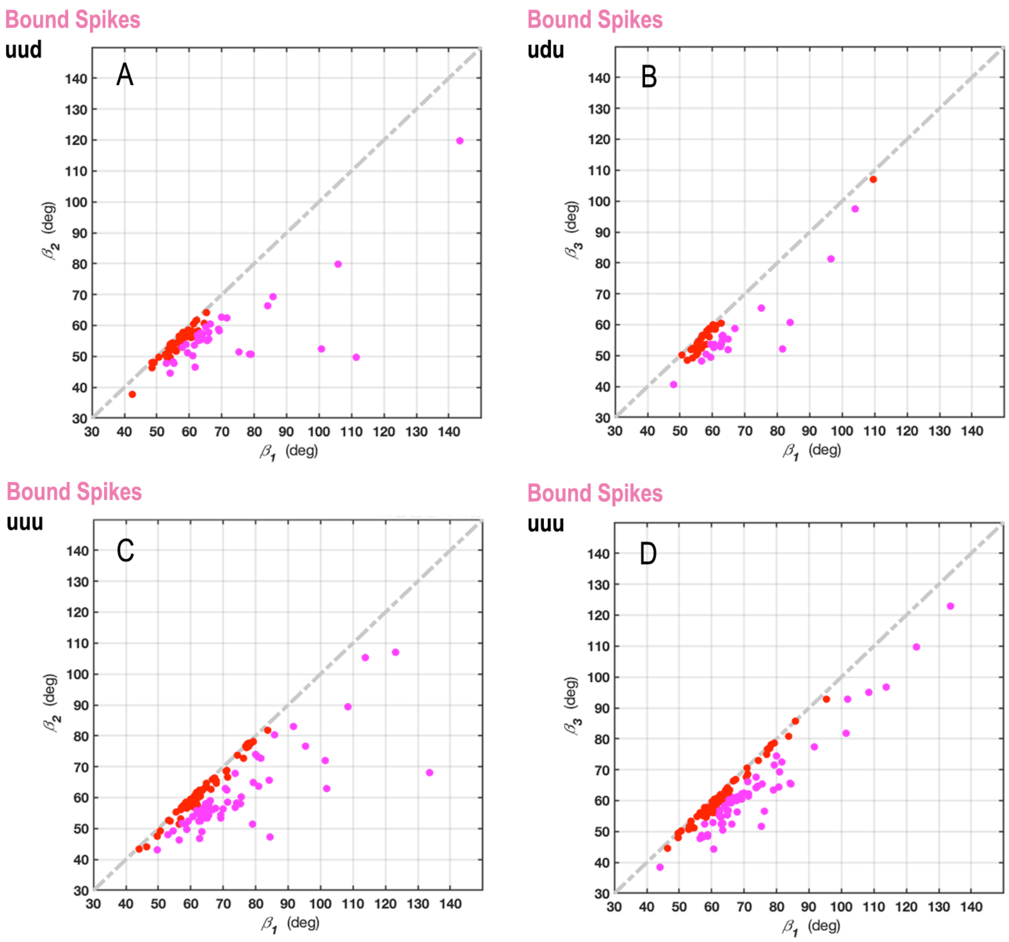


**Figure S7.** Pairwise $\beta$ correlations of RBDs in the partly open (A and B) and fully open (C and D) bound spikes. The data were colored magenta when the difference of the paired $\beta$ was larger than 5º.

### Supplementary Data

The file below contains the Cα-atom positions of the reference spike structure used in this study, adapted from the PDB structure (8DLW.pdb) that was published by Subramaniam group along with the publication:

Mannar, D., Saville, J. W., Sun, Z., Zhu, X., Marti, M. M., Srivastava, S. S., Berezuk, A. M., Zhou, S., Tuttle, K. S., Sobolewski, M. D., Kim, A., Treat, B. R., Da Silva Castanha, P. M., Jacobs, J. L., Barratt-Boyes, S. M., Mellors, J. W., Dimitrov, D. S., Li, W. & Subramaniam, S. SARS-CoV-2 variants of concern: spike protein mutational analysis and epitope for broad neutralization. *Nat Commun* **13**, 4696, doi:10.1038/s41467-022-32262-8 (2022).

```
ATOM      1  CA  CYS A  15      26.467  51.569  11.560  1.00183.98           C
ATOM      2  CA  VAL A  16      26.691  54.835   9.569  1.00185.44           C
ATOM      3  CA  ASN A  17      23.866  56.361   7.506  1.00191.33           C
ATOM      4  CA  LEU A  18      25.085  58.067   4.321  1.00183.17           C
ATOM      5  CA  THR A  19      22.818  60.971   3.363  1.00184.63           C
ATOM      6  CA  THR A  20      24.778  62.937   0.711  1.00180.57           C
ATOM      7  CA  ARG A  21      23.088  61.138  -2.183  1.00175.97           C
ATOM      8  CA  THR A  22      21.077  62.491  -5.105  1.00172.75           C
ATOM      9  CA  GLN A  23      17.648  60.843  -5.427  1.00172.16           C
ATOM     10  CA  LEU A  24      16.917  59.939  -9.056  1.00167.43           C
ATOM     11  CA  PRO A  25      14.473  57.348 -10.486  1.00161.17           C
ATOM     12  CA  PRO A  26      15.677  53.872 -11.564  1.00150.36           C
ATOM     13  CA  ALA A  27      17.025  53.481 -15.094  1.00138.45           C
ATOM     14  CA  TYR A  28      16.063  50.700 -17.496  1.00131.36           C
ATOM     15  CA  THR A  29      17.871  48.851 -20.295  1.00121.19           C
ATOM     16  CA  ASN A  30      17.606  45.915 -22.723  1.00110.45           C
ATOM     17  CA  SER A  31      18.940  42.491 -21.679  1.00107.26           C
ATOM     18  CA  PHE A  32      19.630  41.426 -25.324  1.00105.78           C
ATOM     19  CA  THR A  33      20.884  37.809 -25.266  1.00102.31           C
ATOM     20  CA  ARG A  34      22.682  37.907 -21.900  1.00102.86           C
ATOM     21  CA  GLY A  35      21.979  35.761 -18.862  1.00 98.68           C
ATOM     22  CA  VAL A  36      22.454  32.249 -20.268  1.00 95.03           C
ATOM     23  CA  TYR A  37      24.225  29.823 -17.919  1.00 90.99           C
ATOM     24  CA  TYR A  38      25.116  26.135 -17.886  1.00 87.28           C
ATOM     25  CA  PRO A  39      21.968  24.566 -16.354  1.00 88.76           C
ATOM     26  CA  ASP A  40      23.727  21.426 -15.084  1.00 89.66           C
ATOM     27  CA  LYS A  41      27.108  19.771 -14.458  1.00 89.45           C
ATOM     28  CA  VAL A  42      27.158  17.337 -17.413  1.00 86.23           C
ATOM     29  CA  PHE A  43      29.402  17.732 -20.457  1.00 86.57           C
ATOM     30  CA  ARG A  44      27.763  17.475 -23.853  1.00 85.27           C
ATOM     31  CA  SER A  45      29.108  18.630 -27.180  1.00 86.42           C
ATOM     32  CA  SER A  46      28.017  19.357 -30.776  1.00 87.99           C
ATOM     33  CA  VAL A  47      24.372  19.096 -29.687  1.00 85.53           C
ATOM     34  CA  LEU A  48      21.256  21.162 -29.191  1.00 83.88           C
ATOM     35  CA  HIS A  49      19.776  20.618 -25.731  1.00 83.75           C
ATOM     36  CA  SER A  50      16.287  21.690 -24.660  1.00 84.83           C
ATOM     37  CA  THR A  51      16.059  22.586 -20.975  1.00 86.61           C
ATOM     38  CA  GLN A  52      13.278  23.956 -18.765  1.00 89.96           C
ATOM     39  CA  ASP A  53      14.789  26.110 -16.016  1.00 92.25           C
ATOM     40  CA  LEU A  54      14.805  29.621 -14.566  1.00 91.67           C
```

```
ATOM     41  CA  PHE A  55      16.422  31.894 -17.150  1.00 90.81           C
ATOM     42  CA  LEU A  56      16.580  35.587 -17.880  1.00 94.18           C
ATOM     43  CA  PRO A  57      14.183  36.146 -20.824  1.00 94.13           C
ATOM     44  CA  PHE A  58      15.752  37.494 -23.996  1.00 95.10           C
ATOM     45  CA  PHE A  59      15.327  41.250 -24.602  1.00101.22           C
ATOM     46  CA  SER A  60      13.850  41.718 -21.134  1.00108.56           C
ATOM     47  CA  ASN A  61      13.536  45.005 -19.260  1.00123.41           C
ATOM     48  CA  VAL A  62      16.507  45.177 -16.859  1.00121.93           C
ATOM     49  CA  THR A  63      16.555  47.676 -13.998  1.00126.42           C
ATOM     50  CA  TRP A  64      19.749  49.759 -14.071  1.00133.35           C
ATOM     51  CA  PHE A  65      21.325  51.394 -10.997  1.00132.96           C
ATOM     52  CA  HIS A  66      24.374  53.620 -10.622  1.00137.28           C
ATOM     53  CA  ARG A  78      26.664  57.732  -6.991  1.00156.52           C
ATOM     54  CA  PHE A  79      25.802  55.153  -4.294  1.00158.70           C
ATOM     55  CA  ASP A  80      22.505  53.635  -5.421  1.00147.74           C
ATOM     56  CA  ASN A  81      21.624  50.410  -3.556  1.00144.74           C
ATOM     57  CA  PRO A  82      18.200  49.893  -1.914  1.00133.36           C
ATOM     58  CA  VAL A  83      16.740  46.523  -1.034  1.00124.73           C
ATOM     59  CA  LEU A  84      15.221  44.790  -4.061  1.00119.23           C
ATOM     60  CA  PRO A  85      12.831  41.826  -4.328  1.00113.27           C
ATOM     61  CA  PHE A  86      14.185  38.352  -5.037  1.00110.36           C
ATOM     62  CA  ASN A  87      11.547  36.135  -6.629  1.00108.22           C
ATOM     63  CA  ASP A  88      12.515  32.955  -8.520  1.00102.98           C
ATOM     64  CA  GLY A  89      16.089  34.191  -8.882  1.00100.71           C
ATOM     65  CA  VAL A  90      17.997  37.200 -10.106  1.00101.34           C
ATOM     66  CA  TYR A  91      20.501  38.190 -12.766  1.00105.45           C
ATOM     67  CA  PHE A  92      23.126  40.693 -11.681  1.00115.01           C
ATOM     68  CA  ALA A  93      25.758  42.412 -13.785  1.00121.00           C
ATOM     69  CA  SER A  94      28.369  45.049 -13.032  1.00130.33           C
ATOM     70  CA  THR A  95      30.491  47.287 -15.220  1.00140.35           C
ATOM     71  CA  GLU A  96      33.556  48.546 -13.354  1.00145.05           C
ATOM     72  CA  LYS A  97      37.071  49.825 -13.850  1.00148.80           C
ATOM     73  CA  SER A  98      38.358  50.042 -10.250  1.00148.79           C
ATOM     74  CA  ASN A  99      36.848  47.145  -8.155  1.00146.34           C
ATOM     75  CA  ILE A 100      34.004  48.958  -6.380  1.00147.78           C
ATOM     76  CA  ILE A 101      31.661  45.941  -5.908  1.00145.33           C
ATOM     77  CA  ARG A 102      31.853  44.175  -2.568  1.00144.40           C
ATOM     78  CA  GLY A 103      29.016  41.707  -2.287  1.00132.69           C
ATOM     79  CA  TRP A 104      25.404  40.920  -1.528  1.00122.62           C
ATOM     80  CA  ILE A 105      22.966  40.387   1.349  1.00125.12           C
ATOM     81  CA  PHE A 106      20.115  37.914   0.878  1.00118.46           C
ATOM     82  CA  GLY A 107      17.276  37.325   3.290  1.00120.33           C
ATOM     83  CA  THR A 108      13.680  37.865   4.320  1.00123.56           C
ATOM     84  CA  THR A 109      13.814  40.698   6.874  1.00128.73           C
ATOM     85  CA  LEU A 110      17.586  41.329   7.309  1.00134.15           C
ATOM     86  CA  ASP A 111      17.257  42.346  10.988  1.00136.31           C
ATOM     87  CA  SER A 112      18.404  39.229  12.979  1.00134.42           C
ATOM     88  CA  LYS A 113      14.813  37.897  13.196  1.00132.98           C
ATOM     89  CA  THR A 114      15.396  35.481  10.300  1.00131.08           C
ATOM     90  CA  GLN A 115      18.313  33.641   8.741  1.00129.83           C
ATOM     91  CA  SER A 116      20.322  35.672   6.224  1.00128.72           C
ATOM     92  CA  LEU A 117      23.125  35.076   3.724  1.00122.79           C
ATOM     93  CA  LEU A 118      26.120  37.432   3.519  1.00127.38           C
ATOM     94  CA  ILE A 119      28.562  37.278   0.591  1.00126.06           C
ATOM     95  CA  VAL A 120      31.583  39.614   0.751  1.00132.98           C
ATOM     96  CA  ASN A 121      34.646  39.728  -1.518  1.00137.73           C
ATOM     97  CA  ASN A 122      37.258  41.487   0.580  1.00145.26           C
ATOM     98  CA  ALA A 123      40.844  42.473  -0.277  1.00146.65           C
ATOM     99  CA  THR A 124      42.278  39.006   0.432  1.00142.41           C
ATOM    100  CA  ASN A 125      39.510  36.432  -0.263  1.00140.63           C
```

| | | | | | | | | | | |
|---|---|---|---|---|---|---|---|---|---|---|
| ATOM | 101 | CA | VAL | A | 126 | 35.773 | 35.743 | -0.647 | 1.00131.58 | C |
| ATOM | 102 | CA | VAL | A | 127 | 33.830 | 34.989 | 2.540 | 1.00129.74 | C |
| ATOM | 103 | CA | ILE | A | 128 | 30.293 | 33.563 | 2.638 | 1.00124.53 | C |
| ATOM | 104 | CA | LYS | A | 129 | 28.515 | 33.389 | 5.981 | 1.00126.87 | C |
| ATOM | 105 | CA | VAL | A | 130 | 25.006 | 32.227 | 6.798 | 1.00128.96 | C |
| ATOM | 106 | CA | CYS | A | 131 | 25.127 | 33.531 | 10.333 | 1.00139.43 | C |
| ATOM | 107 | CA | GLU | A | 132 | 22.185 | 35.369 | 11.878 | 1.00141.18 | C |
| ATOM | 108 | CA | PHE | A | 133 | 23.214 | 38.981 | 11.293 | 1.00142.97 | C |
| ATOM | 109 | CA | GLN | A | 134 | 21.745 | 42.222 | 12.593 | 1.00145.79 | C |
| ATOM | 110 | CA | PHE | A | 135 | 22.300 | 44.332 | 9.507 | 1.00149.88 | C |
| ATOM | 111 | CA | CYS | A | 136 | 22.753 | 48.074 | 9.186 | 1.00170.47 | C |
| ATOM | 112 | CA | ASN | A | 137 | 20.141 | 50.166 | 7.429 | 1.00179.27 | C |
| ATOM | 113 | CA | ASP | A | 138 | 22.863 | 51.467 | 5.083 | 1.00179.56 | C |
| ATOM | 114 | CA | PRO | A | 139 | 25.562 | 48.755 | 4.859 | 1.00173.59 | C |
| ATOM | 115 | CA | PHE | A | 140 | 28.801 | 49.420 | 2.964 | 1.00174.04 | C |
| ATOM | 116 | CA | LEU | A | 141 | 32.573 | 48.999 | 2.954 | 1.00169.52 | C |
| ATOM | 117 | CA | GLY | A | 142 | 35.213 | 51.695 | 2.609 | 1.00173.30 | C |
| ATOM | 118 | CA | VAL | A | 143 | 38.543 | 52.040 | 0.839 | 1.00173.38 | C |
| ATOM | 119 | CA | CYS | A | 152 | 43.858 | 52.911 | -1.948 | 1.00182.01 | C |
| ATOM | 120 | CA | MET | A | 153 | 43.538 | 50.392 | 0.892 | 1.00178.37 | C |
| ATOM | 121 | CA | GLU | A | 154 | 40.534 | 48.825 | 2.588 | 1.00172.33 | C |
| ATOM | 122 | CA | SER | A | 155 | 39.676 | 50.158 | 6.042 | 1.00174.66 | C |
| ATOM | 123 | CA | GLU | A | 156 | 36.023 | 50.468 | 7.047 | 1.00173.28 | C |
| ATOM | 124 | CA | PHE | A | 157 | 33.715 | 47.469 | 7.440 | 1.00168.10 | C |
| ATOM | 125 | CA | ARG | A | 158 | 30.195 | 48.575 | 8.448 | 1.00170.19 | C |
| ATOM | 126 | CA | VAL | A | 159 | 27.892 | 45.751 | 7.348 | 1.00161.20 | C |
| ATOM | 127 | CA | TYR | A | 160 | 26.479 | 44.351 | 10.596 | 1.00154.14 | C |
| ATOM | 128 | CA | SER | A | 161 | 26.650 | 44.651 | 14.371 | 1.00153.49 | C |
| ATOM | 129 | CA | SER | A | 162 | 25.436 | 41.548 | 16.207 | 1.00151.04 | C |
| ATOM | 130 | CA | ALA | A | 163 | 26.271 | 38.149 | 14.725 | 1.00145.06 | C |
| ATOM | 131 | CA | ASN | A | 164 | 25.564 | 34.784 | 16.355 | 1.00140.74 | C |
| ATOM | 132 | CA | ASN | A | 165 | 23.885 | 31.434 | 15.530 | 1.00140.35 | C |
| ATOM | 133 | CA | CYS | A | 166 | 26.291 | 30.773 | 12.651 | 1.00140.26 | C |
| ATOM | 134 | CA | THR | A | 167 | 25.204 | 27.856 | 10.468 | 1.00131.35 | C |
| ATOM | 135 | CA | PHE | A | 168 | 27.478 | 27.940 | 7.409 | 1.00121.26 | C |
| ATOM | 136 | CA | GLU | A | 169 | 30.784 | 29.514 | 6.394 | 1.00120.13 | C |
| ATOM | 137 | CA | TYR | A | 170 | 32.808 | 29.258 | 3.187 | 1.00122.25 | C |
| ATOM | 138 | CA | VAL | A | 171 | 36.021 | 30.993 | 2.050 | 1.00124.86 | C |
| ATOM | 139 | CA | SER | A | 172 | 37.626 | 30.804 | -1.404 | 1.00127.70 | C |
| ATOM | 140 | CA | GLN | A | 173 | 39.562 | 32.459 | -4.238 | 1.00127.42 | C |
| ATOM | 141 | CA | PRO | A | 174 | 38.516 | 36.030 | -5.254 | 1.00128.33 | C |
| ATOM | 142 | CA | PHE | A | 175 | 35.216 | 36.533 | -7.198 | 1.00126.86 | C |
| ATOM | 143 | CA | ASN | A | 185 | 34.880 | 53.834 | -20.190 | 1.00151.37 | C |
| ATOM | 144 | CA | PHE | A | 186 | 34.784 | 50.650 | -18.123 | 1.00147.64 | C |
| ATOM | 145 | CA | LYS | A | 187 | 37.043 | 47.623 | -18.556 | 1.00141.06 | C |
| ATOM | 146 | CA | ASN | A | 188 | 35.565 | 44.755 | -16.506 | 1.00138.02 | C |
| ATOM | 147 | CA | LEU | A | 189 | 32.107 | 43.239 | -16.982 | 1.00127.37 | C |
| ATOM | 148 | CA | ARG | A | 190 | 30.990 | 40.704 | -14.380 | 1.00123.22 | C |
| ATOM | 149 | CA | GLU | A | 191 | 27.797 | 38.670 | -14.681 | 1.00109.52 | C |
| ATOM | 150 | CA | PHE | A | 192 | 26.215 | 36.576 | -11.927 | 1.00105.32 | C |
| ATOM | 151 | CA | VAL | A | 193 | 23.084 | 34.445 | -11.656 | 1.00 99.24 | C |
| ATOM | 152 | CA | PHE | A | 194 | 21.640 | 33.805 | -8.186 | 1.00101.74 | C |
| ATOM | 153 | CA | LYS | A | 195 | 19.133 | 30.968 | -7.736 | 1.00 98.96 | C |
| ATOM | 154 | CA | ASN | A | 196 | 17.566 | 29.507 | -4.573 | 1.00105.80 | C |
| ATOM | 155 | CA | ILE | A | 197 | 16.258 | 26.019 | -5.451 | 1.00105.38 | C |
| ATOM | 156 | CA | ASP | A | 198 | 15.801 | 22.842 | -3.331 | 1.00107.63 | C |
| ATOM | 157 | CA | GLY | A | 199 | 18.060 | 23.927 | -0.514 | 1.00105.76 | C |
| ATOM | 158 | CA | TYR | A | 200 | 20.904 | 25.140 | -2.729 | 1.00102.00 | C |
| ATOM | 159 | CA | PHE | A | 201 | 21.804 | 28.726 | -3.491 | 1.00103.72 | C |
| ATOM | 160 | CA | LYS | A | 202 | 23.437 | 28.509 | -6.899 | 1.00 98.16 | C |

| | | | | | | | | | | |
|---|---|---|---|---|---|---|---|---|---|---|
| ATOM | 161 | CA | ILE | A | 203 | 25.867 | 31.198 | -8.034 | 1.00 98.29 | C |
| ATOM | 162 | CA | TYR | A | 204 | 26.892 | 31.337 | -11.690 | 1.00 95.55 | C |
| ATOM | 163 | CA | SER | A | 205 | 29.400 | 33.837 | -13.013 | 1.00100.32 | C |
| ATOM | 164 | CA | LYS | A | 206 | 31.279 | 35.100 | -16.052 | 1.00109.32 | C |
| ATOM | 165 | CA | HIS | A | 207 | 34.064 | 37.690 | -16.298 | 1.00121.21 | C |
| ATOM | 166 | CA | THR | A | 208 | 34.679 | 39.448 | -19.621 | 1.00128.08 | C |
| ATOM | 167 | CA | PRO | A | 209 | 36.322 | 42.627 | -20.919 | 1.00132.86 | C |
| ATOM | 168 | CA | ILE | A | 210 | 34.004 | 45.510 | -21.788 | 1.00137.58 | C |
| ATOM | 169 | CA | ASN | A | 211 | 34.515 | 48.732 | -23.727 | 1.00140.59 | C |
| ATOM | 170 | CA | LEU | A | 212 | 31.289 | 50.737 | -23.518 | 1.00143.26 | C |
| ATOM | 171 | CA | VAL | A | 213 | 30.920 | 53.863 | -21.423 | 1.00145.35 | C |
| ATOM | 172 | CA | ARG | A | 214 | 27.206 | 53.669 | -20.642 | 1.00140.54 | C |
| ATOM | 173 | CA | ASP | A | 215 | 25.766 | 50.287 | -21.704 | 1.00134.33 | C |
| ATOM | 174 | CA | LEU | A | 216 | 25.632 | 46.426 | -21.672 | 1.00125.99 | C |
| ATOM | 175 | CA | PRO | A | 217 | 27.276 | 44.607 | -24.613 | 1.00120.29 | C |
| ATOM | 176 | CA | GLN | A | 218 | 25.448 | 42.798 | -27.377 | 1.00115.95 | C |
| ATOM | 177 | CA | GLY | A | 219 | 26.595 | 39.285 | -28.129 | 1.00107.65 | C |
| ATOM | 178 | CA | PHE | A | 220 | 26.436 | 35.867 | -26.502 | 1.00 98.54 | C |
| ATOM | 179 | CA | SER | A | 221 | 28.425 | 34.229 | -23.719 | 1.00100.00 | C |
| ATOM | 180 | CA | ALA | A | 222 | 27.540 | 31.595 | -21.146 | 1.00 97.92 | C |
| ATOM | 181 | CA | LEU | A | 223 | 27.978 | 31.942 | -17.394 | 1.00 98.30 | C |
| ATOM | 182 | CA | GLU | A | 224 | 29.794 | 29.058 | -15.803 | 1.00 96.37 | C |
| ATOM | 183 | CA | PRO | A | 225 | 28.638 | 27.930 | -12.340 | 1.00 94.04 | C |
| ATOM | 184 | CA | LEU | A | 226 | 30.750 | 29.360 | -9.561 | 1.00 99.76 | C |
| ATOM | 185 | CA | VAL | A | 227 | 29.467 | 27.704 | -6.372 | 1.00102.87 | C |
| ATOM | 186 | CA | ASP | A | 228 | 26.507 | 25.655 | -5.100 | 1.00104.52 | C |
| ATOM | 187 | CA | LEU | A | 229 | 25.839 | 26.545 | -1.456 | 1.00105.57 | C |
| ATOM | 188 | CA | PRO | A | 230 | 23.770 | 23.921 | 0.443 | 1.00101.67 | C |
| ATOM | 189 | CA | ILE | A | 231 | 21.828 | 26.509 | 2.450 | 1.00108.06 | C |
| ATOM | 190 | CA | GLY | A | 232 | 18.067 | 26.230 | 2.978 | 1.00110.40 | C |
| ATOM | 191 | CA | ILE | A | 233 | 17.264 | 29.948 | 3.270 | 1.00114.53 | C |
| ATOM | 192 | CA | ASN | A | 234 | 13.950 | 31.657 | 2.572 | 1.00117.46 | C |
| ATOM | 193 | CA | ILE | A | 235 | 15.149 | 34.466 | 0.281 | 1.00114.69 | C |
| ATOM | 194 | CA | THR | A | 236 | 12.728 | 37.176 | -0.836 | 1.00115.71 | C |
| ATOM | 195 | CA | ARG | A | 237 | 14.873 | 40.336 | -0.612 | 1.00117.69 | C |
| ATOM | 196 | CA | PHE | A | 238 | 18.470 | 41.284 | -1.368 | 1.00119.30 | C |
| ATOM | 197 | CA | GLN | A | 239 | 20.833 | 44.259 | -1.172 | 1.00129.92 | C |
| ATOM | 198 | CA | THR | A | 240 | 24.103 | 45.120 | -2.939 | 1.00132.18 | C |
| ATOM | 199 | CA | LEU | A | 241 | 27.314 | 46.002 | -1.039 | 1.00140.93 | C |
| ATOM | 200 | CA | LEU | A | 242 | 29.485 | 48.705 | -2.653 | 1.00149.38 | C |
| ATOM | 201 | CA | ALA | A | 243 | 32.802 | 50.343 | -1.762 | 1.00153.77 | C |
| ATOM | 202 | CA | TRP | A | 258 | 33.416 | 59.944 | 1.378 | 1.00173.58 | C |
| ATOM | 203 | CA | THR | A | 259 | 33.274 | 59.925 | -2.428 | 1.00170.74 | C |
| ATOM | 204 | CA | ALA | A | 260 | 31.722 | 57.865 | -5.212 | 1.00165.09 | C |
| ATOM | 205 | CA | GLY | A | 261 | 33.587 | 55.675 | -7.680 | 1.00158.37 | C |
| ATOM | 206 | CA | ALA | A | 262 | 32.971 | 54.801 | -11.325 | 1.00153.24 | C |
| ATOM | 207 | CA | ALA | A | 263 | 30.706 | 51.727 | -11.456 | 1.00147.88 | C |
| ATOM | 208 | CA | ALA | A | 264 | 27.251 | 50.670 | -12.609 | 1.00139.65 | C |
| ATOM | 209 | CA | TYR | A | 265 | 25.107 | 47.589 | -12.121 | 1.00132.50 | C |
| ATOM | 210 | CA | TYR | A | 266 | 22.068 | 45.947 | -13.689 | 1.00125.16 | C |
| ATOM | 211 | CA | VAL | A | 267 | 19.433 | 43.709 | -12.063 | 1.00115.30 | C |
| ATOM | 212 | CA | GLY | A | 268 | 17.111 | 41.401 | -13.998 | 1.00106.52 | C |
| ATOM | 213 | CA | TYR | A | 269 | 14.742 | 38.632 | -12.942 | 1.00101.23 | C |
| ATOM | 214 | CA | LEU | A | 270 | 14.584 | 34.966 | -13.893 | 1.00 94.66 | C |
| ATOM | 215 | CA | GLN | A | 271 | 11.433 | 33.262 | -15.160 | 1.00 91.25 | C |
| ATOM | 216 | CA | PRO | A | 272 | 10.522 | 29.625 | -15.902 | 1.00 90.26 | C |
| ATOM | 217 | CA | ARG | A | 273 | 11.538 | 29.257 | -19.552 | 1.00 89.22 | C |
| ATOM | 218 | CA | THR | A | 274 | 12.620 | 26.593 | -22.005 | 1.00 87.08 | C |
| ATOM | 219 | CA | PHE | A | 275 | 15.932 | 27.297 | -23.739 | 1.00 86.46 | C |
| ATOM | 220 | CA | LEU | A | 276 | 17.594 | 25.514 | -26.642 | 1.00 84.00 | C |

| | | | | | | | | | | | |
|---|---|---|---|---|---|---|---|---|---|---|---|
| ATOM | 221 | CA | LEU | A | 277 | 21.343 | 25.528 | -25.947 | 1.00 | 83.79 | C |
| ATOM | 222 | CA | LYS | A | 278 | 23.940 | 24.976 | -28.677 | 1.00 | 86.04 | C |
| ATOM | 223 | CA | TYR | A | 279 | 27.033 | 23.042 | -27.580 | 1.00 | 85.58 | C |
| ATOM | 224 | CA | ASN | A | 280 | 30.076 | 23.184 | -29.877 | 1.00 | 87.98 | C |
| ATOM | 225 | CA | GLU | A | 281 | 32.813 | 20.567 | -30.409 | 1.00 | 89.21 | C |
| ATOM | 226 | CA | ASN | A | 282 | 34.497 | 21.524 | -27.112 | 1.00 | 91.78 | C |
| ATOM | 227 | CA | GLY | A | 283 | 31.222 | 21.516 | -25.176 | 1.00 | 88.06 | C |
| ATOM | 228 | CA | THR | A | 284 | 31.036 | 25.301 | -24.890 | 1.00 | 87.29 | C |
| ATOM | 229 | CA | ILE | A | 285 | 27.627 | 26.940 | -25.202 | 1.00 | 87.42 | C |
| ATOM | 230 | CA | THR | A | 286 | 27.914 | 29.294 | -28.167 | 1.00 | 89.12 | C |
| ATOM | 231 | CA | ASP | A | 287 | 24.273 | 30.152 | -28.863 | 1.00 | 90.08 | C |
| ATOM | 232 | CA | ALA | A | 288 | 20.789 | 29.712 | -27.434 | 1.00 | 86.79 | C |
| ATOM | 233 | CA | VAL | A | 289 | 17.140 | 30.137 | -28.348 | 1.00 | 85.31 | C |
| ATOM | 234 | CA | ASP | A | 290 | 14.603 | 31.474 | -25.844 | 1.00 | 86.75 | C |
| ATOM | 235 | CA | CYS | A | 291 | 11.634 | 29.277 | -26.858 | 1.00 | 86.29 | C |
| ATOM | 236 | CA | ALA | A | 292 | 9.036 | 31.952 | -25.942 | 1.00 | 84.44 | C |
| ATOM | 237 | CA | LEU | A | 293 | 10.700 | 35.159 | -27.201 | 1.00 | 84.62 | C |
| ATOM | 238 | CA | ASP | A | 294 | 8.943 | 35.257 | -30.577 | 1.00 | 82.21 | C |
| ATOM | 239 | CA | PRO | A | 295 | 7.264 | 32.868 | -33.091 | 1.00 | 77.73 | C |
| ATOM | 240 | CA | LEU | A | 296 | 10.547 | 32.138 | -34.911 | 1.00 | 78.03 | C |
| ATOM | 241 | CA | SER | A | 297 | 12.304 | 31.175 | -31.662 | 1.00 | 81.08 | C |
| ATOM | 242 | CA | GLU | A | 298 | 9.419 | 28.832 | -30.841 | 1.00 | 80.50 | C |
| ATOM | 243 | CA | THR | A | 299 | 9.733 | 27.303 | -34.316 | 1.00 | 79.96 | C |
| ATOM | 244 | CA | LYS | A | 300 | 13.482 | 26.818 | -33.818 | 1.00 | 78.61 | C |
| ATOM | 245 | CA | CYS | A | 301 | 12.844 | 25.036 | -30.513 | 1.00 | 82.77 | C |
| ATOM | 246 | CA | THR | A | 302 | 10.060 | 22.890 | -32.009 | 1.00 | 81.42 | C |
| ATOM | 247 | CA | LEU | A | 303 | 12.262 | 21.645 | -34.854 | 1.00 | 80.37 | C |
| ATOM | 248 | CA | LYS | A | 304 | 15.323 | 21.341 | -32.531 | 1.00 | 82.61 | C |
| ATOM | 249 | CA | SER | A | 305 | 17.458 | 23.367 | -34.923 | 1.00 | 81.84 | C |
| ATOM | 250 | CA | PHE | A | 306 | 18.876 | 26.863 | -35.265 | 1.00 | 84.38 | C |
| ATOM | 251 | CA | THR | A | 307 | 18.137 | 26.774 | -39.006 | 1.00 | 84.97 | C |
| ATOM | 252 | CA | VAL | A | 308 | 14.546 | 26.656 | -40.259 | 1.00 | 83.52 | C |
| ATOM | 253 | CA | GLU | A | 309 | 13.686 | 25.791 | -43.848 | 1.00 | 82.29 | C |
| ATOM | 254 | CA | LYS | A | 310 | 10.931 | 27.499 | -45.832 | 1.00 | 77.66 | C |
| ATOM | 255 | CA | GLY | A | 311 | 7.538 | 26.085 | -44.945 | 1.00 | 73.31 | C |
| ATOM | 256 | CA | ILE | A | 312 | 4.727 | 26.024 | -42.419 | 1.00 | 71.55 | C |
| ATOM | 257 | CA | TYR | A | 313 | 5.287 | 24.357 | -39.058 | 1.00 | 73.61 | C |
| ATOM | 258 | CA | GLN | A | 314 | 2.783 | 23.735 | -36.272 | 1.00 | 76.86 | C |
| ATOM | 259 | CA | THR | A | 315 | 4.613 | 24.968 | -33.197 | 1.00 | 78.30 | C |
| ATOM | 260 | CA | SER | A | 316 | 2.176 | 25.669 | -30.357 | 1.00 | 79.14 | C |
| ATOM | 261 | CA | ASN | A | 317 | -1.404 | 25.915 | -29.132 | 1.00 | 79.06 | C |
| ATOM | 262 | CA | PHE | A | 318 | -3.345 | 29.118 | -28.499 | 1.00 | 84.71 | C |
| ATOM | 263 | CA | ARG | A | 319 | -5.780 | 29.222 | -25.586 | 1.00 | 87.71 | C |
| ATOM | 264 | CA | VAL | A | 320 | -7.667 | 32.117 | -24.010 | 1.00 | 87.84 | C |
| ATOM | 265 | CA | GLN | A | 321 | -6.952 | 32.202 | -20.327 | 1.00 | 90.25 | C |
| ATOM | 266 | CA | PRO | A | 322 | -9.721 | 32.727 | -17.762 | 1.00 | 90.18 | C |
| ATOM | 267 | CA | THR | A | 323 | -9.753 | 36.007 | -15.869 | 1.00 | 92.54 | C |
| ATOM | 268 | CA | GLU | A | 324 | -12.435 | 35.385 | -13.221 | 1.00 | 91.55 | C |
| ATOM | 269 | CA | SER | A | 325 | -13.548 | 32.620 | -10.873 | 1.00 | 87.34 | C |
| ATOM | 270 | CA | ILE | A | 326 | -17.322 | 32.111 | -10.848 | 1.00 | 84.69 | C |
| ATOM | 271 | CA | VAL | A | 327 | -18.561 | 30.153 | -7.851 | 1.00 | 83.75 | C |
| ATOM | 272 | CA | ARG | A | 328 | -22.247 | 29.244 | -7.504 | 1.00 | 83.11 | C |
| ATOM | 273 | CA | PHE | A | 329 | -23.753 | 27.180 | -4.685 | 1.00 | 84.56 | C |
| ATOM | 274 | CA | PRO | A | 330 | -27.348 | 26.892 | -3.341 | 1.00 | 85.57 | C |
| ATOM | 275 | CA | ASN | A | 331 | -28.353 | 29.811 | -1.106 | 1.00 | 90.57 | C |
| ATOM | 276 | CA | ILE | A | 332 | -28.850 | 27.729 | 2.032 | 1.00 | 84.78 | C |
| ATOM | 277 | CA | THR | A | 333 | -27.664 | 28.787 | 5.479 | 1.00 | 85.67 | C |
| ATOM | 278 | CA | ASN | A | 334 | -28.586 | 25.956 | 7.872 | 1.00 | 81.86 | C |
| ATOM | 279 | CA | LEU | A | 335 | -25.635 | 24.018 | 9.255 | 1.00 | 80.24 | C |
| ATOM | 280 | CA | CYS | A | 336 | -25.359 | 20.252 | 9.009 | 1.00 | 81.05 | C |

| | | | | | | | | | | | |
|---|---|---|---|---|---|---|---|---|---|---|---|
| ATOM | 281 | CA | PRO | A | 337 | -26.066 | 18.392 | 12.289 | 1.00 | 78.54 | C |
| ATOM | 282 | CA | PHE | A | 338 | -22.546 | 17.002 | 12.663 | 1.00 | 77.49 | C |
| ATOM | 283 | CA | GLY | A | 339 | -22.287 | 18.259 | 16.243 | 1.00 | 82.20 | C |
| ATOM | 284 | CA | GLU | A | 340 | -25.337 | 16.181 | 17.236 | 1.00 | 81.02 | C |
| ATOM | 285 | CA | VAL | A | 341 | -23.589 | 13.045 | 15.964 | 1.00 | 73.01 | C |
| ATOM | 286 | CA | PHE | A | 342 | -20.298 | 13.711 | 17.755 | 1.00 | 70.54 | C |
| ATOM | 287 | CA | ASN | A | 343 | -21.672 | 14.887 | 21.127 | 1.00 | 82.60 | C |
| ATOM | 288 | CA | ALA | A | 344 | -24.529 | 12.396 | 21.609 | 1.00 | 77.54 | C |
| ATOM | 289 | CA | THR | A | 345 | -24.687 | 11.115 | 25.188 | 1.00 | 79.08 | C |
| ATOM | 290 | CA | ARG | A | 346 | -25.063 | 7.460 | 24.168 | 1.00 | 77.31 | C |
| ATOM | 291 | CA | PHE | A | 347 | -23.283 | 5.822 | 21.253 | 1.00 | 65.64 | C |
| ATOM | 292 | CA | ALA | A | 348 | -24.440 | 2.650 | 19.567 | 1.00 | 63.03 | C |
| ATOM | 293 | CA | SER | A | 349 | -22.692 | -0.690 | 19.729 | 1.00 | 64.44 | C |
| ATOM | 294 | CA | VAL | A | 350 | -20.698 | -1.506 | 16.584 | 1.00 | 61.65 | C |
| ATOM | 295 | CA | TYR | A | 351 | -22.940 | -4.513 | 15.810 | 1.00 | 63.69 | C |
| ATOM | 296 | CA | ALA | A | 352 | -25.967 | -2.152 | 15.795 | 1.00 | 62.54 | C |
| ATOM | 297 | CA | TRP | A | 353 | -24.299 | 1.076 | 14.637 | 1.00 | 62.17 | C |
| ATOM | 298 | CA | ASN | A | 354 | -26.290 | 4.283 | 14.210 | 1.00 | 63.67 | C |
| ATOM | 299 | CA | ARG | A | 355 | -26.695 | 6.072 | 10.891 | 1.00 | 65.35 | C |
| ATOM | 300 | CA | LYS | A | 356 | -27.670 | 9.767 | 10.758 | 1.00 | 71.66 | C |
| ATOM | 301 | CA | ARG | A | 357 | -28.470 | 10.698 | 7.165 | 1.00 | 79.71 | C |
| ATOM | 302 | CA | ILE | A | 358 | -27.316 | 14.297 | 6.567 | 1.00 | 75.80 | C |
| ATOM | 303 | CA | SER | A | 359 | -28.880 | 16.325 | 3.741 | 1.00 | 79.36 | C |
| ATOM | 304 | CA | ASN | A | 360 | -30.002 | 19.841 | 2.690 | 1.00 | 81.39 | C |
| ATOM | 305 | CA | CYS | A | 361 | -27.470 | 21.789 | 4.758 | 1.00 | 80.26 | C |
| ATOM | 306 | CA | VAL | A | 362 | -24.098 | 23.544 | 4.794 | 1.00 | 78.23 | C |
| ATOM | 307 | CA | ALA | A | 363 | -21.272 | 21.283 | 5.962 | 1.00 | 75.30 | C |
| ATOM | 308 | CA | ASP | A | 364 | -18.322 | 23.075 | 7.542 | 1.00 | 73.54 | C |
| ATOM | 309 | CA | TYR | A | 365 | -15.625 | 20.405 | 7.687 | 1.00 | 70.88 | C |
| ATOM | 310 | CA | SER | A | 366 | -12.841 | 22.686 | 8.904 | 1.00 | 71.44 | C |
| ATOM | 311 | CA | VAL | A | 367 | -14.214 | 22.505 | 12.447 | 1.00 | 69.58 | C |
| ATOM | 312 | CA | LEU | A | 368 | -13.927 | 18.709 | 12.525 | 1.00 | 67.20 | C |
| ATOM | 313 | CA | TYR | A | 369 | -10.528 | 18.492 | 10.824 | 1.00 | 66.72 | C |
| ATOM | 314 | CA | ASN | A | 370 | -8.798 | 21.243 | 12.789 | 1.00 | 68.44 | C |
| ATOM | 315 | CA | SER | A | 371 | -10.133 | 19.966 | 16.113 | 1.00 | 69.26 | C |
| ATOM | 316 | CA | ALA | A | 372 | -7.620 | 18.708 | 18.659 | 1.00 | 67.89 | C |
| ATOM | 317 | CA | SER | A | 373 | -10.069 | 16.271 | 20.289 | 1.00 | 69.80 | C |
| ATOM | 318 | CA | PHE | A | 374 | -9.693 | 13.658 | 17.504 | 1.00 | 66.31 | C |
| ATOM | 319 | CA | SER | A | 375 | -6.846 | 11.146 | 17.337 | 1.00 | 63.79 | C |
| ATOM | 320 | CA | THR | A | 376 | -7.765 | 9.340 | 14.114 | 1.00 | 62.82 | C |
| ATOM | 321 | CA | PHE | A | 377 | -9.005 | 11.371 | 11.145 | 1.00 | 60.67 | C |
| ATOM | 322 | CA | LYS | A | 378 | -8.386 | 9.360 | 7.980 | 1.00 | 64.14 | C |
| ATOM | 323 | CA | CYS | A | 379 | -10.068 | 10.228 | 4.686 | 1.00 | 63.33 | C |
| ATOM | 324 | CA | TYR | A | 380 | -10.453 | 7.828 | 1.754 | 1.00 | 63.46 | C |
| ATOM | 325 | CA | GLY | A | 381 | -11.483 | 8.981 | -1.706 | 1.00 | 62.44 | C |
| ATOM | 326 | CA | VAL | A | 382 | -11.190 | 12.694 | -0.834 | 1.00 | 63.98 | C |
| ATOM | 327 | CA | SER | A | 383 | -8.610 | 14.749 | 0.896 | 1.00 | 66.50 | C |
| ATOM | 328 | CA | PRO | A | 384 | -9.469 | 16.288 | 4.298 | 1.00 | 66.33 | C |
| ATOM | 329 | CA | THR | A | 385 | -8.317 | 19.796 | 3.303 | 1.00 | 66.45 | C |
| ATOM | 330 | CA | LYS | A | 386 | -10.476 | 20.003 | 0.161 | 1.00 | 67.25 | C |
| ATOM | 331 | CA | LEU | A | 387 | -13.732 | 18.939 | 1.859 | 1.00 | 68.15 | C |
| ATOM | 332 | CA | ASN | A | 388 | -15.056 | 22.487 | 2.181 | 1.00 | 71.17 | C |
| ATOM | 333 | CA | ASP | A | 389 | -14.792 | 22.998 | -1.597 | 1.00 | 74.11 | C |
| ATOM | 334 | CA | LEU | A | 390 | -16.904 | 19.925 | -2.430 | 1.00 | 70.94 | C |
| ATOM | 335 | CA | CYS | A | 391 | -20.606 | 19.174 | -2.700 | 1.00 | 76.80 | C |
| ATOM | 336 | CA | PHE | A | 392 | -22.216 | 15.823 | -2.070 | 1.00 | 75.24 | C |
| ATOM | 337 | CA | THR | A | 393 | -25.740 | 14.592 | -2.652 | 1.00 | 81.75 | C |
| ATOM | 338 | CA | ASN | A | 394 | -25.892 | 12.902 | 0.762 | 1.00 | 80.70 | C |
| ATOM | 339 | CA | VAL | A | 395 | -23.699 | 12.278 | 3.826 | 1.00 | 73.31 | C |
| ATOM | 340 | CA | TYR | A | 396 | -24.124 | 9.178 | 5.999 | 1.00 | 71.54 | C |

| | | | | | | | | | | | |
|---|---|---|---|---|---|---|---|---|---|---|---|
| ATOM | 341 | CA | ALA | A | 397 | -22.880 | 9.898 | 9.555 | 1.00 | 66.54 | C |
| ATOM | 342 | CA | ASP | A | 398 | -22.281 | 6.436 | 11.059 | 1.00 | 64.06 | C |
| ATOM | 343 | CA | SER | A | 399 | -21.440 | 6.170 | 14.752 | 1.00 | 62.66 | C |
| ATOM | 344 | CA | PHE | A | 400 | -20.313 | 3.403 | 17.124 | 1.00 | 60.09 | C |
| ATOM | 345 | CA | VAL | A | 401 | -17.904 | 2.471 | 19.952 | 1.00 | 62.29 | C |
| ATOM | 346 | CA | ILE | A | 402 | -15.021 | 0.050 | 19.327 | 1.00 | 64.88 | C |
| ATOM | 347 | CA | ARG | A | 403 | -11.833 | -1.108 | 21.022 | 1.00 | 71.51 | C |
| ATOM | 348 | CA | GLY | A | 404 | -8.673 | 0.895 | 20.273 | 1.00 | 70.86 | C |
| ATOM | 349 | CA | ASP | A | 405 | -6.860 | -1.978 | 18.530 | 1.00 | 75.73 | C |
| ATOM | 350 | CA | GLU | A | 406 | -9.848 | -2.296 | 16.212 | 1.00 | 71.66 | C |
| ATOM | 351 | CA | VAL | A | 407 | -9.691 | 1.362 | 15.075 | 1.00 | 66.95 | C |
| ATOM | 352 | CA | ARG | A | 408 | -7.432 | 0.271 | 12.197 | 1.00 | 69.62 | C |
| ATOM | 353 | CA | GLN | A | 409 | -10.169 | -2.096 | 10.988 | 1.00 | 67.27 | C |
| ATOM | 354 | CA | ILE | A | 410 | -12.485 | 0.814 | 10.080 | 1.00 | 63.32 | C |
| ATOM | 355 | CA | ALA | A | 411 | -10.915 | 1.158 | 6.629 | 1.00 | 65.46 | C |
| ATOM | 356 | CA | PRO | A | 412 | -11.454 | -0.204 | 3.104 | 1.00 | 67.50 | C |
| ATOM | 357 | CA | GLY | A | 413 | -9.904 | -3.585 | 2.465 | 1.00 | 71.76 | C |
| ATOM | 358 | CA | GLN | A | 414 | -9.579 | -4.702 | 6.094 | 1.00 | 70.52 | C |
| ATOM | 359 | CA | THR | A | 415 | -10.207 | -8.065 | 7.740 | 1.00 | 72.86 | C |
| ATOM | 360 | CA | GLY | A | 416 | -11.009 | -9.143 | 11.272 | 1.00 | 71.96 | C |
| ATOM | 361 | CA | LYS | A | 417 | -14.077 | -9.216 | 13.512 | 1.00 | 68.92 | C |
| ATOM | 362 | CA | ILE | A | 418 | -15.336 | -5.635 | 13.259 | 1.00 | 64.80 | C |
| ATOM | 363 | CA | ALA | A | 419 | -14.710 | -5.478 | 9.520 | 1.00 | 65.53 | C |
| ATOM | 364 | CA | ASP | A | 420 | -15.944 | -8.925 | 8.523 | 1.00 | 70.38 | C |
| ATOM | 365 | CA | TYR | A | 421 | -18.818 | -9.009 | 11.093 | 1.00 | 66.10 | C |
| ATOM | 366 | CA | ASN | A | 422 | -19.698 | -5.584 | 12.530 | 1.00 | 63.95 | C |
| ATOM | 367 | CA | TYR | A | 423 | -18.949 | -2.578 | 10.239 | 1.00 | 62.02 | C |
| ATOM | 368 | CA | LYS | A | 424 | -17.710 | -3.491 | 6.748 | 1.00 | 64.51 | C |
| ATOM | 369 | CA | LEU | A | 425 | -16.625 | -0.602 | 4.484 | 1.00 | 65.40 | C |
| ATOM | 370 | CA | PRO | A | 426 | -16.654 | -0.869 | 0.674 | 1.00 | 69.60 | C |
| ATOM | 371 | CA | ASP | A | 427 | -13.392 | -0.662 | -1.236 | 1.00 | 77.11 | C |
| ATOM | 372 | CA | ASP | A | 428 | -14.556 | 2.387 | -3.238 | 1.00 | 76.97 | C |
| ATOM | 373 | CA | PHE | A | 429 | -15.539 | 4.206 | -0.031 | 1.00 | 67.63 | C |
| ATOM | 374 | CA | THR | A | 430 | -15.502 | 7.998 | -0.127 | 1.00 | 67.15 | C |
| ATOM | 375 | CA | GLY | A | 431 | -15.544 | 9.543 | 3.313 | 1.00 | 64.84 | C |
| ATOM | 376 | CA | CYS | A | 432 | -13.646 | 10.132 | 6.520 | 1.00 | 63.95 | C |
| ATOM | 377 | CA | VAL | A | 433 | -13.123 | 7.856 | 9.537 | 1.00 | 59.33 | C |
| ATOM | 378 | CA | ILE | A | 434 | -12.933 | 10.211 | 12.523 | 1.00 | 60.03 | C |
| ATOM | 379 | CA | ALA | A | 435 | -12.204 | 8.507 | 15.844 | 1.00 | 61.63 | C |
| ATOM | 380 | CA | TRP | A | 436 | -11.248 | 9.543 | 19.362 | 1.00 | 63.00 | C |
| ATOM | 381 | CA | ASN | A | 437 | -10.409 | 7.991 | 22.719 | 1.00 | 66.81 | C |
| ATOM | 382 | CA | SER | A | 438 | -13.479 | 8.086 | 24.971 | 1.00 | 71.08 | C |
| ATOM | 383 | CA | ASN | A | 439 | -12.131 | 6.366 | 28.117 | 1.00 | 75.60 | C |
| ATOM | 384 | CA | ASN | A | 440 | -13.349 | 9.276 | 30.256 | 1.00 | 78.39 | C |
| ATOM | 385 | CA | LEU | A | 441 | -16.909 | 8.783 | 28.933 | 1.00 | 75.29 | C |
| ATOM | 386 | CA | ASP | A | 442 | -17.379 | 5.070 | 28.231 | 1.00 | 74.58 | C |
| ATOM | 387 | CA | SER | A | 443 | -15.378 | 3.354 | 30.998 | 1.00 | 77.85 | C |
| ATOM | 388 | CA | LYS | A | 444 | -16.705 | 2.130 | 34.348 | 1.00 | 81.78 | C |
| ATOM | 389 | CA | VAL | A | 445 | -14.790 | 0.578 | 37.251 | 1.00 | 85.28 | C |
| ATOM | 390 | CA | GLY | A | 446 | -16.303 | -2.874 | 36.998 | 1.00 | 85.76 | C |
| ATOM | 391 | CA | GLY | A | 447 | -16.431 | -2.659 | 33.215 | 1.00 | 82.46 | C |
| ATOM | 392 | CA | ASN | A | 448 | -19.071 | -1.016 | 31.034 | 1.00 | 79.00 | C |
| ATOM | 393 | CA | TYR | A | 449 | -20.850 | -3.984 | 29.455 | 1.00 | 80.62 | C |
| ATOM | 394 | CA | ASN | A | 450 | -23.382 | -1.975 | 27.422 | 1.00 | 76.16 | C |
| ATOM | 395 | CA | TYR | A | 451 | -21.172 | -1.756 | 24.321 | 1.00 | 69.64 | C |
| ATOM | 396 | CA | ARG | A | 452 | -21.249 | -5.048 | 22.412 | 1.00 | 69.13 | C |
| ATOM | 397 | CA | TYR | A | 453 | -19.844 | -6.440 | 19.170 | 1.00 | 66.50 | C |
| ATOM | 398 | CA | ARG | A | 454 | -20.699 | -9.394 | 16.964 | 1.00 | 65.58 | C |
| ATOM | 399 | CA | LEU | A | 455 | -18.075 | -12.105 | 17.287 | 1.00 | 65.46 | C |
| ATOM | 400 | CA | PHE | A | 456 | -19.578 | -14.854 | 15.076 | 1.00 | 67.34 | C |

| | | | | | | | | | | | |
|---|---|---|---|---|---|---|---|---|---|---|---|
| ATOM | 401 | CA | ARG | A | 457 | -21.488 | -15.047 | 11.776 | 1.00 | 68.20 | C |
| ATOM | 402 | CA | LYS | A | 458 | -21.978 | -17.567 | 8.962 | 1.00 | 71.37 | C |
| ATOM | 403 | CA | SER | A | 459 | -20.600 | -15.184 | 6.320 | 1.00 | 69.62 | C |
| ATOM | 404 | CA | ASN | A | 460 | -19.026 | -11.766 | 6.053 | 1.00 | 69.47 | C |
| ATOM | 405 | CA | LEU | A | 461 | -21.155 | -8.622 | 6.168 | 1.00 | 67.26 | C |
| ATOM | 406 | CA | LYS | A | 462 | -21.600 | -6.917 | 2.790 | 1.00 | 68.79 | C |
| ATOM | 407 | CA | PRO | A | 463 | -20.794 | -3.137 | 2.871 | 1.00 | 66.67 | C |
| ATOM | 408 | CA | PHE | A | 464 | -23.105 | -1.066 | 5.134 | 1.00 | 64.34 | C |
| ATOM | 409 | CA | GLU | A | 465 | -25.120 | -4.197 | 6.085 | 1.00 | 69.98 | C |
| ATOM | 410 | CA | ARG | A | 466 | -26.368 | -3.949 | 9.668 | 1.00 | 68.32 | C |
| ATOM | 411 | CA | ASP | A | 467 | -26.973 | -7.230 | 11.501 | 1.00 | 69.46 | C |
| ATOM | 412 | CA | ILE | A | 468 | -28.784 | -7.047 | 14.851 | 1.00 | 68.99 | C |
| ATOM | 413 | CA | SER | A | 469 | -29.808 | -10.681 | 15.285 | 1.00 | 71.13 | C |
| ATOM | 414 | CA | THR | A | 470 | -28.897 | -12.907 | 18.247 | 1.00 | 75.87 | C |
| ATOM | 415 | CA | GLU | A | 471 | -29.255 | -16.277 | 16.532 | 1.00 | 80.68 | C |
| ATOM | 416 | CA | ILE | A | 472 | -27.001 | -18.982 | 17.981 | 1.00 | 79.70 | C |
| ATOM | 417 | CA | TYR | A | 473 | -23.968 | -19.603 | 15.781 | 1.00 | 78.75 | C |
| ATOM | 418 | CA | GLN | A | 474 | -23.412 | -23.184 | 14.638 | 1.00 | 86.46 | C |
| ATOM | 419 | CA | ALA | A | 475 | -19.684 | -23.901 | 14.625 | 1.00 | 87.28 | C |
| ATOM | 420 | CA | GLY | A | 476 | -19.882 | -27.602 | 13.793 | 1.00 | 91.87 | C |
| ATOM | 421 | CA | SER | A | 477 | -22.007 | -29.927 | 11.673 | 1.00 | 95.87 | C |
| ATOM | 422 | CA | THR | A | 478 | -24.648 | -30.534 | 14.383 | 1.00 | 95.31 | C |
| ATOM | 423 | CA | PRO | A | 479 | -27.741 | -28.260 | 14.452 | 1.00 | 95.93 | C |
| ATOM | 424 | CA | CYS | A | 480 | -27.955 | -26.115 | 17.573 | 1.00 | 97.13 | C |
| ATOM | 425 | CA | ASN | A | 481 | -31.771 | -25.686 | 17.751 | 1.00 | 98.17 | C |
| ATOM | 426 | CA | GLY | A | 482 | -31.441 | -22.511 | 19.797 | 1.00 | 93.77 | C |
| ATOM | 427 | CA | VAL | A | 483 | -29.483 | -24.046 | 22.707 | 1.00 | 92.36 | C |
| ATOM | 428 | CA | GLU | A | 484 | -26.113 | -22.714 | 23.896 | 1.00 | 88.89 | C |
| ATOM | 429 | CA | GLY | A | 485 | -23.576 | -25.513 | 24.241 | 1.00 | 84.82 | C |
| ATOM | 430 | CA | PHE | A | 486 | -20.748 | -27.329 | 22.489 | 1.00 | 84.40 | C |
| ATOM | 431 | CA | ASN | A | 487 | -20.143 | -25.856 | 18.986 | 1.00 | 85.43 | C |
| ATOM | 432 | CA | CYS | A | 488 | -23.161 | -23.575 | 19.571 | 1.00 | 85.11 | C |
| ATOM | 433 | CA | TYR | A | 489 | -22.126 | -20.063 | 20.554 | 1.00 | 78.01 | C |
| ATOM | 434 | CA | PHE | A | 490 | -23.749 | -16.744 | 21.368 | 1.00 | 74.59 | C |
| ATOM | 435 | CA | PRO | A | 491 | -22.746 | -14.417 | 18.496 | 1.00 | 70.90 | C |
| ATOM | 436 | CA | LEU | A | 492 | -22.370 | -11.201 | 20.521 | 1.00 | 68.94 | C |
| ATOM | 437 | CA | GLN | A | 493 | -19.756 | -10.341 | 23.133 | 1.00 | 72.02 | C |
| ATOM | 438 | CA | SER | A | 494 | -19.140 | -7.285 | 25.291 | 1.00 | 72.88 | C |
| ATOM | 439 | CA | TYR | A | 495 | -16.051 | -5.129 | 25.700 | 1.00 | 71.91 | C |
| ATOM | 440 | CA | GLY | A | 496 | -15.180 | -4.816 | 29.361 | 1.00 | 78.12 | C |
| ATOM | 441 | CA | PHE | A | 497 | -14.173 | -1.153 | 29.325 | 1.00 | 74.73 | C |
| ATOM | 442 | CA | GLN | A | 498 | -12.220 | -0.322 | 32.495 | 1.00 | 81.84 | C |
| ATOM | 443 | CA | PRO | A | 499 | -10.239 | 2.867 | 33.296 | 1.00 | 81.62 | C |
| ATOM | 444 | CA | THR | A | 500 | -7.128 | 0.803 | 34.132 | 1.00 | 84.02 | C |
| ATOM | 445 | CA | ASN | A | 501 | -7.008 | -0.923 | 30.734 | 1.00 | 79.60 | C |
| ATOM | 446 | CA | GLY | A | 502 | -4.194 | -0.286 | 28.301 | 1.00 | 78.09 | C |
| ATOM | 447 | CA | VAL | A | 503 | -4.595 | 2.171 | 25.430 | 1.00 | 76.46 | C |
| ATOM | 448 | CA | GLY | A | 504 | -5.339 | -0.563 | 22.883 | 1.00 | 74.62 | C |
| ATOM | 449 | CA | TYR | A | 505 | -8.212 | -1.756 | 25.112 | 1.00 | 75.31 | C |
| ATOM | 450 | CA | GLN | A | 506 | -9.664 | 1.655 | 25.931 | 1.00 | 73.75 | C |
| ATOM | 451 | CA | PRO | A | 507 | -12.942 | 2.660 | 24.240 | 1.00 | 69.85 | C |
| ATOM | 452 | CA | TYR | A | 508 | -12.815 | 4.780 | 21.117 | 1.00 | 66.30 | C |
| ATOM | 453 | CA | ARG | A | 509 | -15.845 | 6.487 | 19.649 | 1.00 | 63.83 | C |
| ATOM | 454 | CA | VAL | A | 510 | -15.855 | 6.345 | 15.857 | 1.00 | 59.80 | C |
| ATOM | 455 | CA | VAL | A | 511 | -17.832 | 8.525 | 13.462 | 1.00 | 58.92 | C |
| ATOM | 456 | CA | VAL | A | 512 | -17.573 | 7.586 | 9.800 | 1.00 | 58.97 | C |
| ATOM | 457 | CA | LEU | A | 513 | -18.806 | 9.961 | 7.125 | 1.00 | 62.35 | C |
| ATOM | 458 | CA | SER | A | 514 | -19.793 | 8.449 | 3.801 | 1.00 | 67.78 | C |
| ATOM | 459 | CA | PHE | A | 515 | -20.013 | 10.920 | 0.924 | 1.00 | 70.04 | C |
| ATOM | 460 | CA | GLU | A | 516 | -22.292 | 9.714 | -1.857 | 1.00 | 84.82 | C |

| | | | | | | | | | | | |
|---|---|---|---|---|---|---|---|---|---|---|---|
| ATOM | 461 | CA | LEU | A | 517 | -21.447 | 10.870 | -5.393 | 1.00 | 85.40 | C |
| ATOM | 462 | CA | LEU | A | 518 | -24.075 | 9.959 | -7.997 | 1.00 | 90.69 | C |
| ATOM | 463 | CA | HIS | A | 519 | -25.408 | 12.043 | -10.890 | 1.00 | 96.72 | C |
| ATOM | 464 | CA | ALA | A | 520 | -28.160 | 13.726 | -8.890 | 1.00 | 90.51 | C |
| ATOM | 465 | CA | PRO | A | 521 | -28.699 | 17.147 | -7.229 | 1.00 | 86.05 | C |
| ATOM | 466 | CA | ALA | A | 522 | -26.195 | 17.880 | -4.493 | 1.00 | 81.41 | C |
| ATOM | 467 | CA | THR | A | 523 | -27.677 | 18.808 | -1.118 | 1.00 | 80.28 | C |
| ATOM | 468 | CA | VAL | A | 524 | -24.631 | 18.985 | 1.190 | 1.00 | 76.20 | C |
| ATOM | 469 | CA | CYS | A | 525 | -22.079 | 21.673 | 0.355 | 1.00 | 77.21 | C |
| ATOM | 470 | CA | GLY | A | 526 | -19.121 | 23.254 | 2.114 | 1.00 | 74.90 | C |
| ATOM | 471 | CA | PRO | A | 527 | -19.152 | 26.820 | 3.492 | 1.00 | 76.85 | C |
| ATOM | 472 | CA | LYS | A | 528 | -18.210 | 28.542 | 0.243 | 1.00 | 80.42 | C |
| ATOM | 473 | CA | LYS | A | 529 | -19.316 | 32.033 | -0.696 | 1.00 | 83.33 | C |
| ATOM | 474 | CA | SER | A | 530 | -21.049 | 32.478 | -4.031 | 1.00 | 85.01 | C |
| ATOM | 475 | CA | THR | A | 531 | -20.064 | 35.110 | -6.578 | 1.00 | 85.47 | C |
| ATOM | 476 | CA | ASN | A | 532 | -22.003 | 36.984 | -9.232 | 1.00 | 87.21 | C |
| ATOM | 477 | CA | LEU | A | 533 | -22.663 | 35.100 | -12.447 | 1.00 | 86.00 | C |
| ATOM | 478 | CA | VAL | A | 534 | -20.543 | 36.339 | -15.369 | 1.00 | 86.67 | C |
| ATOM | 479 | CA | LYS | A | 535 | -21.569 | 35.506 | -18.930 | 1.00 | 85.77 | C |
| ATOM | 480 | CA | ASN | A | 536 | -19.864 | 35.705 | -22.348 | 1.00 | 87.25 | C |
| ATOM | 481 | CA | LYS | A | 537 | -16.347 | 35.568 | -20.869 | 1.00 | 88.38 | C |
| ATOM | 482 | CA | CYS | A | 538 | -13.772 | 32.834 | -20.259 | 1.00 | 90.72 | C |
| ATOM | 483 | CA | VAL | A | 539 | -13.969 | 31.964 | -16.543 | 1.00 | 87.13 | C |
| ATOM | 484 | CA | ASN | A | 540 | -13.130 | 29.158 | -14.144 | 1.00 | 85.65 | C |
| ATOM | 485 | CA | PHE | A | 541 | -16.482 | 27.987 | -12.818 | 1.00 | 83.32 | C |
| ATOM | 486 | CA | ASN | A | 542 | -17.724 | 25.813 | -9.969 | 1.00 | 83.32 | C |
| ATOM | 487 | CA | PHE | A | 543 | -21.452 | 24.995 | -10.221 | 1.00 | 82.01 | C |
| ATOM | 488 | CA | ASN | A | 544 | -22.541 | 22.934 | -7.168 | 1.00 | 83.98 | C |
| ATOM | 489 | CA | GLY | A | 545 | -19.280 | 20.987 | -7.150 | 1.00 | 83.42 | C |
| ATOM | 490 | CA | LEU | A | 546 | -18.933 | 20.755 | -10.943 | 1.00 | 83.99 | C |
| ATOM | 491 | CA | THR | A | 547 | -15.702 | 22.510 | -11.907 | 1.00 | 82.61 | C |
| ATOM | 492 | CA | GLY | A | 548 | -14.136 | 23.530 | -15.174 | 1.00 | 83.33 | C |
| ATOM | 493 | CA | THR | A | 549 | -13.080 | 26.338 | -17.471 | 1.00 | 85.69 | C |
| ATOM | 494 | CA | GLY | A | 550 | -15.274 | 27.858 | -20.150 | 1.00 | 85.00 | C |
| ATOM | 495 | CA | VAL | A | 551 | -17.589 | 30.562 | -21.437 | 1.00 | 86.11 | C |
| ATOM | 496 | CA | LEU | A | 552 | -21.097 | 30.761 | -19.988 | 1.00 | 84.47 | C |
| ATOM | 497 | CA | THR | A | 553 | -23.905 | 31.849 | -22.314 | 1.00 | 86.12 | C |
| ATOM | 498 | CA | GLU | A | 554 | -27.679 | 31.919 | -22.078 | 1.00 | 89.96 | C |
| ATOM | 499 | CA | SER | A | 555 | -29.086 | 28.616 | -23.289 | 1.00 | 90.16 | C |
| ATOM | 500 | CA | ASN | A | 556 | -32.055 | 27.849 | -25.522 | 1.00 | 93.26 | C |
| ATOM | 501 | CA | LYS | A | 557 | -32.250 | 24.233 | -24.352 | 1.00 | 90.04 | C |
| ATOM | 502 | CA | LYS | A | 558 | -35.567 | 23.041 | -22.927 | 1.00 | 92.86 | C |
| ATOM | 503 | CA | PHE | A | 559 | -34.419 | 21.394 | -19.716 | 1.00 | 89.55 | C |
| ATOM | 504 | CA | LEU | A | 560 | -36.948 | 19.282 | -17.882 | 1.00 | 90.26 | C |
| ATOM | 505 | CA | PRO | A | 561 | -37.685 | 20.350 | -14.246 | 1.00 | 90.12 | C |
| ATOM | 506 | CA | PHE | A | 562 | -35.476 | 17.564 | -12.797 | 1.00 | 91.20 | C |
| ATOM | 507 | CA | GLN | A | 563 | -32.413 | 18.044 | -15.017 | 1.00 | 90.15 | C |
| ATOM | 508 | CA | GLN | A | 564 | -29.461 | 19.793 | -13.381 | 1.00 | 90.92 | C |
| ATOM | 509 | CA | PHE | A | 565 | -26.899 | 19.815 | -16.202 | 1.00 | 90.11 | C |
| ATOM | 510 | CA | GLY | A | 566 | -26.536 | 18.720 | -19.795 | 1.00 | 90.78 | C |
| ATOM | 511 | CA | ARG | A | 567 | -24.153 | 16.782 | -21.987 | 1.00 | 95.02 | C |
| ATOM | 512 | CA | ASP | A | 568 | -23.162 | 16.865 | -25.645 | 1.00103.17 | | C |
| ATOM | 513 | CA | ILE | A | 569 | -22.923 | 14.057 | -28.221 | 1.00102.95 | | C |
| ATOM | 514 | CA | ALA | A | 570 | -19.393 | 13.522 | -26.818 | 1.00100.39 | | C |
| ATOM | 515 | CA | ASP | A | 571 | -20.862 | 12.779 | -23.330 | 1.00100.43 | | C |
| ATOM | 516 | CA | THR | A | 572 | -19.067 | 15.808 | -21.863 | 1.00100.31 | | C |
| ATOM | 517 | CA | THR | A | 573 | -20.771 | 18.555 | -19.860 | 1.00 | 93.89 | C |
| ATOM | 518 | CA | ASP | A | 574 | -22.113 | 21.362 | -22.057 | 1.00 | 92.91 | C |
| ATOM | 519 | CA | ALA | A | 575 | -24.888 | 23.010 | -20.017 | 1.00 | 87.91 | C |
| ATOM | 520 | CA | VAL | A | 576 | -25.649 | 23.616 | -16.360 | 1.00 | 85.40 | C |

ATOM 521 CA ARG A 577 -28.623 24.789 -14.303 1.00 86.15 C
ATOM 522 CA ASP A 578 -27.551 27.535 -11.893 1.00 86.07 C
ATOM 523 CA PRO A 579 -28.134 26.460 -8.251 1.00 85.92 C
ATOM 524 CA GLN A 580 -29.629 29.806 -7.143 1.00 89.40 C
ATOM 525 CA THR A 581 -31.740 30.938 -10.116 1.00 88.89 C
ATOM 526 CA LEU A 582 -33.139 28.176 -12.314 1.00 90.93 C
ATOM 527 CA GLU A 583 -31.530 29.532 -15.477 1.00 89.96 C
ATOM 528 CA ILE A 584 -30.098 27.169 -18.074 1.00 87.89 C
ATOM 529 CA LEU A 585 -26.586 28.108 -19.171 1.00 85.74 C
ATOM 530 CA ASP A 586 -24.558 26.744 -22.064 1.00 89.32 C
ATOM 531 CA ILE A 587 -20.888 26.049 -21.425 1.00 86.26 C
ATOM 532 CA THR A 588 -18.514 26.358 -24.319 1.00 87.80 C
ATOM 533 CA PRO A 589 -14.752 25.828 -24.055 1.00 87.11 C
ATOM 534 CA CYS A 590 -12.515 28.862 -24.030 1.00 90.03 C
ATOM 535 CA SER A 591 -11.143 29.603 -27.501 1.00 88.40 C
ATOM 536 CA PHE A 592 -8.229 27.424 -28.532 1.00 85.59 C
ATOM 537 CA GLY A 593 -6.515 26.222 -31.652 1.00 80.96 C
ATOM 538 CA GLY A 594 -3.237 25.234 -33.170 1.00 77.15 C
ATOM 539 CA VAL A 595 -0.531 27.822 -33.739 1.00 75.93 C
ATOM 540 CA SER A 596 1.408 27.489 -36.971 1.00 73.64 C
ATOM 541 CA VAL A 597 4.361 29.661 -37.955 1.00 72.67 C
ATOM 542 CA ILE A 598 4.709 30.569 -41.629 1.00 73.23 C
ATOM 543 CA THR A 599 8.235 31.406 -42.531 1.00 77.08 C
ATOM 544 CA PRO A 600 10.749 31.800 -45.330 1.00 81.38 C
ATOM 545 CA GLY A 601 13.988 30.052 -44.483 1.00 87.04 C
ATOM 546 CA THR A 602 16.562 31.573 -42.158 1.00 91.36 C
ATOM 547 CA ASN A 603 18.869 31.882 -45.173 1.00102.88 C
ATOM 548 CA THR A 604 16.557 34.634 -46.488 1.00 92.37 C
ATOM 549 CA SER A 605 14.722 36.358 -43.620 1.00 86.72 C
ATOM 550 CA ASN A 606 13.810 36.160 -39.938 1.00 84.55 C
ATOM 551 CA GLN A 607 10.324 37.591 -40.575 1.00 82.07 C
ATOM 552 CA VAL A 608 7.411 35.267 -39.766 1.00 75.31 C
ATOM 553 CA ALA A 609 3.616 35.143 -39.964 1.00 72.69 C
ATOM 554 CA VAL A 610 1.389 33.285 -37.497 1.00 71.95 C
ATOM 555 CA LEU A 611 -1.746 31.285 -38.289 1.00 73.43 C
ATOM 556 CA TYR A 612 -4.191 30.699 -35.439 1.00 77.15 C
ATOM 557 CA GLN A 613 -5.952 27.625 -36.786 1.00 77.19 C
ATOM 558 CA GLY A 614 -9.759 27.731 -36.623 1.00 80.52 C
ATOM 559 CA VAL A 615 -9.744 30.849 -34.430 1.00 83.87 C
ATOM 560 CA ASN A 616 -11.478 34.185 -35.002 1.00 93.09 C
ATOM 561 CA CYS A 617 -9.395 37.365 -35.255 1.00 92.60 C
ATOM 562 CA THR A 618 -11.465 38.936 -32.440 1.00 94.02 C
ATOM 563 CA GLU A 619 -10.357 36.296 -29.911 1.00 94.07 C
ATOM 564 CA VAL A 620 -6.562 36.225 -30.402 1.00 90.69 C
ATOM 565 CA PRO A 621 -5.563 39.299 -28.395 1.00 95.43 C
ATOM 566 CA ASN A 641 -1.822 42.844 -41.113 1.00 80.92 C
ATOM 567 CA VAL A 642 -4.621 40.756 -39.590 1.00 78.56 C
ATOM 568 CA PHE A 643 -6.404 38.623 -42.209 1.00 81.19 C
ATOM 569 CA GLN A 644 -9.333 36.329 -41.379 1.00 85.02 C
ATOM 570 CA THR A 645 -9.596 33.243 -43.584 1.00 82.48 C
ATOM 571 CA ARG A 646 -11.339 29.867 -43.399 1.00 81.50 C
ATOM 572 CA ALA A 647 -8.137 28.262 -42.106 1.00 76.69 C
ATOM 573 CA GLY A 648 -7.980 30.696 -39.185 1.00 79.89 C
ATOM 574 CA CYS A 649 -6.669 34.120 -38.277 1.00 81.29 C
ATOM 575 CA LEU A 650 -3.422 34.946 -40.107 1.00 75.86 C
ATOM 576 CA ILE A 651 -1.354 37.719 -38.495 1.00 74.59 C
ATOM 577 CA GLY A 652 1.755 39.236 -40.073 1.00 76.40 C
ATOM 578 CA ALA A 653 1.004 38.462 -43.734 1.00 78.70 C
ATOM 579 CA GLU A 654 -0.304 41.192 -46.032 1.00 81.82 C
ATOM 580 CA HIS A 655 -3.363 40.096 -48.000 1.00 82.35 C

| | | | | | | | | | | | |
|---|---|---|---|---|---|---|---|---|---|---|---|
| ATOM | 581 | CA | VAL | A | 656 | -3.173 | 40.992 | -51.697 | 1.00 | 82.31 | C |
| ATOM | 582 | CA | ASN | A | 657 | -5.613 | 40.695 | -54.594 | 1.00 | 88.92 | C |
| ATOM | 583 | CA | ASN | A | 658 | -2.927 | 39.319 | -56.946 | 1.00 | 84.17 | C |
| ATOM | 584 | CA | SER | A | 659 | -2.913 | 35.550 | -57.324 | 1.00 | 79.15 | C |
| ATOM | 585 | CA | TYR | A | 660 | 0.308 | 33.578 | -57.732 | 1.00 | 79.30 | C |
| ATOM | 586 | CA | GLU | A | 661 | 1.443 | 29.971 | -57.644 | 1.00 | 81.75 | C |
| ATOM | 587 | CA | CYS | A | 662 | 1.290 | 28.358 | -54.211 | 1.00 | 78.66 | C |
| ATOM | 588 | CA | ASP | A | 663 | 4.519 | 28.611 | -52.218 | 1.00 | 77.77 | C |
| ATOM | 589 | CA | ILE | A | 664 | 3.795 | 28.008 | -48.522 | 1.00 | 72.37 | C |
| ATOM | 590 | CA | PRO | A | 665 | 0.294 | 26.505 | -48.137 | 1.00 | 69.74 | C |
| ATOM | 591 | CA | ILE | A | 666 | -2.045 | 28.072 | -45.597 | 1.00 | 68.22 | C |
| ATOM | 592 | CA | GLY | A | 667 | -5.286 | 26.290 | -46.389 | 1.00 | 65.18 | C |
| ATOM | 593 | CA | ALA | A | 668 | -8.711 | 26.826 | -48.022 | 1.00 | 64.92 | C |
| ATOM | 594 | CA | GLY | A | 669 | -7.171 | 28.121 | -51.219 | 1.00 | 67.76 | C |
| ATOM | 595 | CA | ILE | A | 670 | -4.845 | 30.530 | -49.403 | 1.00 | 67.99 | C |
| ATOM | 596 | CA | CYS | A | 671 | -1.060 | 30.311 | -49.796 | 1.00 | 74.44 | C |
| ATOM | 597 | CA | ALA | A | 672 | 1.710 | 32.562 | -48.505 | 1.00 | 73.43 | C |
| ATOM | 598 | CA | SER | A | 673 | 5.082 | 33.624 | -49.857 | 1.00 | 78.65 | C |
| ATOM | 599 | CA | TYR | A | 674 | 7.962 | 35.999 | -49.180 | 1.00 | 84.10 | C |
| ATOM | 600 | CA | GLN | A | 675 | 8.192 | 38.752 | -51.805 | 1.00 | 87.45 | C |
| ATOM | 601 | CA | GLN | A | 690 | 10.215 | 42.590 | -48.930 | 1.00 | 87.44 | C |
| ATOM | 602 | CA | SER | A | 691 | 7.549 | 41.024 | -46.719 | 1.00 | 83.56 | C |
| ATOM | 603 | CA | ILE | A | 692 | 5.372 | 37.947 | -46.312 | 1.00 | 79.50 | C |
| ATOM | 604 | CA | ILE | A | 693 | 2.118 | 38.073 | -48.270 | 1.00 | 78.27 | C |
| ATOM | 605 | CA | ALA | A | 694 | -0.969 | 35.873 | -48.409 | 1.00 | 76.38 | C |
| ATOM | 606 | CA | TYR | A | 695 | -3.059 | 35.305 | -51.505 | 1.00 | 74.39 | C |
| ATOM | 607 | CA | THR | A | 696 | -5.553 | 32.996 | -53.106 | 1.00 | 72.06 | C |
| ATOM | 608 | CA | MET | A | 697 | -3.400 | 30.611 | -55.111 | 1.00 | 72.25 | C |
| ATOM | 609 | CA | SER | A | 698 | -3.516 | 30.512 | -58.886 | 1.00 | 72.79 | C |
| ATOM | 610 | CA | LEU | A | 699 | -4.245 | 27.203 | -60.570 | 1.00 | 65.63 | C |
| ATOM | 611 | CA | GLY | A | 700 | -2.299 | 28.197 | -63.684 | 1.00 | 69.85 | C |
| ATOM | 612 | CA | ALA | A | 701 | -2.420 | 30.593 | -66.580 | 1.00 | 72.84 | C |
| ATOM | 613 | CA | GLU | A | 702 | -5.600 | 30.655 | -68.609 | 1.00 | 76.61 | C |
| ATOM | 614 | CA | ASN | A | 703 | -5.432 | 29.396 | -72.186 | 1.00 | 73.07 | C |
| ATOM | 615 | CA | SER | A | 704 | -8.570 | 29.439 | -74.315 | 1.00 | 73.88 | C |
| ATOM | 616 | CA | VAL | A | 705 | -8.040 | 26.909 | -77.097 | 1.00 | 68.83 | C |
| ATOM | 617 | CA | ALA | A | 706 | -9.123 | 28.172 | -80.529 | 1.00 | 66.86 | C |
| ATOM | 618 | CA | TYR | A | 707 | -11.547 | 25.343 | -81.216 | 1.00 | 63.97 | C |
| ATOM | 619 | CA | SER | A | 708 | -13.535 | 25.236 | -84.442 | 1.00 | 65.02 | C |
| ATOM | 620 | CA | ASN | A | 709 | -15.048 | 22.486 | -86.563 | 1.00 | 67.84 | C |
| ATOM | 621 | CA | ASN | A | 710 | -12.442 | 22.777 | -89.344 | 1.00 | 64.71 | C |
| ATOM | 622 | CA | SER | A | 711 | -9.120 | 23.870 | -87.769 | 1.00 | 62.57 | C |
| ATOM | 623 | CA | ILE | A | 712 | -6.320 | 21.580 | -86.593 | 1.00 | 59.22 | C |
| ATOM | 624 | CA | ALA | A | 713 | -2.970 | 22.520 | -85.107 | 1.00 | 57.90 | C |
| ATOM | 625 | CA | ILE | A | 714 | 0.025 | 20.386 | -86.144 | 1.00 | 59.18 | C |
| ATOM | 626 | CA | PRO | A | 715 | 3.573 | 20.714 | -84.768 | 1.00 | 58.94 | C |
| ATOM | 627 | CA | THR | A | 716 | 6.199 | 21.814 | -87.265 | 1.00 | 63.68 | C |
| ATOM | 628 | CA | ASN | A | 717 | 9.129 | 21.289 | -84.898 | 1.00 | 66.22 | C |
| ATOM | 629 | CA | PHE | A | 718 | 9.830 | 19.467 | -81.653 | 1.00 | 58.95 | C |
| ATOM | 630 | CA | THR | A | 719 | 12.128 | 19.378 | -78.661 | 1.00 | 56.69 | C |
| ATOM | 631 | CA | ILE | A | 720 | 13.720 | 16.515 | -76.770 | 1.00 | 54.02 | C |
| ATOM | 632 | CA | SER | A | 721 | 13.041 | 16.803 | -73.049 | 1.00 | 52.61 | C |
| ATOM | 633 | CA | VAL | A | 722 | 14.667 | 14.966 | -70.152 | 1.00 | 52.37 | C |
| ATOM | 634 | CA | THR | A | 723 | 12.584 | 14.835 | -66.968 | 1.00 | 54.92 | C |
| ATOM | 635 | CA | THR | A | 724 | 14.008 | 13.712 | -63.623 | 1.00 | 57.76 | C |
| ATOM | 636 | CA | GLU | A | 725 | 11.892 | 11.387 | -61.466 | 1.00 | 54.47 | C |
| ATOM | 637 | CA | ILE | A | 726 | 13.237 | 10.400 | -58.036 | 1.00 | 53.43 | C |
| ATOM | 638 | CA | LEU | A | 727 | 11.841 | 7.363 | -56.226 | 1.00 | 51.42 | C |
| ATOM | 639 | CA | PRO | A | 728 | 12.743 | 5.766 | -52.883 | 1.00 | 50.51 | C |
| ATOM | 640 | CA | VAL | A | 729 | 13.707 | 2.108 | -53.200 | 1.00 | 52.59 | C |

```
ATOM    641  CA  SER A 730      14.966   1.073 -49.783  1.00 57.05           C
ATOM    642  CA  MET A 731      15.058   2.103 -46.153  1.00 61.91           C
ATOM    643  CA  THR A 732      17.854   1.861 -43.584  1.00 63.16           C
ATOM    644  CA  LYS A 733      17.837  -1.666 -42.160  1.00 61.20           C
ATOM    645  CA  THR A 734      17.694  -1.021 -38.466  1.00 64.70           C
ATOM    646  CA  SER A 735      17.491  -3.670 -35.776  1.00 69.77           C
ATOM    647  CA  VAL A 736      16.848  -3.480 -32.050  1.00 70.58           C
ATOM    648  CA  ASP A 737      17.540  -6.101 -29.418  1.00 74.56           C
ATOM    649  CA  CYS A 738      14.360  -5.880 -27.337  1.00 72.69           C
ATOM    650  CA  THR A 739      15.941  -7.231 -24.155  1.00 70.71           C
ATOM    651  CA  MET A 740      19.024  -5.032 -24.608  1.00 72.60           C
ATOM    652  CA  TYR A 741      16.921  -1.912 -25.189  1.00 63.75           C
ATOM    653  CA  ILE A 742      14.506  -2.513 -22.317  1.00 64.34           C
ATOM    654  CA  CYS A 743      16.658  -4.389 -19.791  1.00 71.51           C
ATOM    655  CA  GLY A 744      20.308  -3.735 -20.715  1.00 73.94           C
ATOM    656  CA  ASP A 745      22.832  -5.838 -18.729  1.00 81.48           C
ATOM    657  CA  SER A 746      20.241  -6.624 -16.026  1.00 73.75           C
ATOM    658  CA  THR A 747      19.227 -10.155 -15.040  1.00 74.26           C
ATOM    659  CA  GLU A 748      16.453  -8.898 -12.720  1.00 73.93           C
ATOM    660  CA  CYS A 749      14.823  -6.770 -15.420  1.00 71.95           C
ATOM    661  CA  SER A 750      15.137  -9.576 -17.991  1.00 70.91           C
ATOM    662  CA  ASN A 751      13.435 -12.002 -15.601  1.00 69.21           C
ATOM    663  CA  LEU A 752      10.632  -9.494 -15.007  1.00 67.16           C
ATOM    664  CA  LEU A 753      10.457  -9.053 -18.809  1.00 65.18           C
ATOM    665  CA  LEU A 754       9.796 -12.779 -19.190  1.00 66.56           C
ATOM    666  CA  GLN A 755       6.324 -12.233 -17.664  1.00 66.06           C
ATOM    667  CA  TYR A 756       5.284 -10.459 -20.898  1.00 64.44           C
ATOM    668  CA  GLY A 757       5.848 -13.599 -22.975  1.00 67.19           C
ATOM    669  CA  SER A 758       6.842 -13.320 -26.626  1.00 69.39           C
ATOM    670  CA  PHE A 759       6.573  -9.512 -27.176  1.00 67.57           C
ATOM    671  CA  CYS A 760      10.320  -9.157 -27.806  1.00 70.84           C
ATOM    672  CA  THR A 761      10.335 -12.118 -30.206  1.00 66.16           C
ATOM    673  CA  GLN A 762       7.571 -10.712 -32.406  1.00 65.62           C
ATOM    674  CA  LEU A 763       9.167  -7.241 -32.383  1.00 63.38           C
ATOM    675  CA  ASN A 764      12.516  -8.719 -33.472  1.00 64.96           C
ATOM    676  CA  ARG A 765      10.699 -10.760 -36.124  1.00 64.20           C
ATOM    677  CA  ALA A 766       8.904  -7.660 -37.436  1.00 57.10           C
ATOM    678  CA  LEU A 767      12.174  -5.726 -37.756  1.00 57.59           C
ATOM    679  CA  THR A 768      13.879  -8.714 -39.416  1.00 59.57           C
ATOM    680  CA  GLY A 769      10.997  -8.889 -41.889  1.00 57.35           C
ATOM    681  CA  ILE A 770      11.424  -5.186 -42.666  1.00 55.99           C
ATOM    682  CA  ALA A 771      15.178  -5.649 -43.302  1.00 57.84           C
ATOM    683  CA  VAL A 772      14.514  -8.619 -45.609  1.00 58.09           C
ATOM    684  CA  GLU A 773      11.854  -6.580 -47.418  1.00 60.34           C
ATOM    685  CA  GLN A 774      14.350  -3.741 -48.002  1.00 57.60           C
ATOM    686  CA  ASP A 775      16.799  -6.162 -49.609  1.00 60.41           C
ATOM    687  CA  LYS A 776      13.973  -7.611 -51.719  1.00 58.18           C
ATOM    688  CA  ASN A 777      12.956  -4.086 -52.788  1.00 55.84           C
ATOM    689  CA  THR A 778      16.499  -3.274 -53.915  1.00 55.36           C
ATOM    690  CA  GLN A 779      16.790  -6.598 -55.766  1.00 58.83           C
ATOM    691  CA  GLU A 780      13.459  -6.112 -57.559  1.00 58.93           C
ATOM    692  CA  VAL A 781      14.291  -2.575 -58.700  1.00 51.87           C
ATOM    693  CA  PHE A 782      17.865  -3.081 -59.864  1.00 52.94           C
ATOM    694  CA  ALA A 783      18.514  -6.795 -60.541  1.00 55.61           C
ATOM    695  CA  GLN A 784      16.268  -6.961 -63.603  1.00 57.79           C
ATOM    696  CA  VAL A 785      19.020  -7.820 -66.111  1.00 63.17           C
ATOM    697  CA  LYS A 786      20.362 -11.352 -66.558  1.00 71.81           C
ATOM    698  CA  GLN A 787      23.756 -10.315 -67.981  1.00 70.05           C
ATOM    699  CA  ILE A 788      26.204  -7.502 -67.227  1.00 67.11           C
ATOM    700  CA  TYR A 789      26.335  -5.488 -70.453  1.00 62.77           C
```

```
ATOM    701  CA  LYS A 790      29.202  -3.173 -71.390  1.00 69.12           C
ATOM    702  CA  THR A 791      29.480  -0.379 -73.938  1.00 70.90           C
ATOM    703  CA  PRO A 792      32.165  -0.541 -76.667  1.00 74.23           C
ATOM    704  CA  PRO A 793      35.454   1.371 -76.244  1.00 77.87           C
ATOM    705  CA  ILE A 794      34.525   3.669 -79.157  1.00 78.03           C
ATOM    706  CA  LYS A 795      31.401   5.539 -78.075  1.00 73.93           C
ATOM    707  CA  ASP A 796      30.216   6.524 -81.555  1.00 72.24           C
ATOM    708  CA  PHE A 797      26.435   6.657 -81.123  1.00 63.88           C
ATOM    709  CA  GLY A 798      25.528   8.788 -84.127  1.00 61.83           C
ATOM    710  CA  GLY A 799      26.138  12.056 -82.294  1.00 59.08           C
ATOM    711  CA  PHE A 800      24.394  11.067 -79.053  1.00 59.11           C
ATOM    712  CA  ASN A 801      26.629  11.705 -76.036  1.00 65.87           C
ATOM    713  CA  PHE A 802      26.112   9.520 -72.956  1.00 60.84           C
ATOM    714  CA  SER A 803      29.199  10.516 -70.904  1.00 67.08           C
ATOM    715  CA  GLN A 804      27.003  12.119 -68.227  1.00 68.69           C
ATOM    716  CA  ILE A 805      25.093   8.915 -67.531  1.00 62.23           C
ATOM    717  CA  LEU A 806      27.820   6.261 -68.298  1.00 64.36           C
ATOM    718  CA  PRO A 807      30.564   5.382 -65.769  1.00 71.03           C
ATOM    719  CA  ASP A 808      33.868   7.284 -65.884  1.00 84.72           C
ATOM    720  CA  PRO A 809      36.593   4.838 -64.651  1.00 88.51           C
ATOM    721  CA  SER A 810      39.075   7.640 -63.759  1.00 92.50           C
ATOM    722  CA  LYS A 811      38.050   7.337 -60.076  1.00 89.86           C
ATOM    723  CA  PRO A 812      37.989   4.302 -57.678  1.00 93.19           C
ATOM    724  CA  SER A 813      34.179   3.952 -57.555  1.00 89.03           C
ATOM    725  CA  LYS A 814      32.890   3.450 -61.092  1.00 84.09           C
ATOM    726  CA  ARG A 815      29.946   5.823 -60.874  1.00 78.24           C
ATOM    727  CA  SER A 816      28.810   8.319 -63.474  1.00 72.03           C
ATOM    728  CA  PRO A 817      29.018  12.139 -62.847  1.00 71.01           C
ATOM    729  CA  ILE A 818      25.220  12.542 -62.368  1.00 70.38           C
ATOM    730  CA  GLU A 819      25.309   9.634 -59.896  1.00 70.39           C
ATOM    731  CA  ASP A 820      28.197  11.384 -58.109  1.00 74.53           C
ATOM    732  CA  LEU A 821      25.974  14.462 -57.826  1.00 74.37           C
ATOM    733  CA  LEU A 822      23.137  12.357 -56.412  1.00 71.97           C
ATOM    734  CA  PHE A 823      25.397  10.730 -53.820  1.00 72.91           C
ATOM    735  CA  ASN A 824      27.066  14.012 -52.803  1.00 75.27           C
ATOM    736  CA  LYS A 825      23.732  15.802 -52.310  1.00 79.62           C
ATOM    737  CA  VAL A 826      22.270  12.982 -50.191  1.00 75.47           C
ATOM    738  CA  LEU A 849      25.437  10.104 -34.276  1.00102.28           C
ATOM    739  CA  ILE A 850      25.758   6.435 -35.269  1.00104.95           C
ATOM    740  CA  CYS A 851      27.652   5.554 -32.061  1.00105.87           C
ATOM    741  CA  ALA A 852      25.074   7.242 -29.794  1.00103.59           C
ATOM    742  CA  GLN A 853      22.278   5.242 -31.415  1.00 98.45           C
ATOM    743  CA  LYS A 854      24.293   1.998 -31.617  1.00 98.19           C
ATOM    744  CA  PHE A 855      25.218   1.974 -27.893  1.00 97.35           C
ATOM    745  CA  ASN A 856      21.620   1.180 -26.785  1.00 88.02           C
ATOM    746  CA  GLY A 857      21.290  -2.096 -28.698  1.00 80.49           C
ATOM    747  CA  LEU A 858      20.220  -0.413 -31.953  1.00 79.33           C
ATOM    748  CA  THR A 859      22.325  -1.772 -34.781  1.00 71.23           C
ATOM    749  CA  VAL A 860      22.186  -1.029 -38.498  1.00 64.33           C
ATOM    750  CA  LEU A 861      22.641  -3.839 -40.981  1.00 62.67           C
ATOM    751  CA  PRO A 862      24.521  -3.108 -44.216  1.00 62.91           C
ATOM    752  CA  PRO A 863      22.563  -3.459 -47.474  1.00 60.95           C
ATOM    753  CA  LEU A 864      23.083  -6.695 -49.369  1.00 61.55           C
ATOM    754  CA  LEU A 865      23.803  -4.808 -52.586  1.00 60.80           C
ATOM    755  CA  THR A 866      26.607  -2.278 -52.196  1.00 62.90           C
ATOM    756  CA  ASP A 867      26.638   1.105 -53.933  1.00 67.88           C
ATOM    757  CA  GLU A 868      29.214  -0.281 -56.383  1.00 68.28           C
ATOM    758  CA  MET A 869      26.845  -3.144 -57.218  1.00 63.60           C
ATOM    759  CA  ILE A 870      23.920  -0.778 -57.765  1.00 60.14           C
ATOM    760  CA  ALA A 871      26.171   1.263 -60.056  1.00 60.21           C
```

| | | | | | | | | | | | | |
|---|---|---|---|---|---|---|---|---|---|---|---|---|
| ATOM | 761 | CA | GLN | A | 872 | 27.128 | -1.938 | -61.917 | 1.00 | 61.33 | | C |
| ATOM | 762 | CA | TYR | A | 873 | 23.448 | -2.854 | -62.348 | 1.00 | 57.33 | | C |
| ATOM | 763 | CA | THR | A | 874 | 22.585 | 0.607 | -63.678 | 1.00 | 56.97 | | C |
| ATOM | 764 | CA | SER | A | 875 | 25.634 | 0.438 | -65.961 | 1.00 | 57.45 | | C |
| ATOM | 765 | CA | ALA | A | 876 | 24.510 | -2.959 | -67.304 | 1.00 | 57.12 | | C |
| ATOM | 766 | CA | LEU | A | 877 | 20.933 | -1.728 | -67.825 | 1.00 | 54.53 | | C |
| ATOM | 767 | CA | LEU | A | 878 | 22.197 | 1.421 | -69.520 | 1.00 | 55.87 | | C |
| ATOM | 768 | CA | ALA | A | 879 | 24.645 | -0.458 | -71.763 | 1.00 | 55.58 | | C |
| ATOM | 769 | CA | GLY | A | 880 | 21.828 | -2.787 | -72.786 | 1.00 | 54.84 | | C |
| ATOM | 770 | CA | THR | A | 881 | 19.456 | 0.112 | -73.478 | 1.00 | 54.48 | | C |
| ATOM | 771 | CA | ILE | A | 882 | 22.049 | 1.980 | -75.581 | 1.00 | 54.71 | | C |
| ATOM | 772 | CA | THR | A | 883 | 23.396 | -0.955 | -77.593 | 1.00 | 55.24 | | C |
| ATOM | 773 | CA | SER | A | 884 | 20.344 | -3.223 | -77.852 | 1.00 | 54.01 | | C |
| ATOM | 774 | CA | GLY | A | 885 | 17.284 | -0.996 | -77.295 | 1.00 | 52.64 | | C |
| ATOM | 775 | CA | TRP | A | 886 | 14.471 | -2.947 | -75.685 | 1.00 | 52.00 | | C |
| ATOM | 776 | CA | THR | A | 887 | 15.552 | -6.358 | -77.005 | 1.00 | 52.58 | | C |
| ATOM | 777 | CA | PHE | A | 888 | 17.657 | -7.176 | -73.920 | 1.00 | 52.54 | | C |
| ATOM | 778 | CA | GLY | A | 889 | 14.530 | -7.003 | -71.737 | 1.00 | 53.77 | | C |
| ATOM | 779 | CA | ALA | A | 890 | 12.609 | -9.659 | -73.648 | 1.00 | 53.08 | | C |
| ATOM | 780 | CA | GLY | A | 891 | 15.349 | -12.171 | -74.442 | 1.00 | 53.14 | | C |
| ATOM | 781 | CA | PRO | A | 892 | 18.883 | -12.163 | -75.853 | 1.00 | 53.32 | | C |
| ATOM | 782 | CA | ALA | A | 893 | 20.263 | -8.699 | -76.479 | 1.00 | 53.09 | | C |
| ATOM | 783 | CA | LEU | A | 894 | 20.303 | -7.898 | -80.189 | 1.00 | 54.32 | | C |
| ATOM | 784 | CA | GLN | A | 895 | 22.765 | -5.249 | -81.314 | 1.00 | 57.70 | | C |
| ATOM | 785 | CA | ILE | A | 896 | 21.372 | -2.253 | -83.168 | 1.00 | 55.39 | | C |
| ATOM | 786 | CA | PRO | A | 897 | 22.751 | 1.144 | -84.257 | 1.00 | 55.99 | | C |
| ATOM | 787 | CA | PHE | A | 898 | 21.400 | 3.815 | -81.908 | 1.00 | 55.53 | | C |
| ATOM | 788 | CA | PRO | A | 899 | 19.822 | 6.105 | -84.609 | 1.00 | 56.26 | | C |
| ATOM | 789 | CA | MET | A | 900 | 17.912 | 3.034 | -85.855 | 1.00 | 57.95 | | C |
| ATOM | 790 | CA | GLN | A | 901 | 16.761 | 2.334 | -82.284 | 1.00 | 55.58 | | C |
| ATOM | 791 | CA | MET | A | 902 | 15.628 | 5.964 | -82.060 | 1.00 | 55.46 | | C |
| ATOM | 792 | CA | ALA | A | 903 | 13.787 | 5.438 | -85.363 | 1.00 | 54.78 | | C |
| ATOM | 793 | CA | TYR | A | 904 | 11.886 | 2.568 | -83.772 | 1.00 | 56.97 | | C |
| ATOM | 794 | CA | ARG | A | 905 | 11.053 | 4.743 | -80.760 | 1.00 | 50.32 | | C |
| ATOM | 795 | CA | PHE | A | 906 | 9.818 | 7.496 | -83.095 | 1.00 | 51.44 | | C |
| ATOM | 796 | CA | ASN | A | 907 | 7.640 | 4.935 | -84.881 | 1.00 | 53.92 | | C |
| ATOM | 797 | CA | GLY | A | 908 | 6.392 | 3.887 | -81.446 | 1.00 | 51.42 | | C |
| ATOM | 798 | CA | ILE | A | 909 | 5.190 | 7.419 | -80.729 | 1.00 | 49.99 | | C |
| ATOM | 799 | CA | GLY | A | 910 | 3.625 | 7.709 | -84.178 | 1.00 | 53.48 | | C |
| ATOM | 800 | CA | VAL | A | 911 | 6.356 | 9.702 | -85.970 | 1.00 | 54.05 | | C |
| ATOM | 801 | CA | THR | A | 912 | 7.923 | 8.421 | -89.187 | 1.00 | 57.95 | | C |
| ATOM | 802 | CA | GLN | A | 913 | 11.600 | 7.440 | -89.443 | 1.00 | 59.44 | | C |
| ATOM | 803 | CA | ASN | A | 914 | 12.597 | 10.181 | -91.891 | 1.00 | 61.47 | | C |
| ATOM | 804 | CA | VAL | A | 915 | 11.750 | 12.867 | -89.325 | 1.00 | 58.92 | | C |
| ATOM | 805 | CA | LEU | A | 916 | 14.475 | 11.457 | -87.089 | 1.00 | 58.51 | | C |
| ATOM | 806 | CA | TYR | A | 917 | 17.030 | 10.837 | -89.851 | 1.00 | 59.24 | | C |
| ATOM | 807 | CA | GLU | A | 918 | 16.666 | 14.223 | -91.582 | 1.00 | 65.72 | | C |
| ATOM | 808 | CA | ASN | A | 919 | 17.031 | 15.983 | -88.205 | 1.00 | 62.30 | | C |
| ATOM | 809 | CA | GLN | A | 920 | 19.513 | 13.535 | -86.563 | 1.00 | 60.59 | | C |
| ATOM | 810 | CA | LYS | A | 921 | 22.084 | 16.244 | -85.773 | 1.00 | 64.57 | | C |
| ATOM | 811 | CA | LEU | A | 922 | 19.475 | 18.560 | -84.237 | 1.00 | 64.53 | | C |
| ATOM | 812 | CA | ILE | A | 923 | 17.966 | 15.714 | -82.188 | 1.00 | 61.13 | | C |
| ATOM | 813 | CA | ALA | A | 924 | 21.393 | 14.625 | -80.911 | 1.00 | 60.90 | | C |
| ATOM | 814 | CA | ASN | A | 925 | 22.179 | 18.233 | -79.947 | 1.00 | 62.72 | | C |
| ATOM | 815 | CA | GLN | A | 926 | 18.817 | 18.658 | -78.175 | 1.00 | 62.13 | | C |
| ATOM | 816 | CA | PHE | A | 927 | 19.360 | 15.405 | -76.273 | 1.00 | 60.47 | | C |
| ATOM | 817 | CA | ASN | A | 928 | 22.903 | 16.327 | -75.205 | 1.00 | 61.89 | | C |
| ATOM | 818 | CA | SER | A | 929 | 21.741 | 19.810 | -74.174 | 1.00 | 64.65 | | C |
| ATOM | 819 | CA | ALA | A | 930 | 18.827 | 18.397 | -72.149 | 1.00 | 63.59 | | C |
| ATOM | 820 | CA | ILE | A | 931 | 21.210 | 16.046 | -70.316 | 1.00 | 65.26 | | C |

| | | | | | | | | | | | |
|---|---|---|---|---|---|---|---|---|---|---|---|
| ATOM | 821 | CA | GLY | A | 932 | 23.455 | 19.041 | -69.584 | 1.00 | 69.20 | C |
| ATOM | 822 | CA | LYS | A | 933 | 20.383 | 20.864 | -68.249 | 1.00 | 71.72 | C |
| ATOM | 823 | CA | ILE | A | 934 | 19.795 | 17.940 | -65.876 | 1.00 | 71.45 | C |
| ATOM | 824 | CA | GLN | A | 935 | 23.426 | 18.184 | -64.743 | 1.00 | 79.08 | C |
| ATOM | 825 | CA | ASP | A | 936 | 23.057 | 21.904 | -64.027 | 1.00 | 83.15 | C |
| ATOM | 826 | CA | SER | A | 937 | 19.662 | 21.469 | -62.329 | 1.00 | 81.52 | C |
| ATOM | 827 | CA | LEU | A | 938 | 21.042 | 18.843 | -59.949 | 1.00 | 83.04 | C |
| ATOM | 828 | CA | SER | A | 939 | 24.232 | 20.875 | -59.378 | 1.00 | 88.32 | C |
| ATOM | 829 | CA | SER | A | 940 | 22.187 | 23.984 | -58.533 | 1.00 | 91.09 | C |
| ATOM | 830 | CA | THR | A | 941 | 18.865 | 23.190 | -56.810 | 1.00 | 91.04 | C |
| ATOM | 831 | CA | PRO | A | 942 | 19.499 | 22.035 | -53.208 | 1.00 | 93.09 | C |
| ATOM | 832 | CA | SER | A | 943 | 15.986 | 20.679 | -52.522 | 1.00 | 90.81 | C |
| ATOM | 833 | CA | ALA | A | 944 | 15.796 | 18.512 | -55.658 | 1.00 | 84.45 | C |
| ATOM | 834 | CA | LEU | A | 945 | 16.453 | 15.158 | -53.925 | 1.00 | 75.38 | C |
| ATOM | 835 | CA | GLY | A | 946 | 13.987 | 15.872 | -51.092 | 1.00 | 70.87 | C |
| ATOM | 836 | CA | LYS | A | 947 | 12.142 | 12.567 | -51.608 | 1.00 | 65.97 | C |
| ATOM | 837 | CA | LEU | A | 948 | 15.198 | 10.554 | -50.517 | 1.00 | 63.69 | C |
| ATOM | 838 | CA | GLN | A | 949 | 16.348 | 13.007 | -47.862 | 1.00 | 68.26 | C |
| ATOM | 839 | CA | ASP | A | 950 | 12.885 | 12.969 | -46.280 | 1.00 | 69.08 | C |
| ATOM | 840 | CA | VAL | A | 951 | 13.086 | 9.173 | -45.898 | 1.00 | 64.98 | C |
| ATOM | 841 | CA | VAL | A | 952 | 16.552 | 9.442 | -44.326 | 1.00 | 64.99 | C |
| ATOM | 842 | CA | ASN | A | 953 | 15.320 | 12.199 | -41.987 | 1.00 | 65.20 | C |
| ATOM | 843 | CA | GLN | A | 954 | 12.207 | 10.243 | -40.952 | 1.00 | 66.67 | C |
| ATOM | 844 | CA | ASN | A | 955 | 14.285 | 7.196 | -40.049 | 1.00 | 65.97 | C |
| ATOM | 845 | CA | ALA | A | 956 | 16.834 | 9.332 | -38.182 | 1.00 | 66.50 | C |
| ATOM | 846 | CA | GLN | A | 957 | 14.026 | 11.015 | -36.217 | 1.00 | 67.20 | C |
| ATOM | 847 | CA | ALA | A | 958 | 12.420 | 7.640 | -35.449 | 1.00 | 64.49 | C |
| ATOM | 848 | CA | LEU | A | 959 | 15.699 | 6.300 | -34.057 | 1.00 | 66.10 | C |
| ATOM | 849 | CA | ASN | A | 960 | 16.407 | 9.496 | -32.108 | 1.00 | 69.44 | C |
| ATOM | 850 | CA | THR | A | 961 | 12.893 | 9.411 | -30.615 | 1.00 | 66.09 | C |
| ATOM | 851 | CA | LEU | A | 962 | 13.267 | 5.745 | -29.652 | 1.00 | 62.43 | C |
| ATOM | 852 | CA | VAL | A | 963 | 16.633 | 6.370 | -27.990 | 1.00 | 65.11 | C |
| ATOM | 853 | CA | LYS | A | 964 | 15.381 | 9.514 | -26.182 | 1.00 | 64.53 | C |
| ATOM | 854 | CA | GLN | A | 965 | 12.526 | 7.476 | -24.679 | 1.00 | 63.58 | C |
| ATOM | 855 | CA | LEU | A | 966 | 15.150 | 5.725 | -22.485 | 1.00 | 61.14 | C |
| ATOM | 856 | CA | SER | A | 967 | 15.604 | 9.005 | -20.561 | 1.00 | 64.30 | C |
| ATOM | 857 | CA | SER | A | 968 | 11.949 | 9.015 | -19.483 | 1.00 | 61.49 | C |
| ATOM | 858 | CA | ASN | A | 969 | 11.013 | 7.908 | -15.987 | 1.00 | 63.24 | C |
| ATOM | 859 | CA | PHE | A | 970 | 7.475 | 6.662 | -16.887 | 1.00 | 60.89 | C |
| ATOM | 860 | CA | GLY | A | 971 | 6.584 | 7.203 | -13.221 | 1.00 | 63.51 | C |
| ATOM | 861 | CA | ALA | A | 972 | 9.674 | 5.453 | -11.843 | 1.00 | 63.19 | C |
| ATOM | 862 | CA | ILE | A | 973 | 12.054 | 7.085 | -9.374 | 1.00 | 62.44 | C |
| ATOM | 863 | CA | SER | A | 974 | 14.787 | 7.118 | -12.045 | 1.00 | 63.53 | C |
| ATOM | 864 | CA | SER | A | 975 | 15.185 | 6.544 | -15.773 | 1.00 | 63.12 | C |
| ATOM | 865 | CA | VAL | A | 976 | 18.460 | 4.695 | -15.089 | 1.00 | 64.73 | C |
| ATOM | 866 | CA | LEU | A | 977 | 17.954 | 0.993 | -14.395 | 1.00 | 67.22 | C |
| ATOM | 867 | CA | ASN | A | 978 | 21.372 | 0.673 | -12.730 | 1.00 | 71.47 | C |
| ATOM | 868 | CA | ASP | A | 979 | 20.530 | 3.367 | -10.166 | 1.00 | 70.73 | C |
| ATOM | 869 | CA | ILE | A | 980 | 17.310 | 1.568 | -9.195 | 1.00 | 65.46 | C |
| ATOM | 870 | CA | LEU | A | 981 | 19.129 | -1.747 | -8.824 | 1.00 | 65.42 | C |
| ATOM | 871 | CA | SER | A | 982 | 21.976 | -0.077 | -6.911 | 1.00 | 66.62 | C |
| ATOM | 872 | CA | ARG | A | 983 | 19.588 | 1.626 | -4.479 | 1.00 | 65.41 | C |
| ATOM | 873 | CA | LEU | A | 984 | 16.644 | -0.749 | -4.018 | 1.00 | 65.38 | C |
| ATOM | 874 | CA | ASP | A | 985 | 16.089 | -4.364 | -2.981 | 1.00 | 71.39 | C |
| ATOM | 875 | CA | PRO | A | 986 | 13.746 | -6.608 | -5.128 | 1.00 | 69.88 | C |
| ATOM | 876 | CA | PRO | A | 987 | 10.390 | -5.826 | -3.323 | 1.00 | 72.68 | C |
| ATOM | 877 | CA | GLU | A | 988 | 10.599 | -2.165 | -4.460 | 1.00 | 72.78 | C |
| ATOM | 878 | CA | ALA | A | 989 | 12.966 | -2.556 | -7.378 | 1.00 | 68.16 | C |
| ATOM | 879 | CA | GLU | A | 990 | 10.519 | -4.856 | -9.197 | 1.00 | 69.71 | C |
| ATOM | 880 | CA | VAL | A | 991 | 7.818 | -2.167 | -9.028 | 1.00 | 65.88 | C |

ATOM 881 CA GLN A 992 10.174 0.506 -10.380 1.00 63.96 C
ATOM 882 CA ILE A 993 11.482 -1.839 -13.086 1.00 63.76 C
ATOM 883 CA ASP A 994 7.921 -2.824 -14.099 1.00 65.91 C
ATOM 884 CA ARG A 995 7.177 0.876 -14.634 1.00 63.77 C
ATOM 885 CA LEU A 996 10.299 1.196 -16.798 1.00 59.63 C
ATOM 886 CA ILE A 997 9.463 -2.010 -18.715 1.00 58.74 C
ATOM 887 CA THR A 998 5.932 -0.790 -19.452 1.00 58.13 C
ATOM 888 CA GLY A 999 7.133 2.570 -20.767 1.00 57.79 C
ATOM 889 CA ARG A1000 10.080 1.200 -22.767 1.00 57.46 C
ATOM 890 CA LEU A1001 7.997 -1.637 -24.228 1.00 58.12 C
ATOM 891 CA GLN A1002 5.300 0.833 -25.291 1.00 60.24 C
ATOM 892 CA SER A1003 7.953 3.054 -26.903 1.00 59.19 C
ATOM 893 CA LEU A1004 9.515 0.113 -28.755 1.00 57.49 C
ATOM 894 CA GLN A1005 6.112 -1.122 -29.987 1.00 58.71 C
ATOM 895 CA THR A1006 5.334 2.398 -31.223 1.00 57.56 C
ATOM 896 CA TYR A1007 8.676 2.493 -33.069 1.00 54.66 C
ATOM 897 CA VAL A1008 8.137 -0.921 -34.692 1.00 54.11 C
ATOM 898 CA THR A1009 4.578 -0.030 -35.777 1.00 53.93 C
ATOM 899 CA GLN A1010 5.824 3.213 -37.370 1.00 55.88 C
ATOM 900 CA GLN A1011 8.603 1.289 -39.133 1.00 54.10 C
ATOM 901 CA LEU A1012 6.117 -1.259 -40.494 1.00 52.18 C
ATOM 902 CA ILE A1013 3.867 1.480 -41.896 1.00 52.73 C
ATOM 903 CA ARG A1014 6.840 3.405 -43.320 1.00 57.77 C
ATOM 904 CA ALA A1015 8.251 0.155 -44.750 1.00 54.68 C
ATOM 905 CA ALA A1016 4.919 -0.535 -46.469 1.00 53.33 C
ATOM 906 CA GLU A1017 5.121 2.939 -48.032 1.00 54.93 C
ATOM 907 CA ILE A1018 8.711 2.260 -49.156 1.00 52.79 C
ATOM 908 CA ARG A1019 7.606 -1.121 -50.565 1.00 53.31 C
ATOM 909 CA ALA A1020 4.885 0.591 -52.613 1.00 49.78 C
ATOM 910 CA SER A1021 7.432 3.119 -53.902 1.00 49.86 C
ATOM 911 CA ALA A1022 9.901 0.324 -54.721 1.00 48.83 C
ATOM 912 CA ASN A1023 7.200 -1.607 -56.597 1.00 48.89 C
ATOM 913 CA LEU A1024 6.478 1.543 -58.593 1.00 47.16 C
ATOM 914 CA ALA A1025 10.206 2.007 -59.269 1.00 47.88 C
ATOM 915 CA ALA A1026 10.535 -1.616 -60.424 1.00 47.77 C
ATOM 916 CA THR A1027 7.513 -1.164 -62.702 1.00 48.74 C
ATOM 917 CA LYS A1028 9.052 2.006 -64.157 1.00 47.53 C
ATOM 918 CA MET A1029 12.377 0.194 -64.595 1.00 49.20 C
ATOM 919 CA SER A1030 10.689 -2.567 -66.592 1.00 49.34 C
ATOM 920 CA GLU A1031 8.335 -0.338 -68.623 1.00 48.97 C
ATOM 921 CA CYS A1032 10.406 2.849 -69.087 1.00 49.27 C
ATOM 922 CA VAL A1033 13.954 1.493 -69.423 1.00 47.89 C
ATOM 923 CA LEU A1034 13.522 -2.026 -70.814 1.00 48.55 C
ATOM 924 CA GLY A1035 10.842 -0.909 -73.253 1.00 46.91 C
ATOM 925 CA GLN A1036 8.866 2.025 -74.566 1.00 47.47 C
ATOM 926 CA SER A1037 5.677 2.975 -72.757 1.00 47.79 C
ATOM 927 CA LYS A1038 2.411 4.234 -74.217 1.00 48.84 C
ATOM 928 CA ARG A1039 1.077 5.101 -70.756 1.00 45.97 C
ATOM 929 CA VAL A1040 0.597 8.874 -70.510 1.00 49.01 C
ATOM 930 CA ASP A1041 2.726 10.592 -67.803 1.00 52.27 C
ATOM 931 CA PHE A1042 4.046 7.253 -66.519 1.00 48.33 C
ATOM 932 CA CYS A1043 7.545 7.682 -67.960 1.00 50.61 C
ATOM 933 CA GLY A1044 7.802 11.479 -67.956 1.00 52.00 C
ATOM 934 CA LYS A1045 5.817 14.404 -69.330 1.00 56.66 C
ATOM 935 CA GLY A1046 5.494 14.126 -73.117 1.00 52.06 C
ATOM 936 CA TYR A1047 5.489 11.230 -75.537 1.00 48.34 C
ATOM 937 CA HIS A1048 7.879 8.683 -74.064 1.00 46.29 C
ATOM 938 CA LEU A1049 10.951 7.763 -76.088 1.00 46.75 C
ATOM 939 CA MET A1050 13.309 6.107 -73.585 1.00 50.03 C
ATOM 940 CA SER A1051 14.532 6.334 -70.008 1.00 47.41 C

```
ATOM    941  CA  PHE A1052      17.840   5.929 -68.218  1.00 48.87           C
ATOM    942  CA  PRO A1053      18.249   4.678 -64.629  1.00 50.39           C
ATOM    943  CA  GLN A1054      20.704   6.221 -62.188  1.00 57.09           C
ATOM    944  CA  SER A1055      21.699   4.956 -58.756  1.00 62.02           C
ATOM    945  CA  ALA A1056      20.931   7.185 -55.789  1.00 61.31           C
ATOM    946  CA  PRO A1057      21.255   6.772 -51.985  1.00 63.26           C
ATOM    947  CA  HIS A1058      18.455   4.378 -50.984  1.00 60.83           C
ATOM    948  CA  GLY A1059      16.740   5.021 -54.295  1.00 56.52           C
ATOM    949  CA  VAL A1060      16.639   5.361 -58.048  1.00 50.53           C
ATOM    950  CA  VAL A1061      16.516   8.387 -60.335  1.00 51.40           C
ATOM    951  CA  PHE A1062      15.035   8.015 -63.802  1.00 48.14           C
ATOM    952  CA  LEU A1063      15.958  10.387 -66.603  1.00 48.51           C
ATOM    953  CA  HIS A1064      13.014  10.195 -68.979  1.00 48.25           C
ATOM    954  CA  VAL A1065      13.674  11.215 -72.571  1.00 48.60           C
ATOM    955  CA  THR A1066      10.443  12.405 -74.142  1.00 50.40           C
ATOM    956  CA  TYR A1067       9.399  13.936 -77.435  1.00 50.72           C
ATOM    957  CA  VAL A1068       7.597  17.251 -76.914  1.00 54.30           C
ATOM    958  CA  PRO A1069       5.805  19.159 -79.712  1.00 55.52           C
ATOM    959  CA  ALA A1070       7.161  22.673 -80.075  1.00 62.42           C
ATOM    960  CA  GLN A1071       6.245  25.156 -82.836  1.00 67.76           C
ATOM    961  CA  GLU A1072       2.536  25.631 -83.553  1.00 71.09           C
ATOM    962  CA  LYS A1073       0.741  26.233 -86.867  1.00 67.09           C
ATOM    963  CA  ASN A1074      -2.949  26.464 -87.709  1.00 69.51           C
ATOM    964  CA  PHE A1075      -4.302  24.469 -90.648  1.00 64.10           C
ATOM    965  CA  THR A1076      -7.723  23.948 -92.170  1.00 64.97           C
ATOM    966  CA  THR A1077      -8.910  20.351 -92.034  1.00 62.92           C
ATOM    967  CA  ALA A1078     -11.575  17.995 -93.332  1.00 63.05           C
ATOM    968  CA  PRO A1079     -12.583  14.442 -92.369  1.00 63.06           C
ATOM    969  CA  ALA A1080     -12.536  13.241 -95.994  1.00 68.00           C
ATOM    970  CA  ILE A1081     -11.964  14.415 -99.559  1.00 71.54           C
ATOM    971  CA  CYS A1082     -14.502  13.976-102.377  1.00 82.14           C
ATOM    972  CA  HIS A1083     -12.702  13.229-105.642  1.00 83.68           C
ATOM    973  CA  ASP A1084     -14.782  11.228-108.168  1.00 85.66           C
ATOM    974  CA  GLY A1085     -17.813  10.986-105.900  1.00 86.10           C
ATOM    975  CA  LYS A1086     -16.019   8.492-103.660  1.00 80.38           C
ATOM    976  CA  ALA A1087     -14.989   9.382-100.107  1.00 74.67           C
ATOM    977  CA  HIS A1088     -11.225   9.259 -99.504  1.00 71.73           C
ATOM    978  CA  PHE A1089      -9.896   8.952 -95.951  1.00 65.59           C
ATOM    979  CA  PRO A1090      -6.244   9.181 -94.830  1.00 64.49           C
ATOM    980  CA  ARG A1091      -4.511   5.928 -93.969  1.00 70.89           C
ATOM    981  CA  GLU A1092      -1.992   7.243 -91.434  1.00 67.39           C
ATOM    982  CA  GLY A1093      -2.282  11.018 -91.081  1.00 62.24           C
ATOM    983  CA  VAL A1094      -4.811  13.829 -91.553  1.00 60.29           C
ATOM    984  CA  PHE A1095      -5.851  16.036 -94.445  1.00 63.02           C
ATOM    985  CA  VAL A1096      -4.745  19.661 -94.066  1.00 66.28           C
ATOM    986  CA  SER A1097      -4.903  22.855 -96.052  1.00 73.88           C
ATOM    987  CA  ASN A1098      -2.711  25.947 -95.918  1.00 85.25           C
ATOM    988  CA  GLY A1099      -5.613  27.947 -97.359  1.00 86.68           C
ATOM    989  CA  THR A1100      -5.145  27.080-101.041  1.00 87.69           C
ATOM    990  CA  HIS A1101      -3.771  23.528-101.428  1.00 83.63           C
ATOM    991  CA  TRP A1102      -4.688  20.263 -99.696  1.00 71.58           C
ATOM    992  CA  PHE A1103      -2.047  17.837 -98.402  1.00 69.51           C
ATOM    993  CA  VAL A1104      -1.928  14.735 -96.208  1.00 65.01           C
ATOM    994  CA  THR A1105       0.450  14.777 -93.242  1.00 61.38           C
ATOM    995  CA  GLN A1106       1.413  12.600 -90.287  1.00 59.32           C
ATOM    996  CA  ARG A1107      -0.478  13.647 -87.151  1.00 61.12           C
ATOM    997  CA  ASN A1108       2.426  14.508 -84.839  1.00 58.99           C
ATOM    998  CA  PHE A1109       4.760  16.352 -87.248  1.00 59.21           C
ATOM    999  CA  TYR A1110       3.863  18.716 -90.075  1.00 62.07           C
ATOM   1000  CA  GLU A1111       5.227  17.166 -93.270  1.00 64.53           C
```

```
ATOM   1001  CA  PRO A1112       2.855  17.836 -96.167  1.00 65.88           C
ATOM   1002  CA  GLN A1113       2.703  15.318 -98.992  1.00 69.62           C
ATOM   1003  CA  ILE A1114       0.680  15.237-102.198  1.00 73.49           C
ATOM   1004  CA  ILE A1115      -2.540  13.297-101.652  1.00 73.46           C
ATOM   1005  CA  THR A1116      -2.457  10.072-103.670  1.00 77.43           C
ATOM   1006  CA  THR A1117      -4.193   6.704-103.814  1.00 79.05           C
ATOM   1007  CA  ASP A1118      -1.222   5.110-101.995  1.00 80.38           C
ATOM   1008  CA  ASN A1119      -1.774   6.970 -98.700  1.00 73.36           C
ATOM   1009  CA  THR A1120      -5.603   7.071 -98.634  1.00 71.32           C
ATOM   1010  CA  PHE A1121      -8.426   4.533 -98.721  1.00 68.43           C
ATOM   1011  CA  VAL A1122     -11.949   4.652-100.147  1.00 70.79           C
ATOM   1012  CA  SER A1123     -15.136   3.781 -98.254  1.00 72.09           C
ATOM   1013  CA  GLY A1124     -18.593   4.926 -99.316  1.00 76.45           C
ATOM   1014  CA  ASN A1125     -19.585   8.071-101.165  1.00 84.51           C
ATOM   1015  CA  CYS A1126     -19.567  11.826-100.555  1.00 85.71           C
ATOM   1016  CA  ASP A1127     -23.178  12.032 -99.319  1.00 86.77           C
ATOM   1017  CA  VAL A1128     -22.570  10.587 -95.840  1.00 77.48           C
ATOM   1018  CA  VAL A1129     -19.446  12.292 -94.417  1.00 72.64           C
ATOM   1019  CA  ILE A1130     -20.177  15.606 -92.696  1.00 69.93           C
ATOM   1020  CA  GLY A1131     -17.584  18.207 -93.641  1.00 69.88           C
ATOM   1021  CA  ILE A1132     -16.175  16.472 -96.736  1.00 71.29           C
ATOM   1022  CA  VAL A1133     -14.409  18.849 -99.130  1.00 74.86           C
ATOM   1023  CA  ASN A1134     -13.845  18.742-102.876  1.00 85.96           C
ATOM   1024  CA  ASN A1135     -10.285  18.181-104.132  1.00 81.80           C
ATOM   1025  CA  THR A1136      -8.160  16.108-106.499  1.00 82.91           C
ATOM   1026  CA  VAL A1137      -6.619  12.733-105.625  1.00 81.81           C
ATOM   1027  CA  TYR A1138      -3.719  11.732-107.861  1.00 86.72           C
ATOM   1028  CA  ASP A1139      -3.110   8.210-109.177  1.00 94.53           C
ATOM   1029  CA  PRO A1140       0.612   7.875-110.276  1.00 95.41           C
TER    1030      PRO A1140
ATOM   1031  CA  CYS B  15     -58.121  -2.810  11.405  1.00181.97           C
ATOM   1032  CA  VAL B  16     -61.053  -4.255   9.405  1.00182.67           C
ATOM   1033  CA  ASN B  17     -60.956  -7.468   7.345  1.00186.60           C
ATOM   1034  CA  LEU B  18     -63.034  -7.269   4.154  1.00178.03           C
ATOM   1035  CA  THR B  19     -64.410 -10.686   3.190  1.00179.08           C
ATOM   1036  CA  THR B  20     -67.089  -9.971   0.533  1.00176.80           C
ATOM   1037  CA  ARG B  21     -64.678 -10.527  -2.356  1.00171.31           C
ATOM   1038  CA  THR B  22     -64.831 -12.939  -5.281  1.00167.86           C
ATOM   1039  CA  GLN B  23     -61.689 -15.087  -5.588  1.00166.12           C
ATOM   1040  CA  LEU B  24     -60.517 -15.269  -9.209  1.00163.65           C
ATOM   1041  CA  PRO B  25     -57.044 -16.095 -10.617  1.00158.95           C
ATOM   1042  CA  PRO B  26     -54.620 -13.319 -11.670  1.00149.07           C
ATOM   1043  CA  ALA B  27     -54.927 -11.948 -15.200  1.00137.15           C
ATOM   1044  CA  TYR B  28     -52.024 -11.398 -17.588  1.00131.11           C
ATOM   1045  CA  THR B  29     -51.308  -8.909 -20.382  1.00121.97           C
ATOM   1046  CA  ASN B  30     -48.619  -7.678 -22.797  1.00112.23           C
ATOM   1047  CA  SER B  31     -46.315  -4.819 -21.744  1.00108.73           C
ATOM   1048  CA  PHE B  32     -45.721  -3.693 -25.387  1.00107.10           C
ATOM   1049  CA  THR B  33     -43.208  -0.805 -25.328  1.00103.80           C
ATOM   1050  CA  ARG B  34     -44.198   0.703 -21.961  1.00104.92           C
ATOM   1051  CA  GLY B  35     -41.997   1.155 -18.916  1.00100.52           C
ATOM   1052  CA  VAL B  36     -39.186   3.322 -20.310  1.00 97.01           C
ATOM   1053  CA  TYR B  37     -37.971   6.064 -17.955  1.00 93.22           C
ATOM   1054  CA  TYR B  38     -35.213   8.668 -17.924  1.00 89.41           C
ATOM   1055  CA  PRO B  39     -32.290   6.714 -16.387  1.00 90.74           C
ATOM   1056  CA  ASP B  40     -30.449   9.805 -15.116  1.00 91.64           C
ATOM   1057  CA  LYS B  41     -30.713  13.557 -14.484  1.00 91.14           C
ATOM   1058  CA  VAL B  42     -28.647  14.834 -17.442  1.00 88.10           C
ATOM   1059  CA  PHE B  43     -30.120  16.584 -20.482  1.00 88.32           C
ATOM   1060  CA  ARG B  44     -29.071  15.301 -23.881  1.00 86.75           C
```

| | | | | | | | | | | |
|---|---|---|---|---|---|---|---|---|---|---|
| ATOM | 1061 | CA | SER | B | 45 | -30.744 | 15.886 | -27.212 | 1.00 87.63 | C |
| ATOM | 1062 | CA | SER | B | 46 | -30.820 | 14.581 | -30.810 | 1.00 89.30 | C |
| ATOM | 1063 | CA | VAL | B | 47 | -28.768 | 11.560 | -29.716 | 1.00 87.11 | C |
| ATOM | 1064 | CA | LEU | B | 48 | -28.986 | 7.828 | -29.224 | 1.00 85.40 | C |
| ATOM | 1065 | CA | HIS | B | 49 | -27.775 | 6.838 | -25.757 | 1.00 86.19 | C |
| ATOM | 1066 | CA | SER | B | 50 | -26.950 | 3.287 | -24.677 | 1.00 87.28 | C |
| ATOM | 1067 | CA | THR | B | 51 | -27.620 | 2.639 | -20.992 | 1.00 88.96 | C |
| ATOM | 1068 | CA | GLN | B | 52 | -27.423 | -0.459 | -18.789 | 1.00 92.51 | C |
| ATOM | 1069 | CA | ASP | B | 53 | -30.052 | -0.242 | -16.046 | 1.00 94.66 | C |
| ATOM | 1070 | CA | LEU | B | 54 | -33.104 | -1.987 | -14.606 | 1.00 94.36 | C |
| ATOM | 1071 | CA | PHE | B | 55 | -35.874 | -1.713 | -17.198 | 1.00 93.23 | C |
| ATOM | 1072 | CA | LEU | B | 56 | -39.153 | -3.414 | -17.934 | 1.00 96.12 | C |
| ATOM | 1073 | CA | PRO | B | 57 | -38.442 | -5.778 | -20.874 | 1.00 95.92 | C |
| ATOM | 1074 | CA | PHE | B | 58 | -40.382 | -5.095 | -24.053 | 1.00 96.34 | C |
| ATOM | 1075 | CA | PHE | B | 59 | -43.429 | -7.329 | -24.663 | 1.00102.54 | C |
| ATOM | 1076 | CA | SER | B | 60 | -43.115 | -8.841 | -21.193 | 1.00109.62 | C |
| ATOM | 1077 | CA | ASN | B | 61 | -45.821 | -10.745 | -19.330 | 1.00123.48 | C |
| ATOM | 1078 | CA | VAL | B | 62 | -47.458 | -8.259 | -16.930 | 1.00122.55 | C |
| ATOM | 1079 | CA | THR | B | 63 | -49.670 | -9.462 | -14.083 | 1.00126.36 | C |
| ATOM | 1080 | CA | TRP | B | 64 | -53.069 | -7.731 | -14.170 | 1.00132.43 | C |
| ATOM | 1081 | CA | PHE | B | 65 | -55.282 | -7.184 | -11.103 | 1.00132.40 | C |
| ATOM | 1082 | CA | HIS | B | 66 | -58.732 | -5.647 | -10.741 | 1.00137.73 | C |
| ATOM | 1083 | CA | ARG | B | 78 | -63.477 | -5.714 | -7.147 | 1.00155.05 | C |
| ATOM | 1084 | CA | PHE | B | 79 | -60.814 | -5.191 | -4.443 | 1.00156.46 | C |
| ATOM | 1085 | CA | ASP | B | 80 | -57.849 | -7.291 | -5.559 | 1.00145.43 | C |
| ATOM | 1086 | CA | ASN | B | 81 | -54.625 | -6.433 | -3.683 | 1.00142.38 | C |
| ATOM | 1087 | CA | PRO | B | 82 | -52.451 | -9.125 | -2.028 | 1.00130.96 | C |
| ATOM | 1088 | CA | VAL | B | 83 | -48.807 | -8.693 | -1.133 | 1.00123.86 | C |
| ATOM | 1089 | CA | LEU | B | 84 | -46.528 | -9.147 | -4.145 | 1.00118.84 | C |
| ATOM | 1090 | CA | PRO | B | 85 | -42.767 | -9.745 | -4.389 | 1.00113.48 | C |
| ATOM | 1091 | CA | PHE | B | 86 | -40.422 | -6.842 | -5.094 | 1.00110.36 | C |
| ATOM | 1092 | CA | ASN | B | 87 | -37.184 | -8.032 | -6.674 | 1.00107.88 | C |
| ATOM | 1093 | CA | ASP | B | 88 | -34.905 | -5.614 | -8.569 | 1.00104.22 | C |
| ATOM | 1094 | CA | GLY | B | 89 | -37.753 | -3.128 | -8.945 | 1.00101.82 | C |
| ATOM | 1095 | CA | VAL | B | 90 | -41.312 | -2.975 | -10.177 | 1.00102.88 | C |
| ATOM | 1096 | CA | TYR | B | 91 | -43.415 | -1.307 | -12.847 | 1.00106.02 | C |
| ATOM | 1097 | CA | PHE | B | 92 | -46.895 | -0.279 | -11.767 | 1.00115.18 | C |
| ATOM | 1098 | CA | ALA | B | 93 | -49.698 | 1.136 | -13.876 | 1.00121.61 | C |
| ATOM | 1099 | CA | SER | B | 94 | -53.287 | 2.080 | -13.133 | 1.00130.35 | C |
| ATOM | 1100 | CA | THR | B | 95 | -56.284 | 2.788 | -15.332 | 1.00140.02 | C |
| ATOM | 1101 | CA | GLU | B | 96 | -58.921 | 4.810 | -13.486 | 1.00144.99 | C |
| ATOM | 1102 | CA | LYS | B | 97 | -61.792 | 7.206 | -14.001 | 1.00147.62 | C |
| ATOM | 1103 | CA | SER | B | 98 | -62.641 | 8.219 | -10.409 | 1.00148.10 | C |
| ATOM | 1104 | CA | ASN | B | 99 | -59.386 | 8.375 | -8.301 | 1.00146.68 | C |
| ATOM | 1105 | CA | ILE | B | 100 | -59.531 | 5.010 | -6.515 | 1.00148.18 | C |
| ATOM | 1106 | CA | ILE | B | 101 | -55.747 | 4.496 | -6.032 | 1.00146.44 | C |
| ATOM | 1107 | CA | ARG | B | 102 | -54.319 | 5.552 | -2.689 | 1.00146.06 | C |
| ATOM | 1108 | CA | GLY | B | 103 | -50.760 | 4.334 | -2.399 | 1.00133.79 | C |
| ATOM | 1109 | CA | TRP | B | 104 | -48.263 | 1.608 | -1.627 | 1.00123.10 | C |
| ATOM | 1110 | CA | ILE | B | 105 | -46.592 | -0.236 | 1.253 | 1.00125.29 | C |
| ATOM | 1111 | CA | PHE | B | 106 | -43.022 | -1.468 | 0.795 | 1.00119.02 | C |
| ATOM | 1112 | CA | GLY | B | 107 | -41.102 | -3.633 | 3.214 | 1.00120.67 | C |
| ATOM | 1113 | CA | THR | B | 108 | -39.778 | -7.019 | 4.248 | 1.00123.76 | C |
| ATOM | 1114 | CA | THR | B | 109 | -42.313 | -8.319 | 6.787 | 1.00129.85 | C |
| ATOM | 1115 | CA | LEU | B | 110 | -44.743 | -5.363 | 7.212 | 1.00135.26 | C |
| ATOM | 1116 | CA | ASP | B | 111 | -45.483 | -6.158 | 10.884 | 1.00137.24 | C |
| ATOM | 1117 | CA | SER | B | 112 | -43.369 | -3.611 | 12.891 | 1.00135.74 | C |
| ATOM | 1118 | CA | LYS | B | 113 | -40.422 | -6.056 | 13.129 | 1.00134.06 | C |
| ATOM | 1119 | CA | THR | B | 114 | -38.604 | -4.347 | 10.242 | 1.00131.56 | C |
| ATOM | 1120 | CA | GLN | B | 115 | -38.466 | -0.902 | 8.675 | 1.00129.59 | C |

| | | | | | | | | | | |
|---|---|---|---|---|---|---|---|---|---|---|
| ATOM | 1121 | CA | SER | B | 116 | -41.217 | -0.175 | 6.145 | 1.00128.41 | C |
| ATOM | 1122 | CA | LEU | B | 117 | -42.089 | 2.558 | 3.647 | 1.00123.48 | C |
| ATOM | 1123 | CA | LEU | B | 118 | -45.625 | 3.972 | 3.430 | 1.00128.09 | C |
| ATOM | 1124 | CA | ILE | B | 119 | -46.708 | 6.162 | 0.498 | 1.00126.74 | C |
| ATOM | 1125 | CA | VAL | B | 120 | -50.243 | 7.606 | 0.645 | 1.00134.37 | C |
| ATOM | 1126 | CA | ASN | B | 121 | -51.874 | 10.192 | -1.636 | 1.00140.13 | C |
| ATOM | 1127 | CA | ASN | B | 122 | -54.716 | 11.576 | 0.444 | 1.00148.88 | C |
| ATOM | 1128 | CA | ALA | B | 123 | -57.362 | 14.183 | -0.436 | 1.00150.19 | C |
| ATOM | 1129 | CA | THR | B | 124 | -55.082 | 17.162 | 0.284 | 1.00145.63 | C |
| ATOM | 1130 | CA | ASN | B | 125 | -51.466 | 16.054 | -0.391 | 1.00142.54 | C |
| ATOM | 1131 | CA | VAL | B | 126 | -48.994 | 13.167 | -0.760 | 1.00133.05 | C |
| ATOM | 1132 | CA | VAL | B | 127 | -47.376 | 11.876 | 2.437 | 1.00130.35 | C |
| ATOM | 1133 | CA | ILE | B | 128 | -44.373 | 9.523 | 2.552 | 1.00124.59 | C |
| ATOM | 1134 | CA | LYS | B | 129 | -43.342 | 8.078 | 5.902 | 1.00127.22 | C |
| ATOM | 1135 | CA | VAL | B | 130 | -40.592 | 5.612 | 6.720 | 1.00128.55 | C |
| ATOM | 1136 | CA | CYS | B | 131 | -41.797 | 5.054 | 10.248 | 1.00139.44 | C |
| ATOM | 1137 | CA | GLU | B | 132 | -41.933 | 1.586 | 11.793 | 1.00141.78 | C |
| ATOM | 1138 | CA | PHE | B | 133 | -45.576 | 0.670 | 11.197 | 1.00142.99 | C |
| ATOM | 1139 | CA | GLN | B | 134 | -47.657 | -2.221 | 12.488 | 1.00145.92 | C |
| ATOM | 1140 | CA | PHE | B | 135 | -49.750 | -2.795 | 9.393 | 1.00149.52 | C |
| ATOM | 1141 | CA | CYS | B | 136 | -53.219 | -4.274 | 9.056 | 1.00168.95 | C |
| ATOM | 1142 | CA | ASN | B | 137 | -53.722 | -7.581 | 7.298 | 1.00176.66 | C |
| ATOM | 1143 | CA | ASP | B | 138 | -56.202 | -5.872 | 4.947 | 1.00177.59 | C |
| ATOM | 1144 | CA | PRO | B | 139 | -55.206 | -2.179 | 4.731 | 1.00172.55 | C |
| ATOM | 1145 | CA | PHE | B | 140 | -57.394 | 0.297 | 2.831 | 1.00173.56 | C |
| ATOM | 1146 | CA | LEU | B | 141 | -58.920 | 3.773 | 2.819 | 1.00170.34 | C |
| ATOM | 1147 | CA | GLY | B | 142 | -62.575 | 4.698 | 2.454 | 1.00174.29 | C |
| ATOM | 1148 | CA | VAL | B | 143 | -64.540 | 7.396 | 0.664 | 1.00174.22 | C |
| ATOM | 1149 | CA | CYS | B | 152 | -67.939 | 11.549 | -2.144 | 1.00186.23 | C |
| ATOM | 1150 | CA | MET | B | 153 | -65.602 | 12.538 | 0.697 | 1.00183.76 | C |
| ATOM | 1151 | CA | GLU | B | 154 | -62.747 | 10.729 | 2.406 | 1.00176.73 | C |
| ATOM | 1152 | CA | SER | B | 155 | -63.471 | 9.330 | 5.864 | 1.00177.43 | C |
| ATOM | 1153 | CA | GLU | B | 156 | -61.916 | 6.015 | 6.885 | 1.00172.67 | C |
| ATOM | 1154 | CA | PHE | B | 157 | -58.167 | 5.514 | 7.293 | 1.00166.76 | C |
| ATOM | 1155 | CA | ARG | B | 158 | -57.368 | 1.916 | 8.315 | 1.00169.27 | C |
| ATOM | 1156 | CA | VAL | B | 159 | -53.766 | 1.333 | 7.229 | 1.00160.99 | C |
| ATOM | 1157 | CA | TYR | B | 160 | -51.859 | 0.811 | 10.485 | 1.00153.54 | C |
| ATOM | 1158 | CA | SER | B | 161 | -52.213 | 0.808 | 14.259 | 1.00152.54 | C |
| ATOM | 1159 | CA | SER | B | 162 | -48.924 | 1.306 | 16.103 | 1.00150.15 | C |
| ATOM | 1160 | CA | ALA | B | 163 | -46.397 | 3.732 | 14.627 | 1.00145.78 | C |
| ATOM | 1161 | CA | ASN | B | 164 | -43.134 | 4.801 | 16.267 | 1.00141.60 | C |
| ATOM | 1162 | CA | ASN | B | 165 | -39.391 | 5.030 | 15.459 | 1.00141.83 | C |
| ATOM | 1163 | CA | CYS | B | 166 | -40.010 | 7.449 | 12.581 | 1.00140.76 | C |
| ATOM | 1164 | CA | THR | B | 167 | -36.926 | 7.978 | 10.421 | 1.00131.25 | C |
| ATOM | 1165 | CA | PHE | B | 168 | -38.111 | 9.910 | 7.353 | 1.00121.21 | C |
| ATOM | 1166 | CA | GLU | B | 169 | -41.116 | 11.986 | 6.312 | 1.00119.62 | C |
| ATOM | 1167 | CA | TYR | B | 170 | -41.899 | 13.858 | 3.098 | 1.00122.05 | C |
| ATOM | 1168 | CA | VAL | B | 171 | -45.007 | 15.764 | 1.942 | 1.00124.03 | C |
| ATOM | 1169 | CA | SER | B | 172 | -45.636 | 17.235 | -1.518 | 1.00127.04 | C |
| ATOM | 1170 | CA | GLN | B | 173 | -48.025 | 18.075 | -4.365 | 1.00127.04 | C |
| ATOM | 1171 | CA | PRO | B | 174 | -50.599 | 15.389 | -5.379 | 1.00128.97 | C |
| ATOM | 1172 | CA | PHE | B | 175 | -49.389 | 12.268 | -7.306 | 1.00127.48 | C |
| ATOM | 1173 | CA | ASN | B | 185 | -64.140 | 3.307 | -20.342 | 1.00150.93 | C |
| ATOM | 1174 | CA | PHE | B | 186 | -61.342 | 4.819 | -18.265 | 1.00146.94 | C |
| ATOM | 1175 | CA | LYS | B | 187 | -59.843 | 8.285 | -18.693 | 1.00140.73 | C |
| ATOM | 1176 | CA | ASN | B | 188 | -56.630 | 8.444 | -16.627 | 1.00137.52 | C |
| ATOM | 1177 | CA | LEU | B | 189 | -53.579 | 6.215 | -17.087 | 1.00127.28 | C |
| ATOM | 1178 | CA | ARG | B | 190 | -50.837 | 6.519 | -14.471 | 1.00123.25 | C |
| ATOM | 1179 | CA | GLU | B | 191 | -47.474 | 4.776 | -14.758 | 1.00110.36 | C |
| ATOM | 1180 | CA | PHE | B | 192 | -44.877 | 4.449 | -11.993 | 1.00106.18 | C |

```
ATOM   1181  CA  VAL B 193     -41.468   2.801 -11.714  1.00100.68           C
ATOM   1182  CA  PHE B 194     -40.203   1.872  -8.241  1.00103.64           C
ATOM   1183  CA  LYS B 195     -36.495   1.111  -7.782  1.00101.39           C
ATOM   1184  CA  ASN B 196     -34.457   0.482  -4.614  1.00107.58           C
ATOM   1185  CA  ILE B 197     -30.777   1.072  -5.478  1.00106.79           C
ATOM   1186  CA  ASP B 198     -27.800   2.238  -3.341  1.00108.34           C
ATOM   1187  CA  GLY B 199     -29.873   3.660  -0.530  1.00106.89           C
ATOM   1188  CA  TYR B 200     -32.317   5.546  -2.751  1.00103.41           C
ATOM   1189  CA  PHE B 201     -35.874   4.547  -3.534  1.00105.83           C
ATOM   1190  CA  LYS B 202     -36.496   6.071  -6.945  1.00100.16           C
ATOM   1191  CA  ILE B 203     -40.040   6.836  -8.086  1.00 99.19           C
ATOM   1192  CA  TYR B 204     -40.681   7.649 -11.744  1.00 97.02           C
ATOM   1193  CA  SER B 205     -44.098   8.575 -13.075  1.00101.50           C
ATOM   1194  CA  LYS B 206     -46.121   9.568 -16.123  1.00110.01           C
ATOM   1195  CA  HIS B 207     -49.756  10.689 -16.385  1.00122.62           C
ATOM   1196  CA  THR B 208     -51.574  10.337 -19.717  1.00128.40           C
ATOM   1197  CA  PRO B 209     -55.143  10.168 -21.031  1.00132.81           C
ATOM   1198  CA  ILE B 210     -56.476   6.720 -21.907  1.00136.57           C
ATOM   1199  CA  ASN B 211     -59.514   5.559 -23.861  1.00139.19           C
ATOM   1200  CA  LEU B 212     -59.648   1.763 -23.655  1.00141.98           C
ATOM   1201  CA  VAL B 213     -62.190  -0.114 -21.577  1.00143.55           C
ATOM   1202  CA  ARG B 214     -60.171  -3.235 -20.785  1.00138.82           C
ATOM   1203  CA  ASP B 215     -56.515  -2.794 -21.820  1.00134.24           C
ATOM   1204  CA  LEU B 216     -53.097  -0.993 -21.761  1.00126.89           C
ATOM   1205  CA  PRO B 217     -52.311   1.335 -24.698  1.00122.21           C
ATOM   1206  CA  GLN B 218     -49.816   0.642 -27.446  1.00117.00           C
ATOM   1207  CA  GLY B 219     -47.337   3.385 -28.192  1.00109.01           C
ATOM   1208  CA  PHE B 220     -44.307   4.963 -26.557  1.00100.57           C
ATOM   1209  CA  SER B 221     -43.893   7.518 -23.783  1.00102.85           C
ATOM   1210  CA  ALA B 222     -41.180   8.064 -21.197  1.00100.51           C
ATOM   1211  CA  LEU B 223     -41.711   8.276 -17.446  1.00100.45           C
ATOM   1212  CA  GLU B 224     -40.114  11.284 -15.848  1.00 98.06           C
ATOM   1213  CA  PRO B 225     -38.578  10.857 -12.374  1.00 95.45           C
ATOM   1214  CA  LEU B 226     -40.886  11.970  -9.606  1.00100.78           C
ATOM   1215  CA  VAL B 227     -38.820  11.699  -6.412  1.00103.57           C
ATOM   1216  CA  ASP B 228     -35.573  10.151  -5.134  1.00106.14           C
ATOM   1217  CA  LEU B 229     -36.024   9.136  -1.490  1.00107.62           C
ATOM   1218  CA  PRO B 230     -32.721   8.647   0.415  1.00103.95           C
ATOM   1219  CA  ILE B 231     -34.008   5.677   2.422  1.00110.05           C
ATOM   1220  CA  GLY B 232     -31.894   2.556   2.968  1.00112.14           C
ATOM   1221  CA  ILE B 233     -34.721   0.011   3.244  1.00116.13           C
ATOM   1222  CA  ASN B 234     -34.540  -3.713   2.547  1.00118.06           C
ATOM   1223  CA  ILE B 235     -37.554  -4.076   0.231  1.00115.41           C
ATOM   1224  CA  THR B 236     -38.698  -7.528  -0.884  1.00115.39           C
ATOM   1225  CA  ARG B 237     -42.508  -7.237  -0.676  1.00116.66           C
ATOM   1226  CA  PHE B 238     -45.111  -4.586  -1.446  1.00118.52           C
ATOM   1227  CA  GLN B 239     -48.868  -4.021  -1.265  1.00128.38           C
ATOM   1228  CA  THR B 240     -51.246  -1.625  -3.043  1.00132.01           C
ATOM   1229  CA  LEU B 241     -53.624   0.713  -1.153  1.00141.55           C
ATOM   1230  CA  LEU B 242     -57.052   1.232  -2.769  1.00149.98           C
ATOM   1231  CA  ALA B 243     -60.134   3.280  -1.882  1.00154.34           C
ATOM   1232  CA  TRP B 258     -68.842  -0.992   1.196  1.00168.84           C
ATOM   1233  CA  THR B 259     -68.716  -1.106  -2.608  1.00166.80           C
ATOM   1234  CA  ALA B 260     -66.128  -1.435  -5.364  1.00162.07           C
ATOM   1235  CA  GLY B 261     -65.136   1.271  -7.825  1.00156.50           C
ATOM   1236  CA  ALA B 262     -64.043   1.175 -11.462  1.00151.43           C
ATOM   1237  CA  ALA B 263     -60.248   0.753 -11.572  1.00146.81           C
ATOM   1238  CA  ALA B 264     -57.594  -1.702 -12.716  1.00138.54           C
ATOM   1239  CA  TYR B 265     -53.856  -2.011 -12.214  1.00131.25           C
ATOM   1240  CA  TYR B 266     -50.910  -3.822 -13.774  1.00125.22           C
```

```
ATOM   1241  CA  VAL B 267     -47.662  -4.994 -12.138  1.00116.42           C
ATOM   1242  CA  GLY B 268     -44.496  -5.855 -14.061  1.00108.01           C
ATOM   1243  CA  TYR B 269     -40.917  -6.535 -13.004  1.00103.34           C
ATOM   1244  CA  LEU B 270     -37.651  -4.855 -13.948  1.00 96.75           C
ATOM   1245  CA  GLN B 271     -34.596  -6.731 -15.210  1.00 94.21           C
ATOM   1246  CA  PRO B 272     -30.986  -5.708 -15.937  1.00 92.94           C
ATOM   1247  CA  ARG B 273     -31.150  -4.629 -19.584  1.00 91.90           C
ATOM   1248  CA  THR B 274     -29.368  -2.363 -22.027  1.00 89.47           C
ATOM   1249  CA  PHE B 275     -31.620   0.159 -23.773  1.00 88.63           C
ATOM   1250  CA  LEU B 276     -30.898   2.488 -26.678  1.00 86.37           C
ATOM   1251  CA  LEU B 277     -32.800   5.721 -25.989  1.00 86.05           C
ATOM   1252  CA  LYS B 278     -33.630   8.243 -28.718  1.00 87.42           C
ATOM   1253  CA  TYR B 279     -33.512  11.888 -27.619  1.00 87.17           C
ATOM   1254  CA  ASN B 280     -35.163  14.448 -29.919  1.00 89.21           C
ATOM   1255  CA  GLU B 281     -34.269  18.127 -30.461  1.00 89.95           C
ATOM   1256  CA  ASN B 282     -35.934  19.116 -27.164  1.00 93.06           C
ATOM   1257  CA  GLY B 283     -34.292  16.283 -25.227  1.00 89.46           C
ATOM   1258  CA  THR B 284     -37.473  14.221 -24.935  1.00 88.86           C
ATOM   1259  CA  ILE B 285     -37.186  10.449 -25.254  1.00 89.14           C
ATOM   1260  CA  THR B 286     -39.365   9.517 -28.220  1.00 90.41           C
ATOM   1261  CA  ASP B 287     -38.277   5.938 -28.914  1.00 91.70           C
ATOM   1262  CA  ALA B 288     -36.148   3.147 -27.481  1.00 88.62           C
ATOM   1263  CA  VAL B 289     -34.681  -0.226 -28.384  1.00 87.11           C
ATOM   1264  CA  ASP B 290     -34.569  -3.087 -25.875  1.00 88.99           C
ATOM   1265  CA  CYS B 291     -31.181  -4.564 -26.884  1.00 88.48           C
ATOM   1266  CA  ALA B 292     -32.200  -8.151 -25.968  1.00 86.04           C
ATOM   1267  CA  LEU B 293     -35.807  -8.309 -27.233  1.00 86.00           C
ATOM   1268  CA  ASP B 294     -35.009  -9.881 -30.607  1.00 83.83           C
ATOM   1269  CA  PRO B 295     -32.097 -10.150 -33.117  1.00 79.13           C
ATOM   1270  CA  LEU B 296     -33.109  -6.944 -34.938  1.00 80.17           C
ATOM   1271  CA  SER B 297     -33.150  -4.942 -31.690  1.00 83.15           C
ATOM   1272  CA  GLU B 298     -29.675  -6.262 -30.873  1.00 82.43           C
ATOM   1273  CA  THR B 299     -28.513  -5.223 -34.349  1.00 81.54           C
ATOM   1274  CA  LYS B 300     -29.979  -1.736 -33.847  1.00 80.14           C
ATOM   1275  CA  CYS B 301     -28.115  -1.395 -30.543  1.00 84.23           C
ATOM   1276  CA  THR B 302     -24.860  -2.726 -32.037  1.00 82.45           C
ATOM   1277  CA  LEU B 303     -24.887  -0.195 -34.882  1.00 81.67           C
ATOM   1278  CA  LYS B 304     -26.164   2.607 -32.563  1.00 84.00           C
ATOM   1279  CA  SER B 305     -28.990   3.433 -34.955  1.00 83.56           C
ATOM   1280  CA  PHE B 306     -32.725   2.906 -35.296  1.00 85.79           C
ATOM   1281  CA  THR B 307     -32.276   2.315 -39.037  1.00 86.63           C
ATOM   1282  CA  VAL B 308     -30.376  -0.732 -40.292  1.00 85.31           C
ATOM   1283  CA  GLU B 309     -29.193  -1.041 -43.881  1.00 83.94           C
ATOM   1284  CA  LYS B 310     -29.287  -4.280 -45.867  1.00 79.32           C
ATOM   1285  CA  GLY B 311     -26.369  -6.514 -44.976  1.00 74.39           C
ATOM   1286  CA  ILE B 312     -24.907  -8.919 -42.453  1.00 72.24           C
ATOM   1287  CA  TYR B 313     -23.740  -7.605 -39.091  1.00 75.21           C
ATOM   1288  CA  GLN B 314     -21.932  -9.452 -36.309  1.00 78.39           C
ATOM   1289  CA  THR B 315     -23.913  -8.488 -33.230  1.00 79.78           C
ATOM   1290  CA  SER B 316     -23.288 -10.947 -30.388  1.00 81.01           C
ATOM   1291  CA  ASN B 317     -21.702 -14.170 -29.166  1.00 80.31           C
ATOM   1292  CA  PHE B 318     -23.501 -17.456 -28.536  1.00 85.15           C
ATOM   1293  CA  ARG B 319     -22.381 -19.615 -25.618  1.00 87.73           C
ATOM   1294  CA  VAL B 320     -23.958 -22.690 -24.039  1.00 86.96           C
ATOM   1295  CA  GLN B 321     -24.390 -22.105 -20.358  1.00 89.03           C
ATOM   1296  CA  PRO B 322     -23.461 -24.769 -17.793  1.00 88.74           C
ATOM   1297  CA  THR B 323     -26.303 -26.417 -15.902  1.00 90.63           C
ATOM   1298  CA  GLU B 324     -24.440 -28.434 -13.247  1.00 89.04           C
ATOM   1299  CA  SER B 325     -21.487 -28.022 -10.900  1.00 84.65           C
ATOM   1300  CA  ILE B 326     -19.171 -31.045 -10.866  1.00 83.06           C
```

| | | | | | | | | | | | |
|---|---|---|---|---|---|---|---|---|---|---|---|
| ATOM | 1301 | CA | VAL | B | 327 | -16.859 | -31.131 | -7.868 | 1.00 | 82.81 | C |
| ATOM | 1302 | CA | ARG | B | 328 | -14.241 | -33.881 | -7.513 | 1.00 | 82.23 | C |
| ATOM | 1303 | CA | PHE | B | 329 | -11.703 | -34.161 | -4.692 | 1.00 | 83.43 | C |
| ATOM | 1304 | CA | PRO | B | 330 | -9.665 | -37.133 | -3.345 | 1.00 | 85.32 | C |
| ATOM | 1305 | CA | ASN | B | 331 | -11.706 | -39.444 | -1.104 | 1.00 | 91.00 | C |
| ATOM | 1306 | CA | ILE | B | 332 | -9.644 | -38.838 | 2.030 | 1.00 | 84.95 | C |
| ATOM | 1307 | CA | THR | B | 333 | -11.152 | -38.325 | 5.475 | 1.00 | 85.55 | C |
| ATOM | 1308 | CA | ASN | B | 334 | -8.241 | -37.714 | 7.872 | 1.00 | 81.45 | C |
| ATOM | 1309 | CA | LEU | B | 335 | -8.031 | -34.188 | 9.251 | 1.00 | 79.59 | C |
| ATOM | 1310 | CA | CYS | B | 336 | -4.898 | -32.083 | 9.013 | 1.00 | 80.39 | C |
| ATOM | 1311 | CA | PRO | B | 337 | -2.946 | -31.776 | 12.301 | 1.00 | 77.99 | C |
| ATOM | 1312 | CA | PHE | B | 338 | -3.488 | -28.029 | 12.674 | 1.00 | 76.70 | C |
| ATOM | 1313 | CA | GLY | B | 339 | -4.719 | -28.430 | 16.249 | 1.00 | 80.82 | C |
| ATOM | 1314 | CA | GLU | B | 340 | -1.396 | -30.036 | 17.238 | 1.00 | 80.03 | C |
| ATOM | 1315 | CA | VAL | B | 341 | 0.453 | -26.961 | 15.962 | 1.00 | 72.15 | C |
| ATOM | 1316 | CA | PHE | B | 342 | -1.763 | -24.436 | 17.752 | 1.00 | 69.61 | C |
| ATOM | 1317 | CA | ASN | B | 343 | -2.092 | -26.217 | 21.125 | 1.00 | 81.50 | C |
| ATOM | 1318 | CA | ALA | B | 344 | 1.492 | -27.443 | 21.619 | 1.00 | 76.28 | C |
| ATOM | 1319 | CA | THR | B | 345 | 2.656 | -26.929 | 25.205 | 1.00 | 78.13 | C |
| ATOM | 1320 | CA | ARG | B | 346 | 6.009 | -25.426 | 24.193 | 1.00 | 75.86 | C |
| ATOM | 1321 | CA | PHE | B | 347 | 6.538 | -23.078 | 21.266 | 1.00 | 65.58 | C |
| ATOM | 1322 | CA | ALA | B | 348 | 9.862 | -22.484 | 19.583 | 1.00 | 63.20 | C |
| ATOM | 1323 | CA | SER | B | 349 | 11.871 | -19.293 | 19.745 | 1.00 | 65.11 | C |
| ATOM | 1324 | CA | VAL | B | 350 | 11.596 | -17.164 | 16.594 | 1.00 | 62.46 | C |
| ATOM | 1325 | CA | TYR | B | 351 | 15.323 | -17.603 | 15.832 | 1.00 | 64.35 | C |
| ATOM | 1326 | CA | ALA | B | 352 | 14.794 | -21.403 | 15.816 | 1.00 | 63.78 | C |
| ATOM | 1327 | CA | TRP | B | 353 | 11.169 | -21.584 | 14.651 | 1.00 | 62.43 | C |
| ATOM | 1328 | CA | ASN | B | 354 | 9.402 | -24.917 | 14.227 | 1.00 | 64.05 | C |
| ATOM | 1329 | CA | ARG | B | 355 | 8.064 | -26.156 | 10.904 | 1.00 | 65.32 | C |
| ATOM | 1330 | CA | LYS | B | 356 | 5.359 | -28.852 | 10.753 | 1.00 | 70.69 | C |
| ATOM | 1331 | CA | ARG | B | 357 | 4.968 | -29.991 | 7.153 | 1.00 | 78.67 | C |
| ATOM | 1332 | CA | ILE | B | 358 | 1.279 | -30.799 | 6.546 | 1.00 | 75.09 | C |
| ATOM | 1333 | CA | SER | B | 359 | 0.305 | -33.165 | 3.716 | 1.00 | 78.71 | C |
| ATOM | 1334 | CA | ASN | B | 360 | -2.187 | -35.893 | 2.684 | 1.00 | 80.81 | C |
| ATOM | 1335 | CA | CYS | B | 361 | -5.143 | -34.672 | 4.752 | 1.00 | 79.60 | C |
| ATOM | 1336 | CA | VAL | B | 362 | -8.352 | -32.626 | 4.785 | 1.00 | 77.61 | C |
| ATOM | 1337 | CA | ALA | B | 363 | -7.813 | -29.049 | 5.956 | 1.00 | 74.91 | C |
| ATOM | 1338 | CA | ASP | B | 364 | -10.845 | -27.398 | 7.531 | 1.00 | 73.14 | C |
| ATOM | 1339 | CA | TYR | B | 365 | -9.866 | -23.735 | 7.701 | 1.00 | 70.39 | C |
| ATOM | 1340 | CA | SER | B | 366 | -13.239 | -22.473 | 8.911 | 1.00 | 70.89 | C |
| ATOM | 1341 | CA | VAL | B | 367 | -12.427 | -23.582 | 12.457 | 1.00 | 69.22 | C |
| ATOM | 1342 | CA | LEU | B | 368 | -9.290 | -21.429 | 12.584 | 1.00 | 67.14 | C |
| ATOM | 1343 | CA | TYR | B | 369 | -10.792 | -18.396 | 10.837 | 1.00 | 66.28 | C |
| ATOM | 1344 | CA | ASN | B | 370 | -14.057 | -18.268 | 12.764 | 1.00 | 67.73 | C |
| ATOM | 1345 | CA | SER | B | 371 | -12.302 | -18.753 | 16.101 | 1.00 | 68.58 | C |
| ATOM | 1346 | CA | ALA | B | 372 | -12.493 | -15.921 | 18.614 | 1.00 | 67.34 | C |
| ATOM | 1347 | CA | SER | B | 373 | -9.179 | -16.813 | 20.279 | 1.00 | 68.34 | C |
| ATOM | 1348 | CA | PHE | B | 374 | -7.076 | -15.208 | 17.504 | 1.00 | 65.49 | C |
| ATOM | 1349 | CA | SER | B | 375 | -6.322 | -11.489 | 17.320 | 1.00 | 64.06 | C |
| ATOM | 1350 | CA | THR | B | 376 | -4.270 | -11.386 | 14.114 | 1.00 | 63.10 | C |
| ATOM | 1351 | CA | PHE | B | 377 | -5.383 | -13.480 | 11.135 | 1.00 | 61.04 | C |
| ATOM | 1352 | CA | LYS | B | 378 | -3.939 | -11.927 | 7.984 | 1.00 | 64.58 | C |
| ATOM | 1353 | CA | CYS | B | 379 | -3.853 | -13.807 | 4.684 | 1.00 | 64.26 | C |
| ATOM | 1354 | CA | TYR | B | 380 | -1.589 | -12.938 | 1.751 | 1.00 | 64.46 | C |
| ATOM | 1355 | CA | GLY | B | 381 | -2.085 | -14.402 | -1.709 | 1.00 | 62.46 | C |
| ATOM | 1356 | CA | VAL | B | 382 | -5.441 | -16.015 | -0.846 | 1.00 | 63.82 | C |
| ATOM | 1357 | CA | SER | B | 383 | -8.501 | -14.815 | 0.893 | 1.00 | 66.41 | C |
| ATOM | 1358 | CA | PRO | B | 384 | -9.412 | -16.332 | 4.291 | 1.00 | 66.46 | C |
| ATOM | 1359 | CA | THR | B | 385 | -13.026 | -17.082 | 3.289 | 1.00 | 66.79 | C |
| ATOM | 1360 | CA | LYS | B | 386 | -12.106 | -19.052 | 0.150 | 1.00 | 66.62 | C |

| | | | | | | | | | | | |
|---|---|---|---|---|---|---|---|---|---|---|---|
| ATOM | 1361 | CA | LEU | B | 387 | -9.566 | -21.339 | 1.864 | 1.00 | 68.04 | C |
| ATOM | 1362 | CA | ASN | B | 388 | -11.974 | -24.266 | 2.169 | 1.00 | 70.37 | C |
| ATOM | 1363 | CA | ASP | B | 389 | -12.540 | -24.299 | -1.612 | 1.00 | 72.74 | C |
| ATOM | 1364 | CA | LEU | B | 390 | -8.825 | -24.594 | -2.445 | 1.00 | 69.83 | C |
| ATOM | 1365 | CA | CYS | B | 391 | -6.328 | -27.431 | -2.702 | 1.00 | 76.16 | C |
| ATOM | 1366 | CA | PHE | B | 392 | -2.622 | -27.152 | -2.069 | 1.00 | 74.39 | C |
| ATOM | 1367 | CA | THR | B | 393 | 0.208 | -29.581 | -2.649 | 1.00 | 80.21 | C |
| ATOM | 1368 | CA | ASN | B | 394 | 1.742 | -28.861 | 0.764 | 1.00 | 79.10 | C |
| ATOM | 1369 | CA | VAL | B | 395 | 1.180 | -26.648 | 3.823 | 1.00 | 72.56 | C |
| ATOM | 1370 | CA | TYR | B | 396 | 4.076 | -25.473 | 5.999 | 1.00 | 71.46 | C |
| ATOM | 1371 | CA | ALA | B | 397 | 2.830 | -24.768 | 9.558 | 1.00 | 65.89 | C |
| ATOM | 1372 | CA | ASP | B | 398 | 5.531 | -22.517 | 11.057 | 1.00 | 64.02 | C |
| ATOM | 1373 | CA | SER | B | 399 | 5.346 | -21.659 | 14.749 | 1.00 | 62.46 | C |
| ATOM | 1374 | CA | PHE | B | 400 | 7.171 | -19.294 | 17.118 | 1.00 | 59.92 | C |
| ATOM | 1375 | CA | VAL | B | 401 | 6.760 | -16.743 | 19.947 | 1.00 | 61.91 | C |
| ATOM | 1376 | CA | ILE | B | 402 | 7.405 | -13.035 | 19.330 | 1.00 | 64.43 | C |
| ATOM | 1377 | CA | ARG | B | 403 | 6.814 | -9.698 | 21.033 | 1.00 | 71.56 | C |
| ATOM | 1378 | CA | GLY | B | 404 | 3.497 | -7.962 | 20.297 | 1.00 | 71.22 | C |
| ATOM | 1379 | CA | ASP | B | 405 | 5.070 | -4.956 | 18.544 | 1.00 | 75.94 | C |
| ATOM | 1380 | CA | GLU | B | 406 | 6.835 | -7.381 | 16.222 | 1.00 | 71.44 | C |
| ATOM | 1381 | CA | VAL | B | 407 | 3.587 | -9.072 | 15.095 | 1.00 | 66.66 | C |
| ATOM | 1382 | CA | ARG | B | 408 | 3.382 | -6.561 | 12.227 | 1.00 | 69.03 | C |
| ATOM | 1383 | CA | GLN | B | 409 | 6.804 | -7.737 | 11.006 | 1.00 | 66.85 | C |
| ATOM | 1384 | CA | ILE | B | 410 | 5.465 | -11.203 | 10.085 | 1.00 | 62.80 | C |
| ATOM | 1385 | CA | ALA | B | 411 | 4.392 | -10.004 | 6.634 | 1.00 | 64.47 | C |
| ATOM | 1386 | CA | PRO | B | 412 | 5.857 | -9.777 | 3.114 | 1.00 | 67.51 | C |
| ATOM | 1387 | CA | GLY | B | 413 | 8.019 | -6.748 | 2.486 | 1.00 | 71.73 | C |
| ATOM | 1388 | CA | GLN | B | 414 | 8.816 | -5.906 | 6.116 | 1.00 | 70.28 | C |
| ATOM | 1389 | CA | THR | B | 415 | 12.047 | -4.781 | 7.759 | 1.00 | 73.27 | C |
| ATOM | 1390 | CA | GLY | B | 416 | 13.384 | -4.943 | 11.290 | 1.00 | 72.15 | C |
| ATOM | 1391 | CA | LYS | B | 417 | 14.964 | -7.577 | 13.527 | 1.00 | 69.29 | C |
| ATOM | 1392 | CA | ILE | B | 418 | 12.483 | -10.447 | 13.270 | 1.00 | 65.44 | C |
| ATOM | 1393 | CA | ALA | B | 419 | 12.055 | -9.983 | 9.527 | 1.00 | 66.23 | C |
| ATOM | 1394 | CA | ASP | B | 420 | 15.664 | -9.334 | 8.544 | 1.00 | 70.71 | C |
| ATOM | 1395 | CA | TYR | B | 421 | 17.160 | -11.783 | 11.121 | 1.00 | 66.93 | C |
| ATOM | 1396 | CA | ASN | B | 422 | 14.637 | -14.251 | 12.548 | 1.00 | 64.42 | C |
| ATOM | 1397 | CA | TYR | B | 423 | 11.655 | -15.121 | 10.272 | 1.00 | 61.83 | C |
| ATOM | 1398 | CA | LYS | B | 424 | 11.829 | -13.597 | 6.775 | 1.00 | 64.30 | C |
| ATOM | 1399 | CA | LEU | B | 425 | 8.783 | -14.093 | 4.510 | 1.00 | 64.96 | C |
| ATOM | 1400 | CA | PRO | B | 426 | 9.039 | -13.990 | 0.699 | 1.00 | 69.27 | C |
| ATOM | 1401 | CA | ASP | B | 427 | 7.245 | -11.260 | -1.210 | 1.00 | 76.80 | C |
| ATOM | 1402 | CA | ASP | B | 428 | 5.183 | -13.779 | -3.225 | 1.00 | 76.67 | C |
| ATOM | 1403 | CA | PHE | B | 429 | 4.089 | -15.552 | -0.028 | 1.00 | 67.00 | C |
| ATOM | 1404 | CA | THR | B | 430 | 0.776 | -17.401 | -0.135 | 1.00 | 66.28 | C |
| ATOM | 1405 | CA | GLY | B | 431 | -0.546 | -18.218 | 3.301 | 1.00 | 64.75 | C |
| ATOM | 1406 | CA | CYS | B | 432 | -1.998 | -16.873 | 6.513 | 1.00 | 64.44 | C |
| ATOM | 1407 | CA | VAL | B | 433 | -0.282 | -15.300 | 9.537 | 1.00 | 59.55 | C |
| ATOM | 1408 | CA | ILE | B | 434 | -2.424 | -16.307 | 12.521 | 1.00 | 60.21 | C |
| ATOM | 1409 | CA | ALA | B | 435 | -1.326 | -14.820 | 15.847 | 1.00 | 61.76 | C |
| ATOM | 1410 | CA | TRP | B | 436 | -2.702 | -14.512 | 19.368 | 1.00 | 62.84 | C |
| ATOM | 1411 | CA | ASN | B | 437 | -1.778 | -13.006 | 22.724 | 1.00 | 66.68 | C |
| ATOM | 1412 | CA | SER | B | 438 | -0.319 | -15.717 | 24.969 | 1.00 | 70.65 | C |
| ATOM | 1413 | CA | ASN | B | 439 | 0.493 | -13.704 | 28.125 | 1.00 | 74.93 | C |
| ATOM | 1414 | CA | ASN | B | 440 | -1.415 | -16.229 | 30.254 | 1.00 | 78.20 | C |
| ATOM | 1415 | CA | LEU | B | 441 | 0.802 | -19.053 | 28.918 | 1.00 | 75.23 | C |
| ATOM | 1416 | CA | ASP | B | 442 | 4.246 | -17.585 | 28.227 | 1.00 | 73.98 | C |
| ATOM | 1417 | CA | SER | B | 443 | 4.731 | -15.002 | 31.002 | 1.00 | 76.88 | C |
| ATOM | 1418 | CA | LYS | B | 444 | 6.451 | -15.558 | 34.352 | 1.00 | 80.64 | C |
| ATOM | 1419 | CA | VAL | B | 445 | 6.862 | -13.152 | 37.277 | 1.00 | 84.24 | C |
| ATOM | 1420 | CA | GLY | B | 446 | 10.615 | -12.805 | 37.046 | 1.00 | 84.42 | C |

| | | | | | | | | | | | |
|---|---|---|---|---|---|---|---|---|---|---|---|
| ATOM | 1421 | CA | GLY | B | 447 | 10.516 | -12.965 | 33.262 | 1.00 | 81.33 | C |
| ATOM | 1422 | CA | ASN | B | 448 | 10.374 | -16.021 | 31.014 | 1.00 | 77.84 | C |
| ATOM | 1423 | CA | TYR | B | 449 | 13.842 | -16.119 | 29.448 | 1.00 | 80.63 | C |
| ATOM | 1424 | CA | ASN | B | 450 | 13.329 | -19.299 | 27.407 | 1.00 | 75.00 | C |
| ATOM | 1425 | CA | TYR | B | 451 | 12.032 | -17.463 | 24.327 | 1.00 | 69.30 | C |
| ATOM | 1426 | CA | ARG | B | 452 | 14.914 | -15.878 | 22.407 | 1.00 | 69.24 | C |
| ATOM | 1427 | CA | TYR | B | 453 | 15.420 | -13.944 | 19.177 | 1.00 | 66.33 | C |
| ATOM | 1428 | CA | ARG | B | 454 | 18.418 | -13.200 | 16.987 | 1.00 | 65.98 | C |
| ATOM | 1429 | CA | LEU | B | 455 | 19.454 | -9.571 | 17.328 | 1.00 | 66.62 | C |
| ATOM | 1430 | CA | PHE | B | 456 | 22.599 | -9.486 | 15.134 | 1.00 | 68.83 | C |
| ATOM | 1431 | CA | ARG | B | 457 | 23.728 | -11.053 | 11.836 | 1.00 | 68.93 | C |
| ATOM | 1432 | CA | LYS | B | 458 | 26.160 | -10.231 | 9.020 | 1.00 | 71.88 | C |
| ATOM | 1433 | CA | SER | B | 459 | 23.412 | -10.231 | 6.371 | 1.00 | 70.27 | C |
| ATOM | 1434 | CA | ASN | B | 460 | 19.663 | -10.579 | 6.090 | 1.00 | 69.68 | C |
| ATOM | 1435 | CA | LEU | B | 461 | 17.997 | -13.994 | 6.192 | 1.00 | 67.59 | C |
| ATOM | 1436 | CA | LYS | B | 462 | 16.754 | -15.238 | 2.810 | 1.00 | 68.07 | C |
| ATOM | 1437 | CA | PRO | B | 463 | 13.077 | -16.427 | 2.883 | 1.00 | 66.06 | C |
| ATOM | 1438 | CA | PHE | B | 464 | 12.436 | -19.462 | 5.151 | 1.00 | 64.49 | C |
| ATOM | 1439 | CA | GLU | B | 465 | 16.156 | -19.638 | 6.109 | 1.00 | 69.18 | C |
| ATOM | 1440 | CA | ARG | B | 466 | 16.562 | -20.843 | 9.693 | 1.00 | 67.92 | C |
| ATOM | 1441 | CA | ASP | B | 467 | 19.704 | -19.718 | 11.523 | 1.00 | 70.06 | C |
| ATOM | 1442 | CA | ILE | B | 468 | 20.449 | -21.383 | 14.870 | 1.00 | 69.57 | C |
| ATOM | 1443 | CA | SER | B | 469 | 24.108 | -20.460 | 15.317 | 1.00 | 71.86 | C |
| ATOM | 1444 | CA | THR | B | 470 | 25.562 | -18.571 | 18.296 | 1.00 | 76.45 | C |
| ATOM | 1445 | CA | GLU | B | 471 | 28.664 | -17.229 | 16.583 | 1.00 | 81.76 | C |
| ATOM | 1446 | CA | ILE | B | 472 | 29.885 | -13.926 | 18.030 | 1.00 | 80.62 | C |
| ATOM | 1447 | CA | TYR | B | 473 | 28.914 | -10.990 | 15.822 | 1.00 | 79.72 | C |
| ATOM | 1448 | CA | GLN | B | 474 | 31.738 | -8.711 | 14.685 | 1.00 | 87.82 | C |
| ATOM | 1449 | CA | ALA | B | 475 | 30.497 | -5.124 | 14.650 | 1.00 | 89.27 | C |
| ATOM | 1450 | CA | GLY | B | 476 | 33.796 | -3.443 | 13.828 | 1.00 | 93.67 | C |
| ATOM | 1451 | CA | SER | B | 477 | 36.883 | -4.126 | 11.736 | 1.00 | 96.64 | C |
| ATOM | 1452 | CA | THR | B | 478 | 38.721 | -6.087 | 14.468 | 1.00 | 96.44 | C |
| ATOM | 1453 | CA | PRO | B | 479 | 38.304 | -9.903 | 14.555 | 1.00 | 97.06 | C |
| ATOM | 1454 | CA | CYS | B | 480 | 36.535 | -11.154 | 17.671 | 1.00 | 98.73 | C |
| ATOM | 1455 | CA | ASN | B | 481 | 38.086 | -14.665 | 17.866 | 1.00 | 98.97 | C |
| ATOM | 1456 | CA | GLY | B | 482 | 35.155 | -15.972 | 19.891 | 1.00 | 94.64 | C |
| ATOM | 1457 | CA | VAL | B | 483 | 35.473 | -13.497 | 22.796 | 1.00 | 93.23 | C |
| ATOM | 1458 | CA | GLU | B | 484 | 32.625 | -11.240 | 23.956 | 1.00 | 89.79 | C |
| ATOM | 1459 | CA | GLY | B | 485 | 33.787 | -7.646 | 24.298 | 1.00 | 86.25 | C |
| ATOM | 1460 | CA | PHE | B | 486 | 33.955 | -4.295 | 22.538 | 1.00 | 85.95 | C |
| ATOM | 1461 | CA | ASN | B | 487 | 32.377 | -4.508 | 19.035 | 1.00 | 86.83 | C |
| ATOM | 1462 | CA | CYS | B | 488 | 31.914 | -8.264 | 19.630 | 1.00 | 87.36 | C |
| ATOM | 1463 | CA | TYR | B | 489 | 28.352 | -9.135 | 20.585 | 1.00 | 79.57 | C |
| ATOM | 1464 | CA | PHE | B | 490 | 26.290 | -12.194 | 21.400 | 1.00 | 75.70 | C |
| ATOM | 1465 | CA | PRO | B | 491 | 23.811 | -12.492 | 18.496 | 1.00 | 71.43 | C |
| ATOM | 1466 | CA | LEU | B | 492 | 20.832 | -13.775 | 20.504 | 1.00 | 69.00 | C |
| ATOM | 1467 | CA | GLN | B | 493 | 18.781 | -11.958 | 23.129 | 1.00 | 72.21 | C |
| ATOM | 1468 | CA | SER | B | 494 | 15.840 | -12.964 | 25.296 | 1.00 | 72.49 | C |
| ATOM | 1469 | CA | TYR | B | 495 | 12.431 | -11.368 | 25.711 | 1.00 | 71.28 | C |
| ATOM | 1470 | CA | GLY | B | 496 | 11.739 | -10.748 | 29.369 | 1.00 | 77.68 | C |
| ATOM | 1471 | CA | PHE | B | 497 | 8.060 | -11.704 | 29.339 | 1.00 | 73.35 | C |
| ATOM | 1472 | CA | GLN | B | 498 | 6.363 | -10.436 | 32.512 | 1.00 | 81.04 | C |
| ATOM | 1473 | CA | PRO | B | 499 | 2.606 | -10.325 | 33.298 | 1.00 | 80.26 | C |
| ATOM | 1474 | CA | THR | B | 500 | 2.807 | -6.597 | 34.138 | 1.00 | 83.23 | C |
| ATOM | 1475 | CA | ASN | B | 501 | 4.234 | -5.615 | 30.739 | 1.00 | 78.71 | C |
| ATOM | 1476 | CA | GLY | B | 502 | 2.267 | -3.498 | 28.310 | 1.00 | 77.18 | C |
| ATOM | 1477 | CA | VAL | B | 503 | 0.342 | -5.064 | 25.430 | 1.00 | 76.58 | C |
| ATOM | 1478 | CA | GLY | B | 504 | 3.081 | -4.335 | 22.887 | 1.00 | 74.36 | C |
| ATOM | 1479 | CA | TYR | B | 505 | 5.546 | -6.222 | 25.123 | 1.00 | 75.34 | C |
| ATOM | 1480 | CA | GLN | B | 506 | 3.322 | -9.182 | 25.945 | 1.00 | 73.73 | C |

| | | | | | | | | | | | |
|---|---|---|---|---|---|---|---|---|---|---|---|
| ATOM | 1481 | CA | PRO | B | 507 | 4.107 | -12.514 | 24.247 | 1.00 | 69.51 | C |
| ATOM | 1482 | CA | TYR | B | 508 | 2.209 | -13.469 | 21.124 | 1.00 | 66.31 | C |
| ATOM | 1483 | CA | ARG | B | 509 | 2.256 | -16.950 | 19.656 | 1.00 | 63.76 | C |
| ATOM | 1484 | CA | VAL | B | 510 | 2.376 | -16.908 | 15.863 | 1.00 | 59.66 | C |
| ATOM | 1485 | CA | VAL | B | 511 | 1.479 | -19.719 | 13.478 | 1.00 | 58.96 | C |
| ATOM | 1486 | CA | VAL | B | 512 | 2.151 | -19.033 | 9.813 | 1.00 | 58.60 | C |
| ATOM | 1487 | CA | LEU | B | 513 | 0.707 | -21.290 | 7.141 | 1.00 | 62.15 | C |
| ATOM | 1488 | CA | SER | B | 514 | 2.515 | -21.363 | 3.817 | 1.00 | 67.36 | C |
| ATOM | 1489 | CA | PHE | B | 515 | 0.498 | -22.780 | 0.925 | 1.00 | 69.53 | C |
| ATOM | 1490 | CA | GLU | B | 516 | 2.687 | -24.150 | -1.856 | 1.00 | 82.99 | C |
| ATOM | 1491 | CA | LEU | B | 517 | 1.262 | -24.002 | -5.391 | 1.00 | 83.99 | C |
| ATOM | 1492 | CA | LEU | B | 518 | 3.350 | -25.827 | -8.009 | 1.00 | 89.06 | C |
| ATOM | 1493 | CA | HIS | B | 519 | 2.201 | -28.025 | -10.899 | 1.00 | 94.97 | C |
| ATOM | 1494 | CA | ALA | B | 520 | 2.152 | -31.252 | -8.898 | 1.00 | 89.11 | C |
| ATOM | 1495 | CA | PRO | B | 521 | -0.519 | -33.445 | -7.226 | 1.00 | 84.68 | C |
| ATOM | 1496 | CA | ALA | B | 522 | -2.408 | -31.638 | -4.494 | 1.00 | 80.91 | C |
| ATOM | 1497 | CA | THR | B | 523 | -2.462 | -33.367 | -1.111 | 1.00 | 79.92 | C |
| ATOM | 1498 | CA | VAL | B | 524 | -4.148 | -30.817 | 1.189 | 1.00 | 76.20 | C |
| ATOM | 1499 | CA | CYS | B | 525 | -7.755 | -29.961 | 0.352 | 1.00 | 77.19 | C |
| ATOM | 1500 | CA | GLY | B | 526 | -10.611 | -28.195 | 2.103 | 1.00 | 74.51 | C |
| ATOM | 1501 | CA | PRO | B | 527 | -13.685 | -29.991 | 3.495 | 1.00 | 76.15 | C |
| ATOM | 1502 | CA | LYS | B | 528 | -15.641 | -30.040 | 0.241 | 1.00 | 78.57 | C |
| ATOM | 1503 | CA | LYS | B | 529 | -18.131 | -32.723 | -0.706 | 1.00 | 83.85 | C |
| ATOM | 1504 | CA | SER | B | 530 | -17.646 | -34.446 | -4.041 | 1.00 | 84.44 | C |
| ATOM | 1505 | CA | THR | B | 531 | -20.415 | -34.876 | -6.597 | 1.00 | 85.05 | C |
| ATOM | 1506 | CA | ASN | B | 532 | -21.078 | -37.483 | -9.257 | 1.00 | 87.00 | C |
| ATOM | 1507 | CA | LEU | B | 533 | -19.107 | -37.124 | -12.468 | 1.00 | 85.47 | C |
| ATOM | 1508 | CA | VAL | B | 534 | -21.233 | -35.917 | -15.401 | 1.00 | 85.38 | C |
| ATOM | 1509 | CA | LYS | B | 535 | -19.991 | -36.408 | -18.956 | 1.00 | 84.58 | C |
| ATOM | 1510 | CA | ASN | B | 536 | -21.001 | -35.036 | -22.382 | 1.00 | 85.32 | C |
| ATOM | 1511 | CA | LYS | B | 537 | -22.626 | -31.911 | -20.912 | 1.00 | 86.74 | C |
| ATOM | 1512 | CA | CYS | B | 538 | -21.533 | -28.319 | -20.295 | 1.00 | 89.51 | C |
| ATOM | 1513 | CA | VAL | B | 539 | -20.701 | -28.063 | -16.575 | 1.00 | 85.05 | C |
| ATOM | 1514 | CA | ASN | B | 540 | -18.691 | -25.942 | -14.168 | 1.00 | 84.04 | C |
| ATOM | 1515 | CA | PHE | B | 541 | -16.010 | -28.267 | -12.840 | 1.00 | 82.19 | C |
| ATOM | 1516 | CA | ASN | B | 542 | -13.514 | -28.257 | -9.985 | 1.00 | 82.27 | C |
| ATOM | 1517 | CA | PHE | B | 543 | -10.937 | -31.072 | -10.229 | 1.00 | 81.56 | C |
| ATOM | 1518 | CA | ASN | B | 544 | -8.620 | -30.988 | -7.168 | 1.00 | 83.15 | C |
| ATOM | 1519 | CA | GLY | B | 545 | -8.568 | -27.191 | -7.136 | 1.00 | 82.03 | C |
| ATOM | 1520 | CA | LEU | B | 546 | -8.531 | -26.764 | -10.927 | 1.00 | 82.51 | C |
| ATOM | 1521 | CA | THR | B | 547 | -11.660 | -24.841 | -11.902 | 1.00 | 81.40 | C |
| ATOM | 1522 | CA | GLY | B | 548 | -13.315 | -23.993 | -15.175 | 1.00 | 82.41 | C |
| ATOM | 1523 | CA | THR | B | 549 | -16.267 | -24.486 | -17.481 | 1.00 | 84.54 | C |
| ATOM | 1524 | CA | GLY | B | 550 | -16.472 | -27.142 | -20.165 | 1.00 | 83.96 | C |
| ATOM | 1525 | CA | VAL | B | 551 | -17.662 | -30.496 | -21.452 | 1.00 | 84.59 | C |
| ATOM | 1526 | CA | LEU | B | 552 | -16.090 | -33.637 | -19.992 | 1.00 | 83.49 | C |
| ATOM | 1527 | CA | THR | B | 553 | -15.635 | -36.619 | -22.313 | 1.00 | 85.53 | C |
| ATOM | 1528 | CA | GLU | B | 554 | -13.813 | -39.925 | -22.069 | 1.00 | 90.07 | C |
| ATOM | 1529 | CA | SER | B | 555 | -10.245 | -39.501 | -23.271 | 1.00 | 88.85 | C |
| ATOM | 1530 | CA | ASN | B | 556 | -8.090 | -41.688 | -25.498 | 1.00 | 91.78 | C |
| ATOM | 1531 | CA | LYS | B | 557 | -4.864 | -40.047 | -24.325 | 1.00 | 88.69 | C |
| ATOM | 1532 | CA | LYS | B | 558 | -2.174 | -42.323 | -22.900 | 1.00 | 91.77 | C |
| ATOM | 1533 | CA | PHE | B | 559 | -1.325 | -40.502 | -19.693 | 1.00 | 88.15 | C |
| ATOM | 1534 | CA | LEU | B | 560 | 1.771 | -41.628 | -17.860 | 1.00 | 89.10 | C |
| ATOM | 1535 | CA | PRO | B | 561 | 1.213 | -42.817 | -14.229 | 1.00 | 88.91 | C |
| ATOM | 1536 | CA | PHE | B | 562 | 2.523 | -39.515 | -12.780 | 1.00 | 89.80 | C |
| ATOM | 1537 | CA | GLN | B | 563 | 0.588 | -37.097 | -15.006 | 1.00 | 88.73 | C |
| ATOM | 1538 | CA | GLN | B | 564 | -2.401 | -35.408 | -13.372 | 1.00 | 89.56 | C |
| ATOM | 1539 | CA | PHE | B | 565 | -3.714 | -33.194 | -16.188 | 1.00 | 88.95 | C |
| ATOM | 1540 | CA | GLY | B | 566 | -2.944 | -32.322 | -19.779 | 1.00 | 89.10 | C |

```
ATOM   1541  CA  ARG B 567      -2.458 -29.285 -21.965  1.00 92.69           C
ATOM   1542  CA  ASP B 568      -2.999 -28.494 -25.635  1.00100.85           C
ATOM   1543  CA  ILE B 569      -0.673 -26.886 -28.199  1.00100.56           C
ATOM   1544  CA  ALA B 570      -2.039 -23.567 -26.845  1.00 97.93           C
ATOM   1545  CA  ASP B 571      -0.687 -24.434 -23.335  1.00 97.64           C
ATOM   1546  CA  THR B 572      -4.212 -24.415 -21.885  1.00 97.45           C
ATOM   1547  CA  THR B 573      -5.711 -27.260 -19.862  1.00 91.62           C
ATOM   1548  CA  ASP B 574      -7.448 -29.842 -22.053  1.00 91.16           C
ATOM   1549  CA  ALA B 575      -7.493 -33.063 -20.007  1.00 86.75           C
ATOM   1550  CA  VAL B 576      -7.647 -34.019 -16.350  1.00 84.55           C
ATOM   1551  CA  ARG B 577      -7.179 -37.180 -14.292  1.00 85.03           C
ATOM   1552  CA  ASP B 578     -10.104 -37.629 -11.897  1.00 85.54           C
ATOM   1553  CA  PRO B 579      -8.902 -37.588  -8.249  1.00 85.46           C
ATOM   1554  CA  GLN B 580     -11.050 -40.559  -7.142  1.00 89.16           C
ATOM   1555  CA  THR B 581     -10.952 -42.956 -10.113  1.00 88.40           C
ATOM   1556  CA  LEU B 582      -7.853 -42.778 -12.297  1.00 89.88           C
ATOM   1557  CA  GLU B 583      -9.828 -42.067 -15.465  1.00 89.33           C
ATOM   1558  CA  ILE B 584      -8.491 -39.652 -18.065  1.00 86.89           C
ATOM   1559  CA  LEU B 585     -11.057 -37.078 -19.167  1.00 83.87           C
ATOM   1560  CA  ASP B 586     -10.885 -34.651 -22.070  1.00 87.98           C
ATOM   1561  CA  ILE B 587     -12.110 -31.121 -21.446  1.00 84.92           C
ATOM   1562  CA  THR B 588     -13.558 -29.229 -24.349  1.00 86.24           C
ATOM   1563  CA  PRO B 589     -14.958 -25.697 -24.108  1.00 86.78           C
ATOM   1564  CA  CYS B 590     -18.699 -25.252 -24.077  1.00 90.13           C
ATOM   1565  CA  SER B 591     -20.036 -24.438 -27.545  1.00 88.40           C
ATOM   1566  CA  PHE B 592     -19.594 -20.830 -28.585  1.00 85.84           C
ATOM   1567  CA  GLY B 593     -19.432 -18.745 -31.706  1.00 81.83           C
ATOM   1568  CA  GLY B 594     -20.224 -15.409 -33.210  1.00 78.28           C
ATOM   1569  CA  VAL B 595     -23.825 -14.371 -33.764  1.00 77.47           C
ATOM   1570  CA  SER B 596     -24.517 -12.531 -36.997  1.00 74.98           C
ATOM   1571  CA  VAL B 597     -27.872 -11.059 -37.980  1.00 73.56           C
ATOM   1572  CA  ILE B 598     -28.828 -11.206 -41.655  1.00 74.40           C
ATOM   1573  CA  THR B 599     -31.312  -8.567 -42.563  1.00 78.12           C
ATOM   1574  CA  PRO B 600     -32.908  -6.587 -45.365  1.00 83.10           C
ATOM   1575  CA  GLY B 601     -33.018  -2.907 -44.530  1.00 89.81           C
ATOM   1576  CA  THR B 602     -35.630  -1.434 -42.215  1.00 94.32           C
ATOM   1577  CA  ASN B 603     -37.048   0.400 -45.237  1.00106.26           C
ATOM   1578  CA  THR B 604     -38.258  -2.988 -46.544  1.00 95.74           C
ATOM   1579  CA  SER B 605     -38.833  -5.431 -43.668  1.00 88.96           C
ATOM   1580  CA  ASN B 606     -38.211  -6.114 -39.983  1.00 87.08           C
ATOM   1581  CA  GLN B 607     -37.710  -9.850 -40.618  1.00 84.49           C
ATOM   1582  CA  VAL B 608     -34.241 -11.212 -39.801  1.00 76.95           C
ATOM   1583  CA  ALA B 609     -32.243 -14.440 -39.994  1.00 74.31           C
ATOM   1584  CA  VAL B 610     -29.528 -15.444 -37.523  1.00 73.38           C
ATOM   1585  CA  LEU B 611     -26.229 -17.161 -38.310  1.00 74.49           C
ATOM   1586  CA  TYR B 612     -24.503 -18.979 -35.456  1.00 78.24           C
ATOM   1587  CA  GLN B 613     -20.963 -18.967 -36.808  1.00 77.70           C
ATOM   1588  CA  GLY B 614     -19.155 -22.317 -36.648  1.00 80.37           C
ATOM   1589  CA  VAL B 615     -21.864 -23.853 -34.453  1.00 83.94           C
ATOM   1590  CA  ASN B 616     -23.894 -27.021 -35.019  1.00 92.62           C
ATOM   1591  CA  CYS B 617     -27.689 -26.802 -35.279  1.00 92.75           C
ATOM   1592  CA  THR B 618     -28.021 -29.369 -32.455  1.00 93.49           C
ATOM   1593  CA  GLU B 619     -26.296 -27.088 -29.919  1.00 94.13           C
ATOM   1594  CA  VAL B 620     -28.125 -23.763 -30.407  1.00 91.41           C
ATOM   1595  CA  PRO B 621     -31.298 -24.437 -28.417  1.00 95.29           C
ATOM   1596  CA  ASN B 641     -36.207 -22.991 -41.148  1.00 83.73           C
ATOM   1597  CA  VAL B 642     -33.000 -24.370 -39.619  1.00 81.22           C
ATOM   1598  CA  PHE B 643     -30.261 -24.851 -42.236  1.00 82.05           C
ATOM   1599  CA  GLN B 644     -26.811 -26.239 -41.406  1.00 84.54           C
ATOM   1600  CA  THR B 645     -24.013 -24.922 -43.613  1.00 82.29           C
```

| | | | | | | | | | | | |
|---|---|---|---|---|---|---|---|---|---|---|---|
| ATOM | 1601 | CA | ARG | B | 646 | -20.217 | -24.745 | -43.429 | 1.00 | 81.02 | C |
| ATOM | 1602 | CA | ALA | B | 647 | -20.419 | -21.170 | -42.132 | 1.00 | 76.70 | C |
| ATOM | 1603 | CA | GLY | B | 648 | -22.606 | -22.255 | -39.211 | 1.00 | 80.02 | C |
| ATOM | 1604 | CA | CYS | B | 649 | -26.228 | -22.827 | -38.307 | 1.00 | 81.69 | C |
| ATOM | 1605 | CA | LEU | B | 650 | -28.568 | -20.424 | -40.134 | 1.00 | 76.61 | C |
| ATOM | 1606 | CA | ILE | B | 651 | -32.009 | -20.029 | -38.526 | 1.00 | 75.69 | C |
| ATOM | 1607 | CA | GLY | B | 652 | -34.882 | -18.100 | -40.103 | 1.00 | 78.06 | C |
| ATOM | 1608 | CA | ALA | B | 653 | -33.834 | -18.353 | -43.765 | 1.00 | 80.54 | C |
| ATOM | 1609 | CA | GLU | B | 654 | -35.533 | -20.855 | -46.067 | 1.00 | 83.13 | C |
| ATOM | 1610 | CA | HIS | B | 655 | -33.044 | -22.944 | -48.035 | 1.00 | 84.05 | C |
| ATOM | 1611 | CA | VAL | B | 656 | -33.911 | -23.226 | -51.733 | 1.00 | 84.83 | C |
| ATOM | 1612 | CA | ASN | B | 657 | -32.437 | -25.188 | -54.633 | 1.00 | 91.08 | C |
| ATOM | 1613 | CA | ASN | B | 658 | -32.579 | -22.174 | -56.984 | 1.00 | 86.02 | C |
| ATOM | 1614 | CA | SER | B | 659 | -29.309 | -20.298 | -57.353 | 1.00 | 80.40 | C |
| ATOM | 1615 | CA | TYR | B | 660 | -29.189 | -16.524 | -57.761 | 1.00 | 80.41 | C |
| ATOM | 1616 | CA | GLU | B | 661 | -26.633 | -13.737 | -57.668 | 1.00 | 83.41 | C |
| ATOM | 1617 | CA | CYS | B | 662 | -25.179 | -13.057 | -54.228 | 1.00 | 80.19 | C |
| ATOM | 1618 | CA | ASP | B | 663 | -27.027 | -10.389 | -52.246 | 1.00 | 78.88 | C |
| ATOM | 1619 | CA | ILE | B | 664 | -26.157 | -10.716 | -48.548 | 1.00 | 72.91 | C |
| ATOM | 1620 | CA | PRO | B | 665 | -23.106 | -12.997 | -48.151 | 1.00 | 69.62 | C |
| ATOM | 1621 | CA | ILE | B | 666 | -23.301 | -15.815 | -45.619 | 1.00 | 68.48 | C |
| ATOM | 1622 | CA | GLY | B | 667 | -20.129 | -17.723 | -46.404 | 1.00 | 65.77 | C |
| ATOM | 1623 | CA | ALA | B | 668 | -18.871 | -20.944 | -48.046 | 1.00 | 64.81 | C |
| ATOM | 1624 | CA | GLY | B | 669 | -20.757 | -20.251 | -51.243 | 1.00 | 67.77 | C |
| ATOM | 1625 | CA | ILE | B | 670 | -24.011 | -19.456 | -49.433 | 1.00 | 68.56 | C |
| ATOM | 1626 | CA | CYS | B | 671 | -25.715 | -16.067 | -49.825 | 1.00 | 74.60 | C |
| ATOM | 1627 | CA | ALA | B | 672 | -29.049 | -14.786 | -48.540 | 1.00 | 73.99 | C |
| ATOM | 1628 | CA | SER | B | 673 | -31.655 | -12.399 | -49.895 | 1.00 | 79.91 | C |
| ATOM | 1629 | CA | TYR | B | 674 | -35.150 | -11.089 | -49.216 | 1.00 | 85.08 | C |
| ATOM | 1630 | CA | GLN | B | 675 | -37.651 | -12.270 | -51.841 | 1.00 | 87.90 | C |
| ATOM | 1631 | CA | GLN | B | 690 | -41.999 | -12.469 | -48.982 | 1.00 | 88.96 | C |
| ATOM | 1632 | CA | SER | B | 691 | -39.308 | -13.984 | -46.768 | 1.00 | 85.30 | C |
| ATOM | 1633 | CA | ILE | B | 692 | -35.553 | -14.313 | -46.349 | 1.00 | 81.32 | C |
| ATOM | 1634 | CA | ILE | B | 693 | -34.025 | -17.189 | -48.305 | 1.00 | 79.73 | C |
| ATOM | 1635 | CA | ALA | B | 694 | -30.579 | -18.767 | -48.436 | 1.00 | 77.62 | C |
| ATOM | 1636 | CA | TYR | B | 695 | -29.037 | -20.308 | -51.522 | 1.00 | 75.60 | C |
| ATOM | 1637 | CA | THR | B | 696 | -25.788 | -21.321 | -53.116 | 1.00 | 72.83 | C |
| ATOM | 1638 | CA | MET | B | 697 | -24.811 | -18.277 | -55.143 | 1.00 | 72.90 | C |
| ATOM | 1639 | CA | SER | B | 698 | -24.665 | -18.344 | -58.917 | 1.00 | 73.52 | C |
| ATOM | 1640 | CA | LEU | B | 699 | -21.431 | -17.317 | -60.590 | 1.00 | 66.12 | C |
| ATOM | 1641 | CA | GLY | B | 700 | -23.271 | -16.143 | -63.708 | 1.00 | 70.57 | C |
| ATOM | 1642 | CA | ALA | B | 701 | -25.273 | -17.460 | -66.605 | 1.00 | 73.42 | C |
| ATOM | 1643 | CA | GLU | B | 702 | -23.713 | -20.242 | -68.621 | 1.00 | 76.62 | C |
| ATOM | 1644 | CA | ASN | B | 703 | -22.707 | -19.481 | -72.196 | 1.00 | 73.46 | C |
| ATOM | 1645 | CA | SER | B | 704 | -21.172 | -22.221 | -74.323 | 1.00 | 74.37 | C |
| ATOM | 1646 | CA | VAL | B | 705 | -19.251 | -20.490 | -77.102 | 1.00 | 69.15 | C |
| ATOM | 1647 | CA | ALA | B | 706 | -19.807 | -22.032 | -80.544 | 1.00 | 67.44 | C |
| ATOM | 1648 | CA | TYR | B | 707 | -16.142 | -22.705 | -81.227 | 1.00 | 64.66 | C |
| ATOM | 1649 | CA | SER | B | 708 | -15.038 | -24.340 | -84.461 | 1.00 | 65.99 | C |
| ATOM | 1650 | CA | ASN | B | 709 | -11.898 | -24.265 | -86.572 | 1.00 | 68.89 | C |
| ATOM | 1651 | CA | ASN | B | 710 | -13.449 | -22.157 | -89.356 | 1.00 | 65.24 | C |
| ATOM | 1652 | CA | SER | B | 711 | -16.055 | -19.819 | -87.788 | 1.00 | 62.80 | C |
| ATOM | 1653 | CA | ILE | B | 712 | -15.481 | -16.249 | -86.613 | 1.00 | 59.35 | C |
| ATOM | 1654 | CA | ALA | B | 713 | -17.977 | -13.820 | -85.138 | 1.00 | 58.44 | C |
| ATOM | 1655 | CA | ILE | B | 714 | -17.603 | -10.163 | -86.167 | 1.00 | 59.78 | C |
| ATOM | 1656 | CA | PRO | B | 715 | -19.674 | -7.263 | -84.795 | 1.00 | 59.38 | C |
| ATOM | 1657 | CA | THR | B | 716 | -21.936 | -5.542 | -87.299 | 1.00 | 64.18 | C |
| ATOM | 1658 | CA | ASN | B | 717 | -22.951 | -2.747 | -84.931 | 1.00 | 66.68 | C |
| ATOM | 1659 | CA | PHE | B | 718 | -21.719 | -1.236 | -81.685 | 1.00 | 58.85 | C |
| ATOM | 1660 | CA | THR | B | 719 | -22.791 | 0.792 | -78.688 | 1.00 | 56.40 | C |

| | | | | | | | | | | | |
|---|---|---|---|---|---|---|---|---|---|---|---|
| ATOM | 1661 | CA | ILE | B | 720 | -21.123 | 3.608 | -76.791 | 1.00 | 53.55 | C |
| ATOM | 1662 | CA | SER | B | 721 | -21.029 | 2.872 | -73.070 | 1.00 | 52.03 | C |
| ATOM | 1663 | CA | VAL | B | 722 | -20.259 | 5.202 | -70.172 | 1.00 | 52.43 | C |
| ATOM | 1664 | CA | THR | B | 723 | -19.100 | 3.476 | -66.986 | 1.00 | 54.63 | C |
| ATOM | 1665 | CA | THR | B | 724 | -18.860 | 5.281 | -63.646 | 1.00 | 57.50 | C |
| ATOM | 1666 | CA | GLU | B | 725 | -15.793 | 4.624 | -61.481 | 1.00 | 54.69 | C |
| ATOM | 1667 | CA | ILE | B | 726 | -15.619 | 6.267 | -58.044 | 1.00 | 53.65 | C |
| ATOM | 1668 | CA | LEU | B | 727 | -12.286 | 6.577 | -56.241 | 1.00 | 51.73 | C |
| ATOM | 1669 | CA | PRO | B | 728 | -11.347 | 8.145 | -52.896 | 1.00 | 50.94 | C |
| ATOM | 1670 | CA | VAL | B | 729 | -8.675 | 10.823 | -53.215 | 1.00 | 52.90 | C |
| ATOM | 1671 | CA | SER | B | 730 | -8.403 | 12.430 | -49.798 | 1.00 | 57.69 | C |
| ATOM | 1672 | CA | MET | B | 731 | -9.335 | 11.993 | -46.165 | 1.00 | 61.89 | C |
| ATOM | 1673 | CA | THR | B | 732 | -10.535 | 14.534 | -43.599 | 1.00 | 62.96 | C |
| ATOM | 1674 | CA | LYS | B | 733 | -7.483 | 16.301 | -42.174 | 1.00 | 61.20 | C |
| ATOM | 1675 | CA | THR | B | 734 | -7.957 | 15.852 | -38.478 | 1.00 | 65.32 | C |
| ATOM | 1676 | CA | SER | B | 735 | -5.556 | 17.009 | -35.794 | 1.00 | 70.85 | C |
| ATOM | 1677 | CA | VAL | B | 736 | -5.394 | 16.350 | -32.068 | 1.00 | 71.11 | C |
| ATOM | 1678 | CA | ASP | B | 737 | -3.496 | 18.262 | -29.420 | 1.00 | 75.23 | C |
| ATOM | 1679 | CA | CYS | B | 738 | -2.098 | 15.400 | -27.338 | 1.00 | 72.92 | C |
| ATOM | 1680 | CA | THR | B | 739 | -1.741 | 17.441 | -24.152 | 1.00 | 71.19 | C |
| ATOM | 1681 | CA | MET | B | 740 | -5.189 | 19.008 | -24.612 | 1.00 | 73.17 | C |
| ATOM | 1682 | CA | TYR | B | 741 | -6.830 | 15.625 | -25.197 | 1.00 | 64.17 | C |
| ATOM | 1683 | CA | ILE | B | 742 | -5.095 | 13.834 | -22.329 | 1.00 | 64.94 | C |
| ATOM | 1684 | CA | CYS | B | 743 | -4.561 | 16.630 | -19.794 | 1.00 | 72.11 | C |
| ATOM | 1685 | CA | GLY | B | 744 | -6.967 | 19.458 | -20.706 | 1.00 | 74.67 | C |
| ATOM | 1686 | CA | ASP | B | 745 | -6.425 | 22.689 | -18.702 | 1.00 | 81.74 | C |
| ATOM | 1687 | CA | SER | B | 746 | -4.423 | 20.830 | -16.024 | 1.00 | 73.77 | C |
| ATOM | 1688 | CA | THR | B | 747 | -0.858 | 21.723 | -15.039 | 1.00 | 74.24 | C |
| ATOM | 1689 | CA | GLU | B | 748 | -0.551 | 18.692 | -12.717 | 1.00 | 74.63 | C |
| ATOM | 1690 | CA | CYS | B | 749 | -1.577 | 16.222 | -15.421 | 1.00 | 72.07 | C |
| ATOM | 1691 | CA | SER | B | 750 | 0.697 | 17.904 | -17.988 | 1.00 | 70.82 | C |
| ATOM | 1692 | CA | ASN | B | 751 | 3.651 | 17.635 | -15.599 | 1.00 | 69.05 | C |
| ATOM | 1693 | CA | LEU | B | 752 | 2.876 | 13.953 | -15.012 | 1.00 | 67.20 | C |
| ATOM | 1694 | CA | LEU | B | 753 | 2.575 | 13.586 | -18.813 | 1.00 | 65.66 | C |
| ATOM | 1695 | CA | LEU | B | 754 | 6.133 | 14.880 | -19.193 | 1.00 | 66.73 | C |
| ATOM | 1696 | CA | GLN | B | 755 | 7.395 | 11.599 | -17.664 | 1.00 | 65.68 | C |
| ATOM | 1697 | CA | TYR | B | 756 | 6.377 | 9.814 | -20.897 | 1.00 | 63.96 | C |
| ATOM | 1698 | CA | GLY | B | 757 | 8.832 | 11.855 | -22.968 | 1.00 | 66.30 | C |
| ATOM | 1699 | CA | SER | B | 758 | 8.107 | 12.577 | -26.621 | 1.00 | 68.30 | C |
| ATOM | 1700 | CA | PHE | B | 759 | 4.931 | 10.458 | -27.168 | 1.00 | 67.11 | C |
| ATOM | 1701 | CA | CYS | B | 760 | 2.769 | 13.539 | -27.804 | 1.00 | 70.62 | C |
| ATOM | 1702 | CA | THR | B | 761 | 5.335 | 15.005 | -30.213 | 1.00 | 65.35 | C |
| ATOM | 1703 | CA | GLN | B | 762 | 5.480 | 11.897 | -32.403 | 1.00 | 65.17 | C |
| ATOM | 1704 | CA | LEU | B | 763 | 1.673 | 11.561 | -32.376 | 1.00 | 63.01 | C |
| ATOM | 1705 | CA | ASN | B | 764 | 1.284 | 15.198 | -33.471 | 1.00 | 65.04 | C |
| ATOM | 1706 | CA | ARG | B | 765 | 3.953 | 14.645 | -36.127 | 1.00 | 63.03 | C |
| ATOM | 1707 | CA | ALA | B | 766 | 2.170 | 11.536 | -37.437 | 1.00 | 56.73 | C |
| ATOM | 1708 | CA | LEU | B | 767 | -1.149 | 13.386 | -37.755 | 1.00 | 56.93 | C |
| ATOM | 1709 | CA | THR | B | 768 | 0.575 | 16.364 | -39.413 | 1.00 | 58.71 | C |
| ATOM | 1710 | CA | GLY | B | 769 | 2.182 | 13.966 | -41.886 | 1.00 | 56.79 | C |
| ATOM | 1711 | CA | ILE | B | 770 | -1.230 | 12.470 | -42.672 | 1.00 | 55.33 | C |
| ATOM | 1712 | CA | ALA | B | 771 | -2.720 | 15.947 | -43.302 | 1.00 | 57.03 | C |
| ATOM | 1713 | CA | VAL | B | 772 | 0.185 | 16.886 | -45.595 | 1.00 | 57.66 | C |
| ATOM | 1714 | CA | GLU | B | 773 | -0.226 | 13.570 | -47.420 | 1.00 | 60.02 | C |
| ATOM | 1715 | CA | GLN | B | 774 | -3.933 | 14.303 | -48.009 | 1.00 | 57.29 | C |
| ATOM | 1716 | CA | ASP | B | 775 | -3.071 | 17.639 | -49.607 | 1.00 | 60.15 | C |
| ATOM | 1717 | CA | LYS | B | 776 | -0.392 | 15.933 | -51.716 | 1.00 | 57.78 | C |
| ATOM | 1718 | CA | ASN | B | 777 | -2.922 | 13.276 | -52.790 | 1.00 | 55.06 | C |
| ATOM | 1719 | CA | THR | B | 778 | -5.405 | 15.926 | -53.921 | 1.00 | 54.92 | C |
| ATOM | 1720 | CA | GLN | B | 779 | -2.676 | 17.849 | -55.770 | 1.00 | 58.52 | C |

| | | | | | | | | | | | |
|---|---|---|---|---|---|---|---|---|---|---|---|
| ATOM | 1721 | CA | GLU | B | 780 | -1.419 | 14.722 | -57.553 | 1.00 | 59.41 | C |
| ATOM | 1722 | CA | VAL | B | 781 | -4.894 | 13.657 | -58.697 | 1.00 | 52.24 | C |
| ATOM | 1723 | CA | PHE | B | 782 | -6.246 | 17.002 | -59.871 | 1.00 | 53.23 | C |
| ATOM | 1724 | CA | ALA | B | 783 | -3.355 | 19.419 | -60.543 | 1.00 | 55.53 | C |
| ATOM | 1725 | CA | GLN | B | 784 | -2.076 | 17.552 | -63.597 | 1.00 | 58.20 | C |
| ATOM | 1726 | CA | VAL | B | 785 | -2.694 | 20.366 | -66.105 | 1.00 | 63.84 | C |
| ATOM | 1727 | CA | LYS | B | 786 | -0.299 | 23.290 | -66.556 | 1.00 | 72.81 | C |
| ATOM | 1728 | CA | GLN | B | 787 | -2.887 | 25.718 | -67.988 | 1.00 | 70.33 | C |
| ATOM | 1729 | CA | ILE | B | 788 | -6.547 | 26.436 | -67.245 | 1.00 | 67.19 | C |
| ATOM | 1730 | CA | TYR | B | 789 | -8.352 | 25.532 | -70.470 | 1.00 | 63.05 | C |
| ATOM | 1731 | CA | LYS | B | 790 | -11.794 | 26.853 | -71.414 | 1.00 | 68.95 | C |
| ATOM | 1732 | CA | THR | B | 791 | -14.356 | 25.692 | -73.959 | 1.00 | 71.00 | C |
| ATOM | 1733 | CA | PRO | B | 792 | -15.549 | 28.097 | -76.692 | 1.00 | 74.56 | C |
| ATOM | 1734 | CA | PRO | B | 793 | -18.837 | 30.012 | -76.275 | 1.00 | 77.57 | C |
| ATOM | 1735 | CA | ILE | B | 794 | -20.369 | 28.062 | -79.188 | 1.00 | 77.82 | C |
| ATOM | 1736 | CA | LYS | B | 795 | -20.457 | 24.424 | -78.101 | 1.00 | 74.34 | C |
| ATOM | 1737 | CA | ASP | B | 796 | -20.711 | 22.901 | -81.579 | 1.00 | 72.29 | C |
| ATOM | 1738 | CA | PHE | B | 797 | -18.954 | 19.552 | -81.140 | 1.00 | 63.90 | C |
| ATOM | 1739 | CA | GLY | B | 798 | -20.344 | 17.702 | -84.145 | 1.00 | 60.92 | C |
| ATOM | 1740 | CA | GLY | B | 799 | -23.478 | 16.594 | -82.313 | 1.00 | 58.99 | C |
| ATOM | 1741 | CA | PHE | B | 800 | -21.739 | 15.580 | -79.077 | 1.00 | 58.62 | C |
| ATOM | 1742 | CA | ASN | B | 801 | -23.409 | 17.195 | -76.058 | 1.00 | 65.85 | C |
| ATOM | 1743 | CA | PHE | B | 802 | -21.254 | 17.860 | -72.982 | 1.00 | 60.61 | C |
| ATOM | 1744 | CA | SER | B | 803 | -23.670 | 20.026 | -70.936 | 1.00 | 66.72 | C |
| ATOM | 1745 | CA | GLN | B | 804 | -23.933 | 17.324 | -68.252 | 1.00 | 68.48 | C |
| ATOM | 1746 | CA | ILE | B | 805 | -20.202 | 17.292 | -67.559 | 1.00 | 62.57 | C |
| ATOM | 1747 | CA | LEU | B | 806 | -19.272 | 20.976 | -68.320 | 1.00 | 64.88 | C |
| ATOM | 1748 | CA | PRO | B | 807 | -19.892 | 23.787 | -65.787 | 1.00 | 73.18 | C |
| ATOM | 1749 | CA | ASP | B | 808 | -23.193 | 25.692 | -65.920 | 1.00 | 86.76 | C |
| ATOM | 1750 | CA | PRO | B | 809 | -22.449 | 29.279 | -64.690 | 1.00 | 90.63 | C |
| ATOM | 1751 | CA | SER | B | 810 | -26.120 | 30.028 | -63.820 | 1.00 | 94.03 | C |
| ATOM | 1752 | CA | LYS | B | 811 | -25.368 | 29.297 | -60.132 | 1.00 | 91.70 | C |
| ATOM | 1753 | CA | PRO | B | 812 | -22.717 | 30.763 | -57.725 | 1.00 | 94.84 | C |
| ATOM | 1754 | CA | SER | B | 813 | -20.513 | 27.635 | -57.589 | 1.00 | 90.81 | C |
| ATOM | 1755 | CA | LYS | B | 814 | -19.414 | 26.764 | -61.118 | 1.00 | 86.21 | C |
| ATOM | 1756 | CA | ARG | B | 815 | -19.994 | 23.028 | -60.888 | 1.00 | 79.59 | C |
| ATOM | 1757 | CA | SER | B | 816 | -21.572 | 20.792 | -63.497 | 1.00 | 72.91 | C |
| ATOM | 1758 | CA | PRO | B | 817 | -24.984 | 19.056 | -62.880 | 1.00 | 71.30 | C |
| ATOM | 1759 | CA | ILE | B | 818 | -23.437 | 15.566 | -62.400 | 1.00 | 70.92 | C |
| ATOM | 1760 | CA | GLU | B | 819 | -20.962 | 17.089 | -59.923 | 1.00 | 70.58 | C |
| ATOM | 1761 | CA | ASP | B | 820 | -23.920 | 18.718 | -58.134 | 1.00 | 75.29 | C |
| ATOM | 1762 | CA | LEU | B | 821 | -25.480 | 15.256 | -57.853 | 1.00 | 74.85 | C |
| ATOM | 1763 | CA | LEU | B | 822 | -22.238 | 13.849 | -56.445 | 1.00 | 72.08 | C |
| ATOM | 1764 | CA | PHE | B | 823 | -21.947 | 16.619 | -53.852 | 1.00 | 73.25 | C |
| ATOM | 1765 | CA | ASN | B | 824 | -25.621 | 16.427 | -52.826 | 1.00 | 76.22 | C |
| ATOM | 1766 | CA | LYS | B | 825 | -25.507 | 12.644 | -52.330 | 1.00 | 80.77 | C |
| ATOM | 1767 | CA | VAL | B | 826 | -22.331 | 12.788 | -50.216 | 1.00 | 76.73 | C |
| ATOM | 1768 | CA | LEU | B | 849 | -21.506 | 16.923 | -34.371 | 1.00107.25 | | C |
| ATOM | 1769 | CA | ILE | B | 850 | -18.490 | 19.043 | -35.348 | 1.00108.98 | | C |
| ATOM | 1770 | CA | CYS | B | 851 | -18.685 | 21.097 | -32.121 | 1.00109.25 | | C |
| ATOM | 1771 | CA | ALA | B | 852 | -18.856 | 18.002 | -29.877 | 1.00106.78 | | C |
| ATOM | 1772 | CA | GLN | B | 853 | -15.702 | 16.618 | -31.490 | 1.00100.50 | | C |
| ATOM | 1773 | CA | LYS | B | 854 | -13.926 | 20.001 | -31.673 | 1.00100.24 | | C |
| ATOM | 1774 | CA | PHE | B | 855 | -14.388 | 20.813 | -27.948 | 1.00100.09 | | C |
| ATOM | 1775 | CA | ASN | B | 856 | -11.893 | 18.105 | -26.832 | 1.00 | 89.22 | C |
| ATOM | 1776 | CA | GLY | B | 857 | -8.885 | 19.485 | -28.719 | 1.00 | 81.46 | C |
| ATOM | 1777 | CA | LEU | B | 858 | -9.765 | 17.721 | -31.990 | 1.00 | 80.28 | C |
| ATOM | 1778 | CA | THR | B | 859 | -9.644 | 20.228 | -34.811 | 1.00 | 71.97 | C |
| ATOM | 1779 | CA | VAL | B | 860 | -10.214 | 19.733 | -38.527 | 1.00 | 64.98 | C |
| ATOM | 1780 | CA | LEU | B | 861 | -8.019 | 21.541 | -41.012 | 1.00 | 62.78 | C |

| | | | | | | | | | | | |
|---|---|---|---|---|---|---|---|---|---|---|---|
| ATOM | 1781 | CA | PRO | B | 862 | -9.589 | 22.799 | -44.248 | 1.00 | 63.39 | C |
| ATOM | 1782 | CA | PRO | B | 863 | -8.286 | 21.277 | -47.497 | 1.00 | 61.27 | C |
| ATOM | 1783 | CA | LEU | B | 864 | -5.744 | 23.359 | -49.379 | 1.00 | 61.42 | C |
| ATOM | 1784 | CA | LEU | B | 865 | -7.728 | 23.040 | -52.601 | 1.00 | 60.14 | C |
| ATOM | 1785 | CA | THR | B | 866 | -11.320 | 24.200 | -52.217 | 1.00 | 62.12 | C |
| ATOM | 1786 | CA | ASP | B | 867 | -14.258 | 22.530 | -53.961 | 1.00 | 67.99 | C |
| ATOM | 1787 | CA | GLU | B | 868 | -14.335 | 25.458 | -56.402 | 1.00 | 68.27 | C |
| ATOM | 1788 | CA | MET | B | 869 | -10.669 | 24.834 | -57.220 | 1.00 | 63.35 | C |
| ATOM | 1789 | CA | ILE | B | 870 | -11.247 | 21.119 | -57.775 | 1.00 | 60.35 | C |
| ATOM | 1790 | CA | ALA | B | 871 | -14.132 | 22.041 | -60.079 | 1.00 | 60.25 | C |
| ATOM | 1791 | CA | GLN | B | 872 | -11.846 | 24.488 | -61.926 | 1.00 | 61.60 | C |
| ATOM | 1792 | CA | TYR | B | 873 | -9.205 | 21.770 | -62.364 | 1.00 | 57.21 | C |
| ATOM | 1793 | CA | THR | B | 874 | -11.766 | 19.290 | -63.694 | 1.00 | 57.63 | C |
| ATOM | 1794 | CA | SER | B | 875 | -13.149 | 22.008 | -65.981 | 1.00 | 57.89 | C |
| ATOM | 1795 | CA | ALA | B | 876 | -9.644 | 22.737 | -67.322 | 1.00 | 56.91 | C |
| ATOM | 1796 | CA | LEU | B | 877 | -8.915 | 19.023 | -67.841 | 1.00 | 54.33 | C |
| ATOM | 1797 | CA | LEU | B | 878 | -12.274 | 18.531 | -69.527 | 1.00 | 55.62 | C |
| ATOM | 1798 | CA | ALA | B | 879 | -11.889 | 21.589 | -71.776 | 1.00 | 55.48 | C |
| ATOM | 1799 | CA | GLY | B | 880 | -8.463 | 20.323 | -72.813 | 1.00 | 54.60 | C |
| ATOM | 1800 | CA | THR | B | 881 | -9.777 | 16.810 | -73.484 | 1.00 | 54.21 | C |
| ATOM | 1801 | CA | ILE | B | 882 | -12.707 | 18.098 | -75.578 | 1.00 | 54.69 | C |
| ATOM | 1802 | CA | THR | B | 883 | -10.861 | 20.739 | -77.608 | 1.00 | 55.17 | C |
| ATOM | 1803 | CA | SER | B | 884 | -7.364 | 19.239 | -77.870 | 1.00 | 53.63 | C |
| ATOM | 1804 | CA | GLY | B | 885 | -7.748 | 15.474 | -77.307 | 1.00 | 52.94 | C |
| ATOM | 1805 | CA | TRP | B | 886 | -4.647 | 14.028 | -75.693 | 1.00 | 51.82 | C |
| ATOM | 1806 | CA | THR | B | 887 | -2.242 | 16.679 | -77.010 | 1.00 | 52.28 | C |
| ATOM | 1807 | CA | PHE | B | 888 | -2.609 | 18.910 | -73.927 | 1.00 | 52.30 | C |
| ATOM | 1808 | CA | GLY | B | 889 | -1.168 | 16.128 | -71.741 | 1.00 | 53.45 | C |
| ATOM | 1809 | CA | ALA | B | 890 | 2.113 | 15.818 | -73.627 | 1.00 | 53.26 | C |
| ATOM | 1810 | CA | GLY | B | 891 | 2.902 | 19.451 | -74.419 | 1.00 | 54.10 | C |
| ATOM | 1811 | CA | PRO | B | 892 | 1.104 | 22.486 | -75.847 | 1.00 | 53.99 | C |
| ATOM | 1812 | CA | ALA | B | 893 | -2.583 | 21.919 | -76.485 | 1.00 | 53.05 | C |
| ATOM | 1813 | CA | LEU | B | 894 | -3.291 | 21.544 | -80.198 | 1.00 | 53.75 | C |
| ATOM | 1814 | CA | GLN | B | 895 | -6.818 | 22.330 | -81.331 | 1.00 | 57.59 | C |
| ATOM | 1815 | CA | ILE | B | 896 | -8.704 | 19.622 | -83.188 | 1.00 | 55.20 | C |
| ATOM | 1816 | CA | PRO | B | 897 | -12.343 | 19.114 | -84.261 | 1.00 | 55.87 | C |
| ATOM | 1817 | CA | PHE | B | 898 | -13.958 | 16.602 | -81.904 | 1.00 | 55.33 | C |
| ATOM | 1818 | CA | PRO | B | 899 | -15.152 | 14.090 | -84.608 | 1.00 | 56.09 | C |
| ATOM | 1819 | CA | MET | B | 900 | -11.537 | 13.963 | -85.858 | 1.00 | 57.63 | C |
| ATOM | 1820 | CA | GLN | B | 901 | -10.355 | 13.313 | -82.290 | 1.00 | 55.56 | C |
| ATOM | 1821 | CA | MET | B | 902 | -12.937 | 10.521 | -82.060 | 1.00 | 55.27 | C |
| ATOM | 1822 | CA | ALA | B | 903 | -11.567 | 9.186 | -85.365 | 1.00 | 54.71 | C |
| ATOM | 1823 | CA | TYR | B | 904 | -8.128 | 8.969 | -83.775 | 1.00 | 56.60 | C |
| ATOM | 1824 | CA | ARG | B | 905 | -9.588 | 7.169 | -80.758 | 1.00 | 50.35 | C |
| ATOM | 1825 | CA | PHE | B | 906 | -11.370 | 4.724 | -83.083 | 1.00 | 50.88 | C |
| ATOM | 1826 | CA | ASN | B | 907 | -8.069 | 4.108 | -84.873 | 1.00 | 53.38 | C |
| ATOM | 1827 | CA | GLY | B | 908 | -6.530 | 3.553 | -81.441 | 1.00 | 50.72 | C |
| ATOM | 1828 | CA | ILE | B | 909 | -8.991 | 0.754 | -80.713 | 1.00 | 49.33 | C |
| ATOM | 1829 | CA | GLY | B | 910 | -8.461 | -0.742 | -84.167 | 1.00 | 52.63 | C |
| ATOM | 1830 | CA | VAL | B | 911 | -11.543 | 0.632 | -85.972 | 1.00 | 53.61 | C |
| ATOM | 1831 | CA | THR | B | 912 | -11.210 | 2.626 | -89.194 | 1.00 | 57.98 | C |
| ATOM | 1832 | CA | GLN | B | 913 | -12.191 | 6.304 | -89.452 | 1.00 | 59.70 | C |
| ATOM | 1833 | CA | ASN | B | 914 | -15.064 | 5.807 | -91.897 | 1.00 | 62.18 | C |
| ATOM | 1834 | CA | VAL | B | 915 | -16.978 | 3.739 | -89.331 | 1.00 | 58.98 | C |
| ATOM | 1835 | CA | LEU | B | 916 | -17.118 | 6.805 | -87.102 | 1.00 | 58.58 | C |
| ATOM | 1836 | CA | TYR | B | 917 | -17.840 | 9.325 | -89.868 | 1.00 | 59.54 | C |
| ATOM | 1837 | CA | GLU | B | 918 | -20.589 | 7.323 | -91.608 | 1.00 | 66.44 | C |
| ATOM | 1838 | CA | ASN | B | 919 | -22.308 | 6.764 | -88.234 | 1.00 | 62.62 | C |
| ATOM | 1839 | CA | GLN | B | 920 | -21.425 | 10.131 | -86.586 | 1.00 | 60.39 | C |
| ATOM | 1840 | CA | LYS | B | 921 | -25.055 | 11.007 | -85.798 | 1.00 | 64.35 | C |

| | | | | | | | | | | | |
|---|---|---|---|---|---|---|---|---|---|---|---|
| ATOM | 1841 | CA | LEU | B | 922 | -25.764 | 7.585 | -84.276 | 1.00 | 64.50 | C |
| ATOM | 1842 | CA | ILE | B | 923 | -22.551 | 7.690 | -82.216 | 1.00 | 60.78 | C |
| ATOM | 1843 | CA | ALA | B | 924 | -23.320 | 11.200 | -80.934 | 1.00 | 60.69 | C |
| ATOM | 1844 | CA | ASN | B | 925 | -26.841 | 10.080 | -79.979 | 1.00 | 62.79 | C |
| ATOM | 1845 | CA | GLN | B | 926 | -25.534 | 6.954 | -78.207 | 1.00 | 62.27 | C |
| ATOM | 1846 | CA | PHE | B | 927 | -22.983 | 9.045 | -76.307 | 1.00 | 60.32 | C |
| ATOM | 1847 | CA | ASN | B | 928 | -25.545 | 11.660 | -75.236 | 1.00 | 61.84 | C |
| ATOM | 1848 | CA | SER | B | 929 | -27.985 | 8.917 | -74.204 | 1.00 | 64.73 | C |
| ATOM | 1849 | CA | ALA | B | 930 | -25.307 | 7.097 | -72.177 | 1.00 | 63.84 | C |
| ATOM | 1850 | CA | ILE | B | 931 | -24.457 | 10.338 | -70.349 | 1.00 | 65.28 | C |
| ATOM | 1851 | CA | GLY | B | 932 | -28.173 | 10.797 | -69.622 | 1.00 | 69.57 | C |
| ATOM | 1852 | CA | LYS | B | 933 | -28.228 | 7.228 | -68.278 | 1.00 | 72.06 | C |
| ATOM | 1853 | CA | ILE | B | 934 | -25.401 | 8.177 | -65.905 | 1.00 | 72.09 | C |
| ATOM | 1854 | CA | GLN | B | 935 | -27.422 | 11.208 | -64.781 | 1.00 | 79.59 | C |
| ATOM | 1855 | CA | ASP | B | 936 | -30.468 | 9.039 | -64.070 | 1.00 | 83.68 | C |
| ATOM | 1856 | CA | SER | B | 937 | -28.408 | 6.309 | -62.366 | 1.00 | 82.43 | C |
| ATOM | 1857 | CA | LEU | B | 938 | -26.822 | 8.814 | -59.983 | 1.00 | 83.76 | C |
| ATOM | 1858 | CA | SER | B | 939 | -30.175 | 10.567 | -59.421 | 1.00 | 88.89 | C |
| ATOM | 1859 | CA | SER | B | 940 | -31.855 | 7.245 | -58.581 | 1.00 | 93.37 | C |
| ATOM | 1860 | CA | THR | B | 941 | -29.513 | 4.759 | -56.855 | 1.00 | 93.24 | C |
| ATOM | 1861 | CA | PRO | B | 942 | -28.834 | 5.888 | -53.252 | 1.00 | 96.36 | C |
| ATOM | 1862 | CA | SER | B | 943 | -25.905 | 3.521 | -52.557 | 1.00 | 93.44 | C |
| ATOM | 1863 | CA | ALA | B | 944 | -23.923 | 4.434 | -55.689 | 1.00 | 86.90 | C |
| ATOM | 1864 | CA | LEU | B | 945 | -21.348 | 6.675 | -53.953 | 1.00 | 77.10 | C |
| ATOM | 1865 | CA | GLY | B | 946 | -20.751 | 4.180 | -51.118 | 1.00 | 72.70 | C |
| ATOM | 1866 | CA | LYS | B | 947 | -16.966 | 4.216 | -51.634 | 1.00 | 67.02 | C |
| ATOM | 1867 | CA | LEU | B | 948 | -16.735 | 7.866 | -50.533 | 1.00 | 64.42 | C |
| ATOM | 1868 | CA | GLN | B | 949 | -19.431 | 7.638 | -47.881 | 1.00 | 69.20 | C |
| ATOM | 1869 | CA | ASP | B | 950 | -17.664 | 4.661 | -46.297 | 1.00 | 69.86 | C |
| ATOM | 1870 | CA | VAL | B | 951 | -14.475 | 6.731 | -45.918 | 1.00 | 65.52 | C |
| ATOM | 1871 | CA | VAL | B | 952 | -16.439 | 9.605 | -44.354 | 1.00 | 65.84 | C |
| ATOM | 1872 | CA | ASN | B | 953 | -18.210 | 7.165 | -42.011 | 1.00 | 65.80 | C |
| ATOM | 1873 | CA | GLN | B | 954 | -14.957 | 5.448 | -40.972 | 1.00 | 67.30 | C |
| ATOM | 1874 | CA | ASN | B | 955 | -13.362 | 8.772 | -40.072 | 1.00 | 66.86 | C |
| ATOM | 1875 | CA | ALA | B | 956 | -16.488 | 9.912 | -38.210 | 1.00 | 67.45 | C |
| ATOM | 1876 | CA | GLN | B | 957 | -16.545 | 6.640 | -36.239 | 1.00 | 68.35 | C |
| ATOM | 1877 | CA | ALA | B | 958 | -12.819 | 6.935 | -35.470 | 1.00 | 65.97 | C |
| ATOM | 1878 | CA | LEU | B | 959 | -13.292 | 10.453 | -34.095 | 1.00 | 66.93 | C |
| ATOM | 1879 | CA | ASN | B | 960 | -16.417 | 9.493 | -32.143 | 1.00 | 69.94 | C |
| ATOM | 1880 | CA | THR | B | 961 | -14.608 | 6.489 | -30.630 | 1.00 | 66.14 | C |
| ATOM | 1881 | CA | LEU | B | 962 | -11.595 | 8.617 | -29.678 | 1.00 | 62.82 | C |
| ATOM | 1882 | CA | VAL | B | 963 | -13.781 | 11.263 | -28.040 | 1.00 | 66.18 | C |
| ATOM | 1883 | CA | LYS | B | 964 | -15.926 | 8.645 | -26.222 | 1.00 | 65.07 | C |
| ATOM | 1884 | CA | GLN | B | 965 | -12.762 | 7.147 | -24.699 | 1.00 | 63.77 | C |
| ATOM | 1885 | CA | LEU | B | 966 | -12.533 | 10.288 | -22.496 | 1.00 | 61.21 | C |
| ATOM | 1886 | CA | SER | B | 967 | -15.605 | 9.050 | -20.576 | 1.00 | 64.34 | C |
| ATOM | 1887 | CA | SER | B | 968 | -13.797 | 5.867 | -19.513 | 1.00 | 62.25 | C |
| ATOM | 1888 | CA | ASN | B | 969 | -12.372 | 5.594 | -16.015 | 1.00 | 63.93 | C |
| ATOM | 1889 | CA | PHE | B | 970 | -9.515 | 3.161 | -16.907 | 1.00 | 61.54 | C |
| ATOM | 1890 | CA | GLY | B | 971 | -9.539 | 2.108 | -13.244 | 1.00 | 64.35 | C |
| ATOM | 1891 | CA | ALA | B | 972 | -9.577 | 5.653 | -11.857 | 1.00 | 64.38 | C |
| ATOM | 1892 | CA | ILE | B | 973 | -12.191 | 6.879 | -9.392 | 1.00 | 64.16 | C |
| ATOM | 1893 | CA | SER | B | 974 | -13.587 | 9.242 | -12.055 | 1.00 | 64.84 | C |
| ATOM | 1894 | CA | SER | B | 975 | -13.289 | 9.893 | -15.781 | 1.00 | 64.12 | C |
| ATOM | 1895 | CA | VAL | B | 976 | -13.320 | 13.656 | -15.100 | 1.00 | 65.67 | C |
| ATOM | 1896 | CA | LEU | B | 977 | -9.856 | 15.056 | -14.407 | 1.00 | 67.96 | C |
| ATOM | 1897 | CA | ASN | B | 978 | -11.270 | 18.185 | -12.743 | 1.00 | 72.31 | C |
| ATOM | 1898 | CA | ASP | B | 979 | -13.213 | 16.123 | -10.188 | 1.00 | 71.54 | C |
| ATOM | 1899 | CA | ILE | B | 980 | -10.063 | 14.208 | -9.211 | 1.00 | 66.34 | C |
| ATOM | 1900 | CA | LEU | B | 981 | -8.086 | 17.432 | -8.830 | 1.00 | 66.21 | C |

```
ATOM   1901  CA  SER B 982     -10.953  19.060  -6.910  1.00 67.54           C
ATOM   1902  CA  ARG B 983     -11.231  16.128  -4.491  1.00 66.27           C
ATOM   1903  CA  LEU B 984      -7.696  14.768  -4.023  1.00 66.53           C
ATOM   1904  CA  ASP B 985      -4.291  16.094  -2.967  1.00 72.78           C
ATOM   1905  CA  PRO B 986      -1.176  15.206  -5.120  1.00 71.83           C
ATOM   1906  CA  PRO B 987      -0.178  11.904  -3.321  1.00 74.45           C
ATOM   1907  CA  GLU B 988      -3.448  10.257  -4.455  1.00 74.29           C
ATOM   1908  CA  ALA B 989      -4.326  12.502  -7.364  1.00 69.76           C
ATOM   1909  CA  GLU B 990      -1.111  11.569  -9.198  1.00 71.42           C
ATOM   1910  CA  VAL B 991      -2.049   7.872  -9.033  1.00 66.94           C
ATOM   1911  CA  GLN B 992      -5.552   8.546 -10.371  1.00 65.20           C
ATOM   1912  CA  ILE B 993      -4.202  10.861 -13.079  1.00 64.96           C
ATOM   1913  CA  ASP B 994      -1.560   8.282 -14.099  1.00 66.62           C
ATOM   1914  CA  ARG B 995      -4.379   5.772 -14.632  1.00 64.49           C
ATOM   1915  CA  LEU B 996      -6.229   8.303 -16.800  1.00 60.59           C
ATOM   1916  CA  ILE B 997      -3.033   9.209 -18.706  1.00 59.32           C
ATOM   1917  CA  THR B 998      -2.299   5.547 -19.449  1.00 58.46           C
ATOM   1918  CA  GLY B 999      -5.799   4.901 -20.783  1.00 58.29           C
ATOM   1919  CA  ARG B1000      -6.081   8.145 -22.777  1.00 58.44           C
ATOM   1920  CA  LEU B1001      -2.581   7.771 -24.233  1.00 58.27           C
ATOM   1921  CA  GLN B1002      -3.360   4.198 -25.301  1.00 60.56           C
ATOM   1922  CA  SER B1003      -6.607   5.381 -26.917  1.00 59.55           C
ATOM   1923  CA  LEU B1004      -4.839   8.203 -28.769  1.00 57.73           C
ATOM   1924  CA  GLN B1005      -2.085   5.857 -30.001  1.00 58.90           C
ATOM   1925  CA  THR B1006      -4.766   3.440 -31.226  1.00 57.12           C
ATOM   1926  CA  TYR B1007      -6.493   6.293 -33.083  1.00 54.66           C
ATOM   1927  CA  VAL B1008      -3.256   7.493 -34.712  1.00 54.14           C
ATOM   1928  CA  THR B1009      -2.286   3.954 -35.792  1.00 53.72           C
ATOM   1929  CA  GLN B1010      -5.723   3.449 -37.380  1.00 55.75           C
ATOM   1930  CA  GLN B1011      -5.409   6.810 -39.148  1.00 54.00           C
ATOM   1931  CA  LEU B1012      -1.966   5.895 -40.508  1.00 51.60           C
ATOM   1932  CA  ILE B1013      -3.238   2.586 -41.905  1.00 52.77           C
ATOM   1933  CA  ARG B1014      -6.383   4.220 -43.327  1.00 57.87           C
ATOM   1934  CA  ALA B1015      -4.254   7.048 -44.759  1.00 54.29           C
ATOM   1935  CA  ALA B1016      -2.011   4.493 -46.480  1.00 52.97           C
ATOM   1936  CA  GLU B1017      -5.136   2.966 -48.045  1.00 55.04           C
ATOM   1937  CA  ILE B1018      -6.311   6.429 -49.162  1.00 52.84           C
ATOM   1938  CA  ARG B1019      -2.825   7.138 -50.570  1.00 52.98           C
ATOM   1939  CA  ALA B1020      -2.978   3.930 -52.621  1.00 49.45           C
ATOM   1940  CA  SER B1021      -6.437   4.897 -53.907  1.00 49.64           C
ATOM   1941  CA  ALA B1022      -5.230   8.425 -54.724  1.00 48.80           C
ATOM   1942  CA  ASN B1023      -2.214   7.037 -56.598  1.00 48.81           C
ATOM   1943  CA  LEU B1024      -4.581   4.834 -58.595  1.00 47.62           C
ATOM   1944  CA  ALA B1025      -6.847   7.830 -59.273  1.00 48.35           C
ATOM   1945  CA  ALA B1026      -3.865   9.922 -60.423  1.00 48.34           C
ATOM   1946  CA  THR B1027      -2.741   7.081 -62.698  1.00 49.00           C
ATOM   1947  CA  LYS B1028      -6.250   6.838 -64.165  1.00 47.59           C
ATOM   1948  CA  MET B1029      -6.329  10.624 -64.603  1.00 49.35           C
ATOM   1949  CA  SER B1030      -3.084  10.533 -66.586  1.00 49.63           C
ATOM   1950  CA  GLU B1031      -3.838   7.387 -68.626  1.00 48.90           C
ATOM   1951  CA  CYS B1032      -7.631   7.593 -69.093  1.00 49.51           C
ATOM   1952  CA  VAL B1033      -8.234  11.342 -69.420  1.00 47.52           C
ATOM   1953  CA  LEU B1034      -4.966  12.725 -70.807  1.00 48.41           C
ATOM   1954  CA  GLY B1035      -4.594   9.848 -73.250  1.00 47.26           C
ATOM   1955  CA  GLN B1036      -6.143   6.675 -74.580  1.00 47.30           C
ATOM   1956  CA  SER B1037      -5.377   3.436 -72.770  1.00 47.97           C
ATOM   1957  CA  LYS B1038      -4.834  -0.021 -74.233  1.00 48.60           C
ATOM   1958  CA  ARG B1039      -4.939  -1.605 -70.770  1.00 45.83           C
ATOM   1959  CA  VAL B1040      -7.963  -3.907 -70.522  1.00 48.90           C
ATOM   1960  CA  ASP B1041     -10.518  -2.907 -67.821  1.00 52.36           C
```

```
ATOM   1961  CA  PHE B1042      -8.280  -0.099 -66.530  1.00 48.39           C
ATOM   1962  CA  CYS B1043     -10.398   2.717 -67.971  1.00 50.68           C
ATOM   1963  CA  GLY B1044     -13.822   1.053 -67.975  1.00 52.65           C
ATOM   1964  CA  LYS B1045     -15.363  -2.127 -69.349  1.00 57.15           C
ATOM   1965  CA  GLY B1046     -14.933  -2.292 -73.128  1.00 51.85           C
ATOM   1966  CA  TYR B1047     -12.418  -0.860 -75.546  1.00 48.10           C
ATOM   1967  CA  HIS B1048     -11.416   2.484 -74.073  1.00 45.82           C
ATOM   1968  CA  LEU B1049     -12.158   5.604 -76.097  1.00 46.37           C
ATOM   1969  CA  MET B1050     -11.911   8.478 -73.597  1.00 49.79           C
ATOM   1970  CA  SER B1051     -12.718   9.438 -70.024  1.00 47.25           C
ATOM   1971  CA  PHE B1052     -14.035  12.508 -68.247  1.00 49.06           C
ATOM   1972  CA  PRO B1053     -13.149  13.482 -64.659  1.00 50.43           C
ATOM   1973  CA  GLN B1054     -15.708  14.821 -62.202  1.00 57.48           C
ATOM   1974  CA  SER B1055     -15.114  16.305 -58.763  1.00 62.11           C
ATOM   1975  CA  ALA B1056     -16.674  14.524 -55.803  1.00 61.49           C
ATOM   1976  CA  PRO B1057     -16.487  15.003 -51.998  1.00 64.01           C
ATOM   1977  CA  HIS B1058     -13.019  13.766 -50.990  1.00 61.44           C
ATOM   1978  CA  GLY B1059     -12.711  11.970 -54.308  1.00 56.50           C
ATOM   1979  CA  VAL B1060     -12.944  11.727 -58.063  1.00 50.19           C
ATOM   1980  CA  VAL B1061     -15.500  10.112 -60.354  1.00 51.63           C
ATOM   1981  CA  PHE B1062     -14.430   9.016 -63.817  1.00 47.66           C
ATOM   1982  CA  LEU B1063     -16.943   8.644 -66.623  1.00 48.26           C
ATOM   1983  CA  HIS B1064     -15.318   6.179 -69.000  1.00 48.37           C
ATOM   1984  CA  VAL B1065     -16.513   6.236 -72.596  1.00 48.57           C
ATOM   1985  CA  THR B1066     -15.923   2.834 -74.150  1.00 50.19           C
ATOM   1986  CA  TYR B1067     -16.724   1.168 -77.445  1.00 50.97           C
ATOM   1987  CA  VAL B1068     -18.681  -2.057 -76.925  1.00 54.12           C
ATOM   1988  CA  PRO B1069     -19.433  -4.554 -79.732  1.00 55.71           C
ATOM   1989  CA  ALA B1070     -23.153  -5.132 -80.098  1.00 62.81           C
ATOM   1990  CA  GLN B1071     -24.844  -7.167 -82.857  1.00 68.25           C
ATOM   1991  CA  GLU B1072     -23.410 -10.619 -83.577  1.00 71.16           C
ATOM   1992  CA  LYS B1073     -23.051 -12.478 -86.891  1.00 67.65           C
ATOM   1993  CA  ASN B1074     -21.397 -15.790 -87.720  1.00 69.80           C
ATOM   1994  CA  PHE B1075     -19.000 -15.966 -90.663  1.00 64.81           C
ATOM   1995  CA  THR B1076     -16.837 -18.669 -92.180  1.00 65.99           C
ATOM   1996  CA  THR B1077     -13.128 -17.902 -92.044  1.00 63.76           C
ATOM   1997  CA  ALA B1078      -9.757 -19.050 -93.329  1.00 62.79           C
ATOM   1998  CA  PRO B1079      -6.177 -18.153 -92.358  1.00 63.17           C
ATOM   1999  CA  ALA B1080      -5.142 -17.508 -95.978  1.00 68.40           C
ATOM   2000  CA  ILE B1081      -6.427 -17.594 -99.549  1.00 70.99           C
ATOM   2001  CA  CYS B1082      -4.769 -19.561-102.370  1.00 80.77           C
ATOM   2002  CA  HIS B1083      -5.014 -17.626-105.634  1.00 83.29           C
ATOM   2003  CA  ASP B1084      -2.230 -18.425-108.150  1.00 85.67           C
ATOM   2004  CA  GLY B1085      -0.500 -20.926-105.884  1.00 84.31           C
ATOM   2005  CA  LYS B1086       0.753 -18.120-103.643  1.00 79.67           C
ATOM   2006  CA  ALA B1087      -0.534 -17.679-100.090  1.00 74.15           C
ATOM   2007  CA  HIS B1088      -2.335 -14.373 -99.487  1.00 71.39           C
ATOM   2008  CA  PHE B1089      -2.739 -13.063 -95.941  1.00 64.87           C
ATOM   2009  CA  PRO B1090      -4.766 -10.013 -94.834  1.00 63.98           C
ATOM   2010  CA  ARG B1091      -2.812  -6.883 -93.984  1.00 70.77           C
ATOM   2011  CA  GLU B1092      -5.208  -5.352 -91.448  1.00 67.46           C
ATOM   2012  CA  GLY B1093      -8.337  -7.486 -91.098  1.00 61.50           C
ATOM   2013  CA  VAL B1094      -9.504 -11.085 -91.575  1.00 59.80           C
ATOM   2014  CA  PHE B1095     -10.901 -13.095 -94.460  1.00 62.21           C
ATOM   2015  CA  VAL B1096     -14.597 -13.939 -94.074  1.00 65.93           C
ATOM   2016  CA  SER B1097     -17.285 -15.671 -96.063  1.00 73.94           C
ATOM   2017  CA  ASN B1098     -21.060 -15.311 -95.953  1.00 85.51           C
ATOM   2018  CA  GLY B1099     -21.346 -18.824 -97.395  1.00 86.71           C
ATOM   2019  CA  THR B1100     -20.799 -17.998-101.074  1.00 87.56           C
ATOM   2020  CA  HIS B1101     -18.396 -15.042-101.455  1.00 83.41           C
```

```
ATOM   2021  CA  TRP B1102     -15.120 -14.210 -99.701  1.00 72.01           C
ATOM   2022  CA  PHE B1103     -14.340 -10.707 -98.414  1.00 69.63           C
ATOM   2023  CA  VAL B1104     -11.730  -9.044 -96.212  1.00 65.38           C
ATOM   2024  CA  THR B1105     -12.955  -6.997 -93.256  1.00 61.41           C
ATOM   2025  CA  GLN B1106     -11.552  -5.078 -90.298  1.00 59.45           C
ATOM   2026  CA  ARG B1107     -11.521  -7.237 -87.158  1.00 61.41           C
ATOM   2027  CA  ASN B1108     -13.716  -5.139 -84.855  1.00 59.05           C
ATOM   2028  CA  PHE B1109     -16.480  -4.051 -87.268  1.00 59.32           C
ATOM   2029  CA  TYR B1110     -18.078  -6.027 -90.085  1.00 62.51           C
ATOM   2030  CA  GLU B1111     -17.411  -4.079 -93.281  1.00 64.94           C
ATOM   2031  CA  PRO B1112     -16.795  -6.470 -96.174  1.00 66.38           C
ATOM   2032  CA  GLN B1113     -14.535  -5.337 -98.994  1.00 69.48           C
ATOM   2033  CA  ILE B1114     -13.451  -7.040-102.202  1.00 73.47           C
ATOM   2034  CA  ILE B1115     -10.161  -8.860-101.658  1.00 73.72           C
ATOM   2035  CA  THR B1116      -7.412  -7.168-103.673  1.00 77.02           C
ATOM   2036  CA  THR B1117      -3.625  -6.987-103.803  1.00 78.22           C
ATOM   2037  CA  ASP B1118      -3.751  -3.624-101.973  1.00 80.32           C
ATOM   2038  CA  ASN B1119      -5.112  -5.047 -98.696  1.00 73.14           C
ATOM   2039  CA  THR B1120      -3.278  -8.407 -98.626  1.00 70.60           C
ATOM   2040  CA  PHE B1121       0.336  -9.561 -98.707  1.00 67.18           C
ATOM   2041  CA  VAL B1122       2.006 -12.663-100.138  1.00 69.76           C
ATOM   2042  CA  SER B1123       4.358 -14.982 -98.245  1.00 71.03           C
ATOM   2043  CA  GLY B1124       5.102 -18.547 -99.304  1.00 75.31           C
ATOM   2044  CA  ASN B1125       2.882 -20.984-101.156  1.00 82.41           C
ATOM   2045  CA  CYS B1126      -0.374 -22.853-100.552  1.00 84.08           C
ATOM   2046  CA  ASP B1127       1.272 -26.068 -99.300  1.00 85.00           C
ATOM   2047  CA  VAL B1128       2.200 -24.811 -95.817  1.00 76.48           C
ATOM   2048  CA  VAL B1129      -0.864 -22.990 -94.407  1.00 72.14           C
ATOM   2049  CA  ILE B1130      -3.357 -25.300 -92.698  1.00 69.21           C
ATOM   2050  CA  GLY B1131      -6.907 -24.359 -93.646  1.00 69.75           C
ATOM   2051  CA  ILE B1132      -6.107 -22.267 -96.738  1.00 71.37           C
ATOM   2052  CA  VAL B1133      -9.046 -21.918 -99.138  1.00 74.80           C
ATOM   2053  CA  ASN B1134      -9.229 -21.365-102.884  1.00 86.07           C
ATOM   2054  CA  ASN B1135     -10.513 -17.994-104.136  1.00 81.55           C
ATOM   2055  CA  THR B1136      -9.772 -15.111-106.493  1.00 82.74           C
ATOM   2056  CA  VAL B1137      -7.611 -12.095-105.615  1.00 81.98           C
ATOM   2057  CA  TYR B1138      -8.181  -9.080-107.853  1.00 86.19           C
ATOM   2058  CA  ASP B1139      -5.420  -6.806-109.168  1.00 93.40           C
ATOM   2059  CA  PRO B1140      -6.963  -3.399-110.258  1.00 92.83           C
TER    2060      PRO B1140
ATOM   2515  CA  CYS E  15      31.968 -48.964  11.436  1.00180.73           C
ATOM   2516  CA  VAL E  16      34.691 -50.778   9.439  1.00181.29           C
ATOM   2517  CA  ASN E  17      37.418 -49.094   7.367  1.00184.98           C
ATOM   2518  CA  LEU E  18      38.283 -51.007   4.183  1.00177.34           C
ATOM   2519  CA  THR E  19      41.932 -50.509   3.217  1.00176.59           C
ATOM   2520  CA  THR E  20      42.654 -53.199   0.573  1.00174.56           C
ATOM   2521  CA  ARG E  21      41.940 -50.849  -2.331  1.00170.43           C
ATOM   2522  CA  THR E  22      44.086 -49.786  -5.276  1.00168.14           C
ATOM   2523  CA  GLN E  23      44.323 -45.990  -5.615  1.00167.91           C
ATOM   2524  CA  LEU E  24      43.817 -44.885  -9.230  1.00163.89           C
ATOM   2525  CA  PRO E  25      42.703 -41.478 -10.609  1.00159.19           C
ATOM   2526  CA  PRO E  26      39.058 -40.786 -11.585  1.00148.01           C
ATOM   2527  CA  ALA E  27      37.958 -41.691 -15.109  1.00135.20           C
ATOM   2528  CA  TYR E  28      36.010 -39.431 -17.470  1.00128.82           C
ATOM   2529  CA  THR E  29      33.495 -40.031 -20.273  1.00120.12           C
ATOM   2530  CA  ASN E  30      31.089 -38.303 -22.690  1.00110.51           C
ATOM   2531  CA  SER E  31      27.456 -37.728 -21.646  1.00107.33           C
ATOM   2532  CA  PHE E  32      26.192 -37.776 -25.292  1.00105.71           C
ATOM   2533  CA  THR E  33      22.432 -37.038 -25.263  1.00101.63           C
ATOM   2534  CA  ARG E  34      21.567 -38.635 -21.900  1.00102.05           C
```

```
ATOM   2535  CA  GLY E  35      20.028 -36.954 -18.875  1.00 97.55           C
ATOM   2536  CA  VAL E  36      16.744 -35.616 -20.284  1.00 93.47           C
ATOM   2537  CA  TYR E  37      13.753 -35.928 -17.935  1.00 90.01           C
ATOM   2538  CA  TYR E  38      10.109 -34.859 -17.906  1.00 87.13           C
ATOM   2539  CA  PRO E  39      10.311 -31.338 -16.391  1.00 87.81           C
ATOM   2540  CA  ASP E  40       6.713 -31.284 -15.117  1.00 89.50           C
ATOM   2541  CA  LYS E  41       3.588 -33.375 -14.466  1.00 89.60           C
ATOM   2542  CA  VAL E  42       1.453 -32.213 -17.421  1.00 86.34           C
ATOM   2543  CA  PHE E  43       0.672 -34.357 -20.465  1.00 86.50           C
ATOM   2544  CA  ARG E  44       1.294 -32.822 -23.862  1.00 85.08           C
ATOM   2545  CA  SER E  45       1.653 -34.591 -27.176  1.00 86.67           C
ATOM   2546  CA  SER E  46       2.839 -34.005 -30.768  1.00 88.98           C
ATOM   2547  CA  VAL E  47       4.427 -30.719 -29.668  1.00 86.56           C
ATOM   2548  CA  LEU E  48       7.761 -29.027 -29.189  1.00 84.21           C
ATOM   2549  CA  HIS E  49       8.016 -27.477 -25.726  1.00 83.76           C
ATOM   2550  CA  SER E  50      10.678 -24.978 -24.653  1.00 84.33           C
ATOM   2551  CA  THR E  51      11.585 -25.233 -20.970  1.00 85.65           C
ATOM   2552  CA  GLN E  52      14.177 -23.521 -18.768  1.00 88.88           C
ATOM   2553  CA  ASP E  53      15.292 -25.914 -16.026  1.00 91.39           C
ATOM   2554  CA  LEU E  54      18.316 -27.700 -14.581  1.00 91.26           C
ATOM   2555  CA  PHE E  55      19.460 -30.243 -17.169  1.00 90.15           C
ATOM   2556  CA  LEU E  56      22.575 -32.233 -17.892  1.00 93.57           C
ATOM   2557  CA  PRO E  57      24.302 -30.426 -20.803  1.00 94.10           C
ATOM   2558  CA  PHE E  58      24.716 -32.440 -23.981  1.00 94.94           C
ATOM   2559  CA  PHE E  59      28.177 -33.974 -24.565  1.00101.73           C
ATOM   2560  CA  SER E  60      29.331 -32.956 -21.091  1.00107.80           C
ATOM   2561  CA  ASN E  61      32.325 -34.367 -19.225  1.00120.36           C
ATOM   2562  CA  VAL E  62      30.975 -37.032 -16.838  1.00119.62           C
ATOM   2563  CA  THR E  63      33.115 -38.356 -13.991  1.00123.08           C
ATOM   2564  CA  TRP E  64      33.350 -42.165 -14.096  1.00129.59           C
ATOM   2565  CA  PHE E  65      33.995 -44.355 -11.029  1.00128.99           C
ATOM   2566  CA  HIS E  66      34.440 -48.105 -10.665  1.00134.22           C
ATOM   2567  CA  ARG E  78      37.110 -52.132  -7.096  1.00151.90           C
ATOM   2568  CA  PHE E  79      35.254 -50.093  -4.433  1.00153.60           C
ATOM   2569  CA  ASP E  80      35.492 -46.475  -5.572  1.00143.21           C
ATOM   2570  CA  ASN E  81      33.068 -44.166  -3.711  1.00140.17           C
ATOM   2571  CA  PRO E  82      34.257 -40.913  -2.060  1.00127.85           C
ATOM   2572  CA  VAL E  83      32.033 -38.002  -1.138  1.00119.63           C
ATOM   2573  CA  LEU E  84      31.264 -35.797  -4.138  1.00115.44           C
ATOM   2574  CA  PRO E  85      29.916 -32.235  -4.382  1.00110.47           C
ATOM   2575  CA  PHE E  86      26.231 -31.648  -5.076  1.00108.08           C
ATOM   2576  CA  ASN E  87      25.646 -28.254  -6.671  1.00106.02           C
ATOM   2577  CA  ASP E  88      22.405 -27.480  -8.551  1.00101.70           C
ATOM   2578  CA  GLY E  89      21.643 -31.184  -8.917  1.00 99.82           C
ATOM   2579  CA  VAL E  90      23.265 -34.353 -10.152  1.00 99.81           C
ATOM   2580  CA  TYR E  91      22.870 -37.005 -12.823  1.00102.60           C
ATOM   2581  CA  PHE E  92      23.737 -40.526 -11.728  1.00111.59           C
ATOM   2582  CA  ALA E  93      23.961 -43.674 -13.815  1.00119.16           C
ATOM   2583  CA  SER E  94      24.991 -47.236 -13.048  1.00128.82           C
ATOM   2584  CA  THR E  95      25.947 -50.175 -15.236  1.00139.08           C
ATOM   2585  CA  GLU E  96      25.570 -53.481 -13.394  1.00144.61           C
ATOM   2586  CA  LYS E  97      25.016 -57.179 -13.908  1.00148.90           C
ATOM   2587  CA  SER E  98      24.585 -58.414 -10.309  1.00150.39           C
ATOM   2588  CA  ASN E  99      22.762 -55.699  -8.218  1.00147.60           C
ATOM   2589  CA  ILE E 100      25.711 -54.062  -6.440  1.00147.05           C
ATOM   2590  CA  ILE E 101      24.194 -50.554  -5.981  1.00144.06           C
ATOM   2591  CA  ARG E 102      22.568 -49.846  -2.636  1.00143.04           C
ATOM   2592  CA  GLY E 103      21.773 -46.165  -2.353  1.00130.83           C
ATOM   2593  CA  TRP E 104      22.831 -42.621  -1.590  1.00119.64           C
ATOM   2594  CA  ILE E 105      23.588 -40.238   1.281  1.00122.34           C
```

| | | | | | | | | |
|---|---|---|---|---|---|---|---|---|
| ATOM | 2595 | CA | PHE E 106 | 22.854 | -36.534 | 0.824 | 1.00116.55 | C |
| ATOM | 2596 | CA | GLY E 107 | 23.797 | -33.791 | 3.239 | 1.00118.54 | C |
| ATOM | 2597 | CA | THR E 108 | 26.086 | -30.959 | 4.253 | 1.00123.10 | C |
| ATOM | 2598 | CA | THR E 109 | 28.523 | -32.517 | 6.742 | 1.00128.26 | C |
| ATOM | 2599 | CA | LEU E 110 | 27.189 | -36.101 | 7.172 | 1.00133.89 | C |
| ATOM | 2600 | CA | ASP E 111 | 28.333 | -36.371 | 10.817 | 1.00136.65 | C |
| ATOM | 2601 | CA | SER E 112 | 25.110 | -35.819 | 12.889 | 1.00135.64 | C |
| ATOM | 2602 | CA | LYS E 113 | 25.765 | -32.048 | 13.131 | 1.00134.03 | C |
| ATOM | 2603 | CA | THR E 114 | 23.332 | -31.313 | 10.285 | 1.00131.52 | C |
| ATOM | 2604 | CA | GLN E 115 | 20.247 | -32.899 | 8.761 | 1.00129.33 | C |
| ATOM | 2605 | CA | SER E 116 | 20.925 | -35.635 | 6.201 | 1.00127.50 | C |
| ATOM | 2606 | CA | LEU E 117 | 18.935 | -37.738 | 3.732 | 1.00121.53 | C |
| ATOM | 2607 | CA | LEU E 118 | 19.490 | -41.503 | 3.488 | 1.00126.00 | C |
| ATOM | 2608 | CA | ILE E 119 | 18.099 | -43.534 | 0.572 | 1.00124.76 | C |
| ATOM | 2609 | CA | VAL E 120 | 18.686 | -47.306 | 0.693 | 1.00131.78 | C |
| ATOM | 2610 | CA | ASN E 121 | 17.306 | -50.038 | -1.577 | 1.00137.17 | C |
| ATOM | 2611 | CA | ASN E 122 | 17.664 | -53.172 | 0.511 | 1.00146.26 | C |
| ATOM | 2612 | CA | ALA E 123 | 16.786 | -56.785 | -0.345 | 1.00148.37 | C |
| ATOM | 2613 | CA | THR E 124 | 13.072 | -56.357 | 0.439 | 1.00143.58 | C |
| ATOM | 2614 | CA | ASN E 125 | 12.136 | -52.686 | -0.226 | 1.00140.62 | C |
| ATOM | 2615 | CA | VAL E 126 | 13.298 | -49.068 | -0.616 | 1.00130.58 | C |
| ATOM | 2616 | CA | VAL E 127 | 13.590 | -47.005 | 2.573 | 1.00129.50 | C |
| ATOM | 2617 | CA | ILE E 128 | 14.057 | -43.218 | 2.681 | 1.00123.66 | C |
| ATOM | 2618 | CA | LYS E 129 | 14.829 | -41.598 | 6.025 | 1.00126.74 | C |
| ATOM | 2619 | CA | VAL E 130 | 15.611 | -37.986 | 6.834 | 1.00128.51 | C |
| ATOM | 2620 | CA | CYS E 131 | 16.792 | -38.785 | 10.329 | 1.00139.93 | C |
| ATOM | 2621 | CA | GLU E 132 | 19.911 | -37.205 | 11.833 | 1.00141.15 | C |
| ATOM | 2622 | CA | PHE E 133 | 22.535 | -39.893 | 11.221 | 1.00142.56 | C |
| ATOM | 2623 | CA | GLN E 134 | 26.096 | -40.227 | 12.493 | 1.00145.13 | C |
| ATOM | 2624 | CA | PHE E 135 | 27.665 | -41.759 | 9.412 | 1.00148.27 | C |
| ATOM | 2625 | CA | CYS E 136 | 30.714 | -43.987 | 9.083 | 1.00166.90 | C |
| ATOM | 2626 | CA | ASN E 137 | 33.827 | -42.759 | 7.320 | 1.00173.92 | C |
| ATOM | 2627 | CA | ASP E 138 | 33.618 | -45.782 | 4.988 | 1.00174.40 | C |
| ATOM | 2628 | CA | PRO E 139 | 29.928 | -46.786 | 4.762 | 1.00169.70 | C |
| ATOM | 2629 | CA | PHE E 140 | 28.900 | -49.934 | 2.879 | 1.00172.19 | C |
| ATOM | 2630 | CA | LEU E 141 | 26.665 | -53.003 | 2.881 | 1.00169.16 | C |
| ATOM | 2631 | CA | GLY E 142 | 27.709 | -56.628 | 2.529 | 1.00175.73 | C |
| ATOM | 2632 | CA | VAL E 143 | 26.380 | -59.691 | 0.745 | 1.00176.06 | C |
| ATOM | 2633 | CA | CYS E 152 | 24.452 | -64.741 | -2.048 | 1.00189.28 | C |
| ATOM | 2634 | CA | MET E 153 | 22.454 | -63.167 | 0.788 | 1.00184.21 | C |
| ATOM | 2635 | CA | GLU E 154 | 22.630 | -59.782 | 2.482 | 1.00176.13 | C |
| ATOM | 2636 | CA | SER E 155 | 24.198 | -59.714 | 5.944 | 1.00177.51 | C |
| ATOM | 2637 | CA | GLU E 156 | 26.288 | -56.710 | 6.974 | 1.00173.52 | C |
| ATOM | 2638 | CA | PHE E 157 | 24.831 | -53.217 | 7.365 | 1.00166.85 | C |
| ATOM | 2639 | CA | ARG E 158 | 27.518 | -50.703 | 8.397 | 1.00169.66 | C |
| ATOM | 2640 | CA | VAL E 159 | 26.185 | -47.315 | 7.284 | 1.00160.11 | C |
| ATOM | 2641 | CA | TYR E 160 | 25.655 | -45.378 | 10.520 | 1.00151.89 | C |
| ATOM | 2642 | CA | SER E 161 | 25.831 | -45.652 | 14.296 | 1.00151.37 | C |
| ATOM | 2643 | CA | SER E 162 | 23.732 | -43.067 | 16.135 | 1.00149.19 | C |
| ATOM | 2644 | CA | ALA E 163 | 20.350 | -42.127 | 14.675 | 1.00145.94 | C |
| ATOM | 2645 | CA | ASN E 164 | 17.795 | -39.850 | 16.339 | 1.00142.49 | C |
| ATOM | 2646 | CA | ASN E 165 | 15.687 | -36.739 | 15.568 | 1.00141.42 | C |
| ATOM | 2647 | CA | CYS E 166 | 13.854 | -38.486 | 12.720 | 1.00140.72 | C |
| ATOM | 2648 | CA | THR E 167 | 11.786 | -36.080 | 10.621 | 1.00130.53 | C |
| ATOM | 2649 | CA | PHE E 168 | 10.639 | -38.041 | 7.556 | 1.00121.04 | C |
| ATOM | 2650 | CA | GLU E 169 | 10.333 | -41.671 | 6.471 | 1.00119.64 | C |
| ATOM | 2651 | CA | TYR E 170 | 9.071 | -43.278 | 3.262 | 1.00122.06 | C |
| ATOM | 2652 | CA | VAL E 171 | 9.032 | -46.920 | 2.089 | 1.00123.79 | C |
| ATOM | 2653 | CA | SER E 172 | 8.084 | -48.211 | -1.372 | 1.00125.40 | C |
| ATOM | 2654 | CA | GLN E 173 | 8.579 | -50.686 | -4.227 | 1.00124.36 | C |

| | | | | | | | | | | |
|---|---|---|---|---|---|---|---|---|---|---|
| ATOM | 2655 | CA | PRO | E | 174 | 12.200 | -51.526 | -5.260 | 1.00125.90 | C |
| ATOM | 2656 | CA | PHE | E | 175 | 14.255 | -48.890 | -7.196 | 1.00124.65 | C |
| ATOM | 2657 | CA | ASN | E | 185 | 29.519 | -57.227 | -20.225 | 1.00150.77 | C |
| ATOM | 2658 | CA | PHE | E | 186 | 26.802 | -55.569 | -18.150 | 1.00147.13 | C |
| ATOM | 2659 | CA | LYS | E | 187 | 23.052 | -56.029 | -18.571 | 1.00141.01 | C |
| ATOM | 2660 | CA | ASN | E | 188 | 21.277 | -53.332 | -16.520 | 1.00137.53 | C |
| ATOM | 2661 | CA | LEU | E | 189 | 21.617 | -49.568 | -16.998 | 1.00126.28 | C |
| ATOM | 2662 | CA | ARG | E | 190 | 19.935 | -47.353 | -14.405 | 1.00121.72 | C |
| ATOM | 2663 | CA | GLU | E | 191 | 19.691 | -43.574 | -14.720 | 1.00107.95 | C |
| ATOM | 2664 | CA | PHE | E | 192 | 18.634 | -41.161 | -11.974 | 1.00102.87 | C |
| ATOM | 2665 | CA | VAL | E | 193 | 18.330 | -37.385 | -11.710 | 1.00 97.37 | C |
| ATOM | 2666 | CA | PHE | E | 194 | 18.504 | -35.817 | -8.240 | 1.00100.46 | C |
| ATOM | 2667 | CA | LYS | E | 195 | 17.321 | -32.224 | -7.781 | 1.00 98.20 | C |
| ATOM | 2668 | CA | ASN | E | 196 | 16.883 | -30.141 | -4.610 | 1.00105.65 | C |
| ATOM | 2669 | CA | ILE | E | 197 | 14.553 | -27.229 | -5.464 | 1.00106.01 | C |
| ATOM | 2670 | CA | ASP | E | 198 | 12.081 | -25.219 | -3.315 | 1.00108.15 | C |
| ATOM | 2671 | CA | GLY | E | 199 | 11.879 | -27.733 | -0.511 | 1.00106.29 | C |
| ATOM | 2672 | CA | TYR | E | 200 | 11.443 | -30.788 | -2.738 | 1.00101.76 | C |
| ATOM | 2673 | CA | PHE | E | 201 | 14.058 | -33.398 | -3.528 | 1.00103.25 | C |
| ATOM | 2674 | CA | LYS | E | 202 | 13.035 | -34.705 | -6.936 | 1.00 97.20 | C |
| ATOM | 2675 | CA | ILE | E | 203 | 14.125 | -38.166 | -8.074 | 1.00 97.35 | C |
| ATOM | 2676 | CA | TYR | E | 204 | 13.743 | -39.130 | -11.736 | 1.00 94.05 | C |
| ATOM | 2677 | CA | SER | E | 205 | 14.670 | -42.554 | -13.061 | 1.00 99.13 | C |
| ATOM | 2678 | CA | LYS | E | 206 | 14.885 | -44.816 | -16.103 | 1.00108.70 | C |
| ATOM | 2679 | CA | HIS | E | 207 | 15.786 | -48.515 | -16.352 | 1.00121.06 | C |
| ATOM | 2680 | CA | THR | E | 208 | 17.076 | -49.912 | -19.655 | 1.00128.32 | C |
| ATOM | 2681 | CA | PRO | E | 209 | 19.056 | -52.906 | -20.937 | 1.00133.34 | C |
| ATOM | 2682 | CA | ILE | E | 210 | 22.717 | -52.344 | -21.789 | 1.00137.28 | C |
| ATOM | 2683 | CA | ASN | E | 211 | 25.252 | -54.415 | -23.710 | 1.00141.03 | C |
| ATOM | 2684 | CA | LEU | E | 212 | 28.605 | -52.627 | -23.512 | 1.00143.87 | C |
| ATOM | 2685 | CA | VAL | E | 213 | 31.517 | -53.861 | -21.436 | 1.00145.28 | C |
| ATOM | 2686 | CA | ARG | E | 214 | 33.204 | -50.536 | -20.693 | 1.00139.33 | C |
| ATOM | 2687 | CA | ASP | E | 215 | 30.967 | -47.611 | -21.733 | 1.00134.09 | C |
| ATOM | 2688 | CA | LEU | E | 216 | 27.687 | -45.554 | -21.661 | 1.00126.27 | C |
| ATOM | 2689 | CA | PRO | E | 217 | 25.240 | -46.041 | -24.566 | 1.00121.48 | C |
| ATOM | 2690 | CA | GLN | E | 218 | 24.570 | -43.542 | -27.318 | 1.00115.46 | C |
| ATOM | 2691 | CA | GLY | E | 219 | 20.955 | -42.733 | -28.041 | 1.00107.17 | C |
| ATOM | 2692 | CA | PHE | E | 220 | 18.039 | -40.894 | -26.462 | 1.00 99.00 | C |
| ATOM | 2693 | CA | SER | E | 221 | 15.573 | -41.808 | -23.724 | 1.00100.97 | C |
| ATOM | 2694 | CA | ALA | E | 222 | 13.685 | -39.732 | -21.176 | 1.00 98.19 | C |
| ATOM | 2695 | CA | LEU | E | 223 | 13.709 | -40.288 | -17.420 | 1.00 97.51 | C |
| ATOM | 2696 | CA | GLU | E | 224 | 10.292 | -40.400 | -15.837 | 1.00 95.33 | C |
| ATOM | 2697 | CA | PRO | E | 225 | 9.900 | -38.869 | -12.353 | 1.00 93.34 | C |
| ATOM | 2698 | CA | LEU | E | 226 | 10.079 | -41.431 | -9.586 | 1.00 98.27 | C |
| ATOM | 2699 | CA | VAL | E | 227 | 9.300 | -39.501 | -6.387 | 1.00101.53 | C |
| ATOM | 2700 | CA | ASP | E | 228 | 9.039 | -35.914 | -5.104 | 1.00103.97 | C |
| ATOM | 2701 | CA | LEU | E | 229 | 10.153 | -35.788 | -1.460 | 1.00105.49 | C |
| ATOM | 2702 | CA | PRO | E | 230 | 8.952 | -32.663 | 0.432 | 1.00101.78 | C |
| ATOM | 2703 | CA | ILE | E | 231 | 12.160 | -32.304 | 2.456 | 1.00108.98 | C |
| ATOM | 2704 | CA | GLY | E | 232 | 13.831 | -28.917 | 2.986 | 1.00112.23 | C |
| ATOM | 2705 | CA | ILE | E | 233 | 17.448 | -30.093 | 3.274 | 1.00115.52 | C |
| ATOM | 2706 | CA | ASN | E | 234 | 20.602 | -28.093 | 2.591 | 1.00117.95 | C |
| ATOM | 2707 | CA | ILE | E | 235 | 22.412 | -30.517 | 0.265 | 1.00113.83 | C |
| ATOM | 2708 | CA | THR | E | 236 | 25.970 | -29.771 | -0.857 | 1.00114.70 | C |
| ATOM | 2709 | CA | ARG | E | 237 | 27.616 | -33.218 | -0.663 | 1.00114.95 | C |
| ATOM | 2710 | CA | PHE | E | 238 | 26.592 | -36.789 | -1.425 | 1.00115.50 | C |
| ATOM | 2711 | CA | GLN | E | 239 | 27.986 | -40.321 | -1.246 | 1.00125.19 | C |
| ATOM | 2712 | CA | THR | E | 240 | 27.128 | -43.582 | -3.024 | 1.00128.24 | C |
| ATOM | 2713 | CA | LEU | E | 241 | 26.344 | -46.816 | -1.123 | 1.00138.25 | C |
| ATOM | 2714 | CA | LEU | E | 242 | 27.649 | -50.035 | -2.723 | 1.00146.67 | C |

| | | | | | | | | | | | |
|---|---|---|---|---|---|---|---|---|---|---|---|
| ATOM | 2715 | CA | ALA | E | 243 | 27.460 | -53.728 | -1.824 | 1.00152.10 | | C |
| ATOM | 2716 | CA | TRP | E | 258 | 35.752 | -59.244 | 1.213 | 1.00169.46 | | C |
| ATOM | 2717 | CA | THR | E | 259 | 35.758 | -59.037 | -2.589 | 1.00167.76 | | C |
| ATOM | 2718 | CA | ALA | E | 260 | 34.734 | -56.595 | -5.308 | 1.00164.58 | | C |
| ATOM | 2719 | CA | GLY | E | 261 | 31.888 | -57.078 | -7.761 | 1.00158.21 | | C |
| ATOM | 2720 | CA | ALA | E | 262 | 31.394 | -56.058 | -11.389 | 1.00152.47 | | C |
| ATOM | 2721 | CA | ALA | E | 263 | 29.786 | -52.593 | -11.488 | 1.00145.85 | | C |
| ATOM | 2722 | CA | ALA | E | 264 | 30.487 | -49.045 | -12.634 | 1.00136.69 | | C |
| ATOM | 2723 | CA | TYR | E | 265 | 28.807 | -45.690 | -12.141 | 1.00128.41 | | C |
| ATOM | 2724 | CA | TYR | E | 266 | 28.864 | -42.234 | -13.710 | 1.00122.28 | | C |
| ATOM | 2725 | CA | VAL | E | 267 | 28.218 | -38.834 | -12.088 | 1.00113.18 | | C |
| ATOM | 2726 | CA | GLY | E | 268 | 27.367 | -35.668 | -14.016 | 1.00105.32 | | C |
| ATOM | 2727 | CA | TYR | E | 269 | 26.147 | -32.230 | -12.966 | 1.00100.84 | | C |
| ATOM | 2728 | CA | LEU | E | 270 | 23.062 | -30.240 | -13.916 | 1.00 | 94.11 | C |
| ATOM | 2729 | CA | GLN | E | 271 | 23.187 | -26.654 | -15.172 | 1.00 | 91.66 | C |
| ATOM | 2730 | CA | PRO | E | 272 | 20.512 | -24.026 | -15.914 | 1.00 | 90.58 | C |
| ATOM | 2731 | CA | ARG | E | 273 | 19.655 | -24.700 | -19.561 | 1.00 | 89.02 | C |
| ATOM | 2732 | CA | THR | E | 274 | 16.796 | -24.276 | -21.999 | 1.00 | 86.14 | C |
| ATOM | 2733 | CA | PHE | E | 275 | 15.720 | -27.484 | -23.744 | 1.00 | 85.72 | C |
| ATOM | 2734 | CA | LEU | E | 276 | 13.341 | -28.021 | -26.647 | 1.00 | 83.91 | C |
| ATOM | 2735 | CA | LEU | E | 277 | 11.483 | -31.280 | -25.955 | 1.00 | 84.24 | C |
| ATOM | 2736 | CA | LYS | E | 278 | 9.712 | -33.264 | -28.682 | 1.00 | 87.03 | C |
| ATOM | 2737 | CA | TYR | E | 279 | 6.498 | -34.988 | -27.588 | 1.00 | 86.14 | C |
| ATOM | 2738 | CA | ASN | E | 280 | 5.114 | -37.701 | -29.886 | 1.00 | 89.03 | C |
| ATOM | 2739 | CA | GLU | E | 281 | 1.486 | -38.774 | -30.444 | 1.00 | 90.49 | C |
| ATOM | 2740 | CA | ASN | E | 282 | 1.456 | -40.707 | -27.145 | 1.00 | 93.60 | C |
| ATOM | 2741 | CA | GLY | E | 283 | 3.058 | -37.848 | -25.208 | 1.00 | 88.94 | C |
| ATOM | 2742 | CA | THR | E | 284 | 6.438 | -39.565 | -24.897 | 1.00 | 88.68 | C |
| ATOM | 2743 | CA | ILE | E | 285 | 9.559 | -37.429 | -25.207 | 1.00 | 88.12 | C |
| ATOM | 2744 | CA | THR | E | 286 | 11.480 | -38.866 | -28.154 | 1.00 | 89.44 | C |
| ATOM | 2745 | CA | ASP | E | 287 | 14.038 | -36.137 | -28.856 | 1.00 | 90.47 | C |
| ATOM | 2746 | CA | ALA | E | 288 | 15.391 | -32.894 | -27.435 | 1.00 | 86.71 | C |
| ATOM | 2747 | CA | VAL | E | 289 | 17.576 | -29.941 | -28.349 | 1.00 | 84.36 | C |
| ATOM | 2748 | CA | ASP | E | 290 | 20.006 | -28.415 | -25.846 | 1.00 | 86.49 | C |
| ATOM | 2749 | CA | CYS | E | 291 | 19.593 | -24.742 | -26.860 | 1.00 | 85.31 | C |
| ATOM | 2750 | CA | ALA | E | 292 | 23.211 | -23.836 | -25.949 | 1.00 | 83.92 | C |
| ATOM | 2751 | CA | LEU | E | 293 | 25.151 | -26.880 | -27.214 | 1.00 | 83.72 | C |
| ATOM | 2752 | CA | ASP | E | 294 | 26.101 | -25.398 | -30.591 | 1.00 | 81.70 | C |
| ATOM | 2753 | CA | PRO | E | 295 | 24.865 | -22.742 | -33.096 | 1.00 | 77.26 | C |
| ATOM | 2754 | CA | LEU | E | 296 | 22.596 | -25.222 | -34.918 | 1.00 | 77.82 | C |
| ATOM | 2755 | CA | SER | E | 297 | 20.888 | -26.260 | -31.666 | 1.00 | 80.30 | C |
| ATOM | 2756 | CA | GLU | E | 298 | 20.289 | -22.591 | -30.849 | 1.00 | 80.24 | C |
| ATOM | 2757 | CA | THR | E | 299 | 18.808 | -22.107 | -34.325 | 1.00 | 79.24 | C |
| ATOM | 2758 | CA | LYS | E | 300 | 16.522 | -25.120 | -33.825 | 1.00 | 78.04 | C |
| ATOM | 2759 | CA | CYS | E | 301 | 15.292 | -23.681 | -30.519 | 1.00 | 82.22 | C |
| ATOM | 2760 | CA | THR | E | 302 | 14.807 | -20.196 | -32.014 | 1.00 | 79.88 | C |
| ATOM | 2761 | CA | LEU | E | 303 | 12.636 | -21.485 | -34.867 | 1.00 | 79.47 | C |
| ATOM | 2762 | CA | LYS | E | 304 | 10.845 | -24.001 | -32.559 | 1.00 | 81.84 | C |
| ATOM | 2763 | CA | SER | E | 305 | 11.554 | -26.855 | -34.957 | 1.00 | 81.94 | C |
| ATOM | 2764 | CA | PHE | E | 306 | 13.895 | -29.813 | -35.297 | 1.00 | 84.64 | C |
| ATOM | 2765 | CA | THR | E | 307 | 14.189 | -29.121 | -39.038 | 1.00 | 84.97 | C |
| ATOM | 2766 | CA | VAL | E | 308 | 15.855 | -25.934 | -40.279 | 1.00 | 82.93 | C |
| ATOM | 2767 | CA | GLU | E | 309 | 15.530 | -24.747 | -43.865 | 1.00 | 81.86 | C |
| ATOM | 2768 | CA | LYS | E | 310 | 18.372 | -23.185 | -45.845 | 1.00 | 77.28 | C |
| ATOM | 2769 | CA | GLY | E | 311 | 18.826 | -19.540 | -44.944 | 1.00 | 72.92 | C |
| ATOM | 2770 | CA | ILE | E | 312 | 20.173 | -17.081 | -42.410 | 1.00 | 71.34 | C |
| ATOM | 2771 | CA | TYR | E | 313 | 18.463 | -16.752 | -39.039 | 1.00 | 73.37 | C |
| ATOM | 2772 | CA | GLN | E | 314 | 19.175 | -14.276 | -36.247 | 1.00 | 76.74 | C |
| ATOM | 2773 | CA | THR | E | 315 | 19.343 | -16.478 | -33.171 | 1.00 | 78.06 | C |
| ATOM | 2774 | CA | SER | E | 316 | 21.179 | -14.719 | -30.335 | 1.00 | 78.49 | C |

| | | | | | | | | | |
|---|---|---|---|---|---|---|---|---|---|
| ATOM | 2775 | CA | ASN E 317 | 23.187 | -11.744 | -29.105 | 1.00 | 78.31 | C |
| ATOM | 2776 | CA | PHE E 318 | 26.937 | -11.652 | -28.498 | 1.00 | 83.54 | C |
| ATOM | 2777 | CA | ARG E 319 | 28.248 | -9.615 | -25.576 | 1.00 | 86.02 | C |
| ATOM | 2778 | CA | VAL E 320 | 31.706 | -9.419 | -24.014 | 1.00 | 86.35 | C |
| ATOM | 2779 | CA | GLN E 321 | 31.447 | -10.102 | -20.329 | 1.00 | 88.28 | C |
| ATOM | 2780 | CA | PRO E 322 | 33.285 | -7.962 | -17.762 | 1.00 | 87.66 | C |
| ATOM | 2781 | CA | THR E 323 | 36.158 | -9.575 | -15.891 | 1.00 | 90.76 | C |
| ATOM | 2782 | CA | GLU E 324 | 36.945 | -6.946 | -13.235 | 1.00 | 89.11 | C |
| ATOM | 2783 | CA | SER E 325 | 35.078 | -4.620 | -10.891 | 1.00 | 85.34 | C |
| ATOM | 2784 | CA | ILE E 326 | 36.498 | -1.087 | -10.853 | 1.00 | 83.95 | C |
| ATOM | 2785 | CA | VAL E 327 | 35.395 | 0.954 | -7.858 | 1.00 | 83.19 | C |
| ATOM | 2786 | CA | ARG E 328 | 36.449 | 4.601 | -7.497 | 1.00 | 82.15 | C |
| ATOM | 2787 | CA | PHE E 329 | 35.425 | 6.934 | -4.672 | 1.00 | 84.03 | C |
| ATOM | 2788 | CA | PRO E 330 | 36.998 | 10.176 | -3.324 | 1.00 | 85.12 | C |
| ATOM | 2789 | CA | ASN E 331 | 40.012 | 9.537 | -1.081 | 1.00 | 90.85 | C |
| ATOM | 2790 | CA | ILE E 332 | 38.473 | 11.039 | 2.052 | 1.00 | 84.57 | C |
| ATOM | 2791 | CA | THR E 333 | 38.760 | 9.462 | 5.494 | 1.00 | 85.53 | C |
| ATOM | 2792 | CA | ASN E 334 | 36.780 | 11.673 | 7.901 | 1.00 | 80.76 | C |
| ATOM | 2793 | CA | LEU E 335 | 33.614 | 10.097 | 9.268 | 1.00 | 79.08 | C |
| ATOM | 2794 | CA | CYS E 336 | 30.221 | 11.757 | 9.028 | 1.00 | 80.64 | C |
| ATOM | 2795 | CA | PRO E 337 | 28.963 | 13.280 | 12.316 | 1.00 | 77.79 | C |
| ATOM | 2796 | CA | PHE E 338 | 25.985 | 10.943 | 12.683 | 1.00 | 76.98 | C |
| ATOM | 2797 | CA | GLY E 339 | 26.948 | 10.078 | 16.260 | 1.00 | 81.76 | C |
| ATOM | 2798 | CA | GLU E 340 | 26.718 | 13.764 | 17.237 | 1.00 | 81.69 | C |
| ATOM | 2799 | CA | VAL E 341 | 23.125 | 13.857 | 15.977 | 1.00 | 73.54 | C |
| ATOM | 2800 | CA | PHE E 342 | 22.048 | 10.689 | 17.788 | 1.00 | 70.43 | C |
| ATOM | 2801 | CA | ASN E 343 | 23.795 | 11.307 | 21.134 | 1.00 | 82.77 | C |
| ATOM | 2802 | CA | ALA E 344 | 23.051 | 15.024 | 21.590 | 1.00 | 77.53 | C |
| ATOM | 2803 | CA | THR E 345 | 22.041 | 15.808 | 25.172 | 1.00 | 79.06 | C |
| ATOM | 2804 | CA | ARG E 346 | 19.046 | 17.940 | 24.161 | 1.00 | 77.19 | C |
| ATOM | 2805 | CA | PHE E 347 | 16.707 | 17.202 | 21.270 | 1.00 | 65.44 | C |
| ATOM | 2806 | CA | ALA E 348 | 14.521 | 19.774 | 19.577 | 1.00 | 63.31 | C |
| ATOM | 2807 | CA | SER E 349 | 10.753 | 19.941 | 19.753 | 1.00 | 64.75 | C |
| ATOM | 2808 | CA | VAL E 350 | 9.033 | 18.623 | 16.618 | 1.00 | 62.28 | C |
| ATOM | 2809 | CA | TYR E 351 | 7.560 | 22.069 | 15.841 | 1.00 | 64.03 | C |
| ATOM | 2810 | CA | ALA E 352 | 11.118 | 23.509 | 15.834 | 1.00 | 62.94 | C |
| ATOM | 2811 | CA | TRP E 353 | 13.091 | 20.463 | 14.664 | 1.00 | 61.66 | C |
| ATOM | 2812 | CA | ASN E 354 | 16.862 | 20.597 | 14.236 | 1.00 | 62.90 | C |
| ATOM | 2813 | CA | ARG E 355 | 18.613 | 20.054 | 10.917 | 1.00 | 64.79 | C |
| ATOM | 2814 | CA | LYS E 356 | 22.295 | 19.040 | 10.785 | 1.00 | 70.37 | C |
| ATOM | 2815 | CA | ARG E 357 | 23.502 | 19.264 | 7.193 | 1.00 | 78.77 | C |
| ATOM | 2816 | CA | ILE E 358 | 26.033 | 16.459 | 6.592 | 1.00 | 74.91 | C |
| ATOM | 2817 | CA | SER E 359 | 28.574 | 16.810 | 3.770 | 1.00 | 79.18 | C |
| ATOM | 2818 | CA | ASN E 360 | 32.182 | 16.031 | 2.726 | 1.00 | 81.32 | C |
| ATOM | 2819 | CA | CYS E 361 | 32.605 | 12.857 | 4.786 | 1.00 | 79.43 | C |
| ATOM | 2820 | CA | VAL E 362 | 32.438 | 9.057 | 4.809 | 1.00 | 77.43 | C |
| ATOM | 2821 | CA | ALA E 363 | 29.068 | 7.740 | 5.978 | 1.00 | 75.37 | C |
| ATOM | 2822 | CA | ASP E 364 | 29.144 | 4.294 | 7.562 | 1.00 | 73.82 | C |
| ATOM | 2823 | CA | TYR E 365 | 25.482 | 3.303 | 7.721 | 1.00 | 71.34 | C |
| ATOM | 2824 | CA | SER E 366 | 26.079 | -0.245 | 8.941 | 1.00 | 71.73 | C |
| ATOM | 2825 | CA | VAL E 367 | 26.624 | 1.023 | 12.485 | 1.00 | 70.07 | C |
| ATOM | 2826 | CA | LEU E 368 | 23.188 | 2.656 | 12.606 | 1.00 | 67.22 | C |
| ATOM | 2827 | CA | TYR E 369 | 21.313 | -0.162 | 10.863 | 1.00 | 66.55 | C |
| ATOM | 2828 | CA | ASN E 370 | 22.833 | -3.061 | 12.782 | 1.00 | 68.16 | C |
| ATOM | 2829 | CA | SER E 371 | 22.389 | -1.292 | 16.121 | 1.00 | 68.95 | C |
| ATOM | 2830 | CA | ALA E 372 | 20.029 | -2.848 | 18.651 | 1.00 | 67.29 | C |
| ATOM | 2831 | CA | SER E 373 | 19.141 | 0.481 | 20.291 | 1.00 | 68.88 | C |
| ATOM | 2832 | CA | PHE E 374 | 16.695 | 1.480 | 17.510 | 1.00 | 65.62 | C |
| ATOM | 2833 | CA | SER E 375 | 13.095 | 0.279 | 17.335 | 1.00 | 63.49 | C |
| ATOM | 2834 | CA | THR E 376 | 11.973 | 2.000 | 14.128 | 1.00 | 62.83 | C |

| | | | | | | | | | | | |
|---|---|---|---|---|---|---|---|---|---|---|---|
| ATOM | 2835 | CA | PHE | E | 377 | 14.335 | 2.083 | 11.143 | 1.00 | 61.29 | C |
| ATOM | 2836 | CA | LYS | E | 378 | 12.273 | 2.549 | 7.987 | 1.00 | 64.58 | C |
| ATOM | 2837 | CA | CYS | E | 379 | 13.869 | 3.570 | 4.696 | 1.00 | 64.04 | C |
| ATOM | 2838 | CA | TYR | E | 380 | 11.982 | 5.091 | 1.757 | 1.00 | 64.13 | C |
| ATOM | 2839 | CA | GLY | E | 381 | 13.500 | 5.388 | -1.700 | 1.00 | 61.99 | C |
| ATOM | 2840 | CA | VAL | E | 382 | 16.572 | 3.287 | -0.824 | 1.00 | 63.01 | C |
| ATOM | 2841 | CA | SER | E | 383 | 17.060 | 0.036 | 0.923 | 1.00 | 65.80 | C |
| ATOM | 2842 | CA | PRO | E | 384 | 18.827 | 0.018 | 4.322 | 1.00 | 66.17 | C |
| ATOM | 2843 | CA | THR | E | 385 | 21.289 | -2.733 | 3.316 | 1.00 | 65.84 | C |
| ATOM | 2844 | CA | LYS | E | 386 | 22.535 | -0.952 | 0.176 | 1.00 | 66.03 | C |
| ATOM | 2845 | CA | LEU | E | 387 | 23.250 | 2.396 | 1.881 | 1.00 | 67.89 | C |
| ATOM | 2846 | CA | ASN | E | 388 | 26.983 | 1.758 | 2.198 | 1.00 | 70.64 | C |
| ATOM | 2847 | CA | ASP | E | 389 | 27.294 | 1.277 | -1.582 | 1.00 | 73.31 | C |
| ATOM | 2848 | CA | LEU | E | 390 | 25.703 | 4.647 | -2.412 | 1.00 | 70.30 | C |
| ATOM | 2849 | CA | CYS | E | 391 | 26.913 | 8.225 | -2.670 | 1.00 | 76.87 | C |
| ATOM | 2850 | CA | PHE | E | 392 | 24.813 | 11.288 | -2.040 | 1.00 | 74.90 | C |
| ATOM | 2851 | CA | THR | E | 393 | 25.500 | 14.955 | -2.627 | 1.00 | 80.74 | C |
| ATOM | 2852 | CA | ASN | E | 394 | 24.119 | 15.930 | 0.789 | 1.00 | 80.08 | C |
| ATOM | 2853 | CA | VAL | E | 395 | 22.483 | 14.336 | 3.848 | 1.00 | 72.47 | C |
| ATOM | 2854 | CA | TYR | E | 396 | 20.009 | 16.247 | 6.024 | 1.00 | 71.56 | C |
| ATOM | 2855 | CA | ALA | E | 397 | 20.010 | 14.822 | 9.584 | 1.00 | 65.87 | C |
| ATOM | 2856 | CA | ASP | E | 398 | 16.702 | 16.039 | 11.068 | 1.00 | 63.73 | C |
| ATOM | 2857 | CA | SER | E | 399 | 16.037 | 15.459 | 14.759 | 1.00 | 61.95 | C |
| ATOM | 2858 | CA | PHE | E | 400 | 13.072 | 15.862 | 17.124 | 1.00 | 60.14 | C |
| ATOM | 2859 | CA | VAL | E | 401 | 11.067 | 14.243 | 19.955 | 1.00 | 61.63 | C |
| ATOM | 2860 | CA | ILE | E | 402 | 7.533 | 12.939 | 19.342 | 1.00 | 64.36 | C |
| ATOM | 2861 | CA | ARG | E | 403 | 4.944 | 10.758 | 21.047 | 1.00 | 71.62 | C |
| ATOM | 2862 | CA | GLY | E | 404 | 5.110 | 7.015 | 20.318 | 1.00 | 70.94 | C |
| ATOM | 2863 | CA | ASP | E | 405 | 1.721 | 6.861 | 18.566 | 1.00 | 76.09 | C |
| ATOM | 2864 | CA | GLU | E | 406 | 2.938 | 9.600 | 16.237 | 1.00 | 71.89 | C |
| ATOM | 2865 | CA | VAL | E | 407 | 6.028 | 7.631 | 15.109 | 1.00 | 66.90 | C |
| ATOM | 2866 | CA | ARG | E | 408 | 3.955 | 6.215 | 12.234 | 1.00 | 69.52 | C |
| ATOM | 2867 | CA | GLN | E | 409 | 3.278 | 9.768 | 11.016 | 1.00 | 66.98 | C |
| ATOM | 2868 | CA | ILE | E | 410 | 6.953 | 10.330 | 10.105 | 1.00 | 62.69 | C |
| ATOM | 2869 | CA | ALA | E | 411 | 6.466 | 8.806 | 6.650 | 1.00 | 64.78 | C |
| ATOM | 2870 | CA | PRO | E | 412 | 5.552 | 9.973 | 3.132 | 1.00 | 67.36 | C |
| ATOM | 2871 | CA | GLY | E | 413 | 1.850 | 10.333 | 2.490 | 1.00 | 71.19 | C |
| ATOM | 2872 | CA | GLN | E | 414 | 0.707 | 10.588 | 6.116 | 1.00 | 70.51 | C |
| ATOM | 2873 | CA | THR | E | 415 | -1.892 | 12.817 | 7.758 | 1.00 | 72.93 | C |
| ATOM | 2874 | CA | GLY | E | 416 | -2.435 | 14.071 | 11.282 | 1.00 | 71.62 | C |
| ATOM | 2875 | CA | LYS | E | 417 | -0.958 | 16.772 | 13.510 | 1.00 | 68.78 | C |
| ATOM | 2876 | CA | ILE | E | 418 | 2.770 | 16.057 | 13.279 | 1.00 | 64.80 | C |
| ATOM | 2877 | CA | ALA | E | 419 | 2.606 | 15.443 | 9.537 | 1.00 | 65.52 | C |
| ATOM | 2878 | CA | ASP | E | 420 | 0.257 | 18.251 | 8.536 | 1.00 | 70.07 | C |
| ATOM | 2879 | CA | TYR | E | 421 | 1.633 | 20.768 | 11.116 | 1.00 | 66.38 | C |
| ATOM | 2880 | CA | ASN | E | 422 | 5.030 | 19.802 | 12.561 | 1.00 | 63.84 | C |
| ATOM | 2881 | CA | TYR | E | 423 | 7.261 | 17.647 | 10.263 | 1.00 | 61.29 | C |
| ATOM | 2882 | CA | LYS | E | 424 | 5.847 | 17.046 | 6.769 | 1.00 | 64.17 | C |
| ATOM | 2883 | CA | LEU | E | 425 | 7.785 | 14.644 | 4.504 | 1.00 | 64.93 | C |
| ATOM | 2884 | CA | PRO | E | 426 | 7.574 | 14.809 | 0.695 | 1.00 | 69.38 | C |
| ATOM | 2885 | CA | ASP | E | 427 | 6.087 | 11.901 | -1.212 | 1.00 | 77.05 | C |
| ATOM | 2886 | CA | ASP | E | 428 | 9.307 | 11.368 | -3.208 | 1.00 | 77.10 | C |
| ATOM | 2887 | CA | PHE | E | 429 | 11.373 | 11.310 | -0.002 | 1.00 | 67.39 | C |
| ATOM | 2888 | CA | THR | E | 430 | 14.629 | 9.362 | -0.106 | 1.00 | 66.63 | C |
| ATOM | 2889 | CA | GLY | E | 431 | 16.008 | 8.644 | 3.328 | 1.00 | 64.97 | C |
| ATOM | 2890 | CA | CYS | E | 432 | 15.575 | 6.715 | 6.538 | 1.00 | 64.56 | C |
| ATOM | 2891 | CA | VAL | E | 433 | 13.345 | 7.396 | 9.556 | 1.00 | 59.96 | C |
| ATOM | 2892 | CA | ILE | E | 434 | 15.296 | 6.063 | 12.539 | 1.00 | 60.90 | C |
| ATOM | 2893 | CA | ALA | E | 435 | 13.450 | 6.271 | 15.858 | 1.00 | 61.91 | C |
| ATOM | 2894 | CA | TRP | E | 436 | 13.877 | 4.932 | 19.377 | 1.00 | 63.05 | C |

| | | | | | | | | | | | |
|---|---|---|---|---|---|---|---|---|---|---|---|
| ATOM | 2895 | CA | ASN | E | 437 | 12.117 | 4.969 | 22.733 | 1.00 | 66.14 | C |
| ATOM | 2896 | CA | SER | E | 438 | 13.738 | 7.586 | 24.973 | 1.00 | 70.44 | C |
| ATOM | 2897 | CA | ASN | E | 439 | 11.595 | 7.279 | 28.132 | 1.00 | 74.08 | C |
| ATOM | 2898 | CA | ASN | E | 440 | 14.740 | 6.898 | 30.256 | 1.00 | 76.65 | C |
| ATOM | 2899 | CA | LEU | E | 441 | 16.069 | 10.235 | 28.925 | 1.00 | 74.28 | C |
| ATOM | 2900 | CA | ASP | E | 442 | 13.072 | 12.477 | 28.230 | 1.00 | 73.73 | C |
| ATOM | 2901 | CA | SER | E | 443 | 10.607 | 11.604 | 31.012 | 1.00 | 76.40 | C |
| ATOM | 2902 | CA | LYS | E | 444 | 10.245 | 13.396 | 34.349 | 1.00 | 80.45 | C |
| ATOM | 2903 | CA | VAL | E | 445 | 7.971 | 12.554 | 37.285 | 1.00 | 84.11 | C |
| ATOM | 2904 | CA | GLY | E | 446 | 5.761 | 15.603 | 37.025 | 1.00 | 84.14 | C |
| ATOM | 2905 | CA | GLY | E | 447 | 5.954 | 15.583 | 33.241 | 1.00 | 81.36 | C |
| ATOM | 2906 | CA | ASN | E | 448 | 8.667 | 17.013 | 31.000 | 1.00 | 78.39 | C |
| ATOM | 2907 | CA | TYR | E | 449 | 7.013 | 20.067 | 29.441 | 1.00 | 80.57 | C |
| ATOM | 2908 | CA | ASN | E | 450 | 10.012 | 21.229 | 27.396 | 1.00 | 75.75 | C |
| ATOM | 2909 | CA | TYR | E | 451 | 9.072 | 19.188 | 24.313 | 1.00 | 69.08 | C |
| ATOM | 2910 | CA | ARG | E | 452 | 6.251 | 20.882 | 22.393 | 1.00 | 69.14 | C |
| ATOM | 2911 | CA | TYR | E | 453 | 4.316 | 20.344 | 19.170 | 1.00 | 66.36 | C |
| ATOM | 2912 | CA | ARG | E | 454 | 2.173 | 22.564 | 16.974 | 1.00 | 65.78 | C |
| ATOM | 2913 | CA | LEU | E | 455 | -1.495 | 21.661 | 17.298 | 1.00 | 66.20 | C |
| ATOM | 2914 | CA | PHE | E | 456 | -3.119 | 24.347 | 15.091 | 1.00 | 68.18 | C |
| ATOM | 2915 | CA | ARG | E | 457 | -2.301 | 26.112 | 11.806 | 1.00 | 68.34 | C |
| ATOM | 2916 | CA | LYS | E | 458 | -4.213 | 27.812 | 8.980 | 1.00 | 71.15 | C |
| ATOM | 2917 | CA | SER | E | 459 | -2.840 | 25.414 | 6.347 | 1.00 | 69.30 | C |
| ATOM | 2918 | CA | ASN | E | 460 | -0.684 | 22.328 | 6.082 | 1.00 | 68.54 | C |
| ATOM | 2919 | CA | LEU | E | 461 | 3.104 | 22.591 | 6.189 | 1.00 | 66.75 | C |
| ATOM | 2920 | CA | LYS | E | 462 | 4.804 | 22.133 | 2.809 | 1.00 | 68.01 | C |
| ATOM | 2921 | CA | PRO | E | 463 | 7.671 | 19.540 | 2.888 | 1.00 | 66.29 | C |
| ATOM | 2922 | CA | PHE | E | 464 | 10.619 | 20.502 | 5.157 | 1.00 | 64.05 | C |
| ATOM | 2923 | CA | GLU | E | 465 | 8.919 | 23.815 | 6.107 | 1.00 | 69.66 | C |
| ATOM | 2924 | CA | ARG | E | 466 | 9.765 | 24.765 | 9.689 | 1.00 | 67.60 | C |
| ATOM | 2925 | CA | ASP | E | 467 | 7.234 | 26.925 | 11.535 | 1.00 | 69.23 | C |
| ATOM | 2926 | CA | ILE | E | 468 | 8.321 | 28.378 | 14.886 | 1.00 | 68.63 | C |
| ATOM | 2927 | CA | SER | E | 469 | 5.696 | 31.082 | 15.362 | 1.00 | 71.47 | C |
| ATOM | 2928 | CA | THR | E | 470 | 3.332 | 31.374 | 18.343 | 1.00 | 75.84 | C |
| ATOM | 2929 | CA | GLU | E | 471 | 0.637 | 33.412 | 16.630 | 1.00 | 81.45 | C |
| ATOM | 2930 | CA | ILE | E | 472 | -2.846 | 32.850 | 18.063 | 1.00 | 80.37 | C |
| ATOM | 2931 | CA | TYR | E | 473 | -4.927 | 30.558 | 15.856 | 1.00 | 79.15 | C |
| ATOM | 2932 | CA | GLN | E | 474 | -8.296 | 31.890 | 14.703 | 1.00 | 86.84 | C |
| ATOM | 2933 | CA | ALA | E | 475 | -10.803 | 29.043 | 14.682 | 1.00 | 87.45 | C |
| ATOM | 2934 | CA | GLY | E | 476 | -13.884 | 31.105 | 13.863 | 1.00 | 92.17 | C |
| ATOM | 2935 | CA | SER | E | 477 | -14.777 | 34.140 | 11.769 | 1.00 | 95.93 | C |
| ATOM | 2936 | CA | THR | E | 478 | -13.939 | 36.710 | 14.489 | 1.00 | 95.08 | C |
| ATOM | 2937 | CA | PRO | E | 479 | -10.392 | 38.177 | 14.562 | 1.00 | 95.43 | C |
| ATOM | 2938 | CA | CYS | E | 480 | -8.446 | 37.245 | 17.683 | 1.00 | 96.85 | C |
| ATOM | 2939 | CA | ASN | E | 481 | -6.095 | 40.280 | 17.846 | 1.00 | 97.69 | C |
| ATOM | 2940 | CA | GLY | E | 482 | -3.548 | 38.359 | 19.898 | 1.00 | 94.02 | C |
| ATOM | 2941 | CA | VAL | E | 483 | -5.865 | 37.479 | 22.816 | 1.00 | 92.45 | C |
| ATOM | 2942 | CA | GLU | E | 484 | -6.455 | 33.904 | 24.006 | 1.00 | 89.19 | C |
| ATOM | 2943 | CA | GLY | E | 485 | -10.157 | 33.166 | 24.367 | 1.00 | 85.83 | C |
| ATOM | 2944 | CA | PHE | E | 486 | -13.172 | 31.666 | 22.626 | 1.00 | 85.79 | C |
| ATOM | 2945 | CA | ASN | E | 487 | -12.241 | 30.386 | 19.117 | 1.00 | 85.46 | C |
| ATOM | 2946 | CA | CYS | E | 488 | -8.741 | 31.831 | 19.684 | 1.00 | 85.66 | C |
| ATOM | 2947 | CA | TYR | E | 489 | -6.224 | 29.163 | 20.631 | 1.00 | 78.89 | C |
| ATOM | 2948 | CA | PHE | E | 490 | -2.545 | 28.874 | 21.419 | 1.00 | 74.70 | C |
| ATOM | 2949 | CA | PRO | E | 491 | -1.099 | 26.859 | 18.504 | 1.00 | 71.07 | C |
| ATOM | 2950 | CA | LEU | E | 492 | 1.496 | 24.912 | 20.510 | 1.00 | 68.60 | C |
| ATOM | 2951 | CA | GLN | E | 493 | 0.941 | 22.238 | 23.146 | 1.00 | 72.18 | C |
| ATOM | 2952 | CA | SER | E | 494 | 3.285 | 20.188 | 25.306 | 1.00 | 72.20 | C |
| ATOM | 2953 | CA | TYR | E | 495 | 3.588 | 16.438 | 25.732 | 1.00 | 71.68 | C |
| ATOM | 2954 | CA | GLY | E | 496 | 3.431 | 15.539 | 29.395 | 1.00 | 77.67 | C |

| | | | | | | | | | | | |
|---|---|---|---|---|---|---|---|---|---|---|---|
| ATOM | 2955 | CA | PHE | E | 497 | 6.082 | 12.816 | 29.350 | 1.00 | 73.54 | C |
| ATOM | 2956 | CA | GLN | E | 498 | 5.861 | 10.726 | 32.533 | 1.00 | 80.50 | C |
| ATOM | 2957 | CA | PRO | E | 499 | 7.643 | 7.416 | 33.316 | 1.00 | 80.33 | C |
| ATOM | 2958 | CA | THR | E | 500 | 4.319 | 5.730 | 34.175 | 1.00 | 82.81 | C |
| ATOM | 2959 | CA | ASN | E | 501 | 2.731 | 6.477 | 30.785 | 1.00 | 78.74 | C |
| ATOM | 2960 | CA | GLY | E | 502 | 1.875 | 3.719 | 28.357 | 1.00 | 77.32 | C |
| ATOM | 2961 | CA | VAL | E | 503 | 4.182 | 2.839 | 25.469 | 1.00 | 76.20 | C |
| ATOM | 2962 | CA | GLY | E | 504 | 2.171 | 4.844 | 22.931 | 1.00 | 74.76 | C |
| ATOM | 2963 | CA | TYR | E | 505 | 2.579 | 7.925 | 25.162 | 1.00 | 75.30 | C |
| ATOM | 2964 | CA | GLN | E | 506 | 6.255 | 7.469 | 25.973 | 1.00 | 73.43 | C |
| ATOM | 2965 | CA | PRO | E | 507 | 8.742 | 9.814 | 24.263 | 1.00 | 69.15 | C |
| ATOM | 2966 | CA | TYR | E | 508 | 10.512 | 8.654 | 21.135 | 1.00 | 65.82 | C |
| ATOM | 2967 | CA | ARG | E | 509 | 13.495 | 10.443 | 19.658 | 1.00 | 63.39 | C |
| ATOM | 2968 | CA | VAL | E | 510 | 13.391 | 10.523 | 15.865 | 1.00 | 59.87 | C |
| ATOM | 2969 | CA | VAL | E | 511 | 16.270 | 11.147 | 13.471 | 1.00 | 59.14 | C |
| ATOM | 2970 | CA | VAL | E | 512 | 15.335 | 11.393 | 9.808 | 1.00 | 58.86 | C |
| ATOM | 2971 | CA | LEU | E | 513 | 18.016 | 11.261 | 7.139 | 1.00 | 62.62 | C |
| ATOM | 2972 | CA | SER | E | 514 | 17.216 | 12.876 | 3.810 | 1.00 | 67.97 | C |
| ATOM | 2973 | CA | PHE | E | 515 | 19.463 | 11.823 | 0.933 | 1.00 | 70.05 | C |
| ATOM | 2974 | CA | GLU | E | 516 | 19.586 | 14.373 | -1.875 | 1.00 | 83.43 | C |
| ATOM | 2975 | CA | LEU | E | 517 | 20.198 | 13.026 | -5.396 | 1.00 | 83.46 | C |
| ATOM | 2976 | CA | LEU | E | 518 | 20.759 | 15.711 | -8.044 | 1.00 | 88.46 | C |
| ATOM | 2977 | CA | HIS | E | 519 | 23.264 | 15.793 | -10.912 | 1.00 | 93.96 | C |
| ATOM | 2978 | CA | ALA | E | 520 | 26.056 | 17.406 | -8.905 | 1.00 | 88.20 | C |
| ATOM | 2979 | CA | PRO | E | 521 | 29.272 | 16.214 | -7.183 | 1.00 | 84.21 | C |
| ATOM | 2980 | CA | ALA | E | 522 | 28.624 | 13.696 | -4.437 | 1.00 | 81.09 | C |
| ATOM | 2981 | CA | THR | E | 523 | 30.124 | 14.536 | -1.049 | 1.00 | 80.33 | C |
| ATOM | 2982 | CA | VAL | E | 524 | 28.740 | 11.800 | 1.238 | 1.00 | 75.98 | C |
| ATOM | 2983 | CA | CYS | E | 525 | 29.803 | 8.252 | 0.391 | 1.00 | 77.24 | C |
| ATOM | 2984 | CA | GLY | E | 526 | 29.688 | 4.891 | 2.130 | 1.00 | 74.86 | C |
| ATOM | 2985 | CA | PRO | E | 527 | 32.783 | 3.125 | 3.514 | 1.00 | 76.35 | C |
| ATOM | 2986 | CA | LYS | E | 528 | 33.811 | 1.461 | 0.259 | 1.00 | 79.86 | C |
| ATOM | 2987 | CA | LYS | E | 529 | 37.383 | 0.655 | -0.683 | 1.00 | 84.71 | C |
| ATOM | 2988 | CA | SER | E | 530 | 38.648 | 1.932 | -4.015 | 1.00 | 85.08 | C |
| ATOM | 2989 | CA | THR | E | 531 | 40.428 | -0.248 | -6.557 | 1.00 | 85.48 | C |
| ATOM | 2990 | CA | ASN | E | 532 | 43.030 | 0.495 | -9.202 | 1.00 | 86.85 | C |
| ATOM | 2991 | CA | LEU | E | 533 | 41.738 | 2.026 | -12.413 | 1.00 | 85.37 | C |
| ATOM | 2992 | CA | VAL | E | 534 | 41.773 | -0.408 | -15.354 | 1.00 | 85.36 | C |
| ATOM | 2993 | CA | LYS | E | 535 | 41.577 | 0.925 | -18.905 | 1.00 | 85.47 | C |
| ATOM | 2994 | CA | ASN | E | 536 | 40.908 | -0.629 | -22.335 | 1.00 | 86.91 | C |
| ATOM | 2995 | CA | LYS | E | 537 | 39.041 | -3.620 | -20.873 | 1.00 | 86.97 | C |
| ATOM | 2996 | CA | CYS | E | 538 | 35.392 | -4.507 | -20.262 | 1.00 | 88.78 | C |
| ATOM | 2997 | CA | VAL | E | 539 | 34.727 | -3.914 | -16.547 | 1.00 | 84.48 | C |
| ATOM | 2998 | CA | ASN | E | 540 | 31.866 | -3.255 | -14.154 | 1.00 | 83.40 | C |
| ATOM | 2999 | CA | PHE | E | 541 | 32.501 | 0.238 | -12.829 | 1.00 | 81.19 | C |
| ATOM | 3000 | CA | ASN | E | 542 | 31.231 | 2.390 | -9.976 | 1.00 | 81.91 | C |
| ATOM | 3001 | CA | PHE | E | 543 | 32.371 | 6.033 | -10.220 | 1.00 | 81.60 | C |
| ATOM | 3002 | CA | ASN | E | 544 | 31.136 | 7.991 | -7.154 | 1.00 | 83.48 | C |
| ATOM | 3003 | CA | GLY | E | 545 | 27.826 | 6.127 | -7.135 | 1.00 | 82.16 | C |
| ATOM | 3004 | CA | LEU | E | 546 | 27.442 | 5.951 | -10.928 | 1.00 | 81.73 | C |
| ATOM | 3005 | CA | THR | E | 547 | 27.349 | 2.279 | -11.902 | 1.00 | 80.55 | C |
| ATOM | 3006 | CA | GLY | E | 548 | 27.455 | 0.425 | -15.176 | 1.00 | 81.61 | C |
| ATOM | 3007 | CA | THR | E | 549 | 29.377 | -1.882 | -17.471 | 1.00 | 84.24 | C |
| ATOM | 3008 | CA | GLY | E | 550 | 31.799 | -0.732 | -20.138 | 1.00 | 83.16 | C |
| ATOM | 3009 | CA | VAL | E | 551 | 35.300 | -0.074 | -21.417 | 1.00 | 84.47 | C |
| ATOM | 3010 | CA | LEU | E | 552 | 37.224 | 2.866 | -19.961 | 1.00 | 83.76 | C |
| ATOM | 3011 | CA | THR | E | 553 | 39.570 | 4.754 | -22.286 | 1.00 | 85.79 | C |
| ATOM | 3012 | CA | GLU | E | 554 | 41.523 | 7.986 | -22.044 | 1.00 | 90.53 | C |
| ATOM | 3013 | CA | SER | E | 555 | 39.379 | 10.865 | -23.252 | 1.00 | 89.09 | C |
| ATOM | 3014 | CA | ASN | E | 556 | 40.210 | 13.819 | -25.483 | 1.00 | 92.52 | C |

| | | | | | | | | | | |
|---|---|---|---|---|---|---|---|---|---|---|
| ATOM | 3015 | CA | LYS | E | 557 | 37.169 | 15.794 | -24.329 | 1.00 88.65 | C |
| ATOM | 3016 | CA | LYS | E | 558 | 37.787 | 19.266 | -22.907 | 1.00 91.43 | C |
| ATOM | 3017 | CA | PHE | E | 559 | 35.771 | 19.114 | -19.707 | 1.00 87.70 | C |
| ATOM | 3018 | CA | LEU | E | 560 | 35.195 | 22.358 | -17.877 | 1.00 88.93 | C |
| ATOM | 3019 | CA | PRO | E | 561 | 36.487 | 22.454 | -14.243 | 1.00 89.10 | C |
| ATOM | 3020 | CA | PHE | E | 562 | 32.969 | 21.935 | -12.797 | 1.00 89.64 | C |
| ATOM | 3021 | CA | GLN | E | 563 | 31.837 | 19.054 | -15.022 | 1.00 88.40 | C |
| ATOM | 3022 | CA | GLN | E | 564 | 31.878 | 15.624 | -13.382 | 1.00 88.98 | C |
| ATOM | 3023 | CA | PHE | E | 565 | 30.615 | 13.373 | -16.188 | 1.00 88.07 | C |
| ATOM | 3024 | CA | GLY | E | 566 | 29.477 | 13.577 | -19.782 | 1.00 88.94 | C |
| ATOM | 3025 | CA | ARG | E | 567 | 26.634 | 12.436 | -21.984 | 1.00 92.85 | C |
| ATOM | 3026 | CA | ASP | E | 568 | 26.292 | 11.539 | -25.655 | 1.00100.06 | C |
| ATOM | 3027 | CA | ILE | E | 569 | 23.776 | 12.682 | -28.290 | 1.00100.05 | C |
| ATOM | 3028 | CA | ALA | E | 570 | 21.614 | 9.781 | -27.006 | 1.00 98.16 | C |
| ATOM | 3029 | CA | ASP | E | 571 | 21.552 | 11.389 | -23.503 | 1.00 97.40 | C |
| ATOM | 3030 | CA | THR | E | 572 | 23.353 | 8.382 | -22.007 | 1.00 97.98 | C |
| ATOM | 3031 | CA | THR | E | 573 | 26.520 | 8.601 | -19.921 | 1.00 91.84 | C |
| ATOM | 3032 | CA | ASP | E | 574 | 29.664 | 8.454 | -22.063 | 1.00 91.38 | C |
| ATOM | 3033 | CA | ALA | E | 575 | 32.431 | 10.058 | -19.986 | 1.00 86.95 | C |
| ATOM | 3034 | CA | VAL | E | 576 | 33.310 | 10.402 | -16.324 | 1.00 84.40 | C |
| ATOM | 3035 | CA | ARG | E | 577 | 35.807 | 12.390 | -14.260 | 1.00 84.67 | C |
| ATOM | 3036 | CA | ASP | E | 578 | 37.650 | 10.079 | -11.863 | 1.00 84.82 | C |
| ATOM | 3037 | CA | PRO | E | 579 | 37.027 | 11.108 | -8.215 | 1.00 84.59 | C |
| ATOM | 3038 | CA | GLN | E | 580 | 40.672 | 10.702 | -7.113 | 1.00 88.63 | C |
| ATOM | 3039 | CA | THR | E | 581 | 42.709 | 11.967 | -10.085 | 1.00 87.64 | C |
| ATOM | 3040 | CA | LEU | E | 582 | 41.018 | 14.576 | -12.261 | 1.00 90.01 | C |
| ATOM | 3041 | CA | GLU | E | 583 | 41.366 | 12.519 | -15.437 | 1.00 89.05 | C |
| ATOM | 3042 | CA | ILE | E | 584 | 38.596 | 12.472 | -18.029 | 1.00 86.36 | C |
| ATOM | 3043 | CA | LEU | E | 585 | 37.667 | 8.960 | -19.137 | 1.00 84.06 | C |
| ATOM | 3044 | CA | ASP | E | 586 | 35.487 | 7.891 | -22.043 | 1.00 88.04 | C |
| ATOM | 3045 | CA | ILE | E | 587 | 33.056 | 5.055 | -21.414 | 1.00 84.07 | C |
| ATOM | 3046 | CA | THR | E | 588 | 32.133 | 2.843 | -24.305 | 1.00 84.83 | C |
| ATOM | 3047 | CA | PRO | E | 589 | 29.798 | -0.153 | -24.028 | 1.00 84.73 | C |
| ATOM | 3048 | CA | CYS | E | 590 | 31.306 | -3.605 | -24.024 | 1.00 88.56 | C |
| ATOM | 3049 | CA | SER | E | 591 | 31.226 | -5.147 | -27.502 | 1.00 86.47 | C |
| ATOM | 3050 | CA | PHE | E | 592 | 27.877 | -6.583 | -28.508 | 1.00 83.84 | C |
| ATOM | 3051 | CA | GLY | E | 593 | 25.965 | -7.455 | -31.624 | 1.00 79.67 | C |
| ATOM | 3052 | CA | GLY | E | 594 | 23.478 | -9.808 | -33.142 | 1.00 76.33 | C |
| ATOM | 3053 | CA | VAL | E | 595 | 24.385 | -13.439 | -33.725 | 1.00 75.22 | C |
| ATOM | 3054 | CA | SER | E | 596 | 23.119 | -14.952 | -36.953 | 1.00 73.12 | C |
| ATOM | 3055 | CA | VAL | E | 597 | 23.517 | -18.593 | -37.943 | 1.00 71.99 | C |
| ATOM | 3056 | CA | ILE | E | 598 | 24.113 | -19.341 | -41.622 | 1.00 73.22 | C |
| ATOM | 3057 | CA | THR | E | 599 | 23.092 | -22.818 | -42.529 | 1.00 76.27 | C |
| ATOM | 3058 | CA | PRO | E | 600 | 22.197 | -25.200 | -45.330 | 1.00 80.81 | C |
| ATOM | 3059 | CA | GLY | E | 601 | 19.085 | -27.176 | -44.509 | 1.00 87.01 | C |
| ATOM | 3060 | CA | THR | E | 602 | 19.141 | -30.181 | -42.204 | 1.00 91.58 | C |
| ATOM | 3061 | CA | ASN | E | 603 | 18.324 | -32.346 | -45.228 | 1.00102.62 | C |
| ATOM | 3062 | CA | THR | E | 604 | 21.863 | -31.665 | -46.514 | 1.00 93.50 | C |
| ATOM | 3063 | CA | SER | E | 605 | 24.244 | -30.917 | -43.628 | 1.00 86.73 | C |
| ATOM | 3064 | CA | ASN | E | 606 | 24.497 | -30.032 | -39.945 | 1.00 83.60 | C |
| ATOM | 3065 | CA | GLN | E | 607 | 27.461 | -27.708 | -40.579 | 1.00 81.93 | C |
| ATOM | 3066 | CA | VAL | E | 608 | 26.861 | -24.027 | -39.778 | 1.00 75.11 | C |
| ATOM | 3067 | CA | ALA | E | 609 | 28.639 | -20.674 | -39.974 | 1.00 72.78 | C |
| ATOM | 3068 | CA | VAL | E | 610 | 28.149 | -17.828 | -37.497 | 1.00 71.62 | C |
| ATOM | 3069 | CA | LEU | E | 611 | 27.989 | -14.112 | -38.275 | 1.00 73.10 | C |
| ATOM | 3070 | CA | TYR | E | 612 | 28.722 | -11.716 | -35.418 | 1.00 77.33 | C |
| ATOM | 3071 | CA | GLN | E | 613 | 26.957 | -8.638 | -36.751 | 1.00 76.27 | C |
| ATOM | 3072 | CA | GLY | E | 614 | 28.984 | -5.413 | -36.586 | 1.00 79.70 | C |
| ATOM | 3073 | CA | VAL | E | 615 | 31.672 | -7.018 | -34.409 | 1.00 83.89 | C |
| ATOM | 3074 | CA | ASN | E | 616 | 35.425 | -7.228 | -34.996 | 1.00 93.42 | C |

| | | | | | | | | | | | |
|---|---|---|---|---|---|---|---|---|---|---|---|
| ATOM | 3075 | CA | CYS | E | 617 | 37.096 | -10.639 | -35.286 | 1.00 | 93.38 | C |
| ATOM | 3076 | CA | THR | E | 618 | 39.525 | -9.683 | -32.483 | 1.00 | 94.26 | C |
| ATOM | 3077 | CA | GLU | E | 619 | 36.732 | -9.323 | -29.898 | 1.00 | 93.97 | C |
| ATOM | 3078 | CA | VAL | E | 620 | 34.718 | -12.545 | -30.366 | 1.00 | 90.55 | C |
| ATOM | 3079 | CA | PRO | E | 621 | 36.914 | -14.991 | -28.448 | 1.00 | 96.24 | C |
| ATOM | 3080 | CA | ASN | E | 641 | 38.049 | -19.901 | -41.123 | 1.00 | 82.16 | C |
| ATOM | 3081 | CA | VAL | E | 642 | 37.620 | -16.436 | -39.593 | 1.00 | 80.01 | C |
| ATOM | 3082 | CA | PHE | E | 643 | 36.675 | -13.821 | -42.210 | 1.00 | 81.95 | C |
| ATOM | 3083 | CA | GLN | E | 644 | 36.165 | -10.134 | -41.389 | 1.00 | 85.45 | C |
| ATOM | 3084 | CA | THR | E | 645 | 33.646 | -8.357 | -43.611 | 1.00 | 82.32 | C |
| ATOM | 3085 | CA | ARG | E | 646 | 31.613 | -5.148 | -43.455 | 1.00 | 80.53 | C |
| ATOM | 3086 | CA | ALA | E | 647 | 28.604 | -7.087 | -42.156 | 1.00 | 76.16 | C |
| ATOM | 3087 | CA | GLY | E | 648 | 30.613 | -8.441 | -39.220 | 1.00 | 80.32 | C |
| ATOM | 3088 | CA | CYS | E | 649 | 32.898 | -11.305 | -38.298 | 1.00 | 81.85 | C |
| ATOM | 3089 | CA | LEU | E | 650 | 31.980 | -14.533 | -40.120 | 1.00 | 75.98 | C |
| ATOM | 3090 | CA | ILE | E | 651 | 33.356 | -17.702 | -38.497 | 1.00 | 74.95 | C |
| ATOM | 3091 | CA | GLY | E | 652 | 33.139 | -21.161 | -40.064 | 1.00 | 77.81 | C |
| ATOM | 3092 | CA | ALA | E | 653 | 32.858 | -20.133 | -43.731 | 1.00 | 79.22 | C |
| ATOM | 3093 | CA | GLU | E | 654 | 35.884 | -20.357 | -46.021 | 1.00 | 82.16 | C |
| ATOM | 3094 | CA | HIS | E | 655 | 36.463 | -17.156 | -47.983 | 1.00 | 83.49 | C |
| ATOM | 3095 | CA | VAL | E | 656 | 37.161 | -17.763 | -51.678 | 1.00 | 83.76 | C |
| ATOM | 3096 | CA | ASN | E | 657 | 38.151 | -15.505 | -54.567 | 1.00 | 90.17 | C |
| ATOM | 3097 | CA | ASN | E | 658 | 35.606 | -17.124 | -56.921 | 1.00 | 84.64 | C |
| ATOM | 3098 | CA | SER | E | 659 | 32.342 | -15.241 | -57.303 | 1.00 | 79.30 | C |
| ATOM | 3099 | CA | TYR | E | 660 | 29.026 | -17.049 | -57.721 | 1.00 | 79.63 | C |
| ATOM | 3100 | CA | GLU | E | 661 | 25.327 | -16.257 | -57.653 | 1.00 | 81.91 | C |
| ATOM | 3101 | CA | CYS | E | 662 | 23.977 | -15.320 | -54.231 | 1.00 | 78.98 | C |
| ATOM | 3102 | CA | ASP | E | 663 | 22.585 | -18.244 | -52.239 | 1.00 | 77.19 | C |
| ATOM | 3103 | CA | ILE | E | 664 | 22.409 | -17.319 | -48.543 | 1.00 | 72.19 | C |
| ATOM | 3104 | CA | PRO | E | 665 | 22.847 | -13.536 | -48.145 | 1.00 | 69.15 | C |
| ATOM | 3105 | CA | ILE | E | 666 | 25.366 | -12.293 | -45.596 | 1.00 | 67.43 | C |
| ATOM | 3106 | CA | GLY | E | 667 | 25.424 | -8.593 | -46.376 | 1.00 | 64.71 | C |
| ATOM | 3107 | CA | ALA | E | 668 | 27.586 | -5.885 | -48.010 | 1.00 | 64.28 | C |
| ATOM | 3108 | CA | GLY | E | 669 | 27.925 | -7.854 | -51.213 | 1.00 | 67.38 | C |
| ATOM | 3109 | CA | ILE | E | 670 | 28.868 | -11.075 | -49.412 | 1.00 | 67.96 | C |
| ATOM | 3110 | CA | CYS | E | 671 | 26.803 | -14.257 | -49.809 | 1.00 | 74.16 | C |
| ATOM | 3111 | CA | ALA | E | 672 | 27.366 | -17.786 | -48.525 | 1.00 | 73.50 | C |
| ATOM | 3112 | CA | SER | E | 673 | 26.627 | -21.243 | -49.881 | 1.00 | 78.78 | C |
| ATOM | 3113 | CA | TYR | E | 674 | 27.262 | -24.921 | -49.190 | 1.00 | 84.45 | C |
| ATOM | 3114 | CA | GLN | E | 675 | 29.573 | -26.482 | -51.790 | 1.00 | 87.82 | C |
| ATOM | 3115 | CA | GLN | E | 690 | 31.963 | -30.125 | -48.916 | 1.00 | 86.76 | C |
| ATOM | 3116 | CA | SER | E | 691 | 31.877 | -27.029 | -46.714 | 1.00 | 83.47 | C |
| ATOM | 3117 | CA | ILE | E | 692 | 30.238 | -23.634 | -46.314 | 1.00 | 79.59 | C |
| ATOM | 3118 | CA | ILE | E | 693 | 31.951 | -20.862 | -48.270 | 1.00 | 78.51 | C |
| ATOM | 3119 | CA | ALA | E | 694 | 31.575 | -17.090 | -48.414 | 1.00 | 76.77 | C |
| ATOM | 3120 | CA | TYR | E | 695 | 32.130 | -14.986 | -51.504 | 1.00 | 74.35 | C |
| ATOM | 3121 | CA | THR | E | 696 | 31.357 | -11.671 | -53.101 | 1.00 | 71.57 | C |
| ATOM | 3122 | CA | MET | E | 697 | 28.228 | -12.360 | -55.120 | 1.00 | 72.23 | C |
| ATOM | 3123 | CA | SER | E | 698 | 28.212 | -12.205 | -58.893 | 1.00 | 72.42 | C |
| ATOM | 3124 | CA | LEU | E | 699 | 25.709 | -9.916 | -60.565 | 1.00 | 65.77 | C |
| ATOM | 3125 | CA | GLY | E | 700 | 25.594 | -12.094 | -63.683 | 1.00 | 70.08 | C |
| ATOM | 3126 | CA | ALA | E | 701 | 27.723 | -13.185 | -66.583 | 1.00 | 72.68 | C |
| ATOM | 3127 | CA | GLU | E | 702 | 29.384 | -10.473 | -68.610 | 1.00 | 76.77 | C |
| ATOM | 3128 | CA | ASN | E | 703 | 28.215 | -9.979 | -72.185 | 1.00 | 73.44 | C |
| ATOM | 3129 | CA | SER | E | 704 | 29.851 | -7.299 | -74.310 | 1.00 | 74.25 | C |
| ATOM | 3130 | CA | VAL | E | 705 | 27.399 | -6.485 | -77.089 | 1.00 | 68.52 | C |
| ATOM | 3131 | CA | ALA | E | 706 | 29.028 | -6.201 | -80.527 | 1.00 | 66.76 | C |
| ATOM | 3132 | CA | TYR | E | 707 | 27.786 | -2.687 | -81.206 | 1.00 | 63.42 | C |
| ATOM | 3133 | CA | SER | E | 708 | 28.667 | -0.902 | -84.431 | 1.00 | 65.25 | C |
| ATOM | 3134 | CA | ASN | E | 709 | 27.037 | 1.790 | -86.533 | 1.00 | 68.60 | C |

| | | | | | | | | | | | |
|---|---|---|---|---|---|---|---|---|---|---|---|
| ATOM | 3135 | CA | ASN | E | 710 | 25.966 | -0.589 | -89.325 | 1.00 | 65.29 | C |
| ATOM | 3136 | CA | SER | E | 711 | 25.263 | -4.026 | -87.770 | 1.00 | 62.98 | C |
| ATOM | 3137 | CA | ILE | E | 712 | 21.892 | -5.324 | -86.577 | 1.00 | 59.35 | C |
| ATOM | 3138 | CA | ALA | E | 713 | 21.051 | -8.709 | -85.114 | 1.00 | 58.55 | C |
| ATOM | 3139 | CA | ILE | E | 714 | 17.697 | -10.222 | -86.143 | 1.00 | 59.95 | C |
| ATOM | 3140 | CA | PRO | E | 715 | 16.213 | -13.469 | -84.785 | 1.00 | 59.78 | C |
| ATOM | 3141 | CA | THR | E | 716 | 15.846 | -16.302 | -87.276 | 1.00 | 64.76 | C |
| ATOM | 3142 | CA | ASN | E | 717 | 13.920 | -18.569 | -84.906 | 1.00 | 66.89 | C |
| ATOM | 3143 | CA | PHE | E | 718 | 11.973 | -18.247 | -81.671 | 1.00 | 58.66 | C |
| ATOM | 3144 | CA | THR | E | 719 | 10.739 | -20.165 | -78.660 | 1.00 | 56.36 | C |
| ATOM | 3145 | CA | ILE | E | 720 | 7.463 | -20.139 | -76.769 | 1.00 | 54.36 | C |
| ATOM | 3146 | CA | SER | E | 721 | 8.049 | -19.673 | -73.050 | 1.00 | 52.37 | C |
| ATOM | 3147 | CA | VAL | E | 722 | 5.648 | -20.177 | -70.151 | 1.00 | 52.21 | C |
| ATOM | 3148 | CA | THR | E | 723 | 6.571 | -18.301 | -66.971 | 1.00 | 54.19 | C |
| ATOM | 3149 | CA | THR | E | 724 | 4.900 | -18.983 | -63.623 | 1.00 | 57.32 | C |
| ATOM | 3150 | CA | GLU | E | 725 | 3.922 | -15.994 | -61.475 | 1.00 | 53.86 | C |
| ATOM | 3151 | CA | ILE | E | 726 | 2.406 | -16.658 | -58.040 | 1.00 | 52.87 | C |
| ATOM | 3152 | CA | LEU | E | 727 | 0.468 | -13.927 | -56.243 | 1.00 | 51.10 | C |
| ATOM | 3153 | CA | PRO | E | 728 | -1.363 | -13.902 | -52.900 | 1.00 | 50.09 | C |
| ATOM | 3154 | CA | VAL | E | 729 | -5.014 | -12.912 | -53.218 | 1.00 | 52.90 | C |
| ATOM | 3155 | CA | SER | E | 730 | -6.536 | -13.495 | -49.799 | 1.00 | 57.36 | C |
| ATOM | 3156 | CA | MET | E | 731 | -5.696 | -14.100 | -46.168 | 1.00 | 62.01 | C |
| ATOM | 3157 | CA | THR | E | 732 | -7.306 | -16.418 | -43.613 | 1.00 | 63.31 | C |
| ATOM | 3158 | CA | LYS | E | 733 | -10.361 | -14.652 | -42.189 | 1.00 | 61.63 | C |
| ATOM | 3159 | CA | THR | E | 734 | -9.741 | -14.860 | -38.492 | 1.00 | 65.40 | C |
| ATOM | 3160 | CA | SER | E | 735 | -11.932 | -13.359 | -35.800 | 1.00 | 70.93 | C |
| ATOM | 3161 | CA | VAL | E | 736 | -11.442 | -12.876 | -32.076 | 1.00 | 71.18 | C |
| ATOM | 3162 | CA | ASP | E | 737 | -14.047 | -12.160 | -29.434 | 1.00 | 74.70 | C |
| ATOM | 3163 | CA | CYS | E | 738 | -12.262 | -9.511 | -27.365 | 1.00 | 73.05 | C |
| ATOM | 3164 | CA | THR | E | 739 | -14.208 | -10.210 | -24.176 | 1.00 | 71.38 | C |
| ATOM | 3165 | CA | MET | E | 740 | -13.832 | -13.979 | -24.625 | 1.00 | 72.69 | C |
| ATOM | 3166 | CA | TYR | E | 741 | -10.081 | -13.700 | -25.204 | 1.00 | 63.91 | C |
| ATOM | 3167 | CA | ILE | E | 742 | -9.410 | -11.299 | -22.336 | 1.00 | 64.91 | C |
| ATOM | 3168 | CA | CYS | E | 743 | -12.105 | -12.240 | -19.807 | 1.00 | 72.18 | C |
| ATOM | 3169 | CA | GLY | E | 744 | -13.343 | -15.738 | -20.724 | 1.00 | 75.01 | C |
| ATOM | 3170 | CA | ASP | E | 745 | -16.417 | -16.895 | -18.733 | 1.00 | 81.92 | C |
| ATOM | 3171 | CA | SER | E | 746 | -15.816 | -14.242 | -16.041 | 1.00 | 74.47 | C |
| ATOM | 3172 | CA | THR | E | 747 | -18.381 | -11.608 | -15.058 | 1.00 | 74.57 | C |
| ATOM | 3173 | CA | GLU | E | 748 | -15.919 | -9.812 | -12.737 | 1.00 | 74.93 | C |
| ATOM | 3174 | CA | CYS | E | 749 | -13.264 | -9.457 | -15.437 | 1.00 | 72.58 | C |
| ATOM | 3175 | CA | SER | E | 750 | -15.855 | -8.334 | -18.007 | 1.00 | 71.11 | C |
| ATOM | 3176 | CA | ASN | E | 751 | -17.109 | -5.650 | -15.616 | 1.00 | 69.67 | C |
| ATOM | 3177 | CA | LEU | E | 752 | -13.540 | -4.465 | -15.024 | 1.00 | 67.35 | C |
| ATOM | 3178 | CA | LEU | E | 753 | -13.069 | -4.540 | -18.826 | 1.00 | 65.75 | C |
| ATOM | 3179 | CA | LEU | E | 754 | -15.976 | -2.115 | -19.211 | 1.00 | 67.25 | C |
| ATOM | 3180 | CA | GLN | E | 755 | -13.774 | 0.627 | -17.680 | 1.00 | 66.36 | C |
| ATOM | 3181 | CA | TYR | E | 756 | -11.711 | 0.637 | -20.907 | 1.00 | 64.86 | C |
| ATOM | 3182 | CA | GLY | E | 757 | -14.703 | 1.739 | -22.987 | 1.00 | 67.69 | C |
| ATOM | 3183 | CA | SER | E | 758 | -14.958 | 0.730 | -26.637 | 1.00 | 69.66 | C |
| ATOM | 3184 | CA | PHE | E | 759 | -11.529 | -0.952 | -27.182 | 1.00 | 67.75 | C |
| ATOM | 3185 | CA | CYS | E | 760 | -13.093 | -4.376 | -27.815 | 1.00 | 70.80 | C |
| ATOM | 3186 | CA | THR | E | 761 | -15.664 | -2.906 | -30.214 | 1.00 | 66.66 | C |
| ATOM | 3187 | CA | GLN | E | 762 | -13.063 | -1.208 | -32.408 | 1.00 | 65.68 | C |
| ATOM | 3188 | CA | LEU | E | 763 | -10.847 | -4.321 | -32.390 | 1.00 | 63.93 | C |
| ATOM | 3189 | CA | ASN | E | 764 | -13.796 | -6.488 | -33.483 | 1.00 | 65.95 | C |
| ATOM | 3190 | CA | ARG | E | 765 | -14.660 | -3.892 | -36.131 | 1.00 | 64.41 | C |
| ATOM | 3191 | CA | ALA | E | 766 | -11.080 | -3.887 | -37.445 | 1.00 | 57.97 | C |
| ATOM | 3192 | CA | LEU | E | 767 | -11.040 | -7.685 | -37.770 | 1.00 | 58.17 | C |
| ATOM | 3193 | CA | THR | E | 768 | -14.482 | -7.667 | -39.425 | 1.00 | 60.22 | C |
| ATOM | 3194 | CA | GLY | E | 769 | -13.201 | -5.079 | -41.899 | 1.00 | 58.10 | C |

| | | | | | | | | | | |
|---|---|---|---|---|---|---|---|---|---|---|
| ATOM | 3195 | CA | ILE | E | 770 | -10.210 | -7.301 | -42.680 | 1.00 56.46 | C |
| ATOM | 3196 | CA | ALA | E | 771 | -12.483 | -10.324 | -43.315 | 1.00 58.03 | C |
| ATOM | 3197 | CA | VAL | E | 772 | -14.728 | -8.262 | -45.614 | 1.00 58.51 | C |
| ATOM | 3198 | CA | GLU | E | 773 | -11.632 | -6.975 | -47.421 | 1.00 60.59 | C |
| ATOM | 3199 | CA | GLN | E | 774 | -10.430 | -10.557 | -48.013 | 1.00 57.83 | C |
| ATOM | 3200 | CA | ASP | E | 775 | -13.747 | -11.455 | -49.631 | 1.00 60.45 | C |
| ATOM | 3201 | CA | LYS | E | 776 | -13.588 | -8.272 | -51.727 | 1.00 58.48 | C |
| ATOM | 3202 | CA | ASN | E | 777 | -10.022 | -9.139 | -52.794 | 1.00 55.81 | C |
| ATOM | 3203 | CA | THR | E | 778 | -11.075 | -12.612 | -53.934 | 1.00 55.14 | C |
| ATOM | 3204 | CA | GLN | E | 779 | -14.101 | -11.205 | -55.786 | 1.00 59.08 | C |
| ATOM | 3205 | CA | GLU | E | 780 | -12.011 | -8.556 | -57.567 | 1.00 59.89 | C |
| ATOM | 3206 | CA | VAL | E | 781 | -9.364 | -11.047 | -58.708 | 1.00 52.28 | C |
| ATOM | 3207 | CA | PHE | E | 782 | -11.597 | -13.888 | -59.873 | 1.00 52.98 | C |
| ATOM | 3208 | CA | ALA | E | 783 | -15.131 | -12.594 | -60.561 | 1.00 55.56 | C |
| ATOM | 3209 | CA | GLN | E | 784 | -14.132 | -10.598 | -63.635 | 1.00 58.04 | C |
| ATOM | 3210 | CA | VAL | E | 785 | -16.256 | -12.559 | -66.132 | 1.00 63.49 | C |
| ATOM | 3211 | CA | LYS | E | 786 | -19.988 | -11.956 | -66.581 | 1.00 72.80 | C |
| ATOM | 3212 | CA | GLN | E | 787 | -20.789 | -15.420 | -67.997 | 1.00 70.24 | C |
| ATOM | 3213 | CA | ILE | E | 788 | -19.567 | -18.943 | -67.250 | 1.00 67.69 | C |
| ATOM | 3214 | CA | TYR | E | 789 | -17.902 | -20.054 | -70.484 | 1.00 63.27 | C |
| ATOM | 3215 | CA | LYS | E | 790 | -17.316 | -23.695 | -71.411 | 1.00 70.02 | C |
| ATOM | 3216 | CA | THR | E | 791 | -15.043 | -25.336 | -73.963 | 1.00 71.40 | C |
| ATOM | 3217 | CA | PRO | E | 792 | -16.505 | -27.625 | -76.669 | 1.00 74.63 | C |
| ATOM | 3218 | CA | PRO | E | 793 | -16.449 | -31.426 | -76.223 | 1.00 77.58 | C |
| ATOM | 3219 | CA | ILE | E | 794 | -13.994 | -31.759 | -79.140 | 1.00 78.37 | C |
| ATOM | 3220 | CA | LYS | E | 795 | -10.825 | -29.961 | -78.068 | 1.00 75.22 | C |
| ATOM | 3221 | CA | ASP | E | 796 | -9.384 | -29.420 | -81.549 | 1.00 73.52 | C |
| ATOM | 3222 | CA | PHE | E | 797 | -7.379 | -26.211 | -81.117 | 1.00 64.56 | C |
| ATOM | 3223 | CA | GLY | E | 798 | -5.087 | -26.511 | -84.122 | 1.00 61.90 | C |
| ATOM | 3224 | CA | GLY | E | 799 | -2.570 | -28.691 | -82.295 | 1.00 59.43 | C |
| ATOM | 3225 | CA | PHE | E | 800 | -2.538 | -26.671 | -79.064 | 1.00 59.10 | C |
| ATOM | 3226 | CA | ASN | E | 801 | -3.101 | -28.911 | -76.034 | 1.00 65.71 | C |
| ATOM | 3227 | CA | PHE | E | 802 | -4.758 | -27.371 | -72.964 | 1.00 60.59 | C |
| ATOM | 3228 | CA | SER | E | 803 | -5.426 | -30.538 | -70.902 | 1.00 66.90 | C |
| ATOM | 3229 | CA | GLN | E | 804 | -2.970 | -29.392 | -68.213 | 1.00 68.17 | C |
| ATOM | 3230 | CA | ILE | E | 805 | -4.836 | -26.158 | -67.543 | 1.00 61.81 | C |
| ATOM | 3231 | CA | LEU | E | 806 | -8.475 | -27.234 | -68.304 | 1.00 64.06 | C |
| ATOM | 3232 | CA | PRO | E | 807 | -10.559 | -29.217 | -65.765 | 1.00 70.49 | C |
| ATOM | 3233 | CA | ASP | E | 808 | -10.480 | -33.029 | -65.880 | 1.00 83.78 | C |
| ATOM | 3234 | CA | PRO | E | 809 | -13.926 | -34.245 | -64.626 | 1.00 87.00 | C |
| ATOM | 3235 | CA | SER | E | 810 | -12.652 | -37.768 | -63.744 | 1.00 90.84 | C |
| ATOM | 3236 | CA | LYS | E | 811 | -12.403 | -36.720 | -60.064 | 1.00 88.08 | C |
| ATOM | 3237 | CA | PRO | E | 812 | -15.025 | -35.197 | -57.662 | 1.00 90.99 | C |
| ATOM | 3238 | CA | SER | E | 813 | -13.497 | -31.687 | -57.555 | 1.00 87.66 | C |
| ATOM | 3239 | CA | LYS | E | 814 | -13.338 | -30.309 | -61.090 | 1.00 83.55 | C |
| ATOM | 3240 | CA | ARG | E | 815 | -9.832 | -28.889 | -60.880 | 1.00 77.58 | C |
| ATOM | 3241 | CA | SER | E | 816 | -7.106 | -29.118 | -63.488 | 1.00 71.20 | C |
| ATOM | 3242 | CA | PRO | E | 817 | -3.880 | -31.180 | -62.874 | 1.00 70.19 | C |
| ATOM | 3243 | CA | ILE | E | 818 | -1.650 | -28.077 | -62.409 | 1.00 69.99 | C |
| ATOM | 3244 | CA | GLU | E | 819 | -4.211 | -26.727 | -59.922 | 1.00 69.38 | C |
| ATOM | 3245 | CA | ASP | E | 820 | -4.099 | -30.108 | -58.140 | 1.00 73.85 | C |
| ATOM | 3246 | CA | LEU | E | 821 | -0.327 | -29.679 | -57.864 | 1.00 73.77 | C |
| ATOM | 3247 | CA | LEU | E | 822 | -0.779 | -26.179 | -56.442 | 1.00 70.84 | C |
| ATOM | 3248 | CA | PHE | E | 823 | -3.296 | -27.372 | -53.847 | 1.00 72.31 | C |
| ATOM | 3249 | CA | ASN | E | 824 | -1.244 | -30.434 | -52.843 | 1.00 75.03 | C |
| ATOM | 3250 | CA | LYS | E | 825 | 1.952 | -28.412 | -52.352 | 1.00 79.83 | C |
| ATOM | 3251 | CA | VAL | E | 826 | 0.220 | -25.753 | -50.227 | 1.00 75.83 | C |
| ATOM | 3252 | CA | LEU | E | 849 | -3.669 | -27.081 | -34.312 | 1.00103.73 | C |
| ATOM | 3253 | CA | ILE | E | 850 | -7.049 | -25.597 | -35.266 | 1.00106.35 | C |
| ATOM | 3254 | CA | CYS | E | 851 | -8.702 | -26.792 | -32.025 | 1.00107.52 | C |

ATOM 3255 CA ALA E 852 -5.983 -25.306 -29.779 1.00104.35 C
ATOM 3256 CA GLN E 853 -6.420 -21.906 -31.420 1.00 98.54 C
ATOM 3257 CA LYS E 854 -10.233 -22.124 -31.641 1.00 99.44 C
ATOM 3258 CA PHE E 855 -10.729 -22.926 -27.919 1.00 99.14 C
ATOM 3259 CA ASN E 856 -9.682 -19.391 -26.809 1.00 89.16 C
ATOM 3260 CA GLY E 857 -12.385 -17.496 -28.711 1.00 81.26 C
ATOM 3261 CA LEU E 858 -10.426 -17.367 -31.986 1.00 79.92 C
ATOM 3262 CA THR E 859 -12.667 -18.516 -34.805 1.00 71.68 C
ATOM 3263 CA VAL E 860 -11.982 -18.739 -38.528 1.00 65.18 C
ATOM 3264 CA LEU E 861 -14.654 -17.728 -41.000 1.00 63.30 C
ATOM 3265 CA PRO E 862 -14.942 -19.690 -44.255 1.00 64.09 C
ATOM 3266 CA PRO E 863 -14.277 -17.786 -47.494 1.00 61.44 C
ATOM 3267 CA LEU E 864 -17.349 -16.637 -49.384 1.00 61.85 C
ATOM 3268 CA LEU E 865 -16.076 -18.192 -52.605 1.00 60.71 C
ATOM 3269 CA THR E 866 -15.267 -21.879 -52.218 1.00 62.32 C
ATOM 3270 CA ASP E 867 -12.352 -23.581 -53.964 1.00 66.65 C
ATOM 3271 CA GLU E 868 -14.835 -25.138 -56.403 1.00 67.16 C
ATOM 3272 CA MET E 869 -16.141 -21.660 -57.237 1.00 63.37 C
ATOM 3273 CA ILE E 870 -12.636 -20.292 -57.788 1.00 59.87 C
ATOM 3274 CA ALA E 871 -11.968 -23.258 -60.078 1.00 59.63 C
ATOM 3275 CA GLN E 872 -15.227 -22.519 -61.935 1.00 61.34 C
ATOM 3276 CA TYR E 873 -14.207 -18.866 -62.373 1.00 57.40 C
ATOM 3277 CA THR E 874 -10.773 -19.827 -63.703 1.00 56.87 C
ATOM 3278 CA SER E 875 -12.423 -22.398 -65.980 1.00 57.02 C
ATOM 3279 CA ALA E 876 -14.809 -19.740 -67.336 1.00 56.80 C
ATOM 3280 CA LEU E 877 -11.960 -17.250 -67.850 1.00 54.16 C
ATOM 3281 CA LEU E 878 -9.848 -19.911 -69.534 1.00 55.30 C
ATOM 3282 CA ALA E 879 -12.689 -21.104 -71.782 1.00 55.06 C
ATOM 3283 CA GLY E 880 -13.296 -17.506 -72.821 1.00 54.46 C
ATOM 3284 CA THR E 881 -9.594 -16.901 -73.491 1.00 54.14 C
ATOM 3285 CA ILE E 882 -9.243 -20.093 -75.575 1.00 54.70 C
ATOM 3286 CA THR E 883 -12.454 -19.817 -77.605 1.00 55.24 C
ATOM 3287 CA SER E 884 -12.914 -16.041 -77.881 1.00 53.76 C
ATOM 3288 CA GLY E 885 -9.474 -14.468 -77.317 1.00 52.97 C
ATOM 3289 CA TRP E 886 -9.789 -11.047 -75.731 1.00 51.79 C
ATOM 3290 CA THR E 887 -13.293 -10.291 -77.042 1.00 52.71 C
ATOM 3291 CA PHE E 888 -15.034 -11.699 -73.946 1.00 52.46 C
ATOM 3292 CA GLY E 889 -13.330 -9.064 -71.767 1.00 53.69 C
ATOM 3293 CA ALA E 890 -14.703 -6.070 -73.655 1.00 53.79 C
ATOM 3294 CA GLY E 891 -18.234 -7.233 -74.442 1.00 54.73 C
ATOM 3295 CA PRO E 892 -19.963 -10.314 -75.860 1.00 54.29 C
ATOM 3296 CA ALA E 893 -17.643 -13.233 -76.499 1.00 53.36 C
ATOM 3297 CA LEU E 894 -16.975 -13.657 -80.214 1.00 54.25 C
ATOM 3298 CA GLN E 895 -15.905 -17.107 -81.346 1.00 57.93 C
ATOM 3299 CA ILE E 896 -12.609 -17.378 -83.196 1.00 55.85 C
ATOM 3300 CA PRO E 897 -10.348 -20.265 -84.290 1.00 56.41 C
ATOM 3301 CA PHE E 898 -7.363 -20.444 -81.938 1.00 55.63 C
ATOM 3302 CA PRO E 899 -4.592 -20.207 -84.639 1.00 56.47 C
ATOM 3303 CA MET E 900 -6.293 -17.008 -85.871 1.00 58.22 C
ATOM 3304 CA GLN E 901 -6.329 -15.671 -82.297 1.00 55.90 C
ATOM 3305 CA MET E 902 -2.621 -16.515 -82.069 1.00 55.97 C
ATOM 3306 CA ALA E 903 -2.152 -14.657 -85.373 1.00 54.86 C
ATOM 3307 CA TYR E 904 -3.679 -11.572 -83.779 1.00 56.83 C
ATOM 3308 CA ARG E 905 -1.383 -11.938 -80.765 1.00 50.40 C
ATOM 3309 CA PHE E 906 1.627 -12.252 -83.092 1.00 51.32 C
ATOM 3310 CA ASN E 907 0.508 -9.086 -84.880 1.00 54.07 C
ATOM 3311 CA GLY E 908 0.218 -7.488 -81.442 1.00 51.09 C
ATOM 3312 CA ILE E 909 3.879 -8.201 -80.723 1.00 49.60 C
ATOM 3313 CA GLY E 910 4.902 -6.967 -84.171 1.00 52.55 C
ATOM 3314 CA VAL E 911 5.279 -10.316 -85.981 1.00 53.75 C

ATOM 3315 CA THR E 912 3.378 -11.028 -89.196 1.00 57.70 C
ATOM 3316 CA GLN E 913 0.697 -13.731 -89.447 1.00 59.38 C
ATOM 3317 CA ASN E 914 2.583 -15.954 -91.887 1.00 61.73 C
ATOM 3318 CA VAL E 915 5.326 -16.567 -89.311 1.00 58.70 C
ATOM 3319 CA LEU E 916 2.731 -18.230 -87.093 1.00 58.42 C
ATOM 3320 CA TYR E 917 0.949 -20.127 -89.876 1.00 59.52 C
ATOM 3321 CA GLU E 918 4.089 -21.491 -91.570 1.00 66.12 C
ATOM 3322 CA ASN E 919 5.391 -22.674 -88.172 1.00 62.30 C
ATOM 3323 CA GLN E 920 2.022 -23.617 -86.557 1.00 60.73 C
ATOM 3324 CA LYS E 921 3.103 -27.195 -85.776 1.00 64.43 C
ATOM 3325 CA LEU E 922 6.405 -26.075 -84.236 1.00 64.27 C
ATOM 3326 CA ILE E 923 4.663 -23.379 -82.170 1.00 60.66 C
ATOM 3327 CA ALA E 924 2.042 -25.845 -80.904 1.00 60.78 C
ATOM 3328 CA ASN E 925 4.807 -28.296 -79.953 1.00 62.83 C
ATOM 3329 CA GLN E 926 6.831 -25.579 -78.178 1.00 62.23 C
ATOM 3330 CA PHE E 927 3.736 -24.450 -76.269 1.00 60.19 C
ATOM 3331 CA ASN E 928 2.790 -27.985 -75.204 1.00 61.66 C
ATOM 3332 CA SER E 929 6.392 -28.688 -74.173 1.00 64.38 C
ATOM 3333 CA ALA E 930 6.595 -25.456 -72.145 1.00 63.01 C
ATOM 3334 CA ILE E 931 3.370 -26.373 -70.319 1.00 64.71 C
ATOM 3335 CA GLY E 932 4.860 -29.808 -69.597 1.00 69.07 C
ATOM 3336 CA LYS E 933 7.961 -28.044 -68.252 1.00 71.23 C
ATOM 3337 CA ILE E 934 5.706 -26.100 -65.870 1.00 71.40 C
ATOM 3338 CA GLN E 935 4.123 -29.386 -64.752 1.00 79.24 C
ATOM 3339 CA ASP E 936 7.547 -30.909 -64.071 1.00 82.82 C
ATOM 3340 CA SER E 937 8.846 -27.755 -62.351 1.00 81.31 C
ATOM 3341 CA LEU E 938 5.899 -27.715 -59.950 1.00 83.35 C
ATOM 3342 CA SER E 939 6.146 -31.495 -59.423 1.00 88.32 C
ATOM 3343 CA SER E 940 9.858 -31.223 -58.590 1.00 92.65 C
ATOM 3344 CA THR E 941 10.784 -27.943 -56.848 1.00 92.49 C
ATOM 3345 CA PRO E 942 9.447 -27.942 -53.253 1.00 94.88 C
ATOM 3346 CA SER E 943 9.984 -24.216 -52.544 1.00 92.48 C
ATOM 3347 CA ALA E 944 8.209 -22.949 -55.678 1.00 86.41 C
ATOM 3348 CA LEU E 945 4.963 -21.848 -53.963 1.00 75.92 C
ATOM 3349 CA GLY E 946 6.797 -20.085 -51.110 1.00 71.18 C
ATOM 3350 CA LYS E 947 4.866 -16.829 -51.625 1.00 65.81 C
ATOM 3351 CA LEU E 948 1.587 -18.454 -50.537 1.00 63.59 C
ATOM 3352 CA GLN E 949 3.115 -20.679 -47.872 1.00 68.68 C
ATOM 3353 CA ASP E 950 4.824 -17.662 -46.297 1.00 69.06 C
ATOM 3354 CA VAL E 951 1.443 -15.920 -45.933 1.00 64.92 C
ATOM 3355 CA VAL E 952 -0.078 -19.044 -44.358 1.00 65.80 C
ATOM 3356 CA ASN E 953 2.916 -19.358 -42.010 1.00 65.64 C
ATOM 3357 CA GLN E 954 2.786 -15.679 -40.985 1.00 67.00 C
ATOM 3358 CA ASN E 955 -0.892 -15.940 -40.085 1.00 66.68 C
ATOM 3359 CA ALA E 956 -0.330 -19.216 -38.216 1.00 67.22 C
ATOM 3360 CA GLN E 957 2.535 -17.635 -36.248 1.00 67.58 C
ATOM 3361 CA ALA E 958 0.425 -14.551 -35.480 1.00 65.58 C
ATOM 3362 CA LEU E 959 -2.387 -16.716 -34.103 1.00 67.02 C
ATOM 3363 CA ASN E 960 0.016 -18.937 -32.158 1.00 69.73 C
ATOM 3364 CA THR E 961 1.703 -15.865 -30.645 1.00 65.82 C
ATOM 3365 CA LEU E 962 -1.653 -14.343 -29.681 1.00 62.59 C
ATOM 3366 CA VAL E 963 -2.822 -17.576 -28.048 1.00 65.94 C
ATOM 3367 CA LYS E 964 0.522 -18.099 -26.235 1.00 65.04 C
ATOM 3368 CA GLN E 965 0.202 -14.613 -24.710 1.00 64.04 C
ATOM 3369 CA LEU E 966 -2.625 -16.005 -22.506 1.00 61.28 C
ATOM 3370 CA SER E 967 -0.010 -18.036 -20.582 1.00 64.57 C
ATOM 3371 CA SER E 968 1.828 -14.866 -19.520 1.00 62.22 C
ATOM 3372 CA ASN E 969 1.338 -13.490 -16.024 1.00 63.65 C
ATOM 3373 CA PHE E 970 2.017 -9.801 -16.914 1.00 61.67 C
ATOM 3374 CA GLY E 971 2.924 -9.298 -13.246 1.00 64.59 C

```
ATOM   3375  CA  ALA E 972      -0.122 -11.117 -11.869  1.00 64.38           C
ATOM   3376  CA  ILE E 973       0.117 -13.985  -9.392  1.00 63.27           C
ATOM   3377  CA  SER E 974      -1.218 -16.382 -12.054  1.00 64.08           C
ATOM   3378  CA  SER E 975      -1.914 -16.447 -15.782  1.00 63.76           C
ATOM   3379  CA  VAL E 976      -5.160 -18.348 -15.103  1.00 65.36           C
ATOM   3380  CA  LEU E 977      -8.104 -16.043 -14.411  1.00 67.86           C
ATOM   3381  CA  ASN E 978     -10.104 -18.833 -12.746  1.00 72.20           C
ATOM   3382  CA  ASP E 979      -7.354 -19.464 -10.180  1.00 71.46           C
ATOM   3383  CA  ILE E 980      -7.286 -15.778  -9.212  1.00 66.34           C
ATOM   3384  CA  LEU E 981     -11.067 -15.686  -8.830  1.00 66.40           C
ATOM   3385  CA  SER E 982     -11.045 -18.992  -6.924  1.00 67.15           C
ATOM   3386  CA  ARG E 983      -8.370 -17.782  -4.494  1.00 66.01           C
ATOM   3387  CA  LEU E 984      -8.962 -14.042  -4.027  1.00 66.59           C
ATOM   3388  CA  ASP E 985     -11.811 -11.749  -2.976  1.00 73.16           C
ATOM   3389  CA  PRO E 986     -12.591  -8.601  -5.126  1.00 72.17           C
ATOM   3390  CA  PRO E 987     -10.232  -6.087  -3.323  1.00 74.36           C
ATOM   3391  CA  GLU E 988      -7.168  -8.093  -4.459  1.00 74.17           C
ATOM   3392  CA  ALA E 989      -8.685  -9.957  -7.374  1.00 69.84           C
ATOM   3393  CA  GLU E 990      -9.462  -6.693  -9.197  1.00 71.58           C
ATOM   3394  CA  VAL E 991      -5.784  -5.686  -9.037  1.00 67.50           C
ATOM   3395  CA  GLN E 992      -4.639  -9.060 -10.384  1.00 65.42           C
ATOM   3396  CA  ILE E 993      -7.330  -9.039 -13.085  1.00 64.87           C
ATOM   3397  CA  ASP E 994      -6.411  -5.463 -14.108  1.00 67.32           C
ATOM   3398  CA  ARG E 995      -2.833  -6.665 -14.642  1.00 64.81           C
ATOM   3399  CA  LEU E 996      -4.102  -9.537 -16.803  1.00 60.68           C
ATOM   3400  CA  ILE E 997      -6.482  -7.234 -18.722  1.00 59.36           C
ATOM   3401  CA  THR E 998      -3.685  -4.763 -19.465  1.00 58.47           C
ATOM   3402  CA  GLY E 999      -1.358  -7.468 -20.776  1.00 58.19           C
ATOM   3403  CA  ARG E1000      -4.007  -9.351 -22.777  1.00 58.25           C
ATOM   3404  CA  LEU E1001      -5.435  -6.138 -24.242  1.00 58.14           C
ATOM   3405  CA  GLN E1002      -1.948  -5.028 -25.300  1.00 60.87           C
ATOM   3406  CA  SER E1003      -1.342  -8.433 -26.912  1.00 59.79           C
ATOM   3407  CA  LEU E1004      -4.665  -8.312 -28.773  1.00 57.78           C
ATOM   3408  CA  GLN E1005      -4.019  -4.745 -29.997  1.00 59.09           C
ATOM   3409  CA  THR E1006      -0.580  -5.840 -31.222  1.00 57.65           C
ATOM   3410  CA  TYR E1007      -2.174  -8.766 -33.084  1.00 55.23           C
ATOM   3411  CA  VAL E1008      -4.844  -6.572 -34.709  1.00 54.73           C
ATOM   3412  CA  THR E1009      -2.281  -3.946 -35.785  1.00 54.30           C
ATOM   3413  CA  GLN E1010      -0.113  -6.655 -37.385  1.00 56.16           C
ATOM   3414  CA  GLN E1011      -3.176  -8.073 -39.149  1.00 54.55           C
ATOM   3415  CA  LEU E1012      -4.128  -4.637 -40.502  1.00 52.46           C
ATOM   3416  CA  ILE E1013      -0.629  -4.075 -41.904  1.00 53.28           C
ATOM   3417  CA  ARG E1014      -0.472  -7.611 -43.331  1.00 58.44           C
ATOM   3418  CA  ALA E1015      -3.989  -7.188 -44.764  1.00 55.19           C
ATOM   3419  CA  ALA E1016      -2.900  -3.965 -46.482  1.00 53.13           C
ATOM   3420  CA  GLU E1017       0.002  -5.890 -48.043  1.00 54.83           C
ATOM   3421  CA  ILE E1018      -2.386  -8.659 -49.165  1.00 52.79           C
ATOM   3422  CA  ARG E1019      -4.758  -6.009 -50.574  1.00 53.16           C
ATOM   3423  CA  ALA E1020      -1.914  -4.521 -52.629  1.00 49.75           C
ATOM   3424  CA  SER E1021      -1.006  -7.999 -53.907  1.00 49.72           C
ATOM   3425  CA  ALA E1022      -4.663  -8.738 -54.724  1.00 49.35           C
ATOM   3426  CA  ASN E1023      -4.980  -5.438 -56.610  1.00 49.27           C
ATOM   3427  CA  LEU E1024      -1.889  -6.402 -58.599  1.00 47.19           C
ATOM   3428  CA  ALA E1025      -3.355  -9.862 -59.270  1.00 47.93           C
ATOM   3429  CA  ALA E1026      -6.652  -8.324 -60.434  1.00 48.03           C
ATOM   3430  CA  THR E1027      -4.743  -5.941 -62.711  1.00 48.39           C
ATOM   3431  CA  LYS E1028      -2.761  -8.855 -64.166  1.00 46.94           C
ATOM   3432  CA  MET E1029      -5.991 -10.834 -64.605  1.00 49.13           C
ATOM   3433  CA  SER E1030      -7.541  -7.983 -66.594  1.00 49.23           C
ATOM   3434  CA  GLU E1031      -4.433  -7.069 -68.632  1.00 48.83           C
```

```
ATOM   3435  CA  CYS E1032      -2.712 -10.458 -69.104  1.00 49.33           C
ATOM   3436  CA  VAL E1033      -5.662 -12.853 -69.441  1.00 48.04           C
ATOM   3437  CA  LEU E1034      -8.489 -10.703 -70.824  1.00 48.13           C
ATOM   3438  CA  GLY E1035      -6.183  -8.933 -73.259  1.00 46.70           C
ATOM   3439  CA  GLN E1036      -2.658  -8.689 -74.581  1.00 47.12           C
ATOM   3440  CA  SER E1037      -0.236  -6.415 -72.765  1.00 47.53           C
ATOM   3441  CA  LYS E1038       2.490  -4.220 -74.224  1.00 48.63           C
ATOM   3442  CA  ARG E1039       3.906  -3.510 -70.756  1.00 45.86           C
ATOM   3443  CA  VAL E1040       7.420  -4.958 -70.501  1.00 48.77           C
ATOM   3444  CA  ASP E1041       7.839  -7.667 -67.796  1.00 51.85           C
ATOM   3445  CA  PHE E1042       4.281  -7.152 -66.520  1.00 48.27           C
ATOM   3446  CA  CYS E1043       2.900 -10.396 -67.971  1.00 50.68           C
ATOM   3447  CA  GLY E1044       6.051 -12.526 -67.971  1.00 52.07           C
ATOM   3448  CA  LYS E1045       9.574 -12.254 -69.339  1.00 56.39           C
ATOM   3449  CA  GLY E1046       9.487 -11.800 -73.119  1.00 51.04           C
ATOM   3450  CA  TYR E1047       6.990 -10.344 -75.541  1.00 47.71           C
ATOM   3451  CA  HIS E1048       3.588 -11.148 -74.077  1.00 45.58           C
ATOM   3452  CA  LEU E1049       1.281 -13.366 -76.106  1.00 46.10           C
ATOM   3453  CA  MET E1050      -1.341 -14.582 -73.610  1.00 49.61           C
ATOM   3454  CA  SER E1051      -1.750 -15.759 -70.032  1.00 47.20           C
ATOM   3455  CA  PHE E1052      -3.744 -18.429 -68.244  1.00 48.46           C
ATOM   3456  CA  PRO E1053      -5.040 -18.170 -64.657  1.00 49.62           C
ATOM   3457  CA  GLN E1054      -4.944 -21.063 -62.208  1.00 56.42           C
ATOM   3458  CA  SER E1055      -6.544 -21.270 -58.775  1.00 61.61           C
ATOM   3459  CA  ALA E1056      -4.235 -21.727 -55.802  1.00 60.44           C
ATOM   3460  CA  PRO E1057      -4.757 -21.789 -51.998  1.00 62.56           C
ATOM   3461  CA  HIS E1058      -5.423 -18.166 -51.001  1.00 60.53           C
ATOM   3462  CA  GLY E1059      -4.014 -17.004 -54.313  1.00 55.69           C
ATOM   3463  CA  VAL E1060      -3.665 -17.084 -58.067  1.00 49.79           C
ATOM   3464  CA  VAL E1061      -0.980 -18.495 -60.347  1.00 50.87           C
ATOM   3465  CA  PHE E1062      -0.554 -17.040 -63.820  1.00 47.15           C
ATOM   3466  CA  LEU E1063       1.054 -19.028 -66.609  1.00 48.18           C
ATOM   3467  CA  HIS E1064       2.343 -16.376 -68.987  1.00 48.16           C
ATOM   3468  CA  VAL E1065       2.896 -17.457 -72.577  1.00 48.25           C
ATOM   3469  CA  THR E1066       5.530 -15.243 -74.150  1.00 49.82           C
ATOM   3470  CA  TYR E1067       7.380 -15.109 -77.439  1.00 50.66           C
ATOM   3471  CA  VAL E1068      11.148 -15.192 -76.907  1.00 53.89           C
ATOM   3472  CA  PRO E1069      13.687 -14.605 -79.716  1.00 55.07           C
ATOM   3473  CA  ALA E1070      16.058 -17.534 -80.066  1.00 62.28           C
ATOM   3474  CA  GLN E1071      18.673 -17.996 -82.820  1.00 68.46           C
ATOM   3475  CA  GLU E1072      20.937 -15.020 -83.551  1.00 71.86           C
ATOM   3476  CA  LYS E1073      22.370 -13.771 -86.863  1.00 68.85           C
ATOM   3477  CA  ASN E1074      24.415 -10.685 -87.701  1.00 70.74           C
ATOM   3478  CA  PHE E1075      23.387  -8.508 -90.642  1.00 65.15           C
ATOM   3479  CA  THR E1076      24.660  -5.282 -92.148  1.00 66.19           C
ATOM   3480  CA  THR E1077      22.139  -2.454 -92.015  1.00 63.94           C
ATOM   3481  CA  ALA E1078      21.439   1.035 -93.308  1.00 63.13           C
ATOM   3482  CA  PRO E1079      18.851   3.671 -92.342  1.00 62.44           C
ATOM   3483  CA  ALA E1080      17.783   4.232 -95.967  1.00 67.87           C
ATOM   3484  CA  ILE E1081      18.531   3.152 -99.527  1.00 71.46           C
ATOM   3485  CA  CYS E1082      19.432   5.563-102.345  1.00 81.73           C
ATOM   3486  CA  HIS E1083      17.921   4.365-105.620  1.00 84.85           C
ATOM   3487  CA  ASP E1084      17.241   7.159-108.156  1.00 86.84           C
ATOM   3488  CA  GLY E1085      18.516   9.920-105.890  1.00 85.13           C
ATOM   3489  CA  LYS E1086      15.445   9.625-103.665  1.00 79.97           C
ATOM   3490  CA  ALA E1087      15.688   8.315-100.102  1.00 74.44           C
ATOM   3491  CA  HIS E1088      13.709   5.112 -99.493  1.00 71.37           C
ATOM   3492  CA  PHE E1089      12.761   4.111 -95.948  1.00 64.86           C
ATOM   3493  CA  PRO E1090      11.135   0.837 -94.827  1.00 63.80           C
ATOM   3494  CA  ARG E1091       7.451   0.984 -93.968  1.00 71.08           C
```

```
ATOM   3495  CA  GLU E1092       7.328  -1.847 -91.421  1.00 67.56           C
ATOM   3496  CA  GLY E1093      10.741  -3.485 -91.076  1.00 62.09           C
ATOM   3497  CA  VAL E1094      14.447  -2.709 -91.539  1.00 60.25           C
ATOM   3498  CA  PHE E1095      16.888  -2.928 -94.423  1.00 62.87           C
ATOM   3499  CA  VAL E1096      19.455  -5.714 -94.037  1.00 66.75           C
ATOM   3500  CA  SER E1097      22.300  -7.178 -96.024  1.00 74.74           C
ATOM   3501  CA  ASN E1098      23.877 -10.624 -95.885  1.00 86.63           C
ATOM   3502  CA  GLY E1099      27.080  -9.117 -97.287  1.00 87.99           C
ATOM   3503  CA  THR E1100      26.144  -9.096-100.980  1.00 88.88           C
ATOM   3504  CA  HIS E1101      22.384  -8.517-101.404  1.00 84.34           C
ATOM   3505  CA  TRP E1102      19.999  -6.086 -99.695  1.00 72.98           C
ATOM   3506  CA  PHE E1103      16.573  -7.151 -98.405  1.00 70.32           C
ATOM   3507  CA  VAL E1104      13.825  -5.685 -96.220  1.00 65.43           C
ATOM   3508  CA  THR E1105      12.651  -7.741 -93.244  1.00 61.67           C
ATOM   3509  CA  GLN E1106      10.262  -7.493 -90.307  1.00 59.54           C
ATOM   3510  CA  ARG E1107      12.093  -6.388 -87.153  1.00 61.78           C
ATOM   3511  CA  ASN E1108      11.365  -9.348 -84.862  1.00 58.86           C
ATOM   3512  CA  PHE E1109      11.816 -12.285 -87.274  1.00 59.35           C
ATOM   3513  CA  TYR E1110      14.334 -12.700 -90.079  1.00 62.85           C
ATOM   3514  CA  GLU E1111      12.320 -13.096 -93.278  1.00 65.13           C
ATOM   3515  CA  PRO E1112      14.104 -11.394 -96.174  1.00 66.73           C
ATOM   3516  CA  GLN E1113      12.010  -9.999 -99.006  1.00 69.64           C
ATOM   3517  CA  ILE E1114      12.970  -8.214-102.210  1.00 74.11           C
ATOM   3518  CA  ILE E1115      12.903  -4.458-101.657  1.00 74.42           C
ATOM   3519  CA  THR E1116      10.071  -2.919-103.678  1.00 77.87           C
ATOM   3520  CA  THR E1117       8.017   0.264-103.823  1.00 78.80           C
ATOM   3521  CA  ASP E1118       5.154  -1.525-102.009  1.00 80.72           C
ATOM   3522  CA  ASN E1119       7.034  -1.969 -98.710  1.00 73.36           C
ATOM   3523  CA  THR E1120       9.016   1.308 -98.637  1.00 71.19           C
ATOM   3524  CA  PHE E1121       8.204   5.015 -98.740  1.00 67.39           C
ATOM   3525  CA  VAL E1122      10.066   8.007-100.169  1.00 70.38           C
ATOM   3526  CA  SER E1123      10.894  11.212 -98.288  1.00 71.02           C
ATOM   3527  CA  GLY E1124      13.617  13.633 -99.345  1.00 75.18           C
ATOM   3528  CA  ASN E1125      16.845  12.913-101.174  1.00 82.63           C
ATOM   3529  CA  CYS E1126      20.083  11.020-100.543  1.00 84.17           C
ATOM   3530  CA  ASP E1127      22.060  14.049 -99.307  1.00 85.67           C
ATOM   3531  CA  VAL E1128      20.501  14.257 -95.828  1.00 76.40           C
ATOM   3532  CA  VAL E1129      20.423  10.701 -94.403  1.00 72.59           C
ATOM   3533  CA  ILE E1130      23.656   9.672 -92.682  1.00 69.71           C
ATOM   3534  CA  GLY E1131      24.590   6.118 -93.623  1.00 69.74           C
ATOM   3535  CA  ILE E1132      22.382   5.774 -96.721  1.00 70.84           C
ATOM   3536  CA  VAL E1133      23.573   3.053 -99.104  1.00 74.66           C
ATOM   3537  CA  ASN E1134      23.238   2.607-102.852  1.00 86.35           C
ATOM   3538  CA  ASN E1135      20.986  -0.200-104.124  1.00 82.28           C
ATOM   3539  CA  THR E1136      18.158  -1.031-106.516  1.00 84.43           C
ATOM   3540  CA  VAL E1137      14.458  -0.709-105.669  1.00 83.27           C
ATOM   3541  CA  TYR E1138      12.183  -2.771-107.902  1.00 88.83           C
ATOM   3542  CA  ASP E1139       8.810  -1.600-109.226  1.00 97.37           C
ATOM   3543  CA  PRO E1140       6.714  -4.697-110.313  1.00 98.95           C
TER    3544      PRO E1140
END
```
